# Cell size reduction scales spindle elongation but not chromosome segregation in *C. elegans*

**DOI:** 10.1101/2025.10.13.681585

**Authors:** Chukwuebuka William Okafornta, Reza Farhadifar, Gunar Fabig, Hai-Yin Wu, Maria Köckert, Martin Vogel, Daniel Baum, Robert Haase, Michael J. Shelley, Daniel J. Needleman, Thomas Müller-Reichert

## Abstract

How embryos adapt their internal cellular machinery to reductions in cell size during development remains a fundamental question in cell biology^1–11^. Here, we use high-resolution lattice light-sheet fluorescence microscopy and automated image analysis to quantify lineage-resolved mitotic spindle and chromosome segregation dynamics from the 2-to 64-cell stages in *Caenorhabditis elegans* embryos. While spindle length scales with cell size across both wild-type and size-perturbed embryos, chromosome segregation dynamics remain largely invariant, suggesting that distinct mechanisms govern these mitotic processes. Combining femtosecond laser ablation^12,13^ with large-scale electron tomography^14^, we find that central spindle microtubules mediate chromosome segregation dynamics and remain uncoupled from cell size across all stages of early development. In contrast, spindle elongation is driven by cortically anchored motor proteins and astral microtubules, rendering it sensitive to cell size^12,13,15–17^. Incorporating these experimental results into an extended stoichiometric model for both the spindle and chromosomes, we find that allowing only cell size and microtubule catastrophe rates to vary reproduces elongation dynamics across development. The same model also accounts for centrosome separation and pronuclear positioning in the one-cell *C. elegans* embryo^18^, spindle-length scaling across nematode species spanning ~100 million years of divergence^17^, and spindle rotation in human cells^19^. Thus, a unified stoichiometric framework provides a predictive, mechanistic account of spindle and nuclear dynamics across scales and species.

## Main

Scaling, the systematic co-variation of biological form and function with size, is a widespread phenomenon, ranging from organismal metabolic rates to subcellular organelle dimensions^20,21,9,3^. Although much is known about how organelles assemble and operate during cell division across model systems, early embryos pose a special challenge: as cells rapidly change in size, geometry, and composition, the mitotic spindle must still ensure faithful chromosome segregation. Spindle behavior must adapt along multiple axes, including elongation dynamics and architecture, orientation within tissues, and interdivision timing. Yet the principles that couple all these adaptations to cell size versus developmental timing remain poorly resolved.

The nematode, *Caenorhabditis elegans*, serves as a premier model system for investigating scaling of cellular components during early embryogenesis^22^. The cell lineage of *C. elegans* is completely annotated^23^, giving access to accurate structural studies, biophysical quantifications, and component-wise comparisons of simultaneous biological processes during development. On top of that, the identity of each cell and its components during embryonic development is easy to follow due to its stereotypic division pattern. The *C. elegans* embryo undergoes a progressive reductive early development in that each of the resulting daughter cells from each division cycle is about half the volume compared to the parental cell. Previous studies have shown that centrosomes^24^, the nucleus and chromosome length^25^, the nucleolus^11^, and the spindle^26^ scale with cell size during development. The spindle, however, must consistently coordinate the movement of the same DNA mass, ensuring that each daughter cell receives an identical, single copy of the genome. This raises a fascinating question: how can one structure dramatically change its morphology yet reliably perform the same function?

Here, we approach this question by analyzing the scaling of both spindle and chromosome dynamics throughout *C. elegans* early embryonic development. Using a combination of live-cell imaging, laser ablation, cellular electron tomography, and mathematical modeling, we reveal that spindle length and dynamics scale with cell size, while chromosome movement and separation remain largely independent of cell size reduction. Our work demonstrates that the mechanisms driving spindle elongation are fundamentally distinct from those governing chromosome segregation, resulting in their differential scaling in the early nematode development.

### Building a spindle atlas for *C. elegans*

Beyond the one-cell embryo, the *C. elegans* cell-division machinery has not been characterized at lineage resolution. Here, we systematically quantified the size and dynamics of the mitotic spindle and separation of the chromosomes across all embryonic lineages from the two-to the 64-cell stages (Fig. S4). We used a custom-built lattice light-sheet fluorescence microscope (LLSM)^27^ and acquired six-dimensional datasets of *C. elegans* embryos (x, y, z, time, and two color channels) expressing protein-tagged GFP::γ-tubulin, mCherry::histone (H2B), and mKate2::PH domain (membrane), resulting in a total of ~3.8 million images (Fig. 1a, Movie S1). We developed a pipeline to automatically segment the cell periphery using U-net^28,29^, a well-established neural network for segmentation of microscopy images, combined with automated 3D tracking of centrosomes in each cell of the developing embryo (Fig. 1b-d, Movie S2). In addition, we tracked the position of the centrosomes and generated kymographs, each aligned with the 3D spindle axis, which has a different orientation in each embryonic cell (Fig. 1e). We then measured pole-to-pole distance (D_P-P_) and chromosome-to-chromosome distance (D_C-C_) over time and determined the anaphase onset (AO), when the two chromatids start to separate (Fig. 1e). Since the embryonic developmental pattern in *C. elegans* is highly stereotypic, we used the position of the cells to annotate cell lineages. We built the spindle lineage atlas for *C. elegans* (Supplementary Material; Fig. S4), which includes spindle elongation dynamics (Fig. 1f), chromosome segregation dynamics (Fig. 1g), and cell size. We then measured six characteristic parameters per cell in each of the 12 wild-type embryos: spindle initial length, spindle final length, spindle elongation rate, chromosome segregation rate, chromosome final separation distance, and cell size, calculated as the cube root of cell volume (Supplementary Material; Fig. 1f, g). Using analysis of variance (ANOVA), we quantified differences in spindle final pole-to-pole distance with the F-statistic, which compares between-group to within-group variability (F = MS_between_ /MS_within_); a larger F indicates greater separation of group means relative to residual noise. Among lineages within the 32-cell stage, we observed clear heterogeneity, F = 4.9. Comparing developmental stages (32 *vs* 64 cells) yielded a much larger effect, F = 517.7. Thus, differences attributable to developmental stage dominate those among lineages within a stage, while the within-stage result confirms that our measurements resolve lineage-specific variation. We, therefore, focused on spindle variation across developmental stages.

**Fig. 1.**
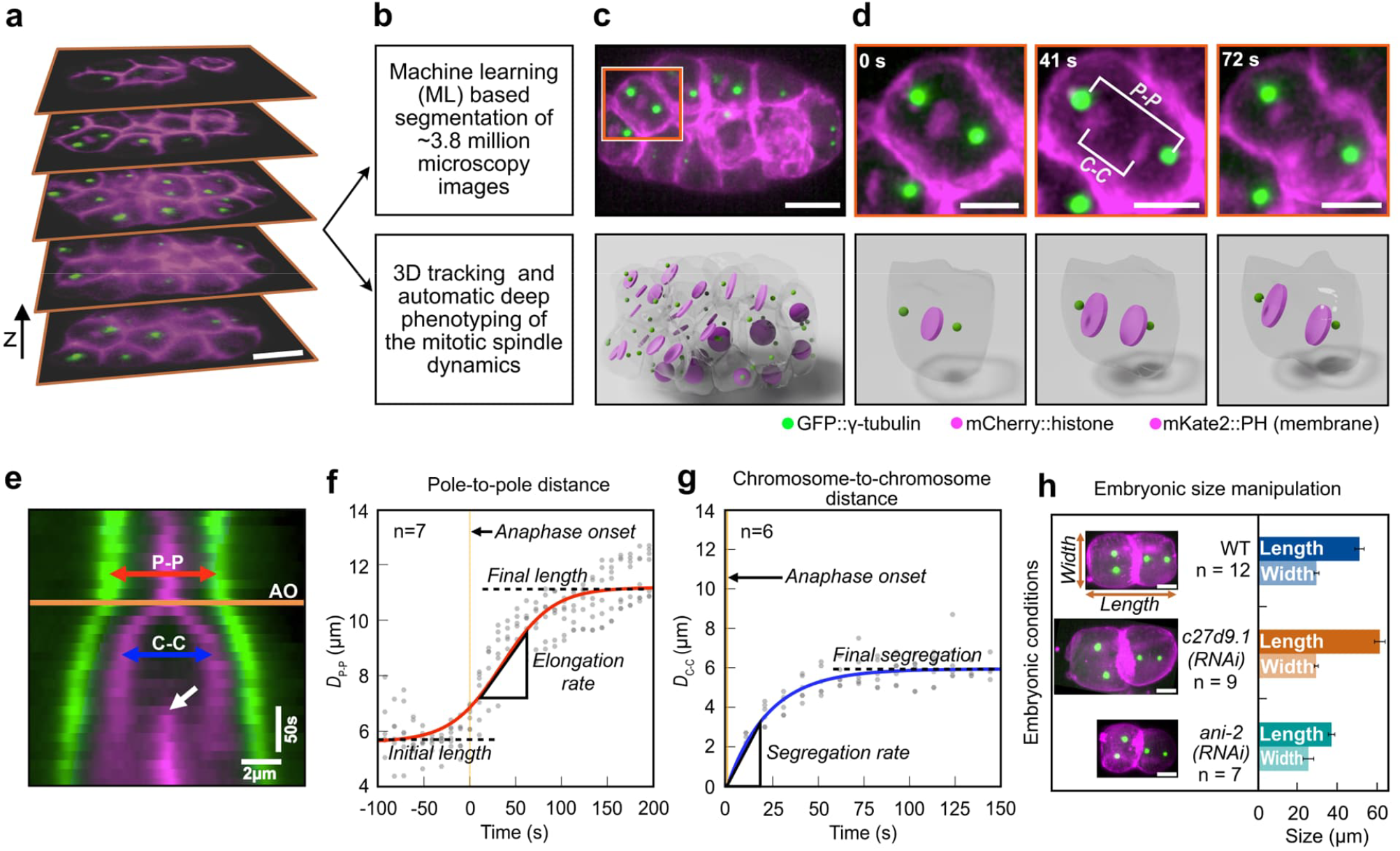
Quantitative measurement of spindle dynamics during *C. elegans* development. **a**, Six-dimensional (x, y, z, time, and two-color channels) lattice light-sheet microscopy of *C. elegans* embryos from 2-to 64-cell stage. Selected z-stacks from a 32-cell stage embryo are shown. Both the chromosomes (mCherry::histone) and the cell membrane (mKate2::PH domain) are shown in magenta, the centrosomes (GPF::γ-tubulin) in green. **b**, Pipeline of image processing and phenotyping using a combination of machine learning approaches for membrane segmentation and automated tracking of centrosomes and chromosomes. **c**, Top panel, z-projection of *C. elegans* embryo at 32-cell stage. Square represents the crop-out cell in **d**. Lower panel, 3D rendering of the embryo shown on top. Cell peripheries are shown in gray, chromosomes/nuclei in magenta, and centrosomes in green. **d**, Upper panel, snapshots from cell ABalpa highlighted in **c** from the onset of chromosome separation to furrow formation. The pole-to-pole distance (P-P) and chromosome-to-chromosome distance (C-C) are indicated. Lower panel, 3D rendering of the cell shown on top. **e**, Kymograph showing centrosomes and chromosomes for the ABalpa cell from **d**. The orange line shows the onset of anaphase (AO), defined as the last time point before separation of the chromosomes. The white arrow shows the ingressing cell membrane. Scale bars for space (horizontal) and time (vertical) are shown. **f**, Pole-to-pole distance (D_P−P_) as a function of time for ABalpa cell from 7 embryos (gray dots) is shown. The origin of time is set to AO (orange line). A sigmoid fit to the data is shown in red. Initial spindle length (5.56 µm), final spindle length (11.17 µm), and elongation rate of the spindle (2.97 µm/min) as measured from the sigmoid fit are shown. **g**, Chromosome-to-chromosome distance (D_c−c_) as a function of time for ABalpa cell from six embryos (gray dots) is shown. The origin of time is set to AO (orange line). An exponential fit to the data is shown in blue. The rate of segregation (15.52 µm/min) and the final segregation distance (5.93 µm) as measured from the exponential fit are shown. **h**, Two-cell stage embryos from wildtype (WT) and two RNAi conditions, c27d9.1 with large embryo and ani-2 with small embryo, are shown. The length and width of the embryo are indicated. The bar plot shows the average length and width of embryos for these three conditions. Error bar indicates the standard deviation. Scale bar: 10 µm.

### Scaling rules for spindle and chromosome dynamics are different

To quantify the average spindle dynamics across 745 cells from the 12 wild-type embryos, we then calculated the average spindle length over all cells in 2-, 4-, 8-, 16-, 32-, and 64-cell stage embryos (Fig. 2a-b; Fig. S2a(i)-g(i)). Consistent with previous studies^2,26,30^, we observed that the spindle dynamics dramatically change as the embryo proceeds in development and cells get progressively smaller (Fig. 2c). We quantified spindle length and spindle elongation rate by fitting a sigmoid function and estimated cell size by calculating the cube root of cell volume for all wild-type datasets up to the 64-cell stage (see Methods). We observed that spindle length decreased by ~2-fold from the 2-to the 64-cell stage (~55 % decrease in length), and found that the spindle final length scales with estimated cell size with slope = 0.75 ± 0.03 (mean ± standard deviation; Fig. 2d).

**Fig. 2.**
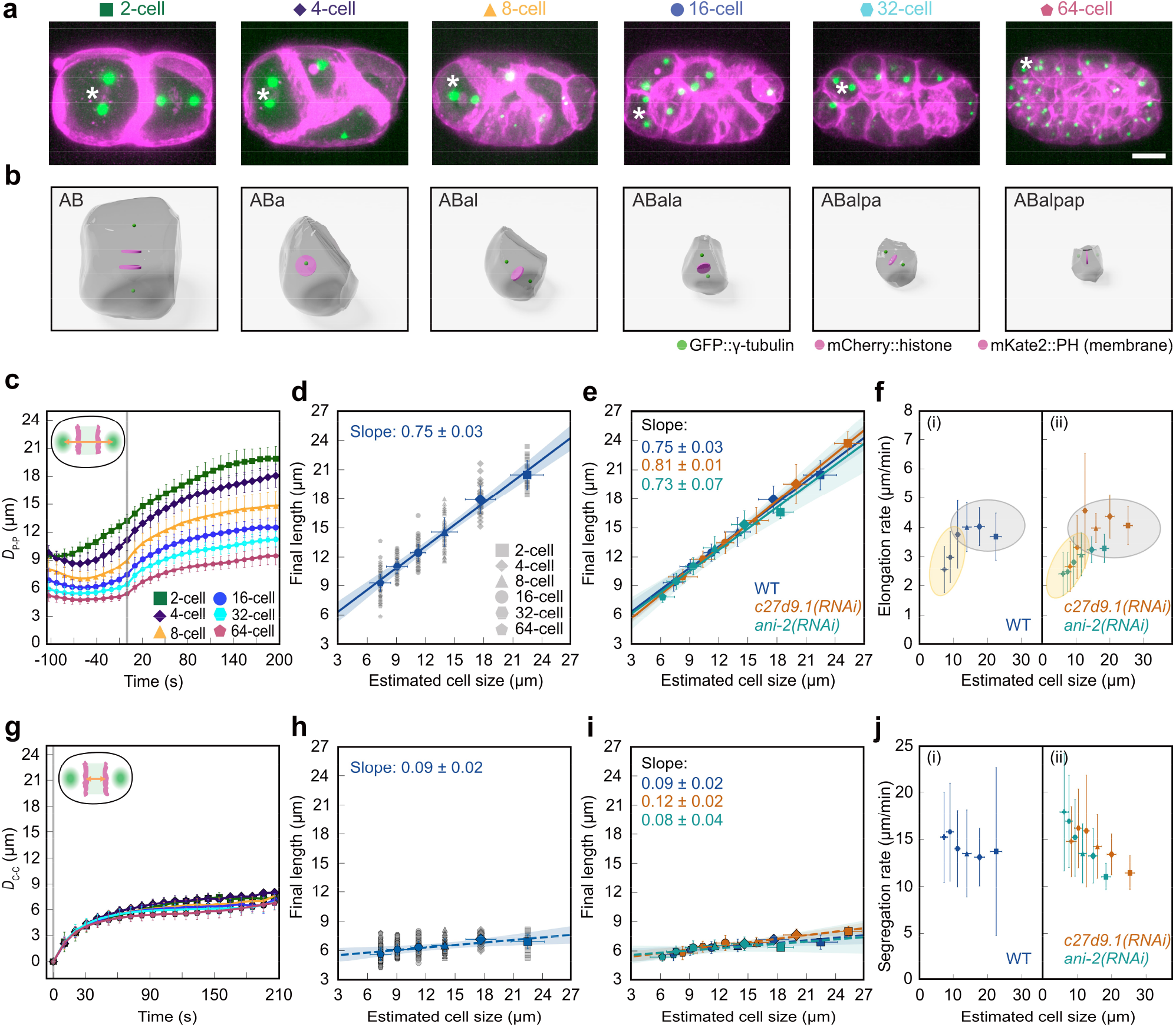
Spindle and chromosome dynamics scale differently with cell size. **a**, Partial z-projection of an embryo at different cell stages during development. The asterisk highlights the cells rendered in **b**. Chromosomes (mCherry::histone, magenta), cell membrane (mKate2::PH domain, magenta), and centrosomes (GPF::γ-tubulin, green) are shown. Scale bar, 10 µm. **b**, 3D rendering of the cells highlighted with asterisks in **a** with cell peripheries (gray), chromosomes (magenta), and centrosomes (green) are shown. **c**, Average pole-to-pole distance (D_P−P_) as a function of time for different cell stages of WT embryos (error bars, standard deviation). The gray line shows the onset of anaphase. Inset is the schematic of the embryo indicating D_P−P_. **d**, Final pole-to-pole distance as a function of cell size for WT embryos. Gray symbols show the measurements from individual embryos, and blue symbols show the average (vertical and horizontal error bars: standard deviation and error propagation, respectively). The blue line shows the linear regression to the average, with the blue shadow as the 95% confidence interval. **e**, Final pole-to-pole distance as a function of cell size for WT, *c27d9*.*1(RNAi)*, and *ani-2(RNAi)* (vertical and horizontal error bars: standard deviation and error propagation, respectively). The colored lines show the linear regression to the average, with the shadows as the 95% confidence interval. **f**, Spindle elongation rate as a function of cell size for WT, *c27d9*.*1(RNAi)*, and *ani-2(RNAi)* embryos (vertical and horizontal error bars: standard deviation and error propagation, respectively). **g**, Average chromosome-to-chromosome distance (D_c−c_) as a function of time for different cell stages of WT embryos (error bar, standard deviation). The gray line shows the onset of anaphase. Inset is the schematic of the embryo indicating D_c−c_. **h**, Final chromosome-to-chromosome distance as a function of cell size. Gray symbols show the measurements from individual embryos, and blue symbols show the average (vertical and horizontal error bars: standard deviation and error propagation, respectively). The blue line shows the linear regression to the average, with the blue shadow as the 95% confidence interval. **i**, Final chromosome-to-chromosome distance as a function of cell size for wild-type (WT), *c27d9*.*1(RNAi)*, and *ani-2(RNAi)* embryos (vertical and horizontal error bars: standard deviation and error propagation, respectively). The colored lines show the linear regression to the average, with the shadows as the 95% confidence interval. **j**, Chromosome segregation rate as a function of cell size for WT, *c27d9*.*1(RNAi)*, and *ani-2(RNAi)* embryos (vertical and horizontal error bars: standard deviation and error propagation, respectively).

We sought to test whether spindle scaling is governed by developmental progression or directly by cell size. Under a developmental clock model, final spindle length is prescribed solely by stage; thus, increasing or decreasing embryo size without changing developmental stage would leave the spindle length at that stage unchanged. By contrast, a cell-size–dependent model posits that spindle length is set by cellular dimensions; thus, at a given stage, larger cells should assemble longer spindles and smaller cells shorter spindles. Accordingly, when data are plotted, the developmental-clock model predicts collapse by developmental stage but not by cell size, whereas the cell size-dependent model predicts collapse by cell size but not by stage (Fig. S5a-d).

We used *c27d9*.*1(RNAi)* to generate larger embryos compared to wild type (n = 9) with a length of 62 µm, and *ani-2(RNAi)* to obtain smaller embryos (n = 7) with a length of 37 µm (wild-type embryo length was 51 µm (Fig. 1h, Movie S3 & S4). By analyzing a total of 1099 cells, we built the spindle lineage atlas for both RNAi conditions from the 2-cell stage up to the 64-cell stage (see Supplementary Material; Fig. S1a-f). Similar to wild type, we measured the average spindle dynamics in the 2-to 64-cell stage and quantified the final spindle length by fitting a sigmoid function (Fig. S2a(ii)-g(ii). In both cases, we observed that the spindle length scales with cell size with a slope similar to the wild-type (slope: WT = 0.75 ± 0.03; *c27d9*.*1(RNAi)* = 0.81 ± 0.01; *ani-2(RNAi)* = 0.73 ± 0.07; mean ± SD; Fig. 2e). Therefore, the scaling of the final spindle length in *C. elegans* is most likely driven by cell size.

To further test the origin of spindle scaling (developmental clock vs cell size), we also measured how the elongation rate of the spindle scales (see Methods). We found that, in wild-type, from the 2- to 8-cell stage, the elongation rate remains almost independent of cell size (Fig. 2f(i), gray color), but after the 8-cell stage, when cell size is ~12 µm, the elongation rate decreases with cell size (Fig. 2f(i), yellow color). If this change was due to a developmental clock, we expect that in *c27d9*.*1(RNAi)* and *ani-2(RNAi)* embryos, the transition happens in the 8-cell stage. However, if this transition in elongation rate was due to changes in cell size, we expect it to happen when cells are ~12 µm in both *c27d9*.*1(RNAi)* and *ani-2(RNAi)* embryos, irrespective of the stage. We repeated these measurements in *c27d9*.*1(RNAi)* and *ani-2(RNAi)* backgrounds, fitted a piecewise linear function, and measured the transition in elongation rate (see Methods and Fig S6a-c). We observed that in both cases this transition roughly happens when cells are ~12 µm in size, which is the 16-cell stage in *c27d9*.*1(RNAi)* and the 4-cell stage in *ani-2(RNAi)* (Fig. 2f(ii)). Similar to the final spindle length, the scaling of spindle elongation rate in *C. elegans* is most likely driven by cell size.

We next asked how chromosome motion changes as cells, and consequently their spindles, dramatically decrease in size during *C. elegans* development. We measured the distance between the segregating chromatids, D_C-C_, as a function of time from the onset of anaphase to the early stage of cytokinesis for 2-to 64-cell stages (Fig. 2g, inset). Surprisingly, we observed that chromosome segregation dynamics remained mostly unchanged during embryonic development (Fig. 2g). By fitting an exponential function to the data, we measured the final distance between the chromatids as a function of cell size. The slope of scaling for WT chromosome distance was 0.09 ± 0.02 (mean ± SD; Fig. 2h). Thus, during *C. elegans* development, a ~3-fold reduction in cell size results in ~225% reduction in spindle final length, but only ~30% reduction in final chromosome separation. Repeating this analysis for *c27d9*.*1(RNAi)* and *ani-2(RNAi)* embryos, we found that in both cases, the distance between chromosomes scales with cell size at a similar rate to the wild-type (slope: *c27d9*.*1(RNAi)* = 0.12 ± 0.02; *ani-2(RNAi)* = 0.08 ± 0.04; mean ± SD; Fig. 2i; Supplementary Material, Fig. S1d-f; Table S4). Interestingly, the dynamics of chromosome segregation (both the speed of separation and the final distance) remained mostly unchanged with cell size reduction during development.

### Examining forces on the spindle and chromosomes throughout development

We next asked what force mechanism might regulate spindle pole dynamics and chromosome separation during development. Previous studies have used laser ablation to investigate forces acting on the spindle in the one-cell stage *C. elegans* embryo^12–15,17,31^. The first mitotic spindle is large and parallel to the imaging plane, so it is amenable to ablation; later-stage spindles are smaller and variously oriented, making microtubule ablation challenging. To selectively sever sub-populations of microtubules in these smaller spindles (Fig. 3a; see Methods), we developed and utilized a femtosecond stereotactic laser ablation system (FESLA), which enables precise cutting of structures at near-diffraction limits^13^. We ablated microtubules between the spindle pole and chromosome in a 16-cell-stage embryo, where the distance between the pole and chromosome is only about one micron (Fig. 3b–c), thus demonstrating the precision of FESLA. We then monitored the motion of spindle poles and chromosomes in response to the ablation (Fig. 3d). Measuring the pole-to-pole (D_P-P_; Fig. 3e, khaki), pole-to-chromosome (D_P-C_; Fig. 3e, slate blue), and chromosome-to-chromosome distance (D_C-C_; Fig. 3e, coral), we found that after ablation, the pole moved away from the rest of the spindle toward the cell periphery. Subsequently, likely due to microtubule regrowth, the pole returned approximately to its original position. The motion of the chromatids, however, remained largely unchanged during this time (Fig. 3e, coral).

**Fig. 3.**
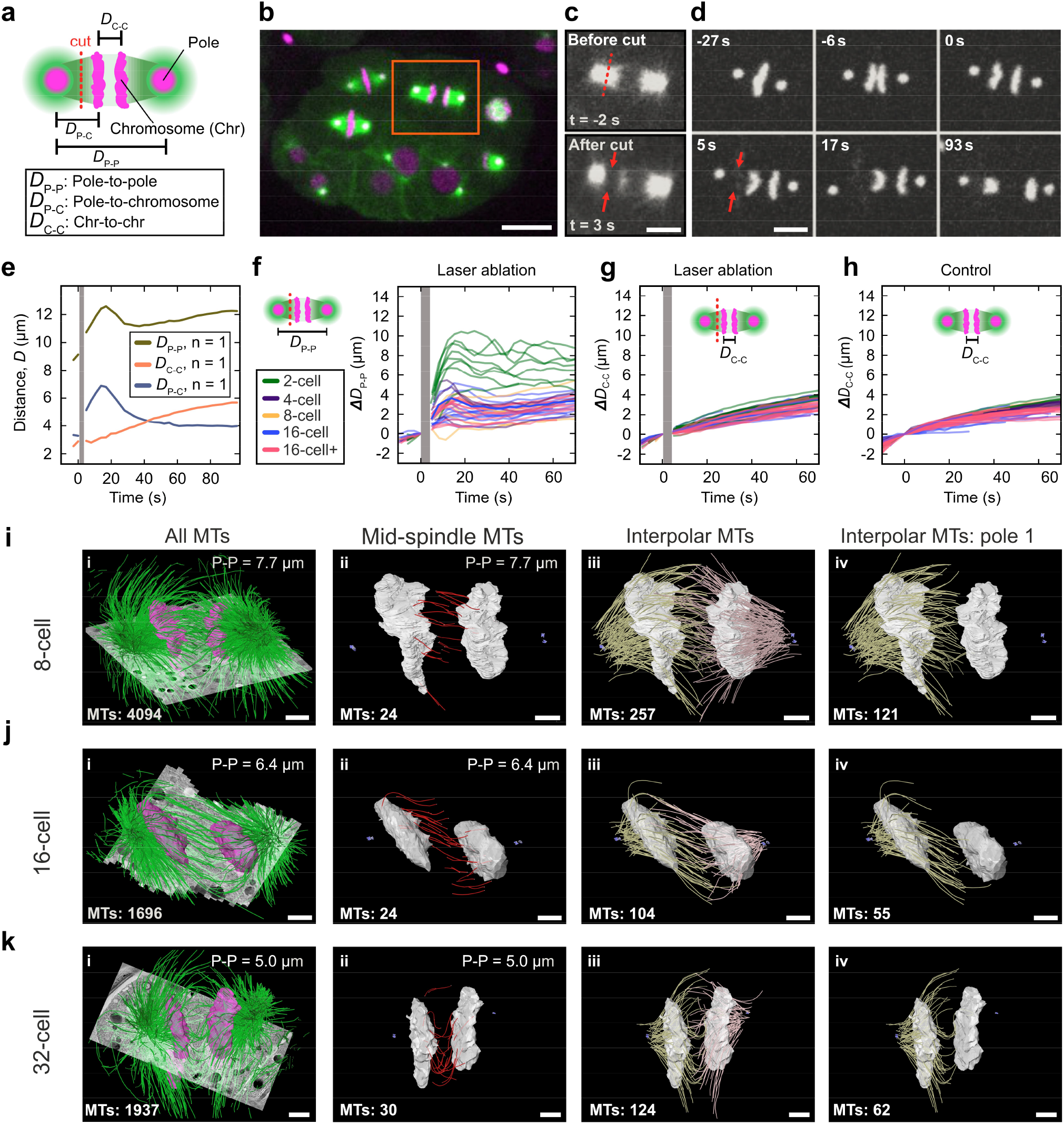
Different forces regulate spindle dynamics and chromosome segregation. **a**, Schematic of laser ablation of the spindle and corresponding measured quantities (magenta, spindle poles and chromosomes; green, microtubules, MTs; red dashed line: ablation position). **b**, Snapshot of a 16-cell stage embryo before ablation (magenta, chromosome and centrosome; green, microtubule; orange box, ablated spindle). Scale bar, 10 µm. **c**, Top panel, spindle microtubules before ablation (red dashed line, ablation position). Lower panel, spindle microtubules after ablation (red arrows, ablation position). Scale bar, 5 µm. **d**, Snapshots of centrosomes and chromosomes before and after ablation from **c** (red arrows, ablation position). Scale bar, 5 µm. **e**, Measured displacements (D_P−P_, D_P−c_, and D_c−c_) as a function of time for the ablation in **c**. The Gray bar shows the time of ablation. **f**, Relative pole-to-pole distance (ΔD_P−P_) as a function of time for ablation experiments of different cell stages (gray bar, ablation time). The schematic shows the position of ablation. **g**, Relative chromosome-to-chromosome distance (ΔD_c−c_) as a function of time for ablation experiments of different cell stages (gray bar, ablation time). The schematic shows the position of ablation. **h**, Relative chromosome-to-chromosome distance for unperturbed cells. **i**-**k**, Three-dimensional models of microtubules (green) and chromosomes (magenta) overlayed on tomographic slices (gray) for 8-, 16-, and 32-cell stage cells are shown **(i)**. Mid-spindle microtubules (red) with both ends in between chromosomes (gray) are shown in ii. Interpolar microtubules (yellow and pink) spanning from each pole together with chromosomes (gray) are shown in iii. Interpolar microtubules from one pole (yellow), together with chromosomes (gray), are shown in iv. Centrioles are shown in purple, pole-to-pole distance (top, right) and the number of microtubules (bottom, left) for each panel are indicated. Scale bar, 1 µm.

To extend these findings, we systematically performed FESLA experiments from the 2-to 16+ cell stages, measuring pole-to-chromosome and chromosome-to-chromosome distances over time. Across all stages, spindle poles consistently moved away from the spindle body toward the cell periphery following ablation (Fig. 3f, Movie S5). In addition, across all cell stages, we observed more than a 30% increase in the elongation rate of the poles in the first 15 s after ablation and a gradual recovery as aforementioned (Table S6). In contrast, the chromosome segregation remained unaffected (Fig. 3g–h, and Fig. S3a-d). These results demonstrate that spindle pole dynamics at later stages are driven by pulling forces generated from cortically anchored motor proteins. Furthermore, they reveal that distinct force mechanisms regulate spindle pole and chromosome motion, as ablation perturbs only the former.

Previously, we combined such ablation experiments with electron tomography and showed that forces that segregate the chromosomes during the first mitosis in *C. elegans* are generated by a sub-population of microtubules nucleating and growing in between the chromosomes^12^. We next investigated whether a similar microtubule population exists at later developmental stages. For this, we utilized serial-section electron tomography after high-pressure freezing to investigate the inter-chromosomal 3D organization of microtubules in three spindles of 8-, 16-, and 32-cell stages at anaphase. We identified the centrioles as markers for spindle poles, traced all microtubules, and segmented the chromosomes (Fig. 3i-k(i)). We found that there is a ~10-fold reduction in the total number of microtubules from the one-cell stage^14^ to the 32-cell stage (~20,000 microtubules for the one-cell stage vs. ~2000 microtubules at the 32-cell stage). We then classified the microtubules based on their associations with spindle poles: those with one end near a centriole are classified as interpolar microtubules, and those with both ends in between the chromosomes are classified as mid-spindle microtubules. We observed that in all three tomograms, there is a population of mid-spindle microtubules, similar to what we found in our previous study in the one-cell stage embryo (Fig. 3i-k(ii)). The number of mid-spindle microtubules in these later stages was fewer than those in the first spindle (~200 in the one-cell stage^12^ vs. ~30 in the 32-cell stage). In addition, we observed ~100-200 interpolar microtubules, with some passing through the nearest chromosomal plate and reaching out to the other plate (Fig. 3i-k(iii-iv)). Our ablation experiments and electron tomography showed that such interpolar microtubules have little or no effect on the segregation dynamics of the chromosomes. Most likely, the mid-spindle microtubules segregate and regulate the separation of the chromosomes in later developmental stages in *C. elegans*, similar to the one-cell embryo.

Taken together, our results suggest that the dynamics of the pole separation are different from the dynamics of the chromosome separation during embryonic development. Spindle length elongation is controlled by the interaction of astral microtubules with cortically bound molecular motors and is thus dependent on cell size^17^. In contrast, chromosome segregation in *C. elegans* is only regulated by the mid-spindle microtubules^12^ and is independent of cell size. We, therefore, sought to theoretically investigate if such mechanisms could indeed result in the observed scaling of the spindle.

### Microtubule length scales with cell size to regulate spindle dynamics behavior

Previously, we have developed and used a stoichiometric model as a quantitative framework to study the dynamics of spindle elongation in one-cell stage nematode embryos and human cells^17,32^. In this model, the interaction of a cortically anchored dynein motor with microtubules is stoichiometric – each motor can only bind one microtubule at a time. This constraint causes a competition between centrosomal asters for binding to the motor, resulting in a stable positioning of the asters in the two hemispheres of the embryo, achieved even if only pulling forces are present. Here, we extend this stoichiometric model to incorporate the internal mechanics of the spindle and chromosome dynamics by adding viscous elements between spindle poles and their associated chromosomes (Fig. 4a; see Methods). Therefore, the dynamics of spindle poles are governed by a balance of pulling forces from cortical motors, cytoplasmic drag forces on the poles (with coefficient η), and viscous friction arising from the relative speed of the spindle pole and the chromosome (with coefficient ν)

**Fig. 4.**
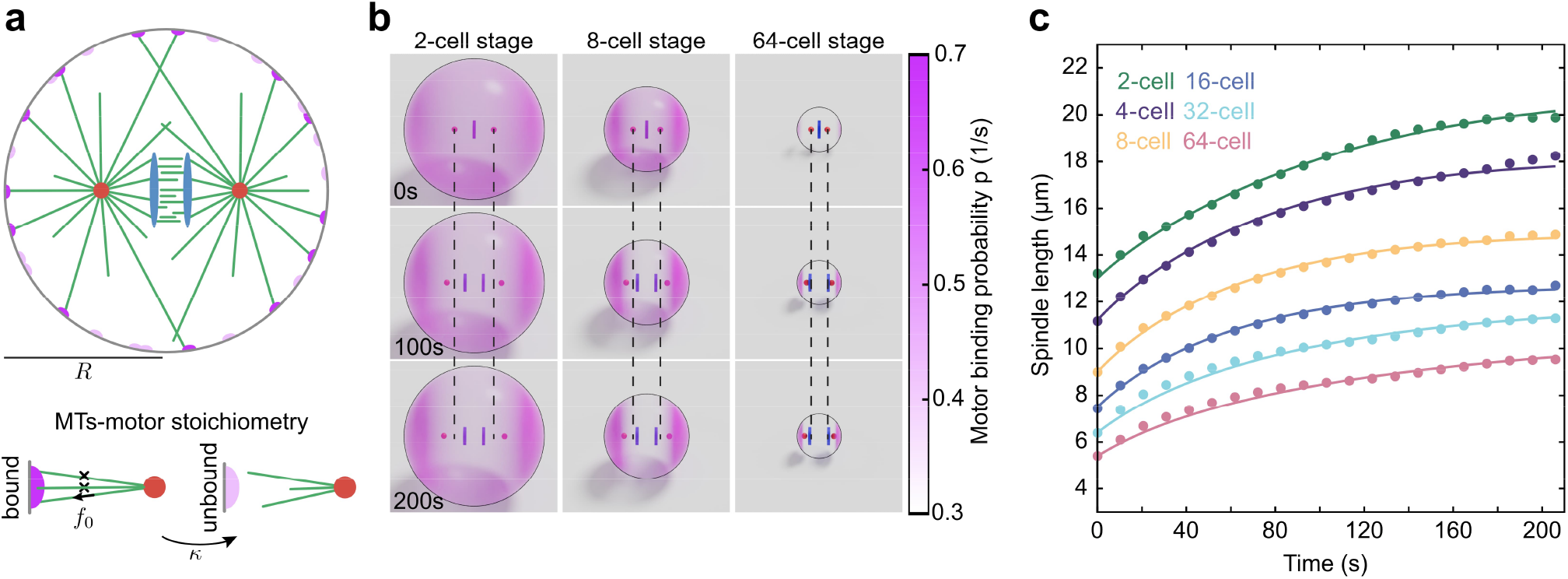
Theory quantitatively explains spindle and chromosome dynamics. **a**, Schematic of the Stoichiometric model for a spindle in a sphere of radius, R, where two centrosomal asters (red) nucleate microtubules (green) in all directions. Some microtubules reach cortical motors (dark purple) and get pulled upon. The chromosomal plates are shown in blue. The lower panel indicates the stoichiometric interaction of microtubules and a motor. Once a motor is bound, it exerts a pulling force of f_0_ in the direction of the microtubule. The microtubule detaches from the motor at a rate κ. **b**, Snapshots from simulations of 2-, 8-, and 64-cell stage cells. The binding probability of the motors is shown in pink. Dashed lines show the initial position of centrosomes. **c**, Spindle length as a function of time for different cell stages (lines show simulation, and dots indicate experimental measurements of wild-type embryos). The time t=0 corresponds to the anaphase onset.

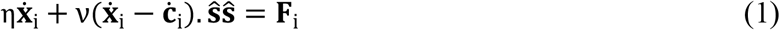

Here, 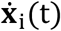 and 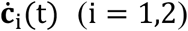 are the velocities of spindle poles and chromosomes, respectively, and **F**_i_ is the total pulling force from cortical motors on the *i*th pole. The vector **ŝ = (x**_**1**_ **− x**_**2**_**)/**|**x**_**1**_ **− x**_**2**_| is the spindle axis orientation vector. The force **F**_i_ is evaluated from an evolving coarse-grained surface field for the probability of a motor being bound to a microtubule from the *i*th centrosome (see Methods).

Our laser ablation experiments indicated that the forces that segregate the chromosomes are generated from the motor/microtubule complex between the chromosomes. Our live-cell microscopy experiments show that the dynamics of the separation vector of the two chromosomal sets, Δ**c**(t), is relatively independent of cell size and is well approximated by an exponential function, Δ**c**(t) = c_f_(1 − e^−t/τ^)**ŝ**, with c_f_ ~ 6.2 μm and τ ~ 29 s. For simplicity, we do not explicitly model the biophysics of chromosome separation, but instead impose the dynamics in our simulations (see Methods). For each cell stage, we performed simulations in spherical cells of radii given by the cube root of experimentally measured average cell volume at that stage, and used the experimentally measured average spindle initial length. Within an optimization problem, we used the microtubule catastrophe rate as the sole fitting parameter of the model to the data at each stage. All other model parameters were kept fixed and estimated from previous measurements^17^ (see Methods).

Representative simulation snapshots, showing the positions of centrosomal asters, chromosomal plates, and the spatial distribution of motor binding probability on the cell surface for the 2-, 4-, and 64-cell stages, are shown in Figure 4b. As a result, the dynamics of spindle elongation for both simulation and experiment show remarkable concordance across time at each cell stage (Fig. 4c). We find that the fit catastrophe rate increases as cells divide, leading to increasingly smaller spindles within the increasingly smaller cells. The consequence of this is that while the cortical motors at the two cell poles are increasingly engaged, those encircling the spindle become less so. By contrast, holding the catastrophe rate constant across stages fails to produce proper spindle elongation (Fig. S7a). In conclusion, these results show that regulation of microtubule dynamics over cell division can serve as a central mechanism for controlling both spindle scaling and chromosome separation through the changes of cell size in early embryonic development in *C. elegans*.

## Discussion

Our work sheds new light on spindle and chromosome dynamics during development. By combining long-term volumetric imaging with automated analysis, we established a mitotic spindle lineage atlas for early *C. elegans* development. We found that as cells progressively decrease in volume during embryogenesis, spindle length scales accordingly, yet the dynamics and extent of chromosome separation remain largely insensitive to cell size. Through an integrated approach combining femtosecond laser ablation, electron tomography, and mathematical modeling, we reveal that distinct force-generating mechanisms independently regulate spindle elongation and chromosome segregation. While many factors can change in cells during development, here we show that changes in cell size and microtubule catastrophe rate, and thus the microtubule average length, are enough to explain spindle scaling dynamics.

We present a spindle lineage atlas that systematically quantifies spindle and chromosome dynamics together with cell morphology from the 2-to 64-cell stage in wild-type and two mutant *C. elegans* backgrounds at single-cell resolution. This dataset enables quantitative analysis of lineage-specific regulation of spindle dynamics in early development and creates a foundation for future mechanistic studies. Complementing these data, our electron tomography analyses, covering later developmental stages and referencing prior work on the one-cell embryo^12,14^, provide quantitative measures of spindle microtubule number and spatial organization, offering detailed structural insights into how spindle architecture evolves and revealing new possibilities for investigating spatial regulation and geometrical constriction of microtubules in development.

Development and evolution are deeply intertwined, and insights often emerge from studying them in parallel. We have previously examined the evolutionary aspects of the first mitotic spindle by quantifying its dynamics both within *C. elegans* and across nematode species separated by about 100 million years of evolution^10^. Across this wide evolutionary span, embryo size varies dramatically between species, and spindle size and dynamics scale accordingly^33^. Others have examined spindle scaling during development across a broad range of species, including yeast^34^, flies^35^, frog^36^, sea urchin^2^, fish^1^, and mouse^37^, and consistently observed similar relationships. These converging observations suggest that the underlying scaling principles are both universal and conserved. The stoichiometric model we describe here quantitatively captures both spindle positioning across *C. elegans* cell populations and among diverse nematode species, as well as spindle scaling as described in this study. Moreover, our experiments and modeling indicate that the same framework can also account for nuclear positioning prior to spindle formation^18^, highlighting its broader applicability to understanding cell division and development.

Looking forward, quantitative measurements of microtubule dynamics and organization, characterization of chromosome dynamics, and mapping of the spatial and temporal distribution in the context of key molecular players, such as PRC-1, regulating central spindle organization and motor proteins generating forces on the spindle, will enable the construction of a holistic biophysical model of spindle positioning and dynamics across both development and evolution.

## Methods

### Maintenance of *C. elegans*

Worms were cultured and maintained on standard nematode growth medium (NGM) plates at 20°C, seeded with OP50 feeding bacteria. Between 5 - 10 adult hermaphrodites^22^ were periodically transferred to fresh OP50-seeded NGM plates every 3 - 5 days to maintain the worm strain^38–40^.

### Worm strains

The *C. elegans* strains **TMR31**^41^ (unc-119(ed3) III; ddIs6[tbg-1::GFP + unc-119(+)]; ltIs37[pAA64; pie-1::mCherry::HIS-58 + unc-119(+)] IV; cpIs54 [mex-5p::mKate2::PLC(delta)-PH(A735T)::tbb-2 3’UTR + unc-119 (+)] II), a cross between **LP307**^42^ (unc-119(ed3) III; cpIs54 [mex-5p::mKate2::PLC(delta)-PH(A735T)::tbb-2 3’UTR + unc-119 (+)] II) and **TMR17**^43^ (unc-119(ed3) III; ddIs6[tbg-1::GFP + unc-119(+)]; ltIs37[pAA64; pie-1::mCherry::HIS-58 + unc-119(+)] IV), were used for live-cell imaging. To quantify spindle dynamics, centrosomes (GFP::γ-tubulin), chromosomes (mCherry::histone), and the plasma membrane (mKate2::PH domain) were visualized. In addition, *C. elegans* strain **SA250**^12,13^ (tjIs54 [pie-1p::GFP::tbb-2 + pie-1p::2xmCherry::tbg-1 + unc-119(+)]; tjIs57 [pie-1p::mCherry::his-48 + unc-119(+)]) was utilized for FESLA experiments in this study. Wild-type worms (N2, Bristol)^22^ were used for electron tomography.

### Manipulation of embryonic size

Embryonic size manipulation was done by RNAi feeding^44^. Approximately 20 L1 and non-gravid L4 wild-type worms were transferred to NGM plates seeded with an E. coli feeding strain expressing double-stranded RNA to knock down either ani-2^45,46^ or c27d9.1^11,47^ to generate smaller vs. bigger embryos, respectively. The RNAi worm plates were incubated at 20°C for 24 hours until the worms started laying eggs. By visual inspection, changes in embryo size with reference to wild-type embryos (length: ~50 µm, width: ~30 µm) were determined by using polystyrene beads (Ø 20 µm) in distilled water added to the dissection M9 buffer.

### Imaging of *C. elegans* embryos

On a depressed microscope glass slide, gravid worms in M9 buffer were dissected around the gonad region using a syringe needle to release early embryos into the buffer solution. The embryos were constantly kept moisturized to avoid dehydration. For imaging, early embryos were selected and transferred to poly-L-lysine-coated coverslips (Ø 5mm). Coverslips were mounted on custom-designed sample holders. For pressure-free imaging (i.e., without any compression) of both wild-type and RNAi-treated embryos at a controlled temperature of 20°C, a custom-designed lattice light-sheet fluorescence microscope^17^ (LLSM; MPI-CBG, Dresden) was used. Six-dimensional (6D) high-resolution images were acquired with a 25x detection objective (Nikon, 25x/NA 1.1 water immersion) and a 28.6x illumination objective (Special Optics, 28.6x/NA 0.7; custom objective). Images were detected using an ORCA Flash v4.0 camera (Dual Hamamatsu) with a time resolution of 10.3 s, almost without any damage caused by phototoxicity or photobleaching. mCherry and mKate2 fluorescence were excited with 5% of 25 mW laser power at 589 nm wavelength and detection at 642 nm wavelength using a 560 nm longpass beamsplitter as secondary dichroic (FF560-FDi01 Semrock BrightLine 560 edge). GFP fluorescence was excited with 3% of 25 mW laser power at 488 nm wavelength and detected using a longpass filter at 530 nm wavelength. The embryos were imaged continuously for about 3 hours (± 10 minutes). The raw data were saved as individual z-stacks for each frame.

### Data processing

The z-stacks of raw, unprocessed image datasets were stitched together using the “Import Image Sequence” function in Fiji/ImageJ^48,49^. Because of the *C. elegans* stereotypic pattern of early embryo development^23,50,51^, the arivis Vision4D software package was used for 6D cropping of the embryonic cells in each recorded embryo, starting from 10 frames before anaphase onset to the early stage of cytokinesis. To maintain the identity of each cell, accurate annotation was achieved by using the WormGUIDE software^52^.

### Automatic deep phenotyping and three-dimensional tracking of mitotic spindles

A supervised automatic tracking of the centrosomes over time was carried out for all cropped spindles by using an ImageJ macro custom script. This custom script took the 6D cropped cells as input and detected the bright GFP::γ-tubulin (i.e., the centrosomes) as designed within the script algorithm. The script algorithm is based on the CLIJ2 library that enables the use of a GPU for accelerated processing of images, especially biological images^53–55^. From the CLIJ2 library, a Gaussian blur was implemented to reduce the pixelated noise in x, y, and z. Then, automatic thresholding using the Phansalkar method was applied to improve the contrast of the LLSM images for spot detection^56^. The GFP::γ-tubulin bright spots were detected with a maxima box filter, converted to a point-list, and the two points closest to the center of the image stack were identified. The two GFP::γ-tubulin bright spots were then temporally tracked. In addition, the script was adapted to manually correct the wrongly detected GFP::γ-tubulin spots before saving the final 3D coordinate positions. Three-dimensional coordinate values in x, y and z for each timepoint of the dataset were extracted to a table as numbers.

### Generation and processing of kymographs

To quantitatively analyze spindle dynamics, a custom-written CLIJ2 library-based^53–55^ Fiji/ImageJ macro script was used to generate a 2D kymograph for two channels, i.e., the GPF channel for the centrosomes and the mCherry/mKate2 channel for the chromosomes and the cell membrane. The kymographs were adapted to determine a 3D Euclidean line through the center of mass of the two centrosomes along the spindle axis using the 3D coordinate information from the spot tracking custom algorithm and the 6D cropped cell dataset as input, taking into account the voxel size of the dataset in x, y, and z, i.e., 0.1 µm, 0.1 µm and 0.5 µm, respectively. Generation of the kymographs was achieved by using CLIJ2 affineTransform3D^53–55^ after precise isotropic transformation and translation of the input data to have a similar spindle orientation, thus solving the challenge of random orientation of the spindle axis over time. The maximum intensities of the fluorescent signals along the spindle axis were extracted and displayed as a kymograph in 1D. This process was repeated for all time series of each input dataset. Subsequently, the generated kymographs from each frame were stacked to form a continuous 2D kymograph for both channels to visualize spatiotemporal spindle dynamics over time.

The fluorescence intensity readouts for the spindle pole and chromosome positions from the kymographs were implemented using a supervised automated custom MATLAB script (The MathWorks Inc., Natick, USA). This customized MATLAB intensity readout script used the findpeaks function to locate the initial positions of the spindle poles. A Gaussian function was implemented to fit a curve to each spindle pole position using four variable parameters (a, b, c, d) to refine the peak intensity of each spindle pole as follows:

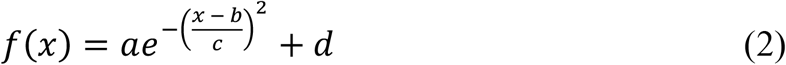

The accurate positions of the spindle poles were then extracted for all frames. Furthermore, the positions of the chromosomes were analyzed within the boundaries of the detected spindle poles. In addition, the script algorithm was designed to account for the chromosome movement during the metaphase-to-anaphase transition. The positions of the spindle poles and the chromosomes were then outputted to a table for further analysis.

### Machine learning-based segmentation of the cell membrane

For the segmentation of the mKate2-tagged cell membrane^42^, an iterative process between manual segmentation, training of a U-net based neural network, and automatic segmentation was used. First, we used Fiji^48,49^ to segment the cell membrane manually. For this, we used six different representative time points in three data sets. Then we used this manual segmentation to train a U-net^28^. This model was then used to segment all the membranes in all planes of the three example data sets and create a binary membrane segmentation. We then used ZIB Amira^57^ to create an image processing pipeline to create further training data for all time points of all available data sets. We used a combination of dilation/erosion to close gaps in the membrane after the automatic U-net segmentation. Then we filtered out small objects that were disconnected from the membrane signal (objects smaller than 7.5 × 10^8^ Å). After creating a binary image again, we used the Distance map algorithm^58,59^ (type: Euclid). After that, we inverted the image and applied a sigmoid intensity remapping (parameters: α=50.000, β=50.000, range: −25.000-15.000). After inverting the image again, we used the algorithm “contour tree segmentation”^71^ to segment the cells (parameters: threshold: −10.000, persistence mode: absolute, persistence options: range, persistence value: 1300, minimal segment size: 0, sort by neighbors: majority vote, fast segmentation: on). Then we filtered out all objects with more than 5000 voxels on the image border (BorderVoxelCount) and a cell volume greater than 1 × 10^13^ Å^3^. The resulting image volume contained all cells that could be segmented with confidence. We smoothed the outline of the cells by using a majority filter (kernel size X, Y, Z: 9, 9, 3). Next, we extracted the cell boundaries by using the function “Label boundaries” (XY planes) and converted them to binary images. These images were then utilized to retrain the U-net and segment the cell membrane again in all acquired images.

### Analysis of the mitotic spindle dynamics

Mitotic spindle dynamics were quantitatively analyzed with customized scripts and Jupyter Notebooks written in Python. The Python script algorithm was implemented to run a batch analysis process. A one-dimensional Euclidean distance for each frame was calculated using the saved read-out positions from the kymograph showing the spindle poles and the chromosomes for each cell in each recorded embryo. The 1D Euclidean distance was determined as follows:

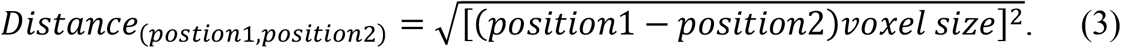

The calculated spindle lengths were given by the pole-to-pole distances, and the chromosome distances by the chromosome-to-chromosome distances. The final pole-to-pole length (FL_PP_) and the elongation rate (*ER*_*PP*_), when the inflection point (*t*_0_) is at its maximum rate (*τ*), were extracted for each cell in each embryo by fitting a sigmoid curve to the spatiotemporal distance as follows:

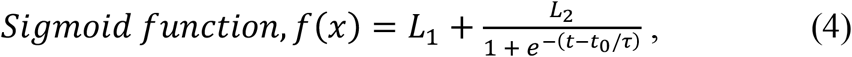

where *L*_1_ is the initial pole-to-pole length, *L*_2_ the total change in the pole-to-pole length, and t the time.

The final pole-to-pole length was calculated by determining the maximum length travelled by the poles (*FL*_*PP*_ = *L*_1_ + *L*_2_), and the pole-to-pole elongation rate was derived at the inflection point (*t* = *t*_0_), when the slope is at its maximum:

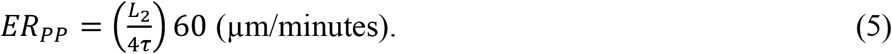

The final chromosome-to-chromosome length (*FL*_*cc*_) and the segregation rate (*SR*_*cc*_) of chromosomes for each cell of each embryo were extracted by fitting an exponential curve to the spatiotemporal distance using the following exponential function:

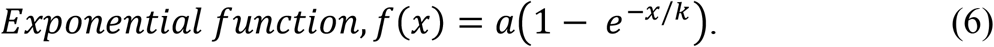

The final chromosome-to-chromosome length was derived from the exponential function as the amplitude (*a*), and the rate of chromosome segregation during anaphase was derived when time is equal to zero using the following expression:

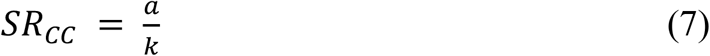

where, *k* is the time constant.

### Femtosecond stereotactic laser ablation (FESLA) experiments

Gravid *C. elegans* hermaphrodites were dissected, and early embryos were transferred by mouth pipette onto flat and trimmed 4% agarose (Bio-Rad) pads. After moisturizing with M9 buffer, embryos were mounted and held in place by the sunk agarose pad, which was sealed between a coverslip and a microscopic slide.

Confocal epifluorescent images were acquired through an inverted microscope (Nikon, TE2000) equipped with a 60X water-immersion objective (Nikon, CFI Plan Apo VC 60X WI, NA 1.2), a spinning disk unit (Yokogawa, CSU-X1), and an EM-CCD camera (Hamamatsu, ImagEM Enhanced C9100-13). The dynamics of spindle poles and chromosomes, through mCherry::γ-tubulin and mCherry::histone labelling, respectively, were derived from 3D time-lapse movies. GFP::β-tubulin images were also acquired to ensure microtubule severing. To perform FESLA^13^, a femtosecond laser pulse train (16-KHz repetition rate, ~70-fs pulse width, and 800-nm center wavelength) was introduced into the microscope and focused onto the sample by the same objective. The pulse train was obtained from a femtosecond Ti:sapphire laser (Spectra-Physics, Mai Tai DeepSee) after repetition reduction through a pulse picker (KMLabs, Eclipse). During FESLA execution, precise 3D sample movement and synchronized laser exposure were achieved through a 3-axis piezo stage (Physik Instrumente, P-545 PInano XYZ) and a mechanical fast shutter (Newport). Image acquisition and FESLA were controlled through a self-developed LabVIEW (National Instruments) interface.

The image analyses and data quantification for FESLA experiments were carried out in MATLAB (MathWorks). The features (poles and chromosomes) in the z-stack (with 1-μm spacing) time-lapse images were identified by the modified codes^13^ based on the 3D feature finding algorithms written by Yongxiang Gao and Maria Kilfoil^60^. With the derived 3D positions of features in different time frames, we tracked them using the codes from MATLAB Particle Tracking Code Repository by Daniel Blair and Eric Dufresne (http://site.physics.georgetown.edu/matlab/) (adapted from IDL Particle Tracking software from David D. Grier, John C. Crocker, and Eric R. Weeks)^61^. The extracted poles and chromosomes locations were then used to calculate the pole-to-pole distance D_P-P_, the pole-to-chromosome distance D_P-C_, and the chromosome-to-chromosome distance D_C-C_ over time.

To analyze the impact of FESLA, the pole-to-pole separation velocity 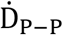 and the chromosome-to-chromosome separation velocity 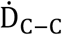 were calculated. The separation velocities before and after FESLA were derived from the slope of the linear fit using the time and distance data within the interval −8 < t ≪ 0 sec (for 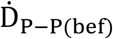 and 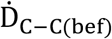) and 4 < t < 12 sec (for 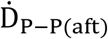 and 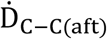), respectively. The separation velocities of the control set were calculated using the data within the intervals of the same conditions (−8 < t ≪ 0 sec for 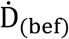 and 4 < t < 12 sec for 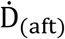). The differences in the velocities were expressed by 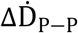 and 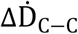.

### Electron tomography and ultrastructural analysis of *C. elegans* spindles

Wild-type hermaphrodites were cryo-immobilized using a Compact 03 high-pressure freezer (Wohlwend GmbH, Sennwald, Switzerland). Freeze substitution was performed over 3 days at −90°C in anhydrous acetone containing 1% OsO_4_ and 0.1% uranyl acetate using an automatic freeze substitution machine (EM AFS2, Leica Microsystems, Vienna, Austria). Epon/Araldite infiltrated samples were embedded in a thin layer of resin and polymerized for three days at 60°C^14^. Hermaphrodites with an appropriate number of developing embryos were selected by light microscopy for re-mounting on dummy blocks. Serial semi-thick sections (330 nm) were cut using an Ultracut UCT Microtome (Leica Microsystems, Vienna, Austria). Ribbons of serial sections through the adult hermaphrodite gonad were collected on Formvar-coated copper slot grids and poststained with 2% uranyl acetate in 70% methanol followed by Reynold’s lead citrate. Colloidal gold particles (20 nm; BBI, UK) were attached to both sides of the semi-thick sections to serve as fiducial markers for subsequent electron tomography. To select embryonic cells in anaphase in 8-, 16- and 32-cell embryos, serial sections were pre-inspected at low magnification (~2900x) using a transmission electron microscope (Morgagni, Thermo Fisher Scientific) operated at 80 kV. Ribbons containing embryos of interest were then transferred to a TECNAI F30 transmission electron microscope (Thermo Fischer Scientific, USA) operated at 300 kV and equipped with a CMOS camera (4k × 4k; Gatan, USA). Tilt series (a-stacks) were acquired over a ±60° range with 1° increments at a magnification of 4700x (pixel size 2.572 nm). Specimens were then rotated 90° to acquire second tilt series (b-stacks). Tomograms were calculated using the IMOD software package (http://bio3d.colourado.edu/imod)^62^ by applying an R-weighted back-projection algorithm, and a- and b-stacks were joined to generate double-tilt electron tomograms^63^. Microtubules were automatically segmented using the ZIBAmira (Zuse Institute Berlin, Germany) software package^57,64,65^ and TARDIS^66^. The automatic segmentation of the microtubules was followed by a visual inspection of the traced microtubules and correction of the individual microtubule tracings. Corrections included: manual tracing of undetected microtubules, connection of microtubules and deletions of tracing artifacts (for example, membranes of vesicles). For each data set, we quantitatively determined the total number of microtubules with either both ends or one end (i.e., the putative plus end) positioned in between the segregated chromatids in late anaphase^12^. Additionally, the chromatids from the tomograms were segmented using µSAM software^67^ and the Zeiss arivis cloud/APEER software (https://www.arivis.cloud/home/), which are AI-based segmentation tools for automatic segmentation of biological structures.

### Biophysical model analysis of the *C. elegans* spindle dynamics

We previously derived the stoichiometric model for asters under pulling forces from cortically anchored motors^17,32^. Here, we briefly go through the stoichiometric model and then present the modification to incorporate the interaction of asters with the chromosomes.

Consider two asters, where microtubules symmetrically nucleate at a rate 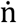, grow with a velocity V_g_, and undergo catastrophe and disassemble at a rate λ. Also, consider minus-end-directed motors of size r randomly distributed on the surface of the cell. A motor can bind to only one microtubule at a time (stoichiometry). Once bound, the motor exerts a pulling force f_0_ along the microtubule direction on the centrosome. The equation of motion of each centrosome (i = 1,2) is given by

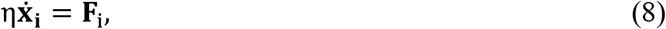

where η is the drag coefficient on the centrosome, and **F**_i_ is the total pulling force on the centrosome from the motors. The force is calculated from a surface field P_i_(**Y**, t) indicating the probability of binding a motor at a position **Y** at the time t to a microtubule from the ith centrosome and is given by

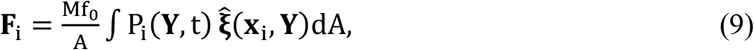

where M is the number of motors and A is the cell area. The integration is over the cell surface and 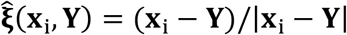 is a unit vector from the centrosome’s position to a point **Y** on the cell surface. For a low density of motors, where the overlap between them is negligible, the time evolution of the binding probability P_i_(**Y**, t) is given by

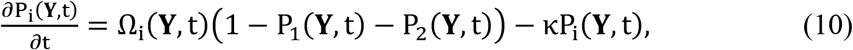

where Ω_i_(**Y**, t) is the rate of microtubule impingement from the ith centrosome on point **Y**, and κ is the motor unbinding rate. The first term on the right-hand side accounts for the stoichiometric binding of motors to microtubules (for the formulation of the model for overlapping or stacking motor distribution, see Young et al.^32^). To account for the interaction between centrosomes and chromosomes, we considered a viscous element of coefficient ν and the equation of motion is given by;

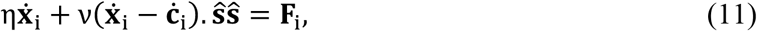

where 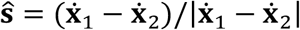 is a unit vector along the spindle axis. The equation for chromosome motion is presented in the main text.

We set up the simulation of spindle elongation at a given developmental stage as follows. The initial distance between the two asters is set by experimental measurements at the onset of anaphase, averaged over cells of that stage, with the distance of the two chromosomal plates set to zero. The radius *R* of the cell is set as the cube root of the experimentally measured averaged cell volume in that stage. For all developmental stages, we use the parameter values given in Table 1, except for the microtubule catastrophe rate *λ*, which we inferred from an optimization by minimizing the error function

**Table 1:** Model parameters for simulation of spindle and chromosome dynamics for all developmental stages.

| Parameter | Symbol | Range | Used value |
| --- | --- | --- | --- |
| Number of MTs per aster | $N_T$ | 3000-8000 | 5000 |
| MT growth velocity | $V_g$ | 0.5-2 [$\mu\text{m}/\text{s}$] | 1 [$\mu\text{m}/\text{s}$] |
| Motors density | $\rho$ | 0.05-0.2 [$1/\mu\text{m}^2$] | 0.12 [$1/\mu\text{m}^2$] |
| Motor size | $r$ | 0.5-2 [$\mu\text{m}$] | 1.5 [$\mu\text{m}$] |
| Motor detachment rate | $\kappa$ | 1/10-1/30 [1/s] | 1/20 [1/s] |
| Motor force | $f_0$ | 1-10 [pN] | 5 [pN] |
| Aster drag coefficient | $\eta$ | 300-600 [pNs/$\mu\text{m}$] | 450 [pNs/$\mu\text{m}$] |
| Spindle viscous coefficient | $\nu$ | 20-100 [pNs/$\mu\text{m}$] | 50 [pNs/$\mu\text{m}$] |

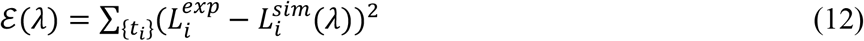

Here, the summation is over all time points *t*_*i*_ where we have experimental measurements, 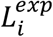 is the experimental measurement of pole-to-pole distance at time point *t*_*i*_, and 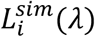 is the pole-to-pole distance at time *t* = *t*_*i*_ from a simulation where the microtubule catastrophe rate is set to *λ*. We use differential evolution, a global derivative-free optimizer, and search for *λ* ∈ [0.001,10]. In each generation, we perform 50 simulations and select candidates to minimize *ℰ*(*λ*). The values of pole-to-pole distance at the onset of anaphase, estimated cell radius, and microtubule catastrophe rate for developmental stages are given in Table 2.

**Table 2:** Model parameters for simulation of spindle and chromosome dynamics for each developmental stages.

| Cell stage | Cell radius | Initial pole-to-pole distance | Inferred MT catastrophe rate |
| --- | --- | --- | --- |
| 2 | 23 [$\mu m$] | 13 [$\mu m$] | 0.038 [1/s] |
| 4 | 18 [$\mu m$] | 11.2 [$\mu m$] | 0.086 [1/s] |
| 8 | 14 [$\mu m$] | 9 [$\mu m$] | 0.164 [1/s] |
| 16 | 11 [$\mu m$] | 7.5 [$\mu m$] | 0.332 [1/s] |
| 32 | 9 [$\mu m$] | 6.4 [$\mu m$] | 0.615 [1/s] |
| 64 | 7.3 [$\mu m$] | 5.4 [$\mu m$] | 0.945 [1/s] |

## Supporting information

Supplementary material for spindle lineage atlas

## Acknowledgement

The authors would like to thank Dr. Nicola Maghelli and the Advanced Light Microscopy Facility at MPI-CBG (Dresden, Germany) for technical assistance in using the custom-built LLSM. We thank Dr. Bob Goldstein’s lab (UNC Chapel Hill, USA) and Tony Hyman’s lab (MPI-CBG, Dresden, Germany) for providing reagents. Some strains were provided by the CGC, which is funded by NIH Office of Research Infrastructure Programs (P40 OD010440). The authors are grateful to Dr. Tobias Fürstenhaupt (Electron Microscopy Facility, MPI-CBG, Dresden, Germany) for his technical support in electron tomography, Dr. Robert Kiewisz (NYSBC, USA) for his assistance with developing and updating new functions on the TARDIS software package^66^ to optimize our data analysis, and to Hanna-Margareta Schwarzbach for her assistance in tomographic reconstruction and microtubule segmentation. The computations in this work were performed at facilities supported by the Scientific Computing Core at the Flatiron Institute, a division of the Simons Foundation. We acknowledge support from the CCB_X_ program of the Center for Computational Biology of the Flatiron Institute. Research in the Müller-Reichert lab was also supported by the Deutsche Forschungsgemeinschaft (DFG, grant number 258577783).

