## Supplementary Information for "Cell size reduction scales spindle elongation but not chromosome segregation in *C. elegans*"

### Index of supplementary information

#### Supplementary Figures:

Figure S1: Spindle length and the chromosome dynamics of different embryonic conditions.
Figure S2: Cell lineage quantitative analysis of spindle dynamics in a single embryo.
Figure S3: Laser ablation probes different forces in the early *C. elegans* embryo.
Figure S4: Mitotic lineage atlas of early *C. elegans* embryonic cell stage.
Figure S5: Models for spindle length dynamics.
Figure S6: Spindle elongation rate as a function of cell size.
Figure S7: Model simulation for spindle length dynamics.

#### Supplementary Tables:

Table S1: *P*-value for the pole-to-pole elongation derived from the Welch t-test rate for the analyzed embryonic conditions.
Table S2: *P*-value for the chromosome-to-chromosome segregation rate derived from the Welch t-test rate for the analyzed embryonic conditions.
Table S3: Summary of the *C. elegans* mitotic spindle dynamics parameters.
Table S4: Mitotic spindle dynamics parameter comparison for different *C. elegans* embryonic conditions.
Table S5: Anaphase spindle microtubule quantification for different cell stages.
Table S6: Quantification of the poles elongation and the chromosome separation rate during and after laser ablation for the spindles at different embryonic cell stages.

#### Figure S1

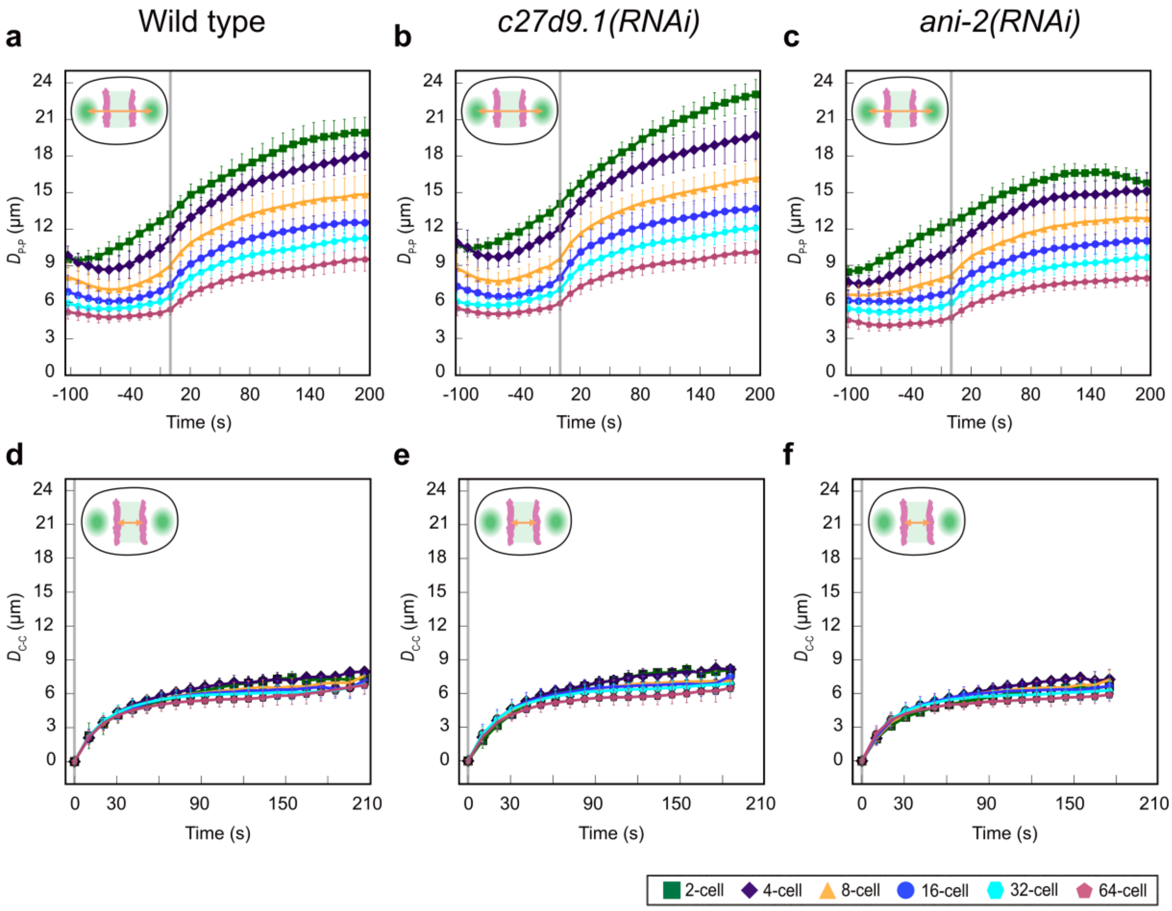


**Figure S1| Spindle length and the chromosome dynamics of different embryonic conditions. a-c,** Pole-to-pole distance, $D_{P\text{-}P}$, given as a function of time for the wild-type, *c27d9.1(RNAi)*, and *ani-2(RNAi)* embryonic conditions, respectively, of different embryonic cell stages. Sample size, n, is indicated in Supplementary Table 1. **d-f,** Chromosome-to-chromosome distance, $D_{C\text{-}C}$, given as a function of time for the wild-type, *c27d9.1(RNAi)*, and *ani-2(RNAi)* embryonic conditions of different embryonic cell stages. Sample size(n), is indicated in Supplementary Table 2. Each cell stage is indicated with a unique color code and symbol as shown in the plot legend. The inset on the top left of each plot indicates a schematic of the spindle with the poles in green and chromosomes in magenta, with the orange arrow showing the measured distances.

#### Figure S2

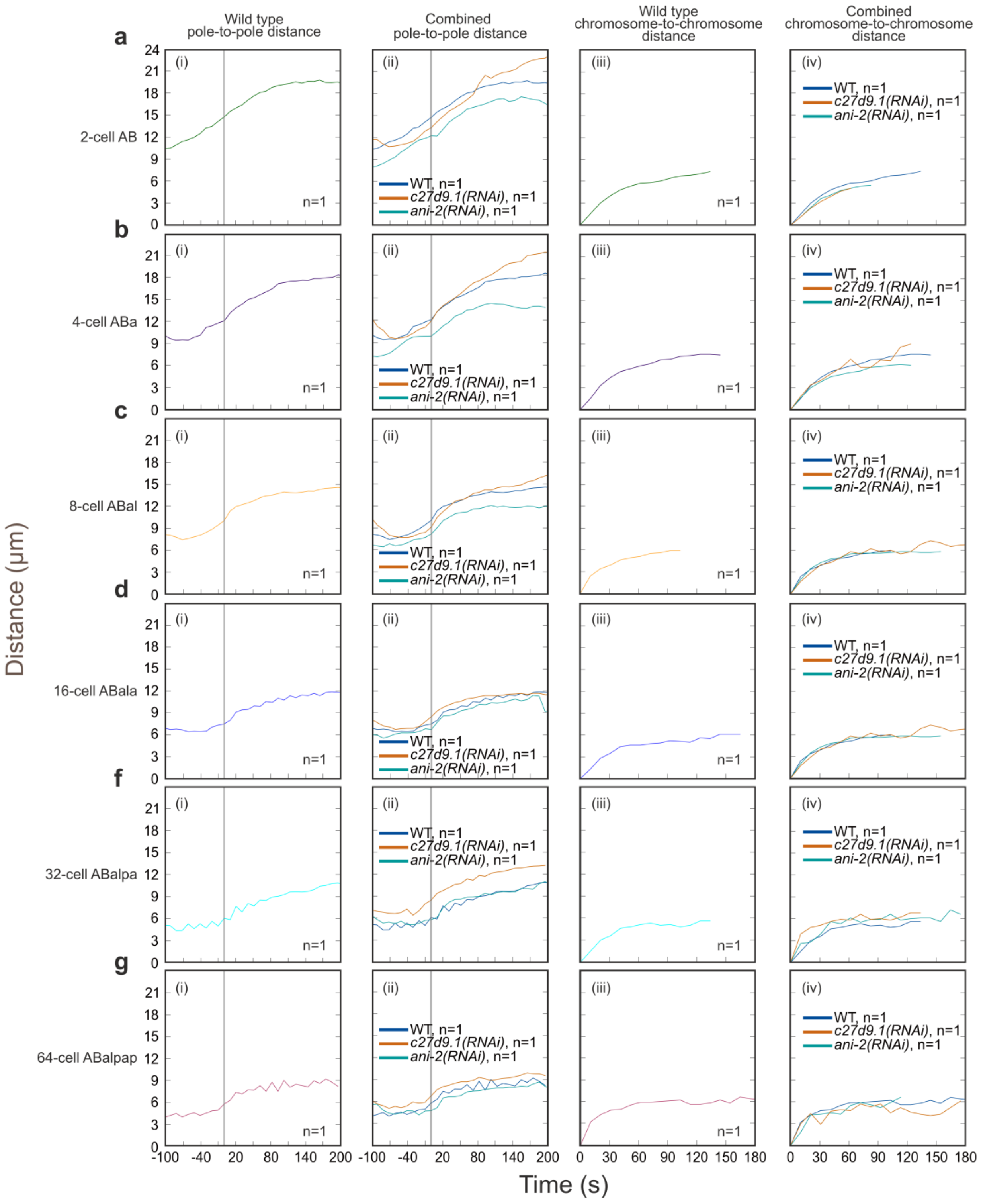


**Figure S2| Cell lineage quantitative analysis of spindle dynamics in a single embryo. a**(i)-**g**(i)**,** Wild-type embryo spindle pole-to-pole distance given as a function of time for a 2-cell AB, 4-cell ABa, 8-cell ABal, 16-cell ABala, 32-cell ABalpa, and 64-cell ABalpap. n = 1, for each cell type of each embryonic cell stage. **a**(ii)**-g**(ii)**,** Spindle pole-to-pole distance, given as a function of time for the cell types in (a(i)-g(i)), of the wild-type, *c27d9.1(RNAi)* and *ani-*

*2(RNAi)* embryonic conditions (n = 1, for each cell type of each embryonic cell stage). **a**(iii)-**g**(iii)**,** Wild type embryo spindle chromosome-to-chromosome distance given as a function of time for a 2-cell AB, 4-cell ABa, 8-cell ABal, 16-cell ABala, 32-cell ABalpa, and 64-cell ABalpap (n = 1, for each cell type of each embryonic cell stage). **a**(iv)-**g**(iv)**,** Spindle chromosome-to-chromosome distance, given as a function of time for the cell types in (a(iii)-g(iii)), of the Wild type, *c27d9.1(RNAi)*, and *ani-2(RNAi)* embryonic conditions (n = 1, for each cell type of each embryonic cell stage).

### Figure S3

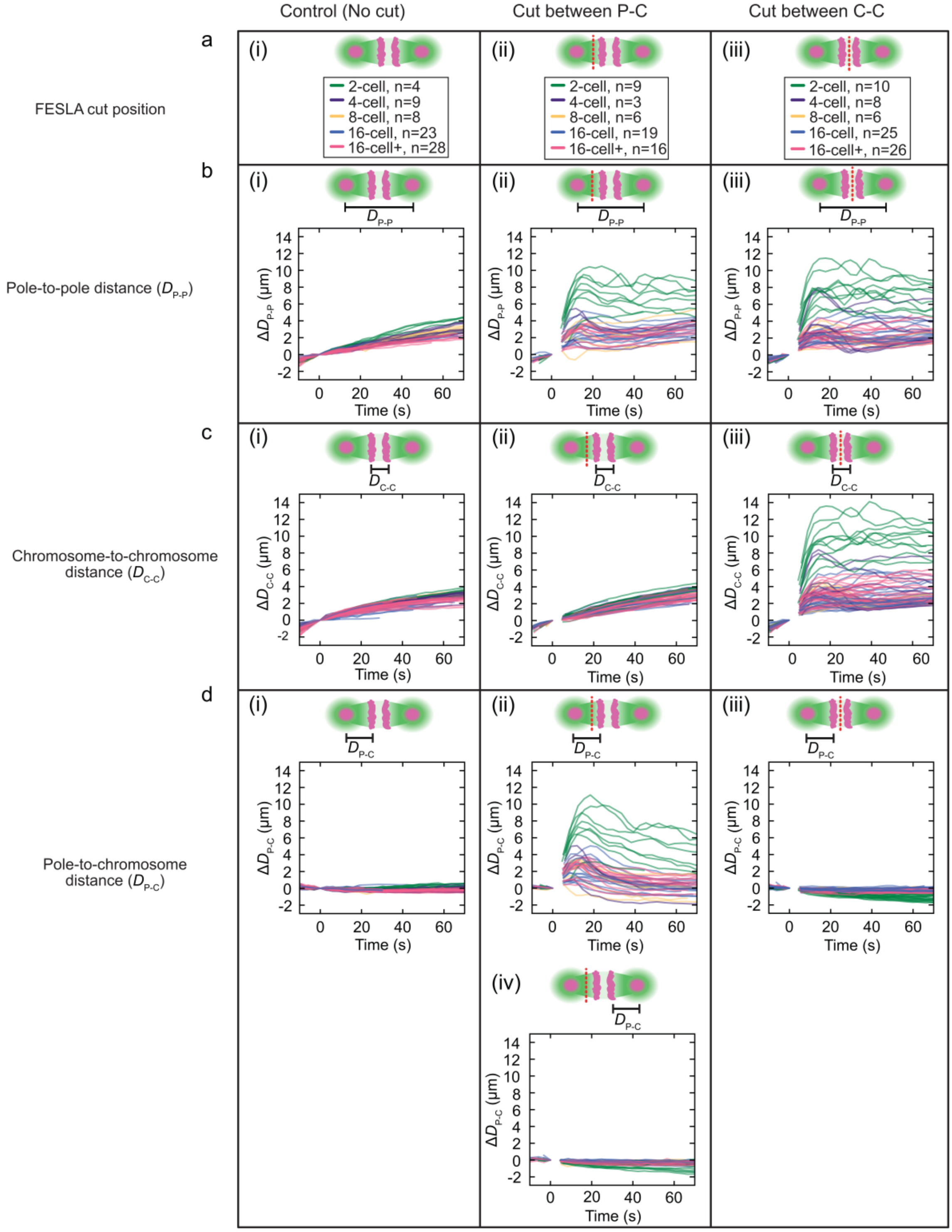


**Figure S3| Laser ablation probes different forces in the early *C. elegans* embryo. a,** Schematics illustrating the positions of laser ablation on the mitotic spindle during FESLA experiment, where (i) shows the control when there is no laser cut on the spindle, and the number of sample, n, for spindles in each cell stage with the corresponding color code, (ii) shows the laser cut, in red dash line, between the pole and chromosome of one half of the bipolar spindle, all in magenta, and the number of sample, n, for spindles in each cell stage with the corresponding color code, (iii) shows the laser cut in red dash line, between the chromosomes, in magenta, and the number of sample, n, for spindles in each cell stage with the corresponding color code. The microtubules are shown in green. **b,** Relative pole-to-pole distance, $D_{P-P}$, to the time of ablation cut at t = 0, given as a function of time; showing the (i) control, when there is no laser cut, (ii) dynamics of the spindles at each cell stage when there is a laser cut between the pole and chromosome on the first half of the bipolar spindle, and (iii) dynamics of the spindles at each cell stage when there is a laser cut between the chromosomes. The microtubules are shown in green. The black bar on the spindle schematics indicates the spindle region that is quantified on the corresponding plot. **c,** Relative chromosome-to-chromosome distance, $D_{C-C}$, to the time of ablation cut at t = 0, given as a function of time; showing the (i) control, when there is no laser cut, (ii) dynamics of the spindles at each cell stage when there is a laser cut between the pole and chromosome on the first half of the bipolar spindle, and (iii) dynamics of the spindles at each cell stage when there is a laser cut between the chromosomes. The microtubules are shown in green. The black bar on the spindle schematics indicates the spindle region that is quantified on the corresponding plot. **d,** Relative pole-to-chromosome distance, $D_{P-C}$, of the first half of the spindle, given as a function of time; showing the (i) control, when there is no laser cut, (ii) dynamics of the spindles at each cell stage when there is a laser cut between the pole and chromosome of first half of the bipolar spindle, and (iii) dynamics of the spindles at each cell stage when there is a laser cut between the chromosomes. (iv) Relative pole-to-chromosome distance, $D_{P-C}$, of the second half of the spindle, given as a function of time when there is a laser cut between the pole and chromosomes on the first half of the spindle. The microtubules are shown in green. The black bar on the spindle schematics indicates the spindle region that is quantified on the corresponding plot.

### Figure S4

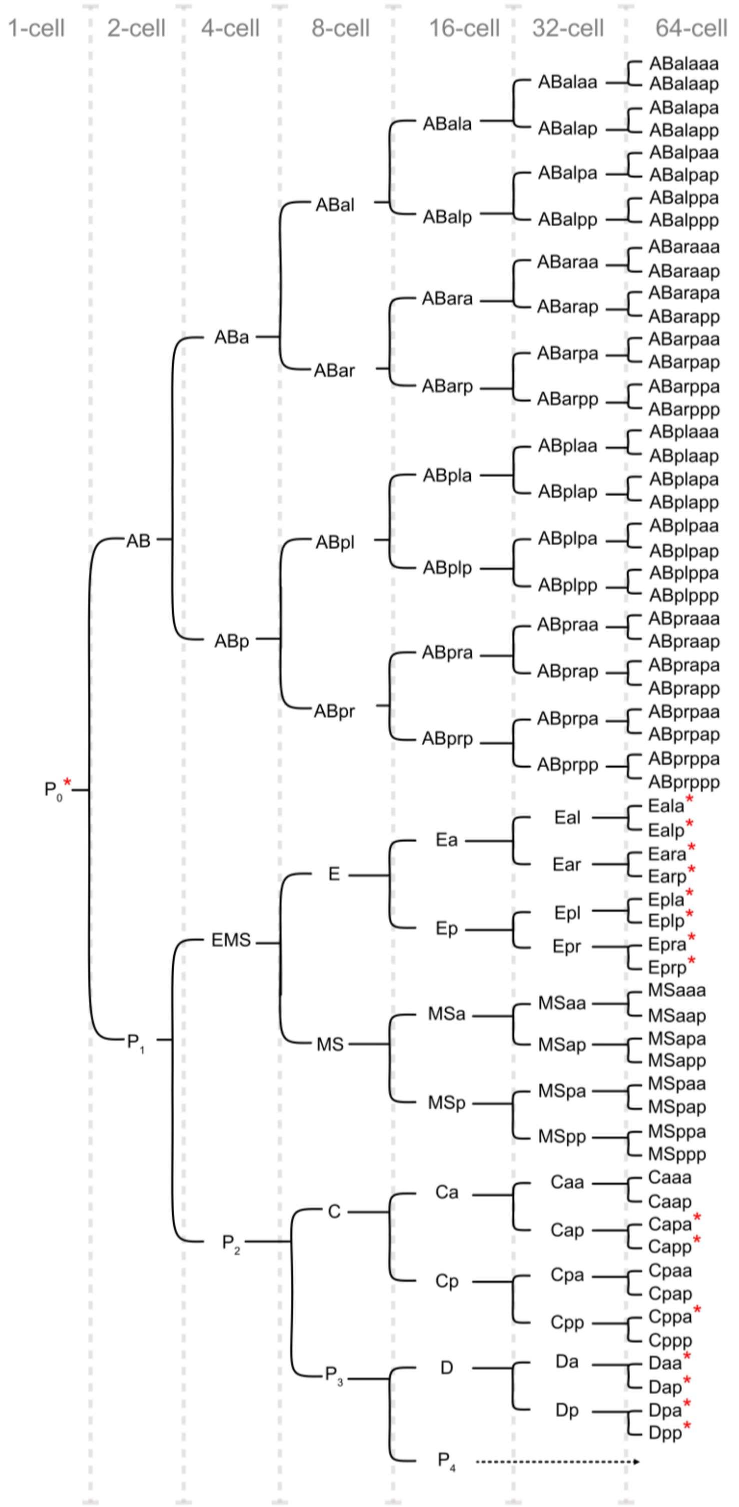


**Figure S4| Mitotic lineage atlas of early *C. elegans* embryonic cell stage.** Lineage atlas of the developing *C. elegans* embryo, reconstructed from Sulston et al., 1983[1]. The lineage atlas shows the identity of the cells in each embryonic cell stage whose spindles were analyzed. The cells marked with red asterisks were not analyzed or included in this study (cell within the center of the embryo were more challenging to resolve). The 16-cell stage cell, $P_4$, does not go into division until the 120-cell stage (black dashed arrow).

#### Figure S5

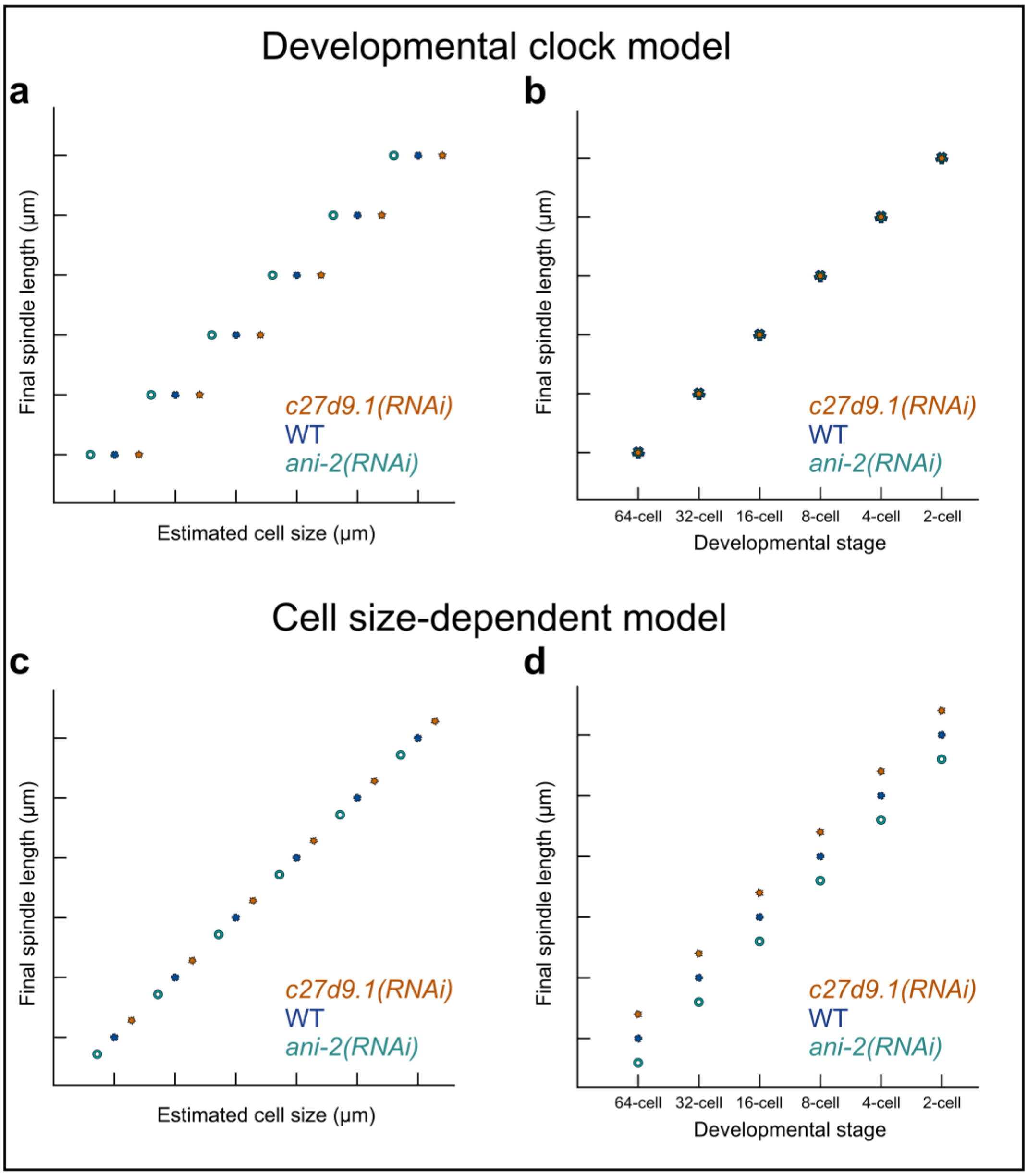


**Figure S5| Models for spindle length dynamics. a,b**, Developmental clock model showing (**a**) final spindle length as a function of estimated cell size and (**b**) final spindle length as a function of developmental stage for three embryonic conditions. **c,d**, Cell size–dependent model showing (**c**) final spindle length as a function of estimated cell size and (**d**) final spindle length as a function of developmental stage for the same three embryonic conditions. The three conditions are: wild type, blue; *c27d9.1(RNAi),* orange; *ani-2(RNAi)*, cyan.

### Figure S6

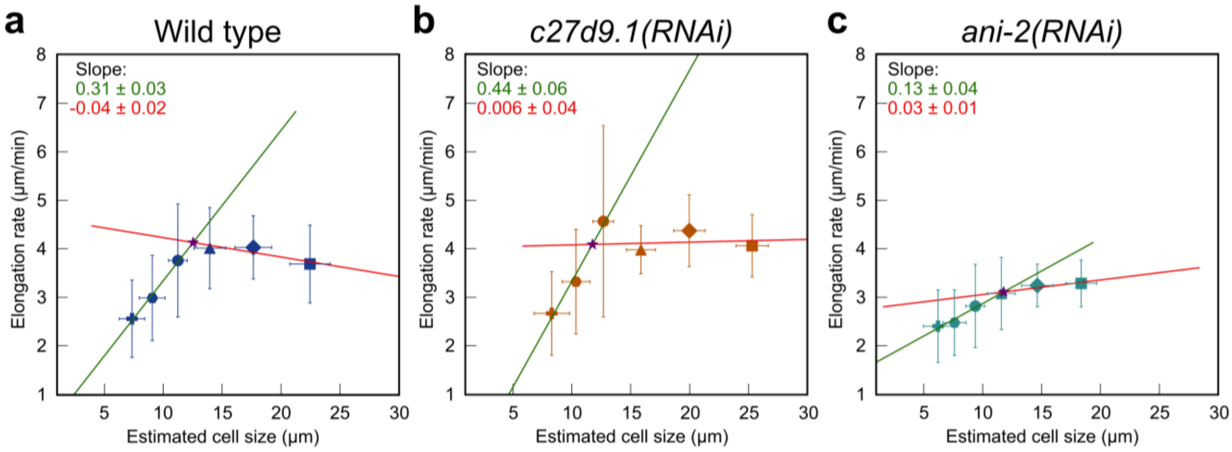


**Figure S6| Spindle elongation rate as a function of cell size.** Spindle elongation rates are plotted against cell size for **a,** wild type; **b,** *c27d9.1(RNAi)*; and **c,** *ani-2(RNAi)* embryos. Each plot shows a two-piecewise linear regression fit with corresponding slopes. The intersection point is indicated by a purple star.

### Figure S7

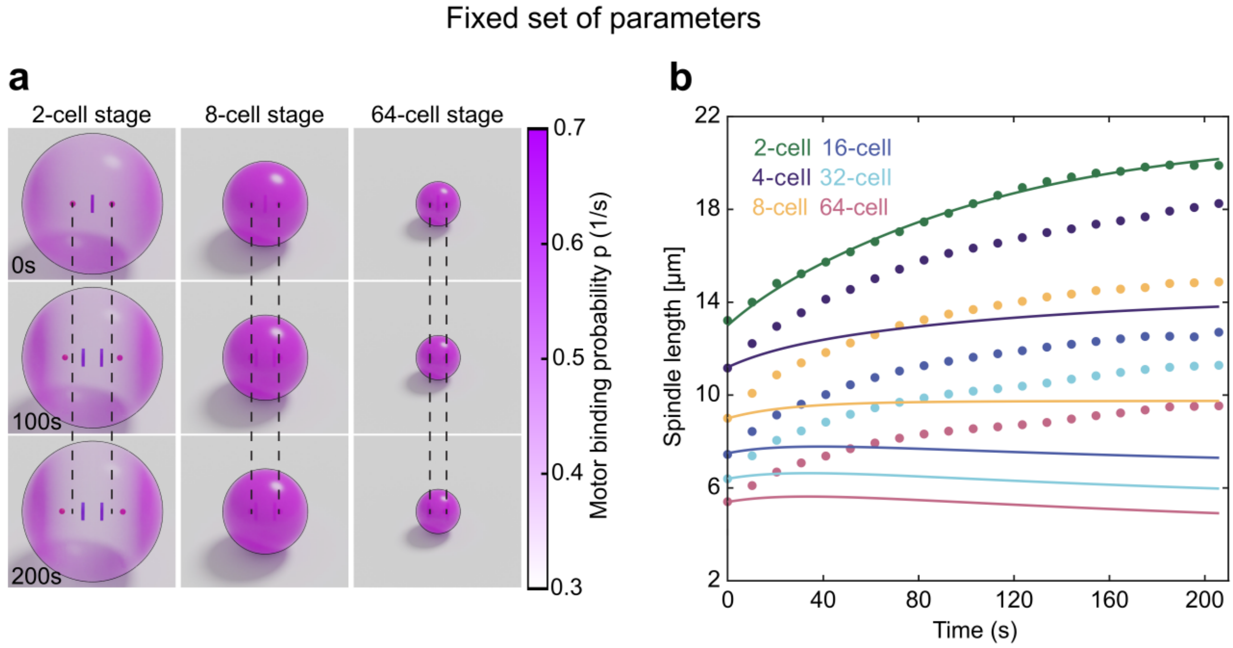


**Figure S7| Model simulation for spindle length dynamics. a,** Snapshots from simulations of 2-, 8-, and 64-cell stage cells. The binding probability of the motors is shown in pink. Dashed lines show the initial position of centrosomes. **b,** Spindle length as a function of time for different cell stages (lines show simulation, and dots indicate experimental measurements). The time t=0 corresponds to the anaphase onset.

#### Supplementary Movie Legends

**Movie S1:** LLSM movie of *C. elegans* WT embryo from the 2-cell stage to the late embryonic cell stage. The upper panel shows the two-dimensional view of the embryo to visualize the weak chromosome signal, and the lower panel shows the maximum Z-projection view. Time resolution: 10.3 s.

**Movie S2:** Movie of a 3D segmented and rendered mitotic *C. elegans* embryo in 32-cell stage. The one-frame movie was created using the data from the wide-type embryonic data. It shows the cells at different stages (interphase, prophase, metaphase, and anaphase) during development. The spindles in their corresponding cells are highlighted in green for the centrosomes and magenta for the chromosomes.

**Movie S3:** LLSM movie of *C. elegans ani-2(RNAi)* embryo from the 2-cell stage to the late embryonic cell stage. The upper panel shows the two-dimensional view of the embryo to visualize the weak chromosome signal, and the lower panel shows the maximum Z-projection view. Time resolution: 10.3 s.

**Movie S4:** LLSM movie of *C. elegans c27d9.1(RNAi)* embryo from the 2-cell stage to the late embryonic cell stage. The upper panel shows the two-dimensional view of the embryo to visualize the weak chromosome signal, and the lower panel shows the maximum Z-projection view. Time resolution: 10.3 s.

**Movie S5:** FESLA experiment with a wide-type *C. elegans* embryo. The laser cut between the pole and chromosomes is indicated with two red arrows at t = 0 sec. Scale bar, 10 µm.

#### Supplementary Material

Extended supplementary material for the mitotic spindle atlas for Fig. 2. This folder contains subdirectories with datasets quantifying spindle length and chromosome dynamics across all cell types from the 2- to 64-cell stages, for each analyzed embryonic condition (wild type, *ani-2(RNAi)*, and *c27d9.1(RNAi)* embryos). Each subfolder includes a spreadsheet summarizing the analyzed results for the corresponding datasets.

**Table S1.** ***P*-value for the pole-to-pole elongation derived from the Welch t-test rate for the analyzed embryonic conditions**

| Wild type | | | | | | |
|---|---|---|---|---|---|---|
| | 2-cell | 4-cell | 8-cell | 16-cell | 32-cell | 64-cell |
| 2-cell | 1 | | | | | |
| 4-cell | 0.17312239 | 1 | | | | |
| 8-cell | 0.18201251 | 0.920051 | 1 | | | |
| 16-cell | 0.76458859 | 0.080965 | 0.0751 | 1 | | |
| 32-cell | 0.00612913 | 1.33E-10 | 1.38E-14 | 3.41E-10 | 1 | |
| 64-cell | 0.00010947 | 3.46E-15 | 1.48E-23 | 1.62E-21 | 4.69E-08 | 1 |
| n | 15 | 33 | 70 | 135 | 213 | 279 |
| *c27d9.1(RNAi)* | | | | | | |
| 2-cell | 1 | | | | | |
| 4-cell | 0.243987 | 1 | | | | |
| 8-cell | 0.714287 | 0.014885 | 1 | | | |
| 16-cell | 0.094467 | 0.422763 | 0.005316 | 1 | | |
| 32-cell | 0.007267 | 3.83E-08 | 1.51E-09 | 3.10E-08 | 1 | |
| 64-cell | 7.57E-05 | 9.05E-14 | 1.22E-31 | 9.56E-16 | 2.76E-10 | 1 |
| n | 10 | 29 | 58 | 102 | 180 | 208 |
| *ani-2(RNAi)* | | | | | | |
| 2-cell | 1 | | | | | |
| 4-cell | 0.001966 | 1 | | | | |
| 8-cell | 1.92E-08 | 2.56E-10 | 1 | | | |
| 16-cell | 8.61E-09 | 1.56E-17 | 3.83E-19 | 1 | | |
| 32-cell | 3.50E-09 | 4.55E-21 | 4.18E-36 | 7.10E-33 | 1 | |
| 64-cell | 1.68E-09 | 1.30E-23 | 1.88E-45 | 1.46E-72 | 4.80E-55 | 1 |
| n | 8 | 30 | 62 | 107 | 157 | 148 |

**Table S2. *P*-value for the chromosome-to-chromosome segregation rate derived from the Welch t-test rate for the analyzed embryonic conditions**

| Wild type | | | | | | |
|---|---|---|---|---|---|---|
| | 2-cell | 4-cell | 8-cell | 16-cell | 32-cell | 64-cell |
| 2-cell | 1 | | | | | |
| 4-cell | 0.80017772 | 1 | | | | |
| 8-cell | 0.93054264 | 0.634831 | 1 | | | |
| 16-cell | 0.89190577 | 0.18432 | 0.452379 | 1 | | |
| 32-cell | 0.37002153 | 0.000208 | 0.00115 | 0.000597 | 1 | |
| 64-cell | 0.5083043 | 0.001862 | 0.009554 | 0.009784 | 0.232402 | 1 |
| n | 17 | 30 | 65 | 130 | 206 | 263 |
| *c27d9.1(RNAi)* | | | | | | |
| 2-cell | 1 | | | | | |
| 4-cell | 0.01236 | 1 | | | | |
| 8-cell | 0.000853 | 0.193446 | 1 | | | |
| 16-cell | 4.16E-06 | 0.001115 | 0.033454 | 1 | | |
| 32-cell | 1.48E-06 | 3.53E-06 | 0.001017 | 0.672624 | 1 | |
| 64-cell | 0.000103 | 0.010033 | 0.334798 | 0.092195 | 0.000827 | 1 |
| n | 11 | 26 | 52 | 98 | 174 | 204 |
| *ani-2(RNAi)* | | | | | | |
| 2-cell | 1 | | | | | |
| 4-cell | 0.007965 | 1 | | | | |
| 8-cell | 0.002069 | 0.687348 | 1 | | | |
| 16-cell | 1.27E-05 | 0.003707 | 0.002827 | 1 | | |
| 32-cell | 1.37E-07 | 4.79E-07 | 1.34E-08 | 0.002517 | 1 | |
| 64-cell | 1.32E-09 | 1.72E-08 | 1.16E-09 | 8.47E-05 | 0.140927 | 1 |
| n | 8 | 31 | 62 | 113 | 148 | 126 |

##### Table S3. Summary of the *C. elegans* mitotic spindle dynamics parameters

| **Wild type** | | | | | | |
|---|---|---|---|---|---|---|
| | 2-cell | 4-cell | 8-cell | 16-cell | 32-cell | 64-cell |
| **Cell volume (μm³)** | 11337.00 ± 2597.08 | 5499.32 ± 1455.04 | 2715.33 ± 784.27 | 1423.68 ± 302.11 | 745.53 ± 268.61 | 397.08 ± 175.15 |
| **Estimated cell size (μm)** | 22.46 | 17.65 | 13.95 | 11.25 | 9.07 | 7.35 |
| **Error propagation** | 1.72 | 1.56 | 1.34 | 0.80 | 1.09 | 1.08 |
| **Initial pole-to-pole distance (μm)** | 7.38 ± 1.91 | 8.38 ± 0.87 | 7.01 ± 0.86 | 5.87 ± 1.13 | 5.20 ± 0.65 | 4.67 ± 0.44 |
| **Final pole-to-pole distance (μm)** | 20.46 ± 1.52 | 17.91 ± 1.43 | 14.57 ± 1.47 | 12.44 ± 0.89 | 11.01 ± 0.76 | 9.29 ± 0.88 |
| **Final chromosome-to-chromosome distance (μm)** | 6.89 ± 0.73 | 7.14 ± 0.53 | 6.54 ± 0.65 | 6.33 ± 0.61 | 6.10 ± 0.56 | 5.6 ± 0.67 |
| **Metaphase pole-to-pole distance (μm)** | 13.27 ± 0.77 | 11.27 ± 1.06 | 9.13 ± 0.86 | 7.66 ± 0.67 | 6.78 ± 0.43 | 5.71 ± 0.36 |
| **Elongation rate of poles (μm/min)** | 3.69 ± 0.80 | 4.03 ± 0.65 | 4.01 ± 0.84 | 3.76 ± 1.17 | 2.99 ± 0.88 | 2.56 ± 0.80 |
| **Segregation speed of chromosomes (μm/min)** | 13.69 ± 8.90 | 13.10 ± 3.05 | 13.49 ± 4.63 | 14.00 ± 4.06 | 15.76 ± 5.18 | 15.20 ± 4.78 |
| ***ani-2(RNAi)*** | | | | | | |
| **Cell volume (μm³)** | 6195.37 ± 1342.83 | 3137.68 ± 867.48 | 1556.91 ± 475.71 | 822.81 ± 213.12 | 441.56 ± 171.12 | 239.37 ± 143.89 |
| **Estimated cell size (μm)** | 18.37 | 14.64 | 11.59 | 9.37 | 7.61 | 6.21 |
| **Error propagation** | 1.33 | 1.35 | 1.18 | 0.81 | 0.98 | 1.24 |
| **Initial pole-to-pole distance (μm)** | 7.52 ± 0.75 | 6.67 ± 1.80 | 6.23 ± 0.83 | 5.78 ± 0.79 | 5.01 ± 0.68 | 4.04 ± 0.43 |
| **Final pole-to-pole distance (μm)** | 16.60 ± 0.67 | 15.33 ± 1.41 | 12.93 ± 1.17 | 11.02 ± 0.91 | 9.44 ± 0.83 | 7.83 ± 0.58 |
| **Final chromosome-to-chromosome distance (μm)** | 6.36 ± 0.42 | 6.72 ± 0.69 | 6.54 ± 0.58 | 6.33 ± 0.70 | 5.93 ± 0.57 | 5.37 ± 0.52 |
| **Metaphase pole-to-pole distance (μm)** | 12.70 ± 0.89 | 10.55 ± 0.81 | 8.50 ± 0.88 | 7.20 ± 0.69 | 6.16 ± 0.50 | 4.91 ± 0.37 |
| **Elongation rate of poles (μm/min)** | 3.29 ± 0.48 | 3.24 ± 0.44 | 3.08 ± 0.74 | 2.82 ± 0.86 | 2.48 ± 0.68 | 2.40 ± 0.75 |
| **Segregation speed of chromosomes (μm/min)** | 11.03 ± 1.43 | 13.23 ± 2.86 | 13.50 ± 3.13 | 15.18 ± 4.08 | 16.88 ± 4.88 | 17.90 ± 6.24 |
| ***c27d9.1(RNAi)*** | | | | | | |
| **Cell volume (μm³)** | 16228.51 ± 2647.68 | 7973.19 ± 1594.60 | 4000.03 ± 934.68 | 2037.04 ± 428.12 | 1111.67 ± 380.67 | 570.53 ± 311.46 |
| **Estimated cell size (μm)** | 25.32 | 19.98 | 15.87 | 12.68 | 10.36 | 8.29 |
| **Error propagation** | 1.38 | 1.33 | 1.24 | 0.89 | 1.18 | 1.51 |
| **Initial pole-to-pole distance (μm)** | 8.58 ± 1.07 | 9.29 ± 1.22 | 7.51 ± 0.84 | 6.23 ± 1.14 | 5.36 ± 0.78 | 4.87 ± 0.44 |
| **Final pole-to-pole distance (μm)** | 23.72 ± 1.20 | 19.56 ± 2.02 | 15.75 ± 1.35 | 13.45 ± 1.13 | 11.77 ± 0.83 | 9.90 ± 0.74 |
| **Final chromosome-to-chromosome distance (μm)** | 7.99 ± 0.53 | 7.62 ± 0.60 | 6.87 ± 0.49 | 6.81 ± 0.78 | 6.43 ± 0.56 | 5.78 ± 0.68 |
| **Metaphase pole-to-pole distance (μm)** | 14.16 ± 0.65 | 12.42 ± 0.98 | 9.90 ± 0.90 | 8.26 ± 0.73 | 7.36 ± 0.43 | 6.17 ± 0.43 |
| **Elongation rate of poles (μm/min)** | 4.06 ± 0.64 | 4.37 ± 0.74 | 3.98 ± 0.50 | 4.57 ± 1.97 | 3.32 ± 1.08 | 2.67 ± 0.86 |
| **Segregation speed of chromosomes (μm/min)** | 11.43 ± 1.81 | 13.39 ± 2.16 | 14.23 ± 3.36 | 15.86 ± 5.92 | 16.15 ± 4.18 | 14.75 ± 3.72 |

**Table S4. Mitotic spindle dynamics parameter comparison for different *C. elegans* embryonic conditions**

| | Dynamic parameters | Slope | p-value of the slope | t-statistic of the slope | $R^2$ |
|---|---|---|---|---|---|
| **Wild type** | Pole-to-pole final distance | 0.75 ± 0.03 | 0.0000214 | 22.93 | 0.99 |
| | Chromosome-to-chromosome final distance | 0.09 ± 0.02 | 0.0184743 | 3.84 | 0.79 |
| ***c27d9.1(RNAi)*** | Pole-to-pole final distance | 0.81± 0.01 | 0.0000005 | 58.94 | 1.00 |
| | Chromosome-to-chromosome final distance | 0.12 ± 0.02 | 0.0014016 | 7.88 | 0.94 |
| ***ani-2(RNAi)*** | Pole-to-pole final distance | 0.73 ± 0.07 | 0.0003732 | 11.11 | 0.97 |
| | Chromosome-to-chromosome final distance | 0.08 ± 0.04 | 0.1135826 | 2.02 | 0.50 |

**Table S5. Anaphase spindle microtubule quantification for different cell stages**

| MT classes | Measurements | 8-cell | 16-cell | 32-cell |
|---|---|---|---|---|
| **All MTs** | | 4094 | 1696 | 1937 |
| **Interpolar MT** | Number of MTs | 257 | 104 | 124 |
| | Avg. MT length (µm) ± std | 3.95 ± 0.75 | 3.55 ± 0.98 | 2.97 ± 0.77 |
| **Mid-spindle MT** | Number of MTs | 24 | 24 | 30 |
| | Avg. MT length (µm) ± std | 1.1 ± 0.6 | 1.92 ± 0.78 | 1.28 ± 0.56 |
| **KMT** | Number of MTs | 460 | 213 | 295 |
| | Avg. MT length (µm) ± std | 2.07 ± 0.5 | 1.32 ± 0.38 | 1.32 ± 0.39 |

**Table S6. Quantification of the poles elongation and the chromosome separation rate during and after laser ablation for the spindles at different embryonic cell stages**

| Pole-to-pole: $D_{\text{P-P}}$ 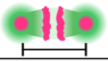 | | | | |
|---|---|---|---|---|
| | Stage | $\dot{D}_{\text{P-P(before)}}$ (μm/s) | $\dot{D}_{\text{P-P(after)}}$ (μm/s) | $\Delta\dot{D}_{\text{P-P}}$ (μm/s) |
| Control 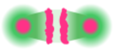 | 2-cell | 0.077±0.049 (n=4) | 0.062±0.039 (n=4) | -0.015±0.020 (n=4) |
| | 4-cell | 0.072±0.015 (n=7) | 0.054±0.021 (n=8) | -0.018±0.018 (n=7) |
| | 8cell | 0.074±0.028 (n=8) | 0.062±0.040 (n=7) | -0.005±0.054 (n=7) |
| | 16-cell | 0.058±0.025 (n=11) | 0.042±0.027 (n=13) | -0.014±0.038 (n=11) |
| | 16-cell+ | 0.065±0.036 (n=13) | 0.037±0.021 (n=17) | -0.026±0.036 (n=13) |
| Cut btw P-C 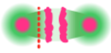 | 2-cell | 0.103±0.050 (n=7) | 0.691±0.179 (n=8) | 0.605±0.181 (n=7) |
| | 4-cell | 0.096±0.007 (n=2) | 0.246±0.142 (n=3) | 0.231±0.004 (n=2) |
| | 8cell | 0.105±0.043 (n=4) | 0.129±0.189 (n=5) | 0.099±0.096 (n=4) |
| | 16-cell | 0.059±0.026 (n=13) | 0.176±0.082 (n=13) | 0.112±0.083 (n=12) |
| | 16-cell+ | 0.090±0.029 (n=7) | 0.134±0.059 (n=11) | 0.041±0.076 (n=7) |
| Cut btw C-C 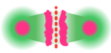 | 2-cell | 0.084±0.024 (n=10) | 0.638±0.161 (n=10) | 0.555±0.144 (n=10) |
| | 4-cell | 0.080±0.020 (n=8) | 0.386±0.224 (n=8) | 0.305±0.234 (n=8) |
| | 8cell | 0.075±0.016 (n=5) | 0.209±0.115 (n=5) | 0.135±0.124 (n=5) |
| | 16-cell | 0.076±0.035 (n=18) | 0.157±0.091 (n=16) | 0.081±0.092 (n=16) |
| | 16-cell+ | 0.069±0.039 (n=18) | 0.098±0.071 (n=18) | 0.029±0.080 (n=18) |
| Chromosome-to-chromosome: $D_{\text{C-C}}$ 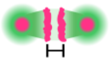 | | | | |
| | Stage | $\dot{D}_{\text{C-C(before)}}$ (μm/s) | $\dot{D}_{\text{C-C(after)}}$ (μm/s) | $\Delta\dot{D}_{\text{C-C}}$ (μm/s) |
| Control 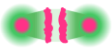 | 2-cell | 0.122±0.005 (n=4) | 0.074±0.007 (n=4) | -0.048±0.005 (n=4) |
| | 4-cell | 0.093±0.025 (n=9) | 0.074±0.018 (n=9) | -0.018±0.014 (n=9) |
| | 8cell | 0.091±0.018 (n=8) | 0.079±0.024 (n=8) | -0.012±0.021 (n=8) |
| | 16-cell | 0.089±0.034 (n=20) | 0.065±0.025 (n=20) | -0.014±0.028 (n=18) |
| | 16-cell+ | 0.100±0.028 (n=25) | 0.068±0.022 (n=25) | -0.031±0.026 (n=25) |
| Cut btw P-C 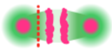 | 2-cell | 0.136±0.008 (n=7) | 0.064±0.033 (n=9) | -0.081±0.027 (n=7) |
| | 4-cell | 0.090±0.000 (n=1) | 0.039±0.041 (n=3) | -0.031±0.000 (n=1) |
| | 8cell | 0.129±0.085 (n=4) | 0.054±0.018 (n=6) | -0.082±0.076 (n=4) |
| | 16-cell | 0.094±0.035 (n=15) | 0.050±0.022 (n=18) | -0.048±0.036 (n=15) |
| | 16-cell+ | 0.118±0.035 (n=11) | 0.035±0.021 (n=15) | -0.090±0.037 (n=10) |
| Cut btw C-C 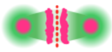 | 2-cell | 0.106±0.022 (n=10) | 0.745±0.187 (n=10) | 0.639±0.182 (n=10) |
| | 4-cell | 0.106±0.017 (n=8) | 0.412±0.210 (n=8) | 0.306±0.204 (n=8) |
| | 8cell | 0.098±0.036 (n=6) | 0.235±0.128 (n=6) | 0.137±0.107 (n=6) |
| | 16-cell | 0.093±0.036 (n=25) | 0.149±0.094 (n=25) | 0.056±0.093 (n=25) |
| | 16-cell+ | 0.116±0.030 (n=24) | 0.145±0.083 (n=24) | 0.029±0.081 (n=24) |
