## Supplementary figures and images for "Cell size reduction scales spindle elongation but not chromosome segregation in *C. elegans*"

### 2-cell_AB.png

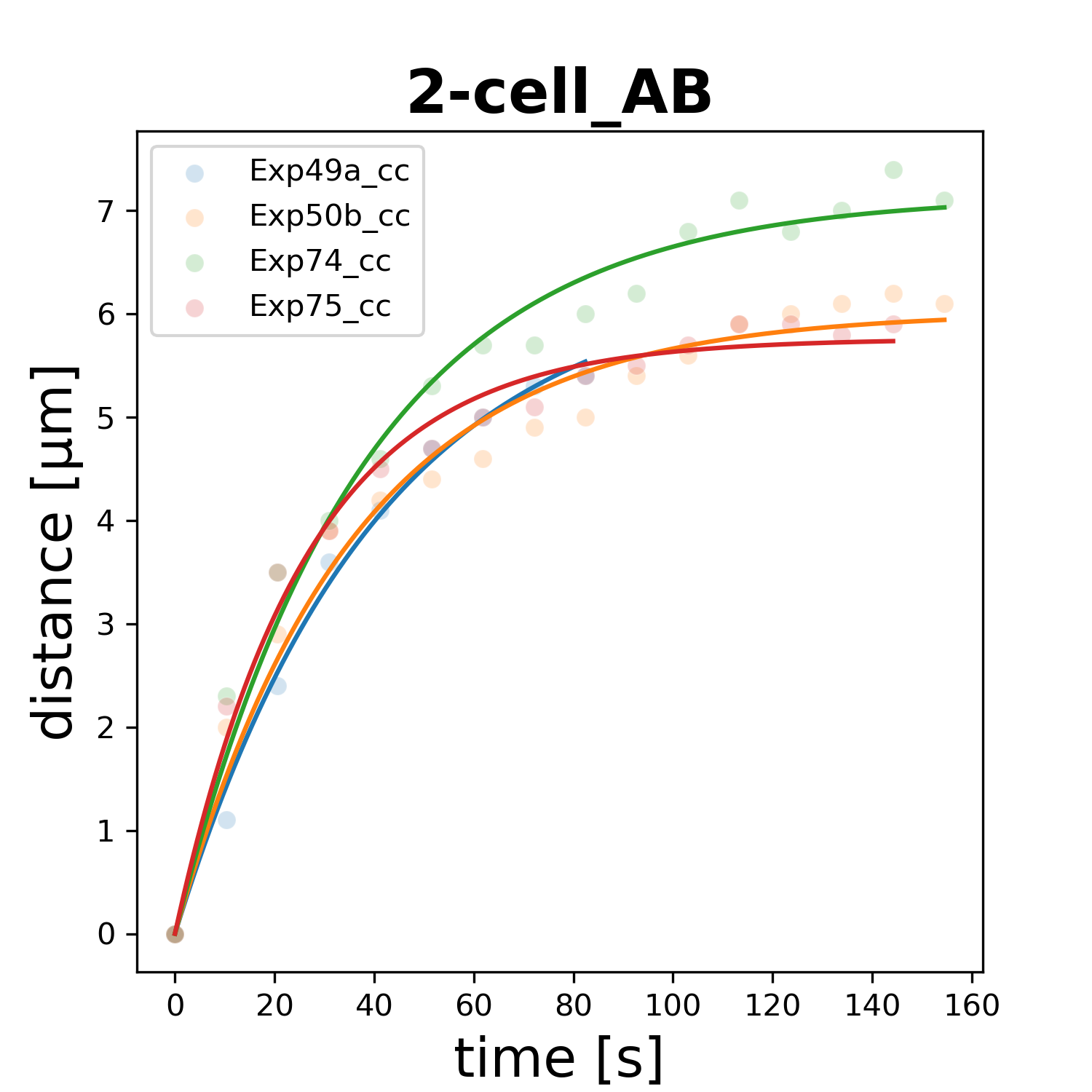

### 2-cell_P1.png

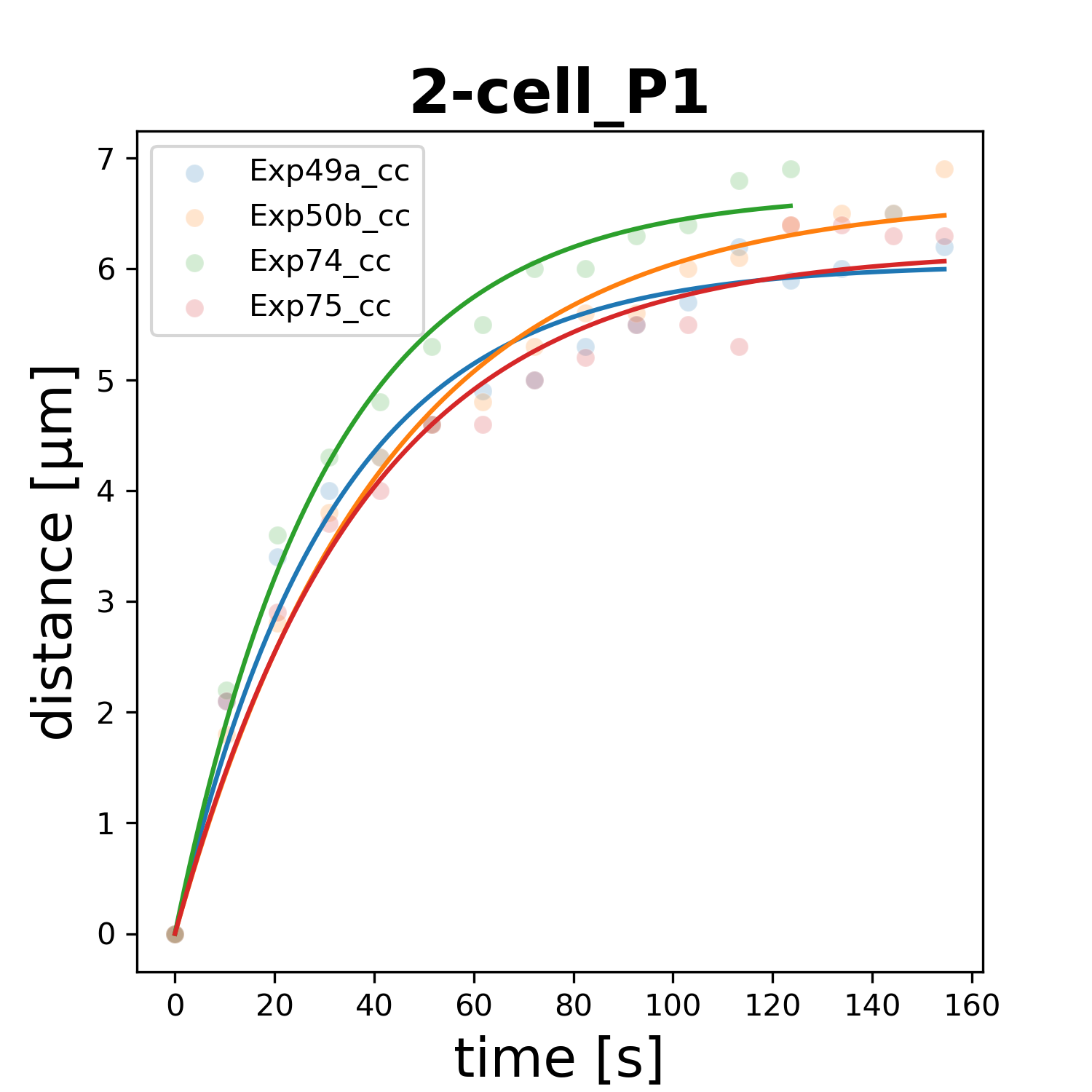

### 4-cell_ABa.png

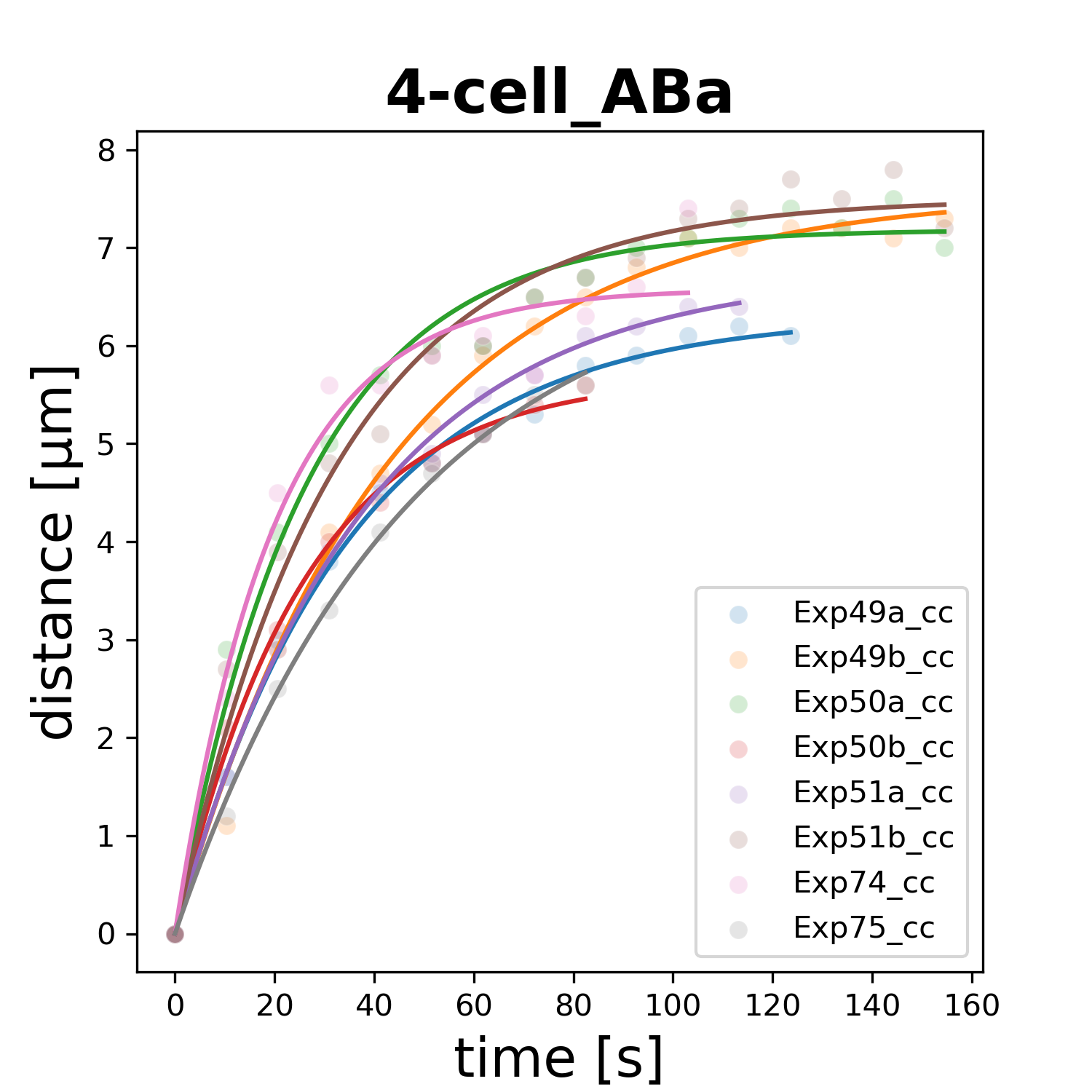

### 4-cell_ABp.png

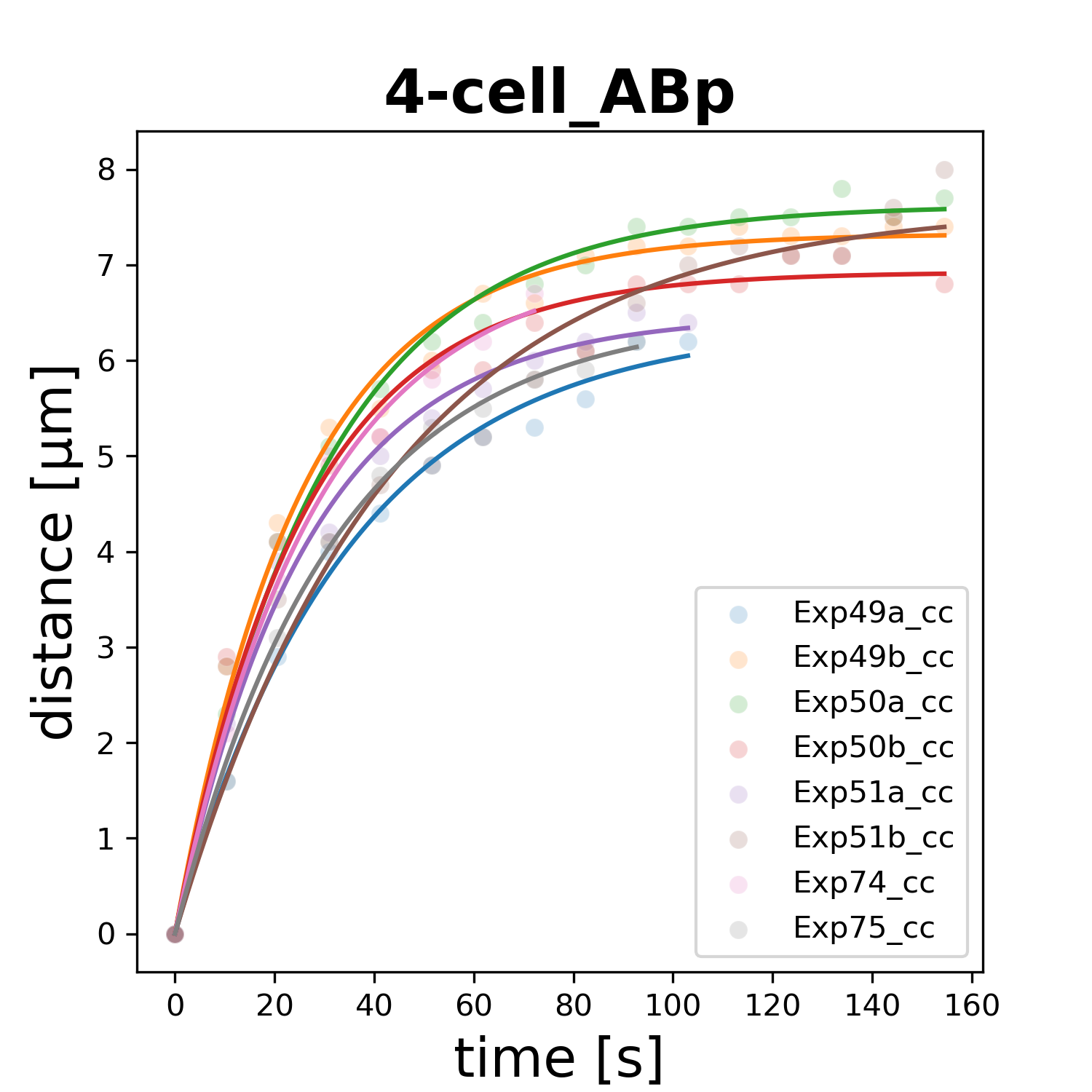

### 4-cell_EMS.png

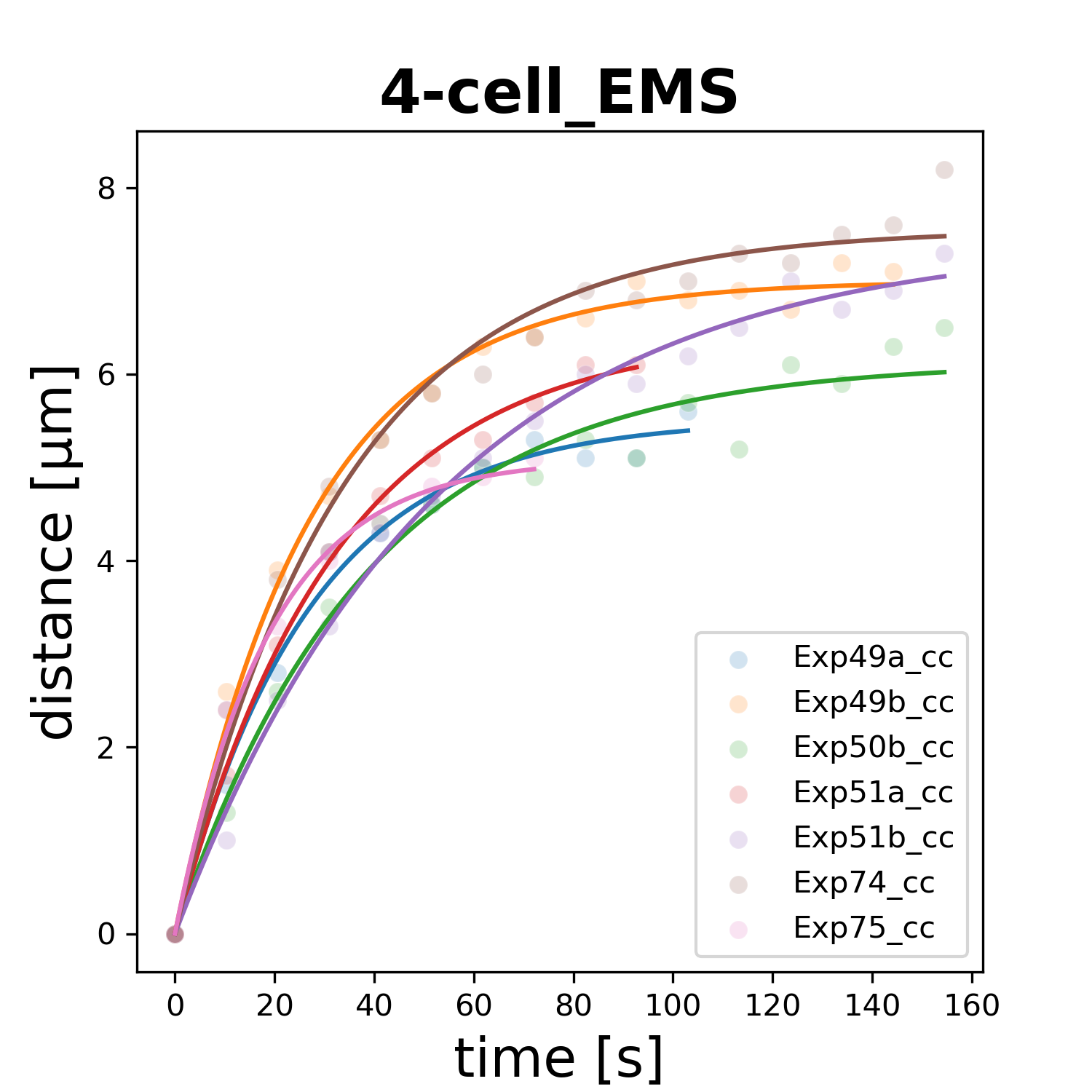

### 4-cell_P2.png

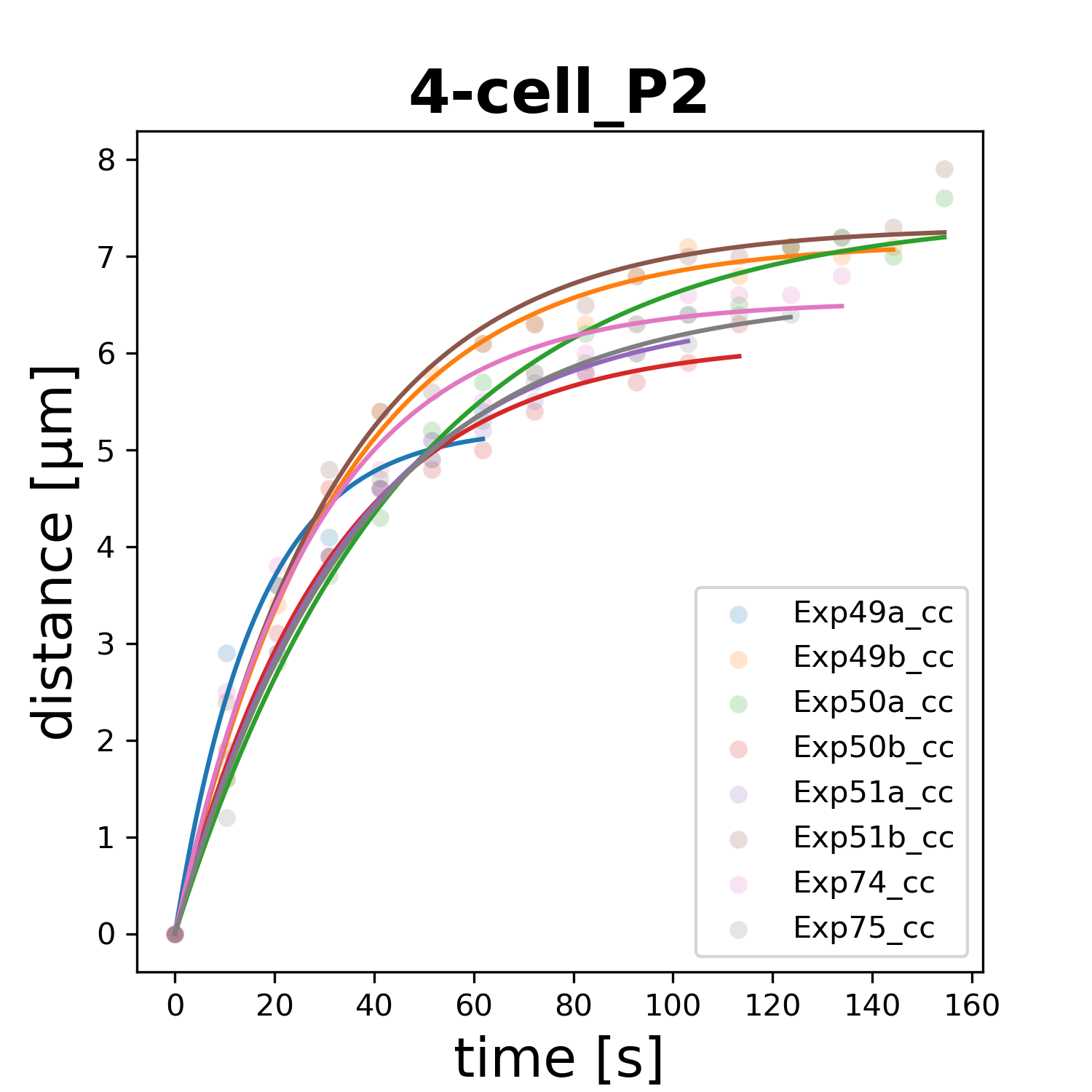

### 8-cell_ABal.png

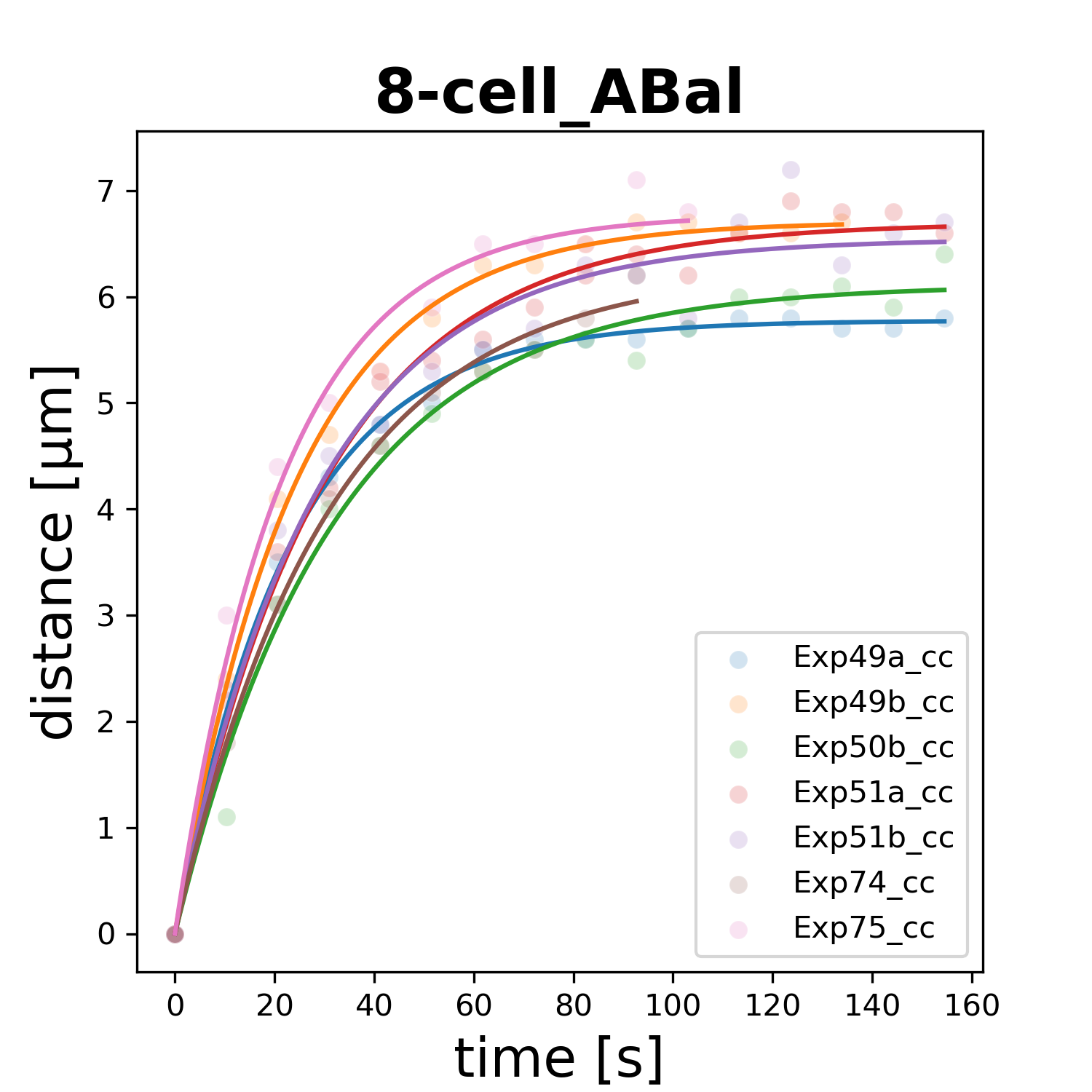

### 8-cell_ABar.png

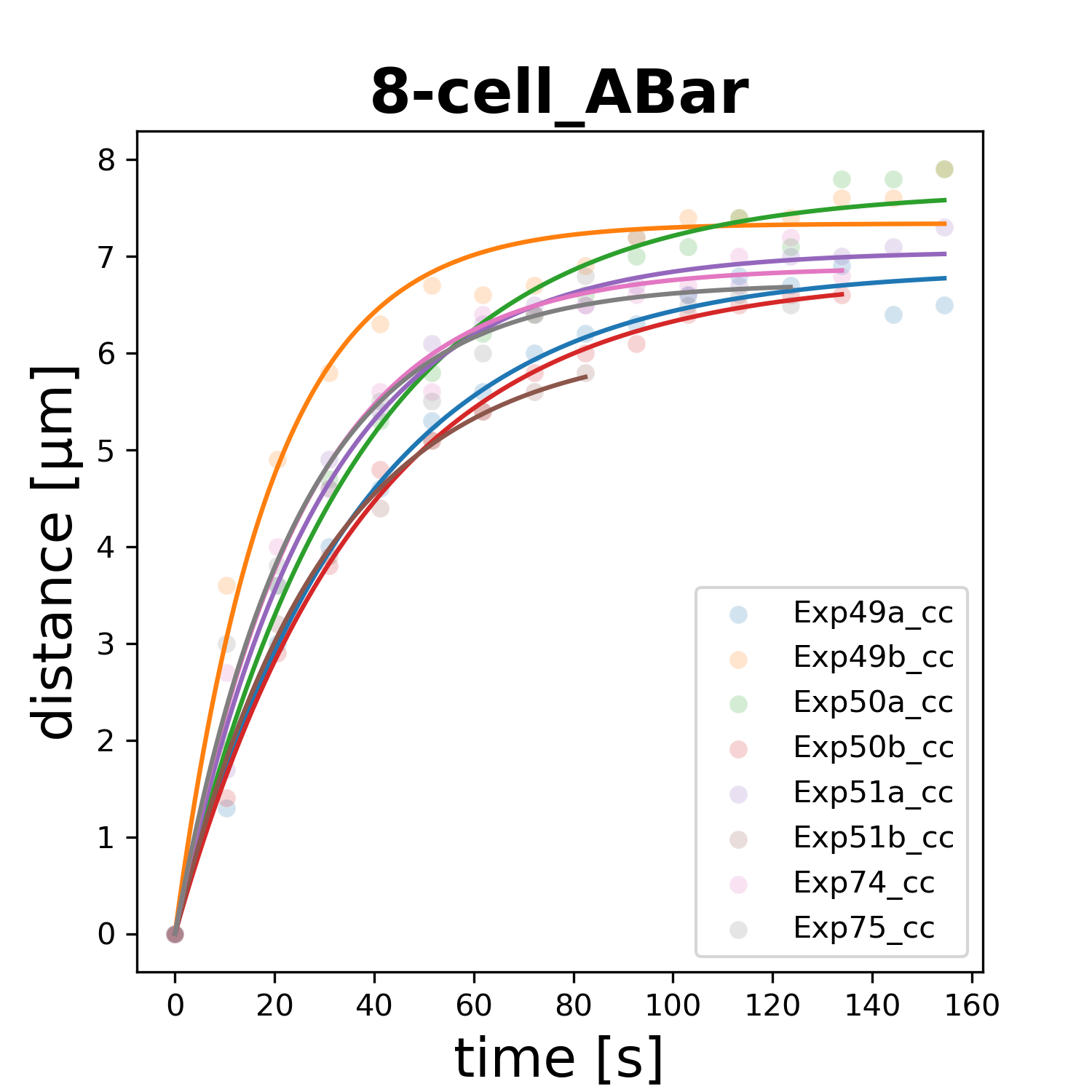

### 8-cell_ABpl.png

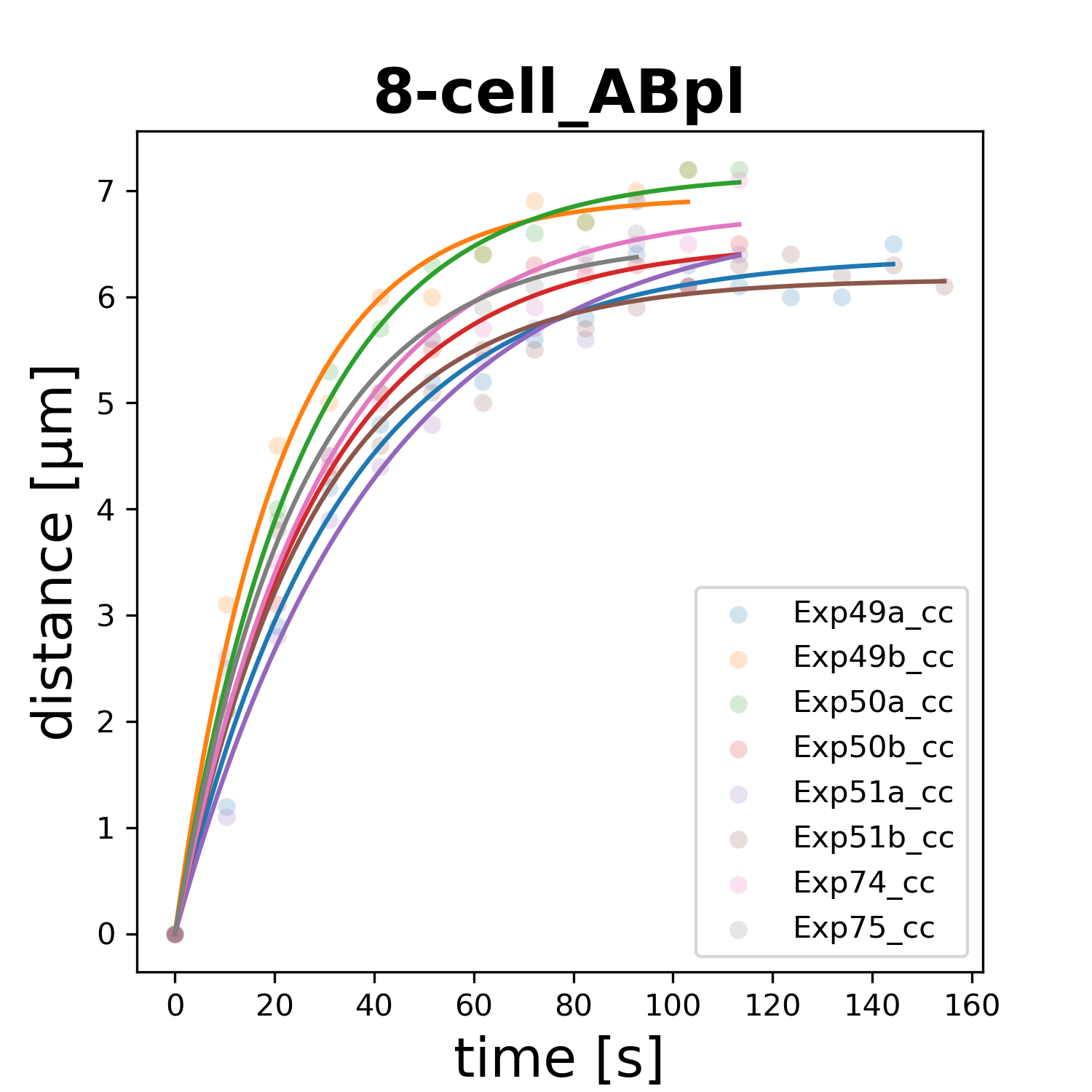

### 8-cell_ABpr.png

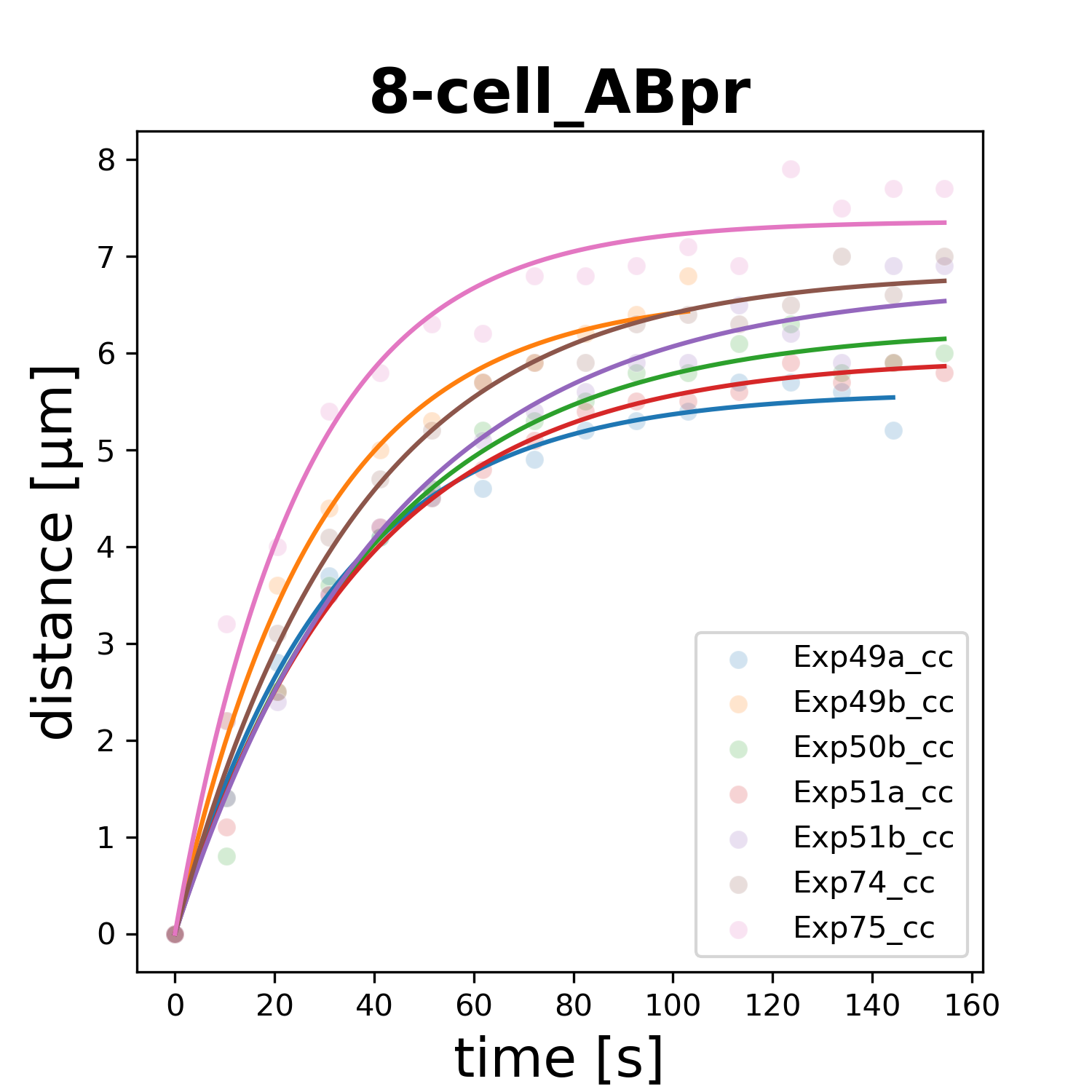

### 8-cell_C.png

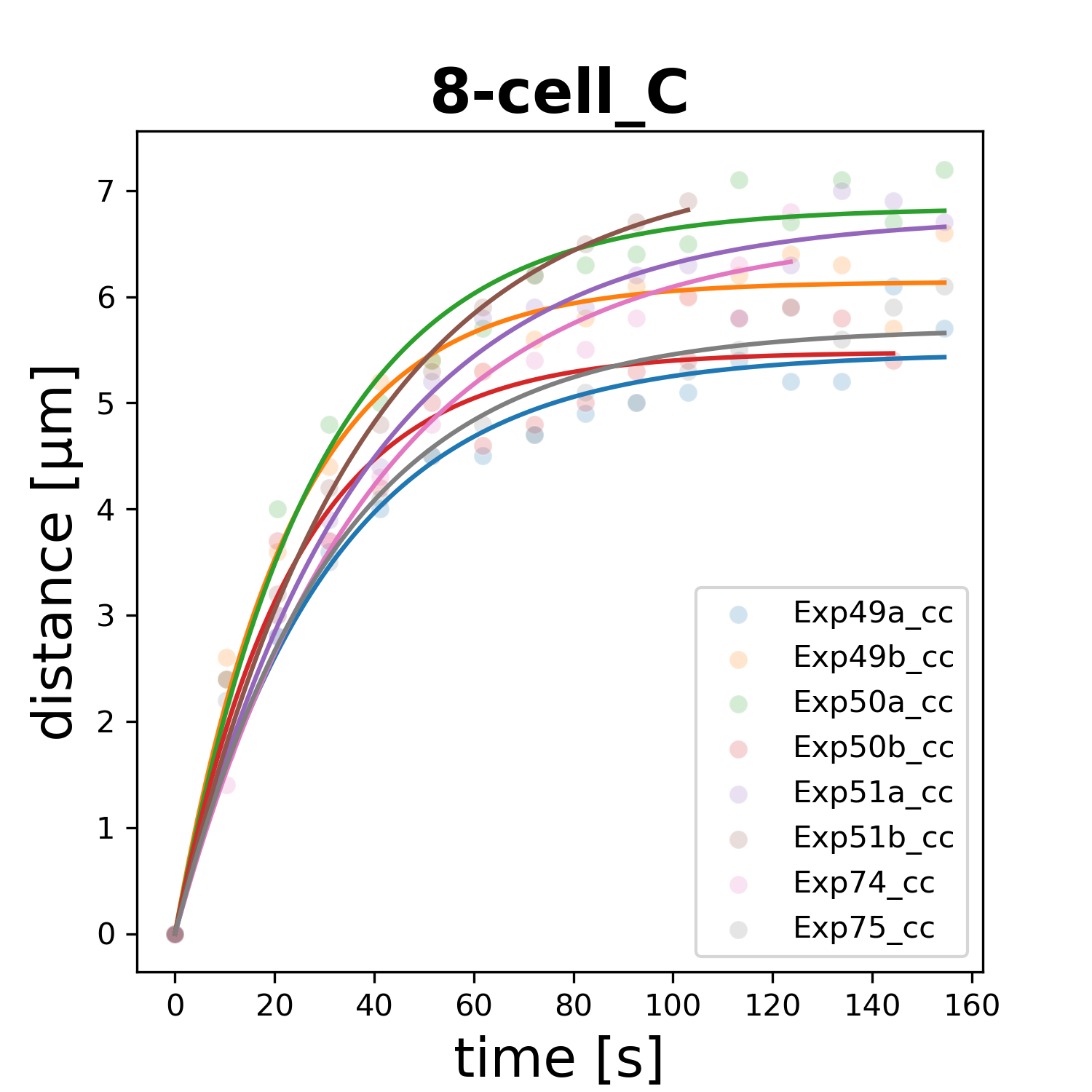

### 8-cell_E.png

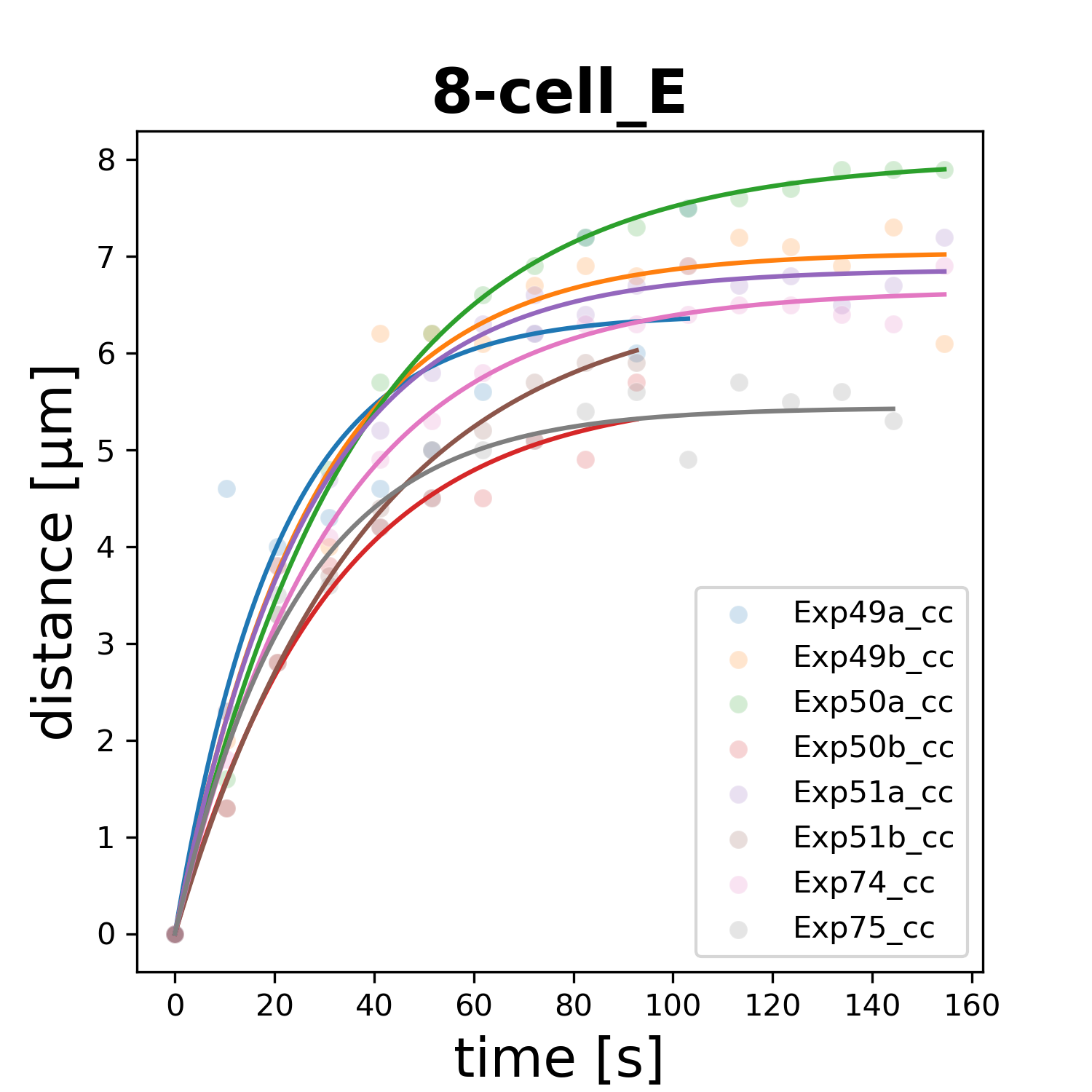

### 8-cell_MS.png

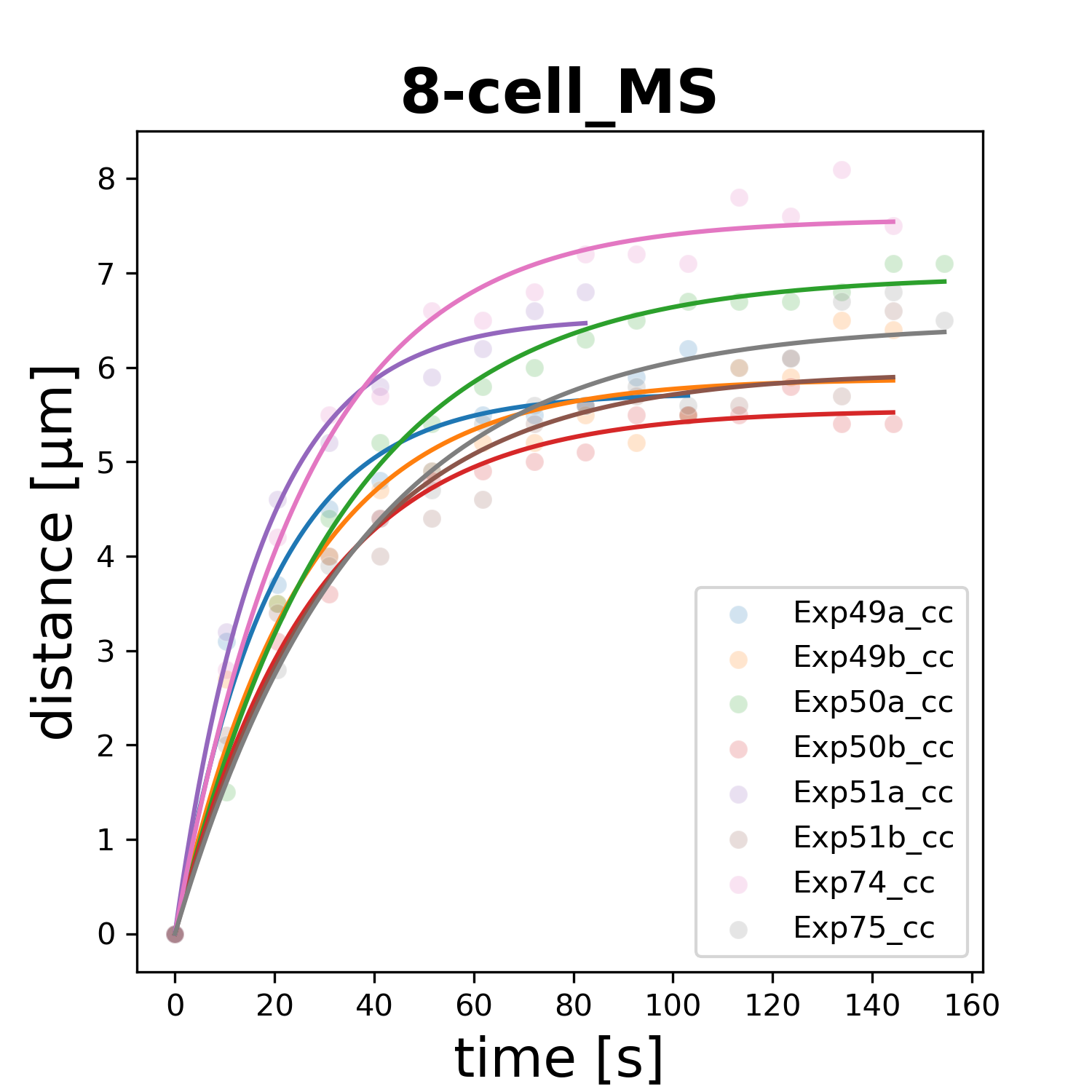

### 8-cell_P3.png

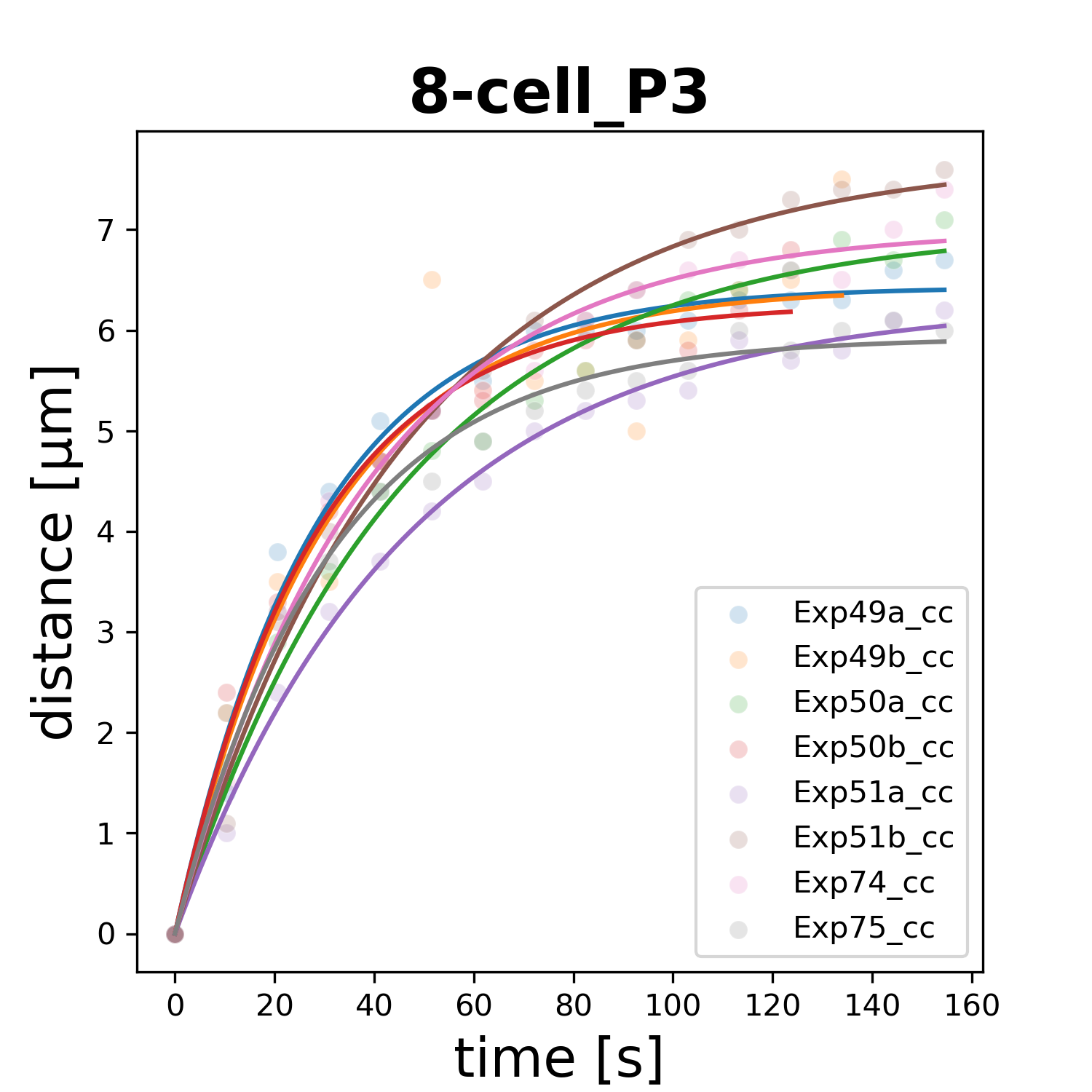

### 16-cell_ABala.png

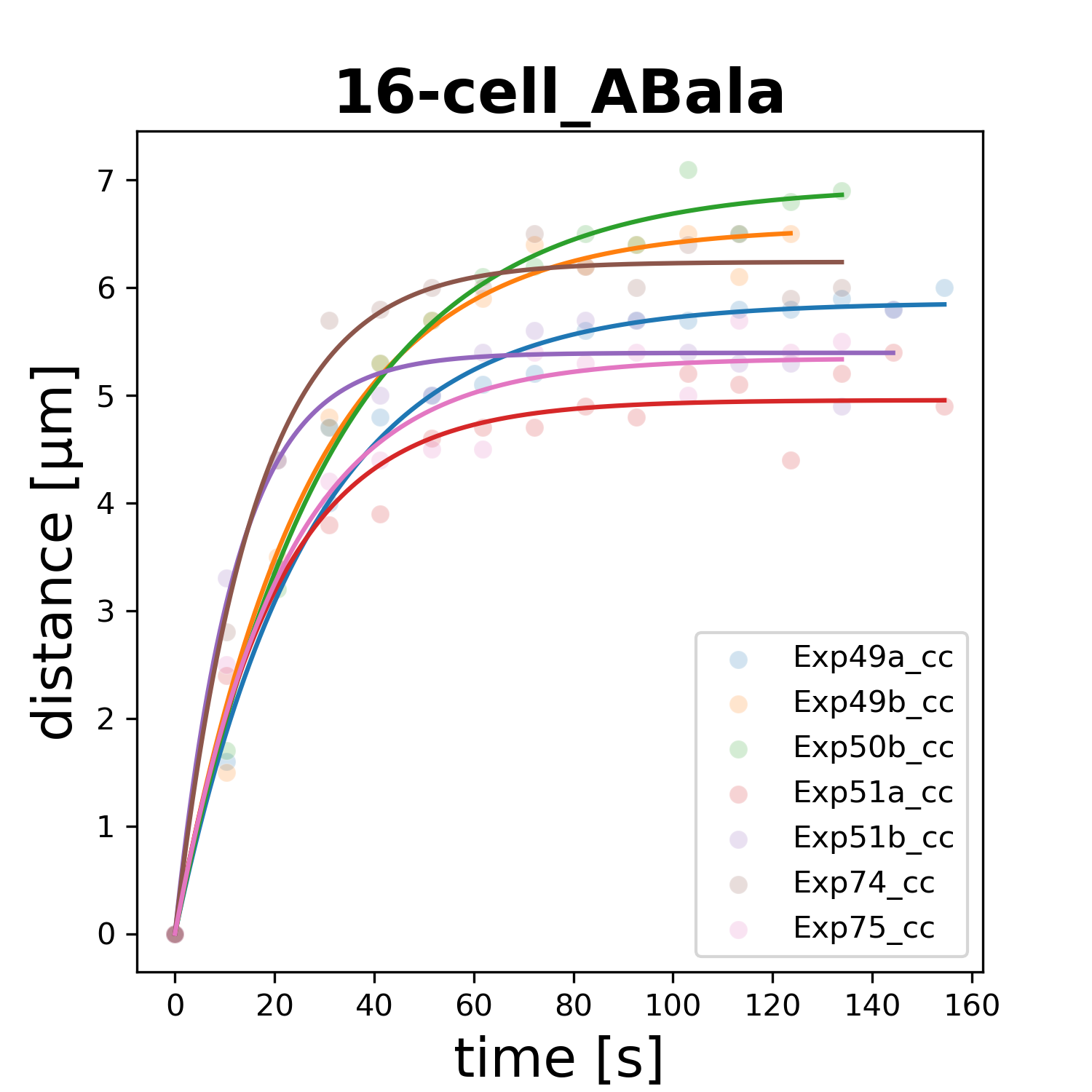

### 16-cell_ABala.png

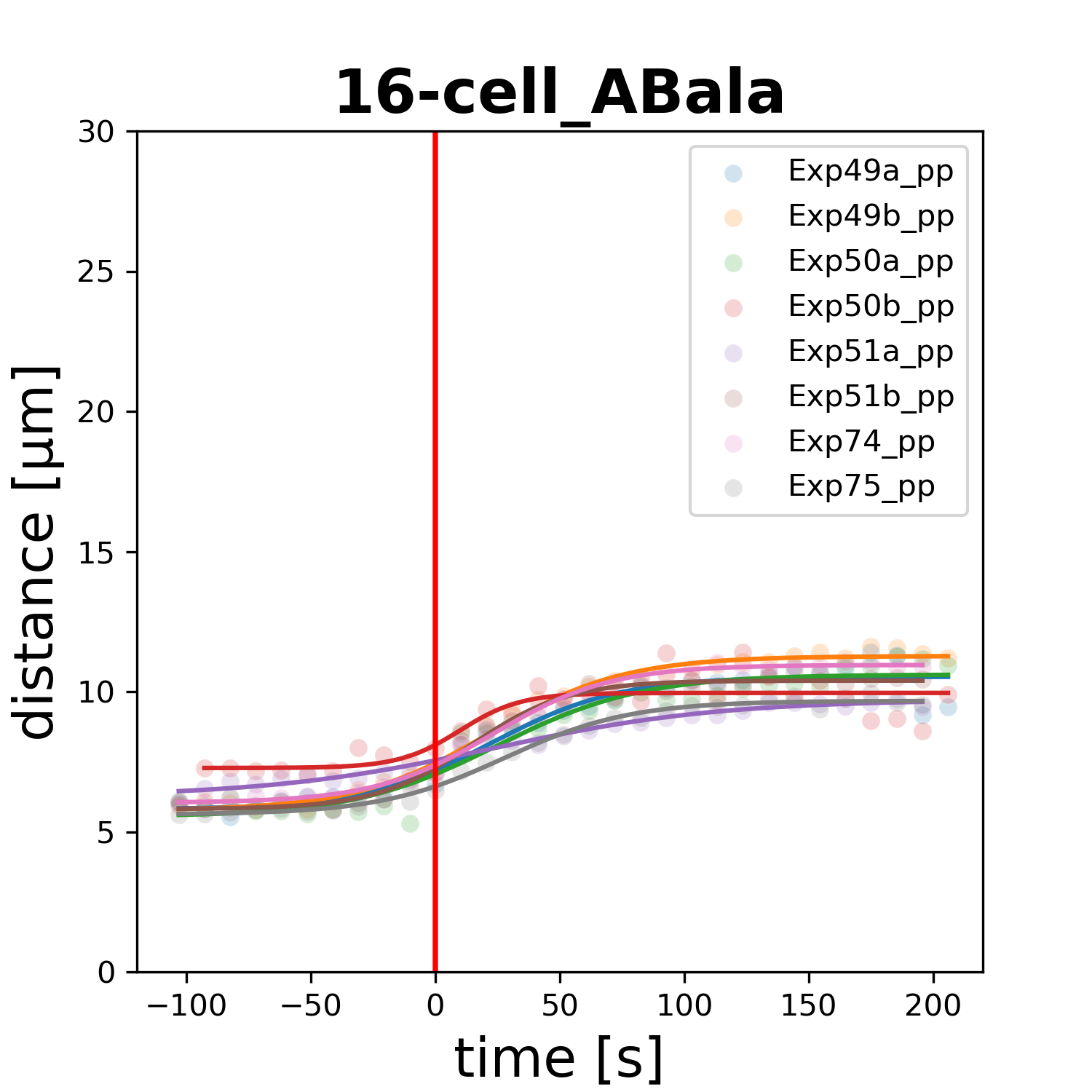

### 16-cell_ABalp.png

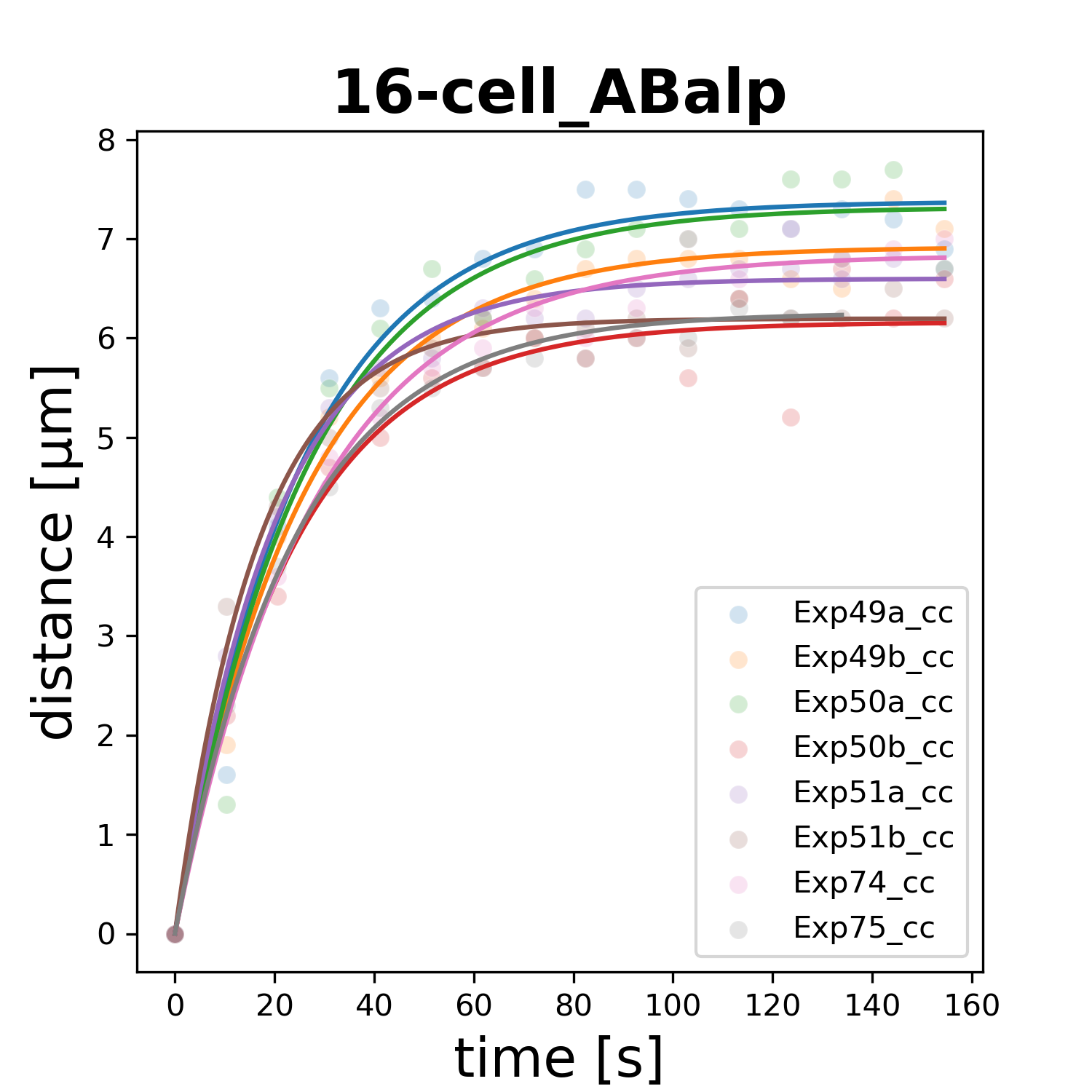

### 16-cell_ABalp.png

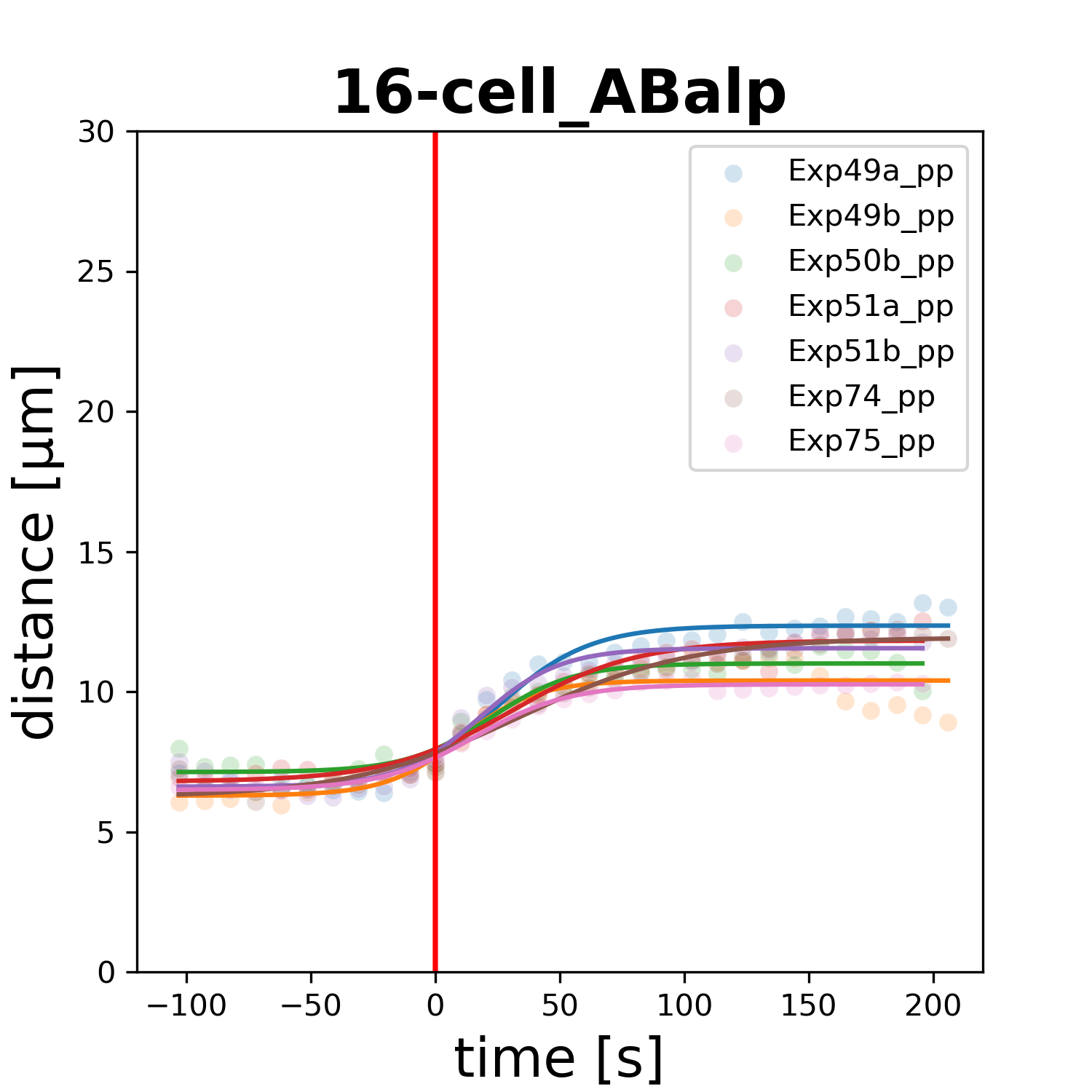

### 16-cell_ABara.png

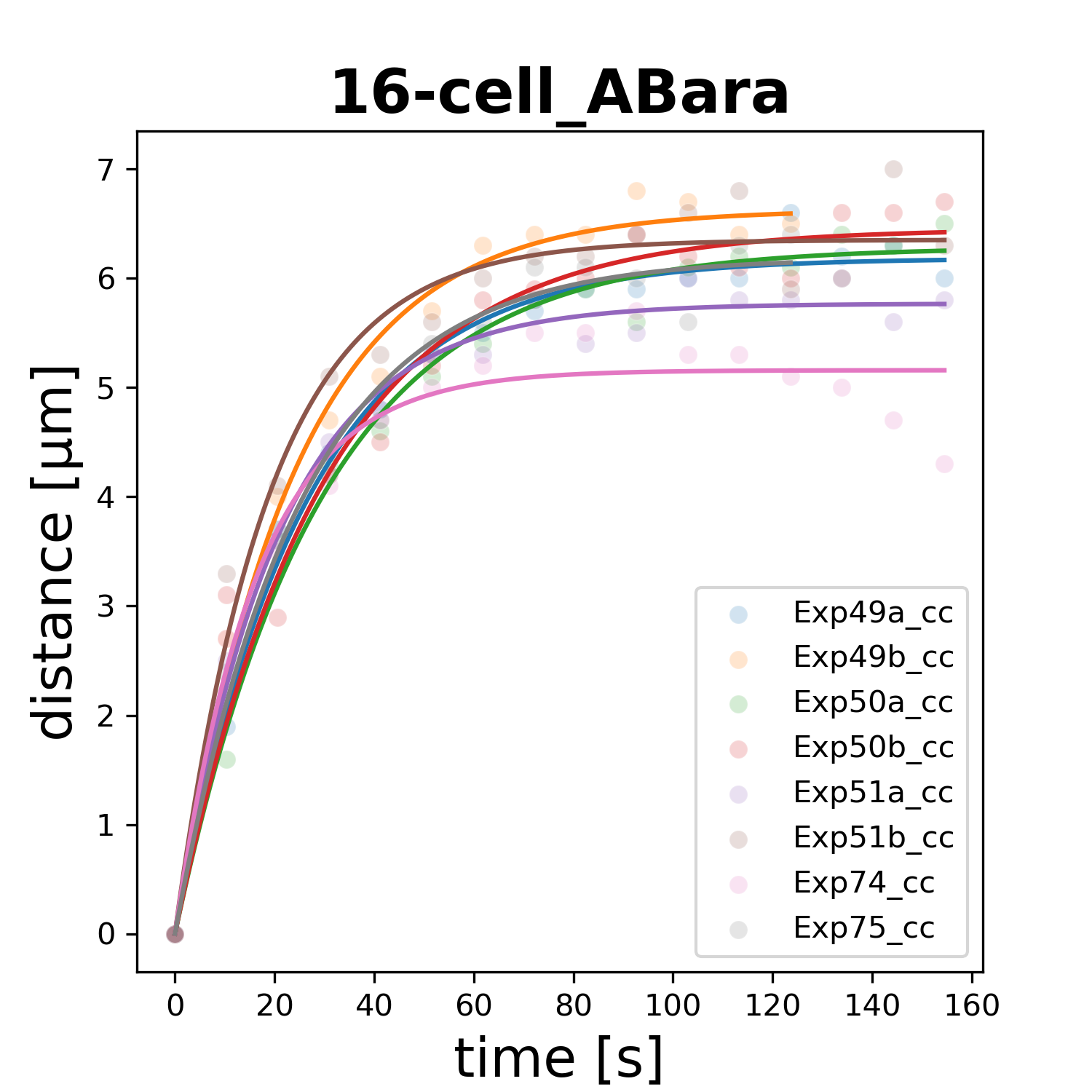

### 16-cell_ABara.png

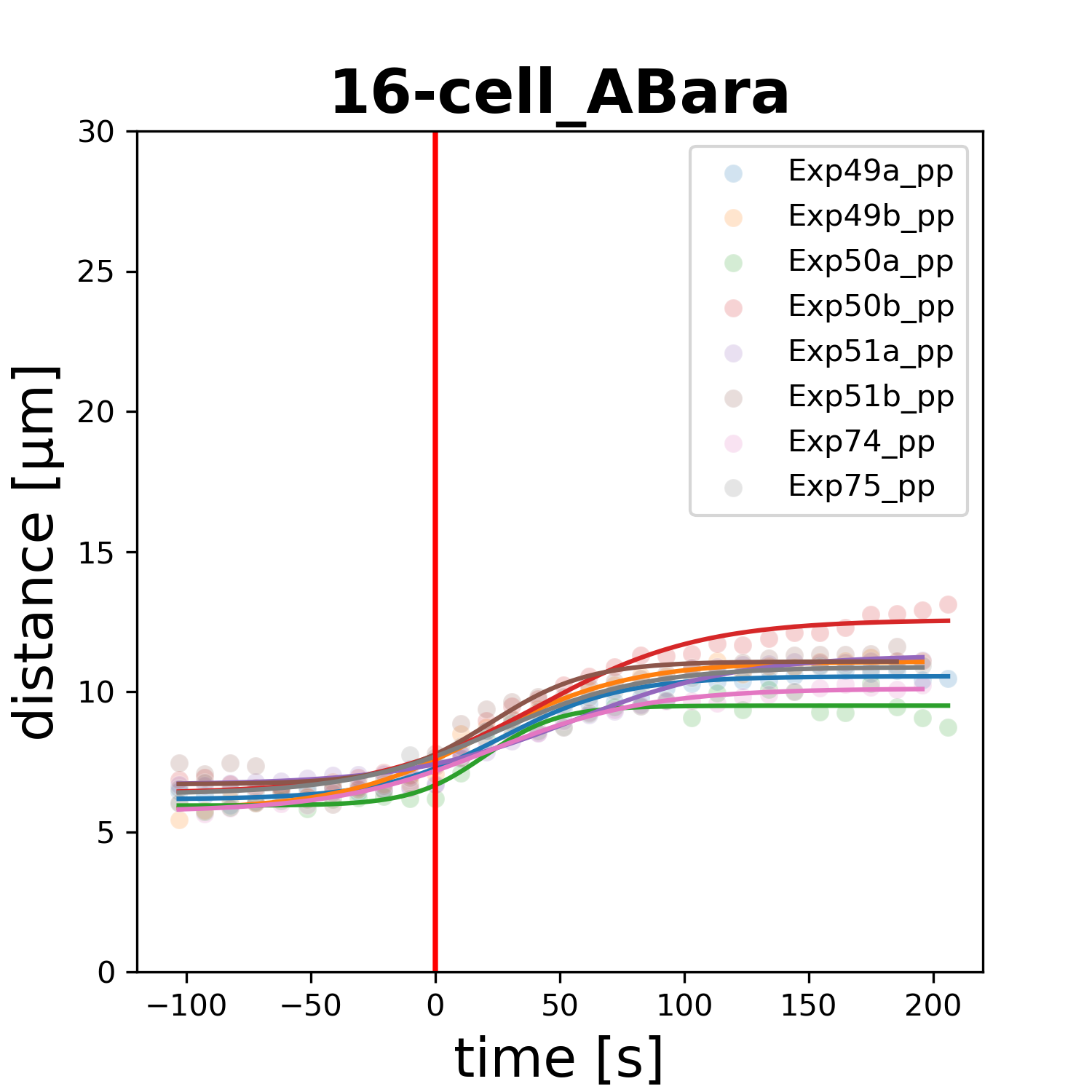

### 16-cell_ABarp.png

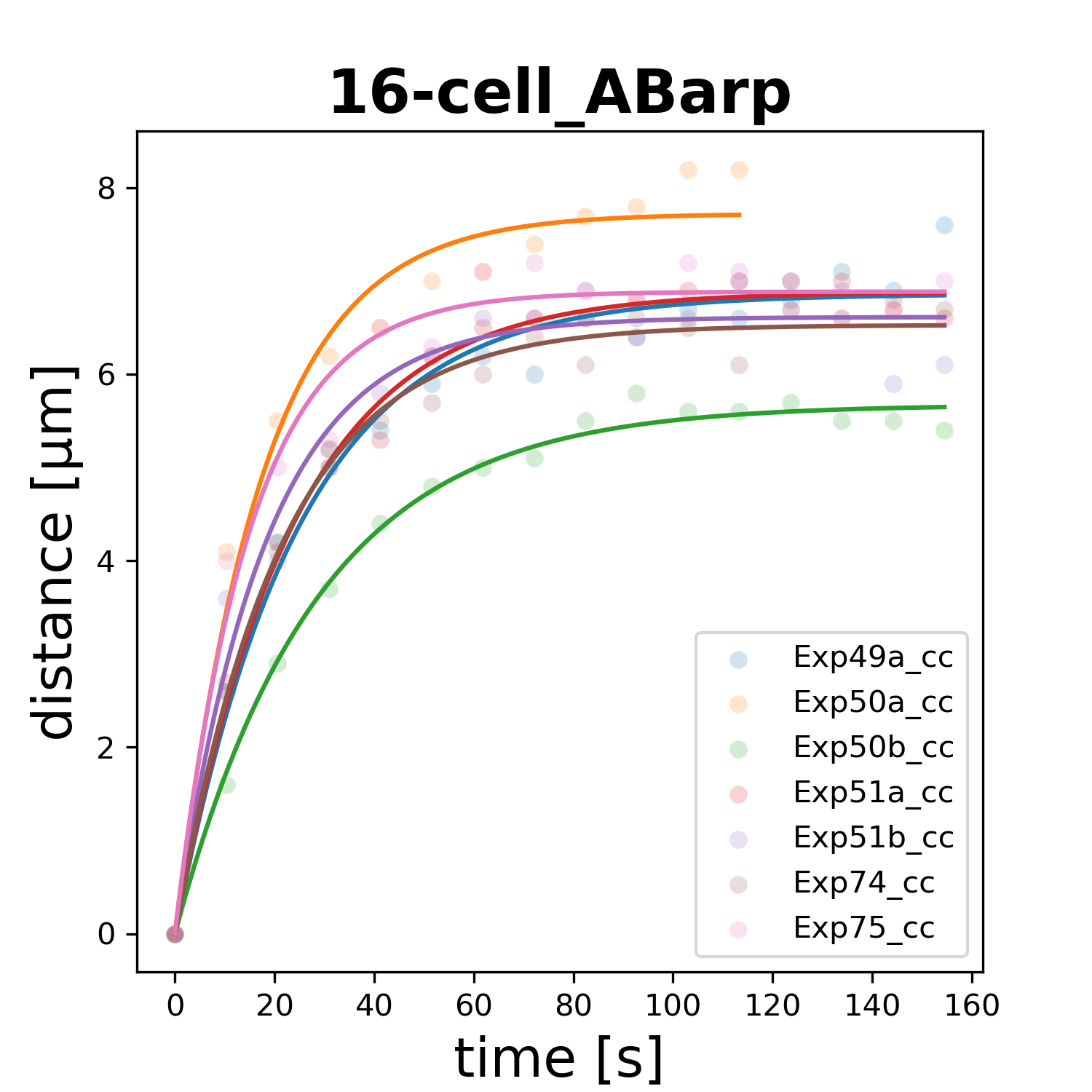

### 16-cell_ABarp.png

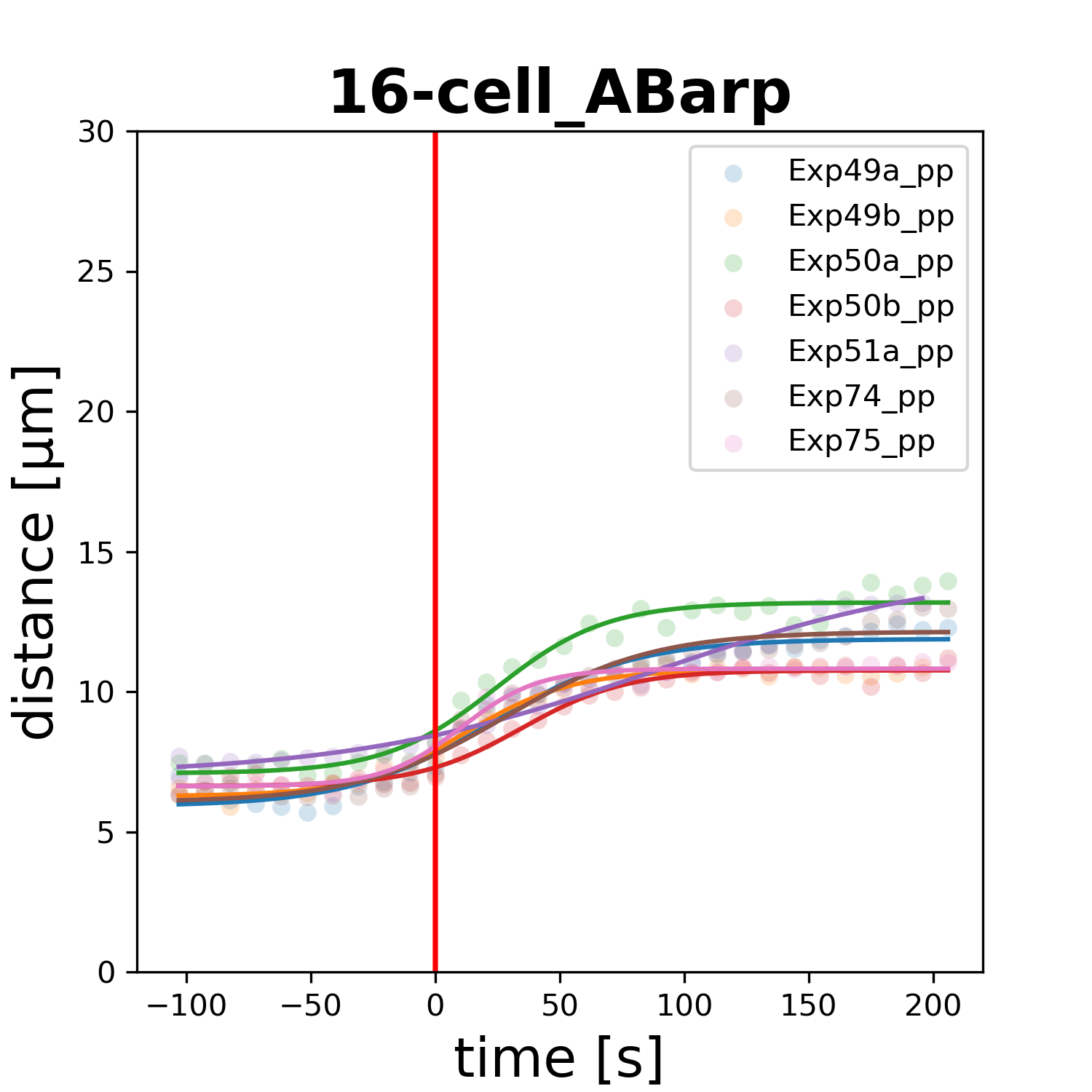

### 16-cell_ABpla.png

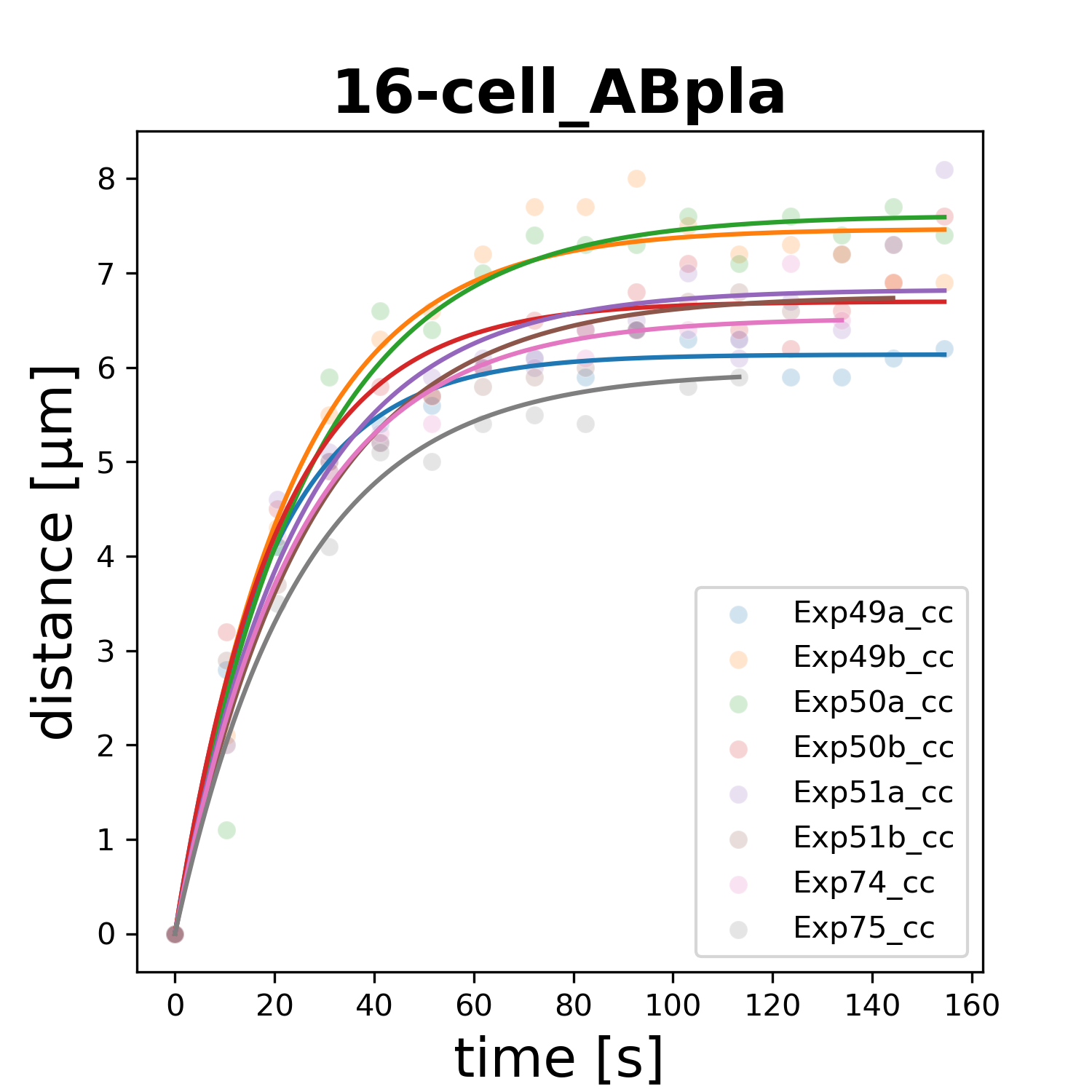

### 16-cell_ABpla.png

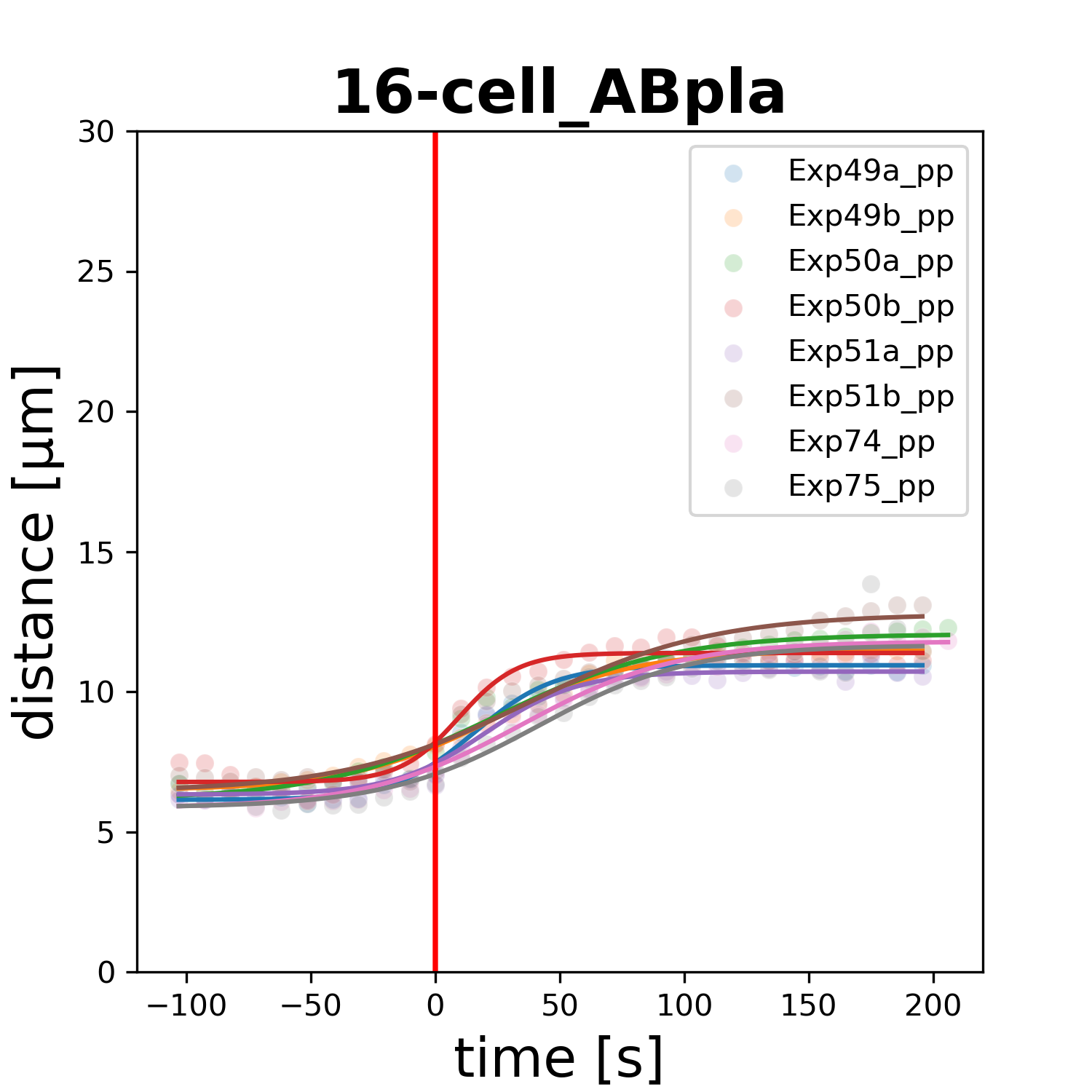

### 16-cell_ABplp.png

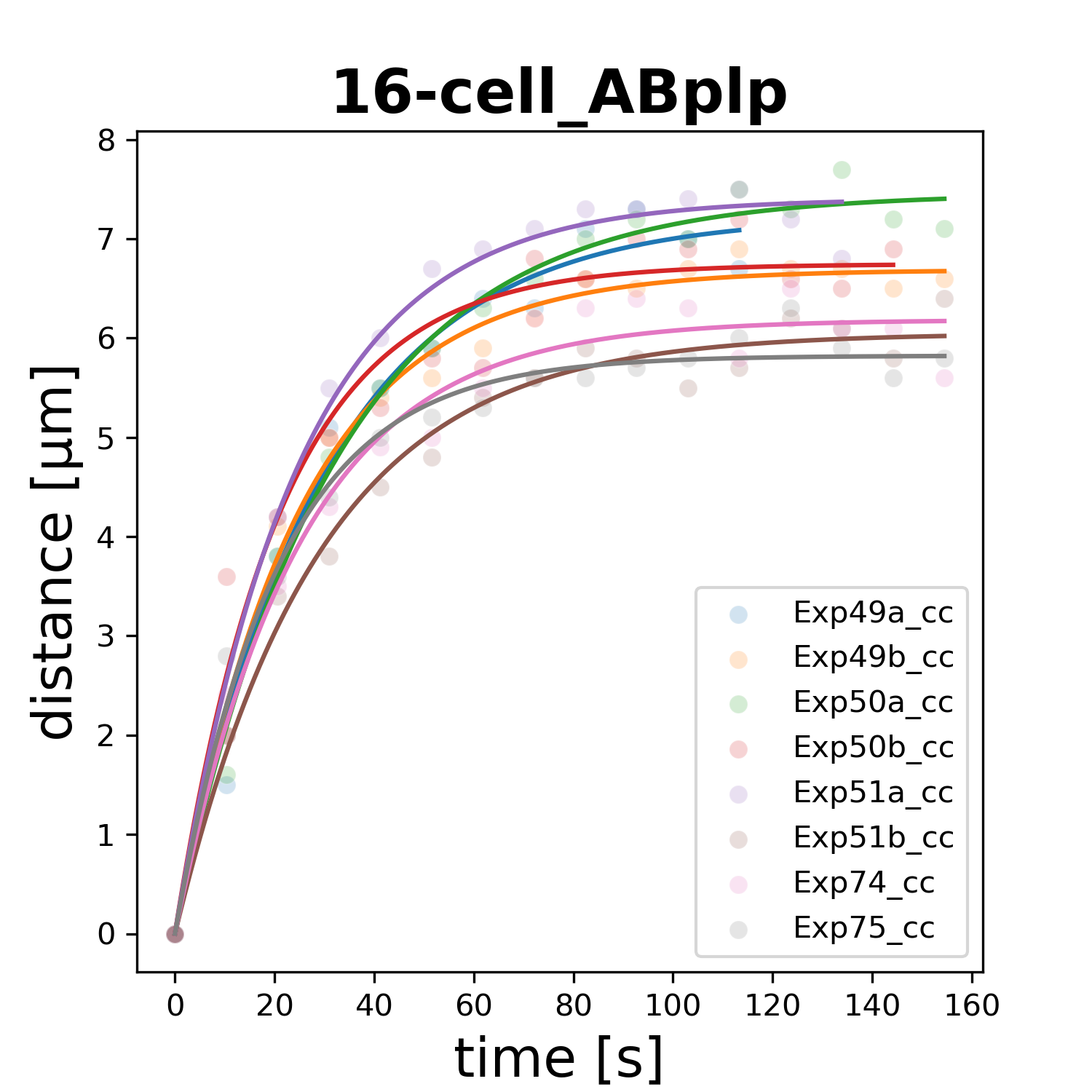

### 16-cell_ABpra.png

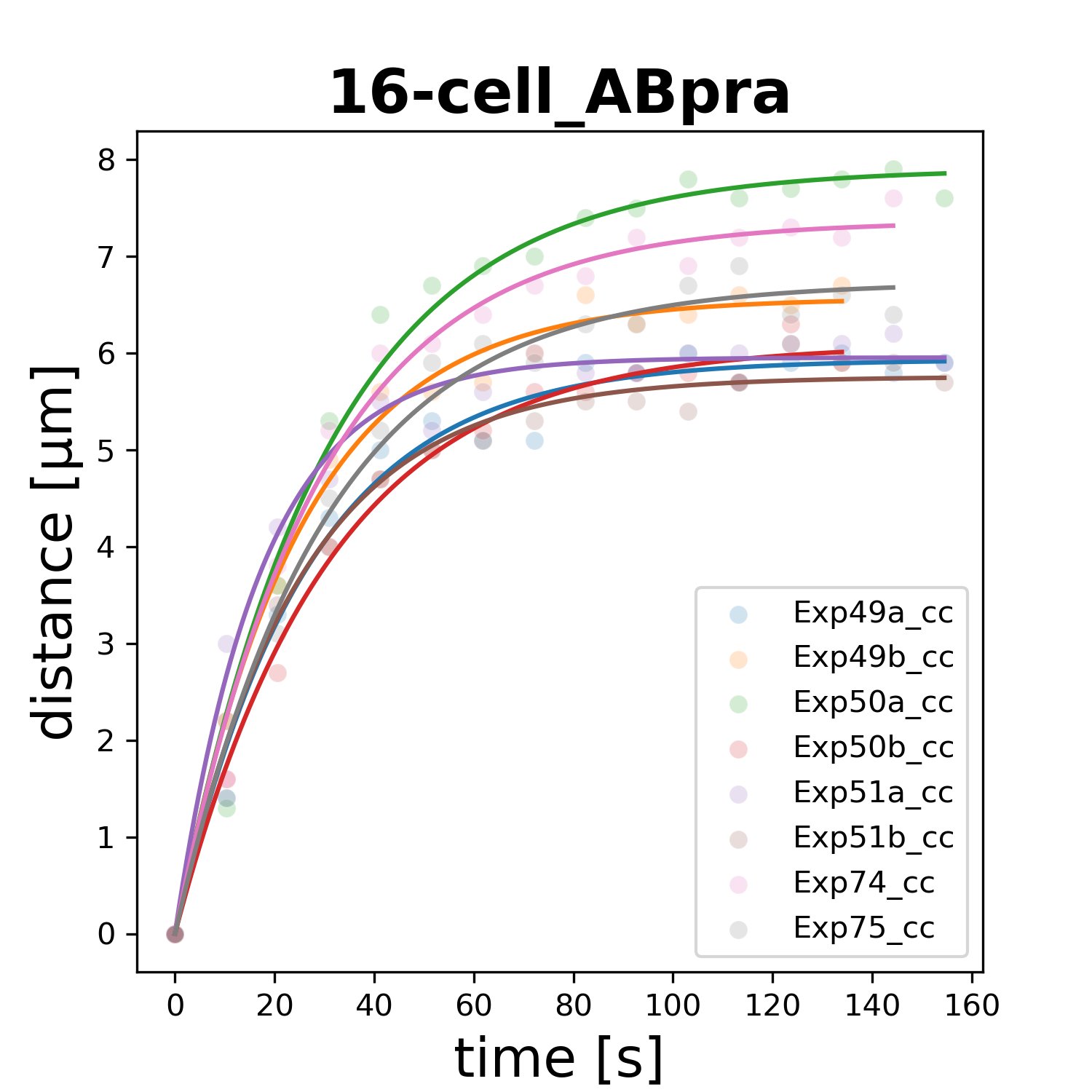

### 16-cell_ABprp.png

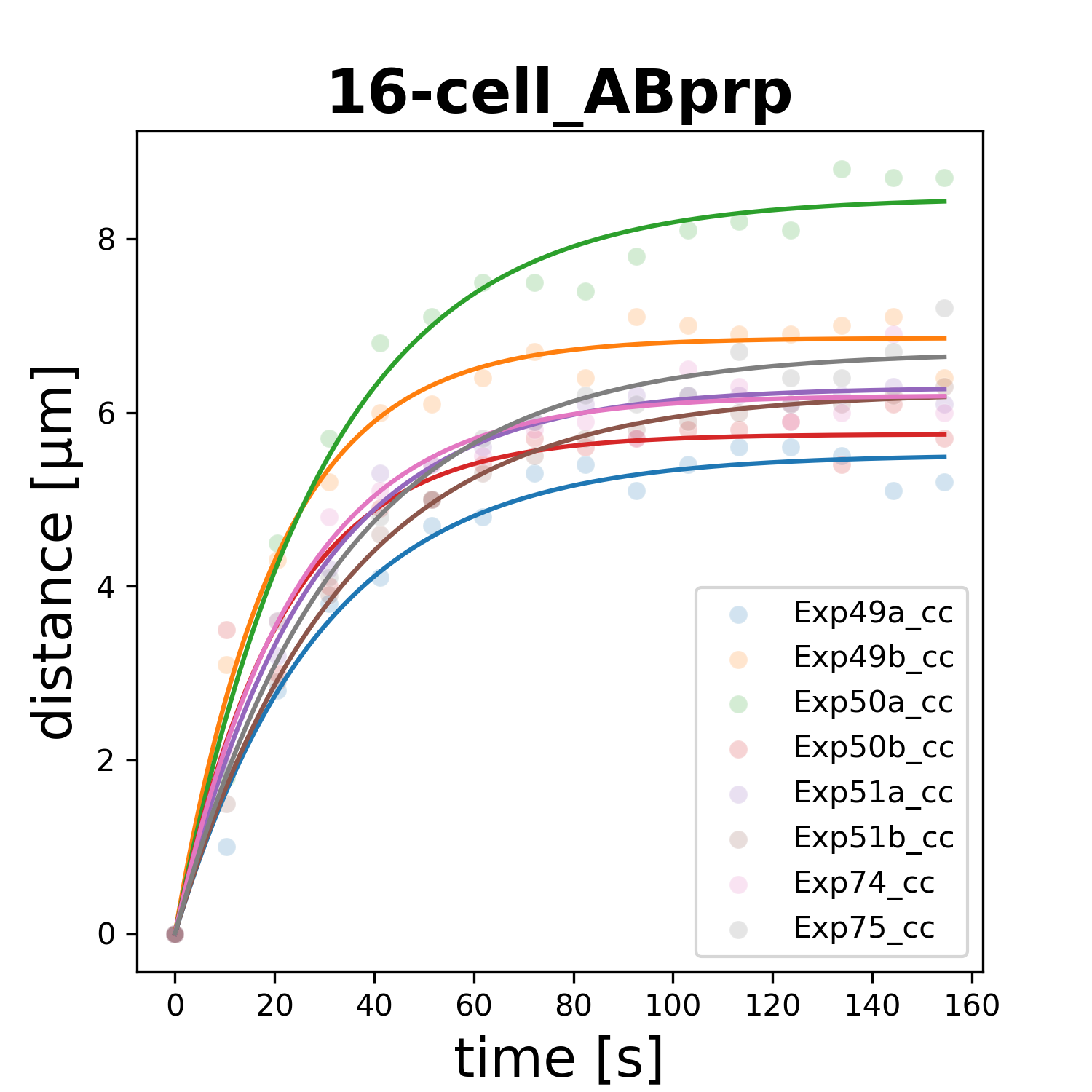

### 16-cell_Ca.png

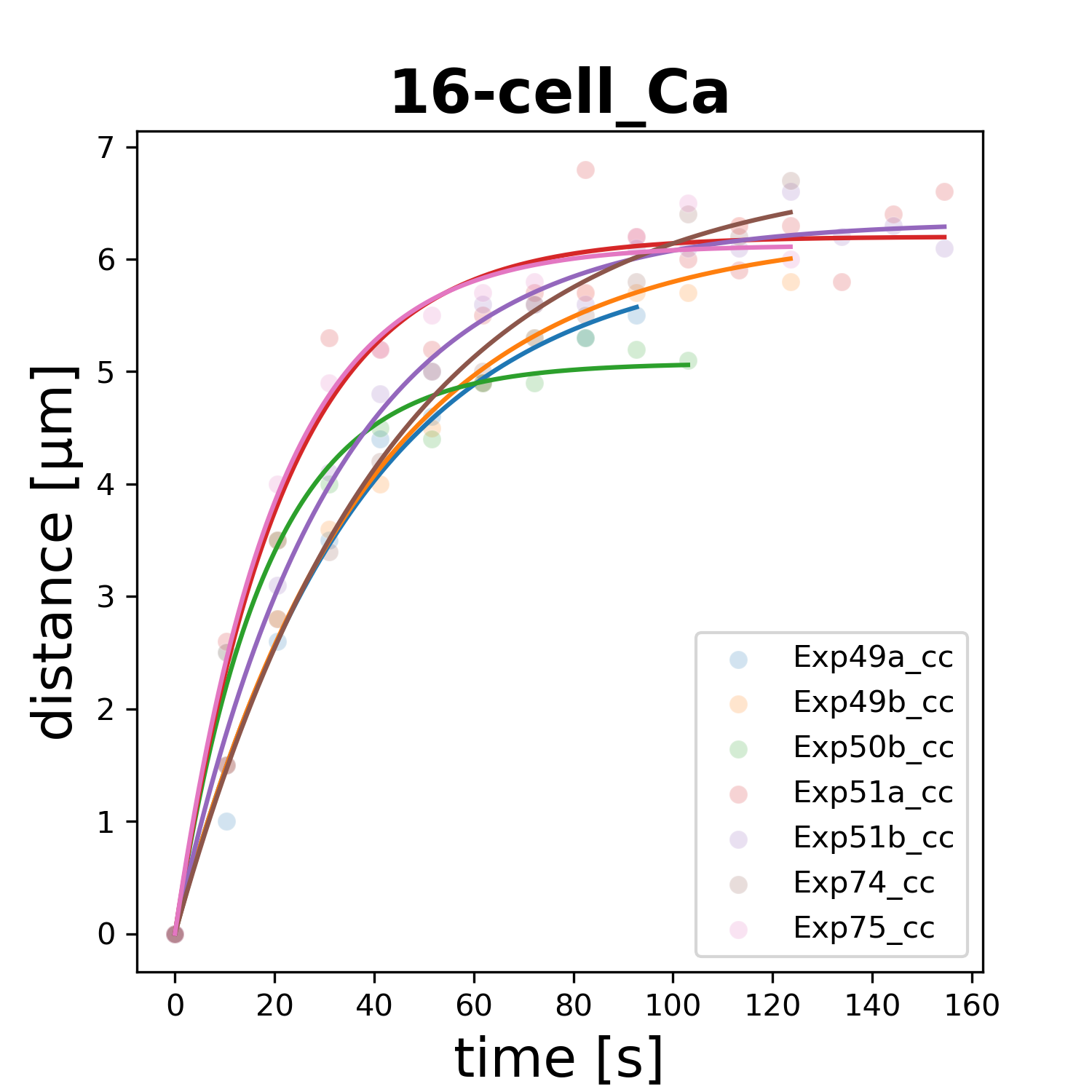

### 16-cell_Cp.png

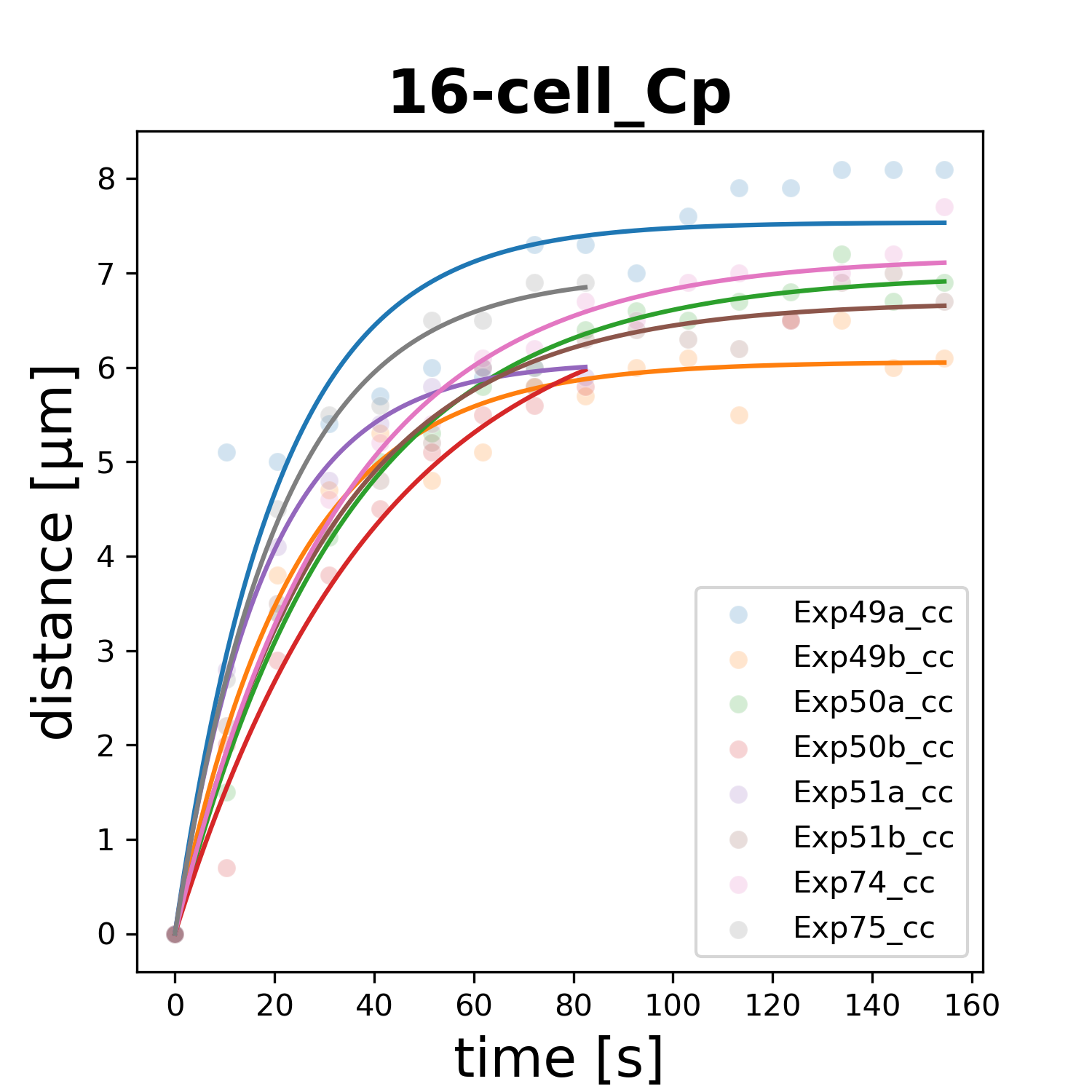

### 16-cell_D.png

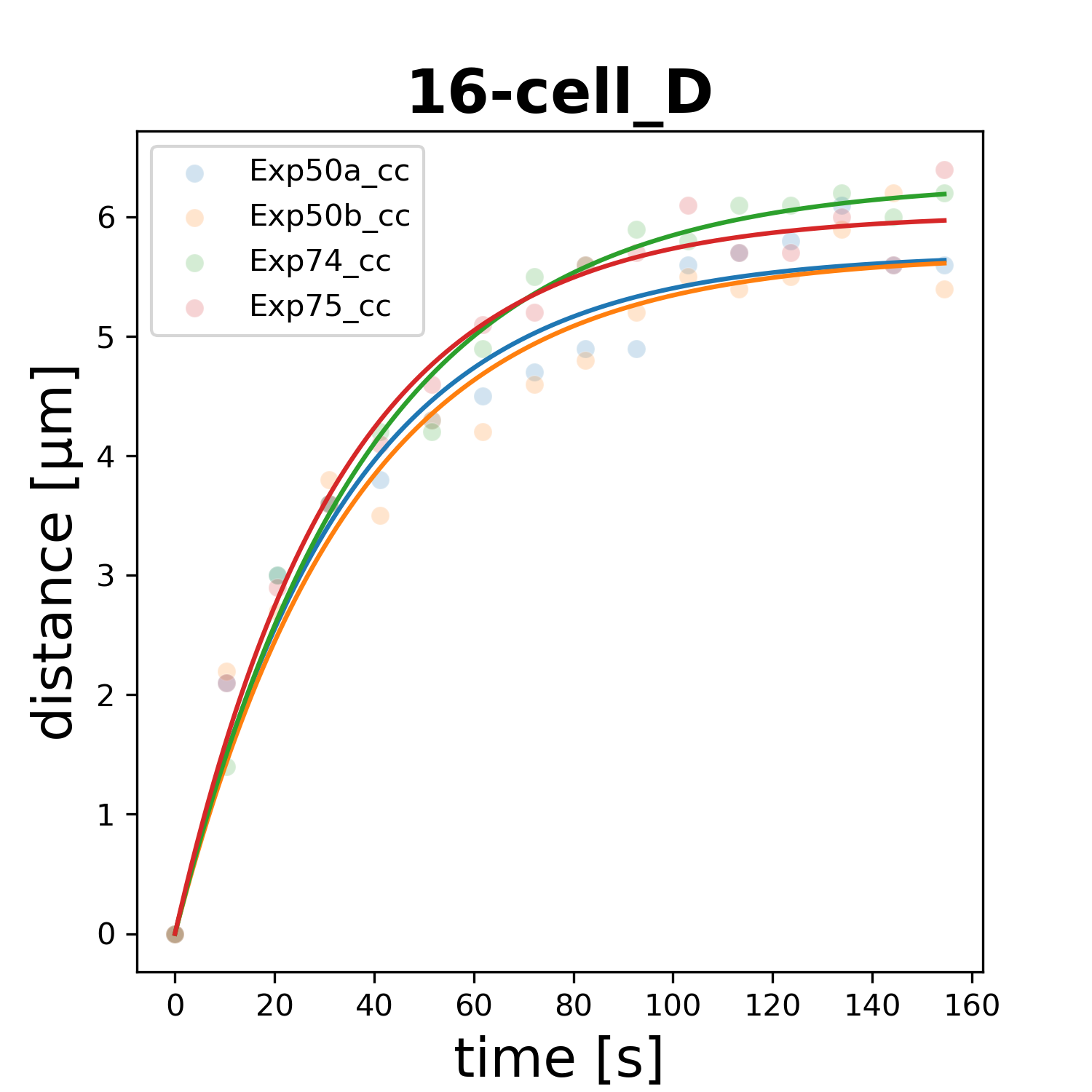

### 16-cell_Ea.png

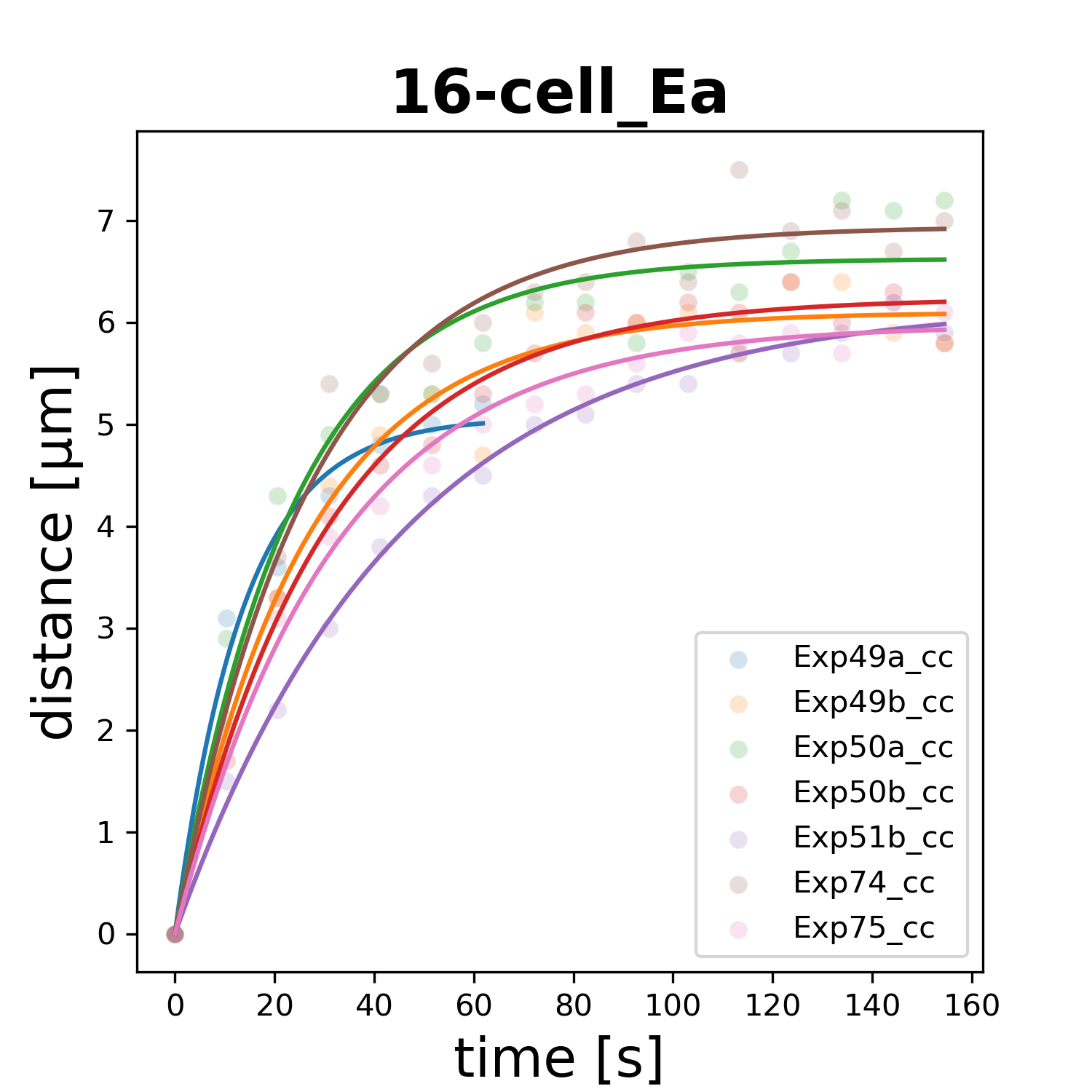

### 16-cell_Ep.png

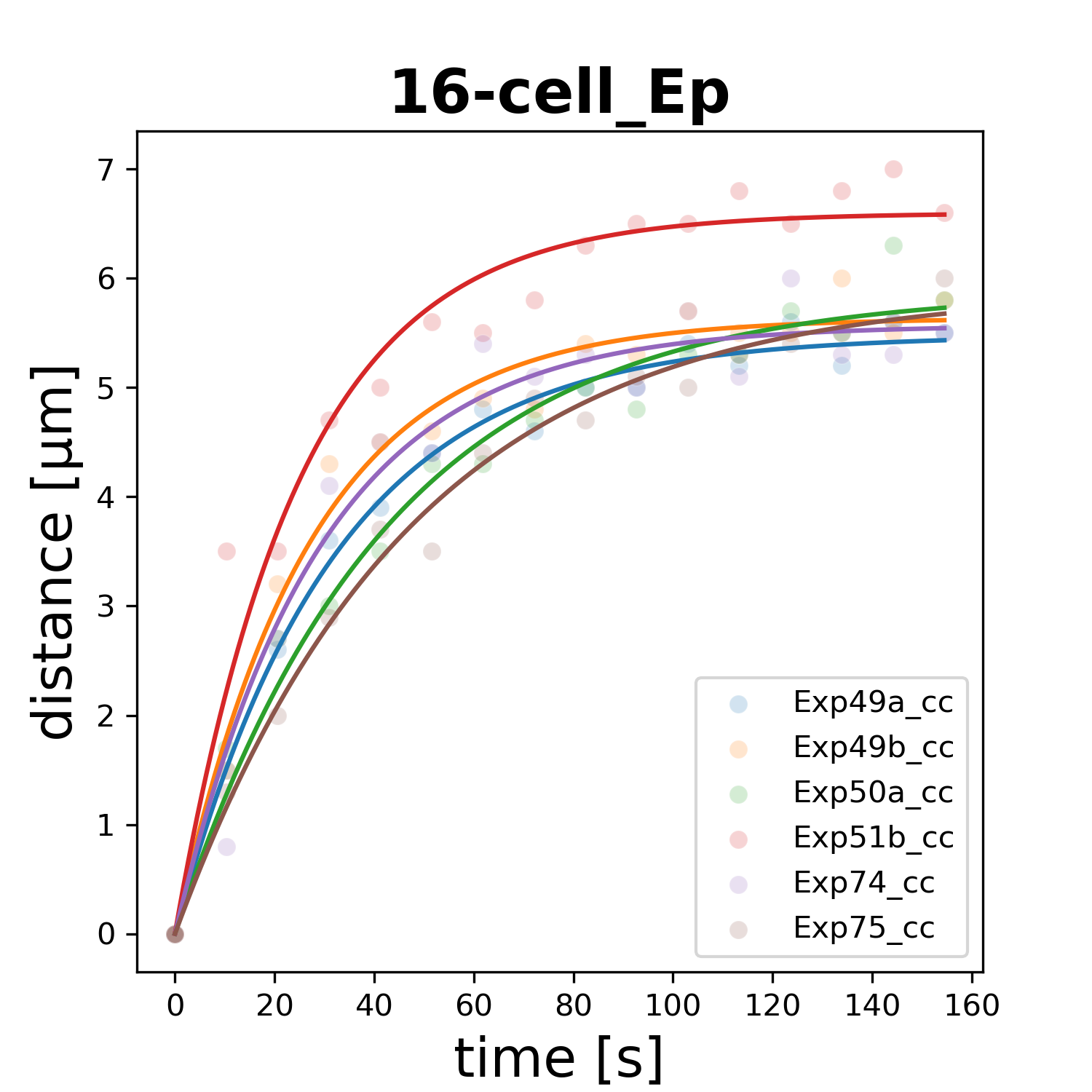

### 16-cell_MSa.png

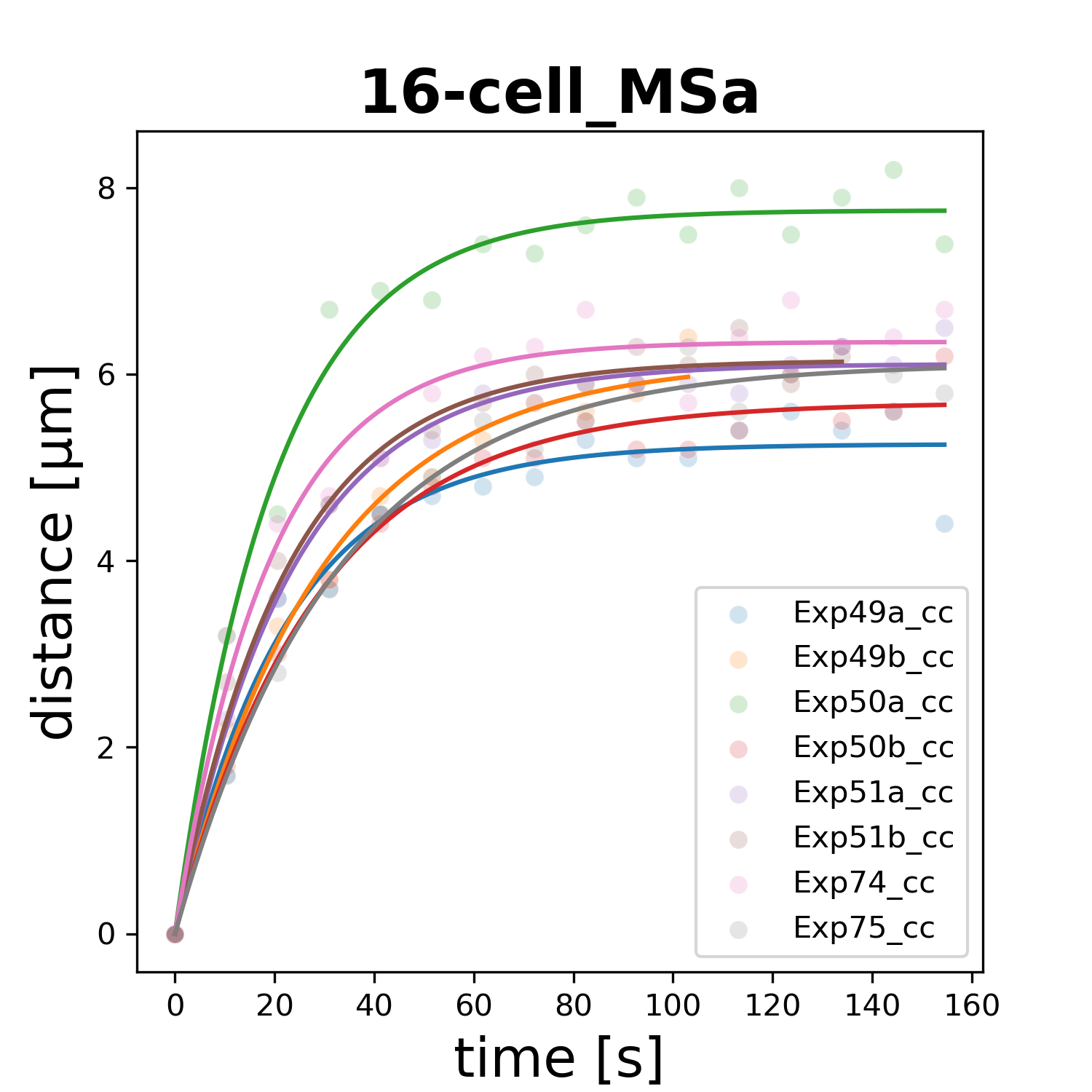

### 16-cell_MSp.png

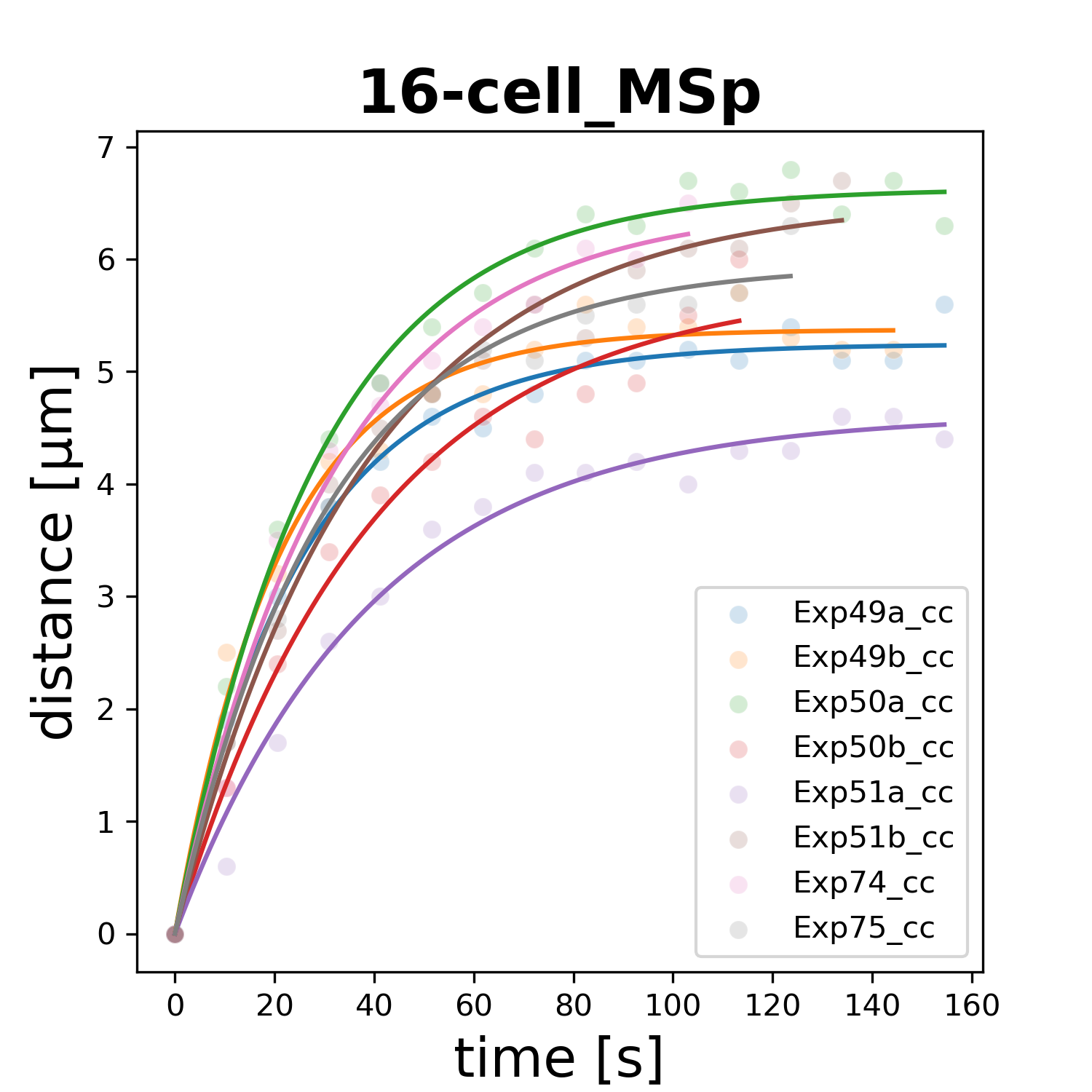

### 16-cell_P4.png

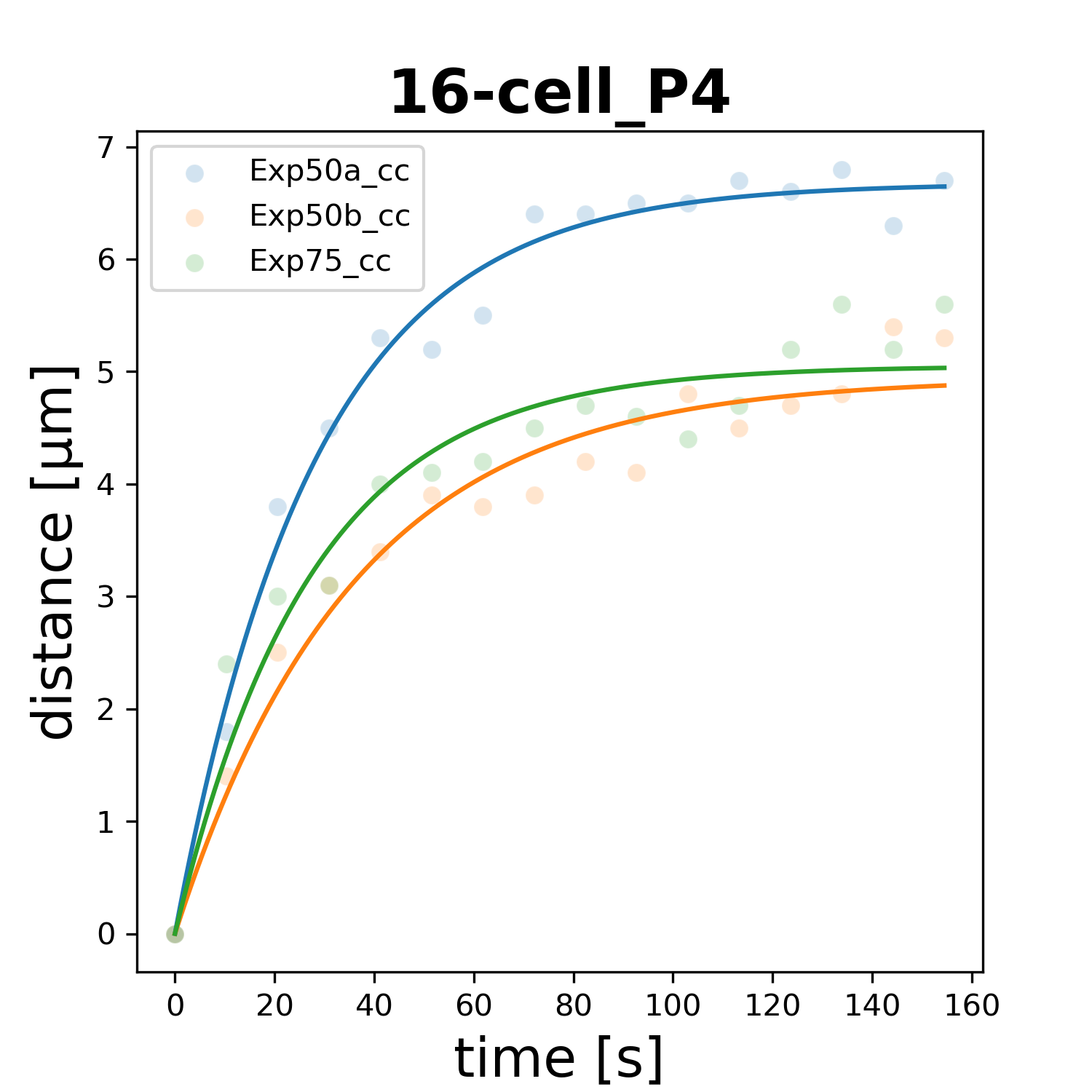

### 32-cell_ABalaa.png

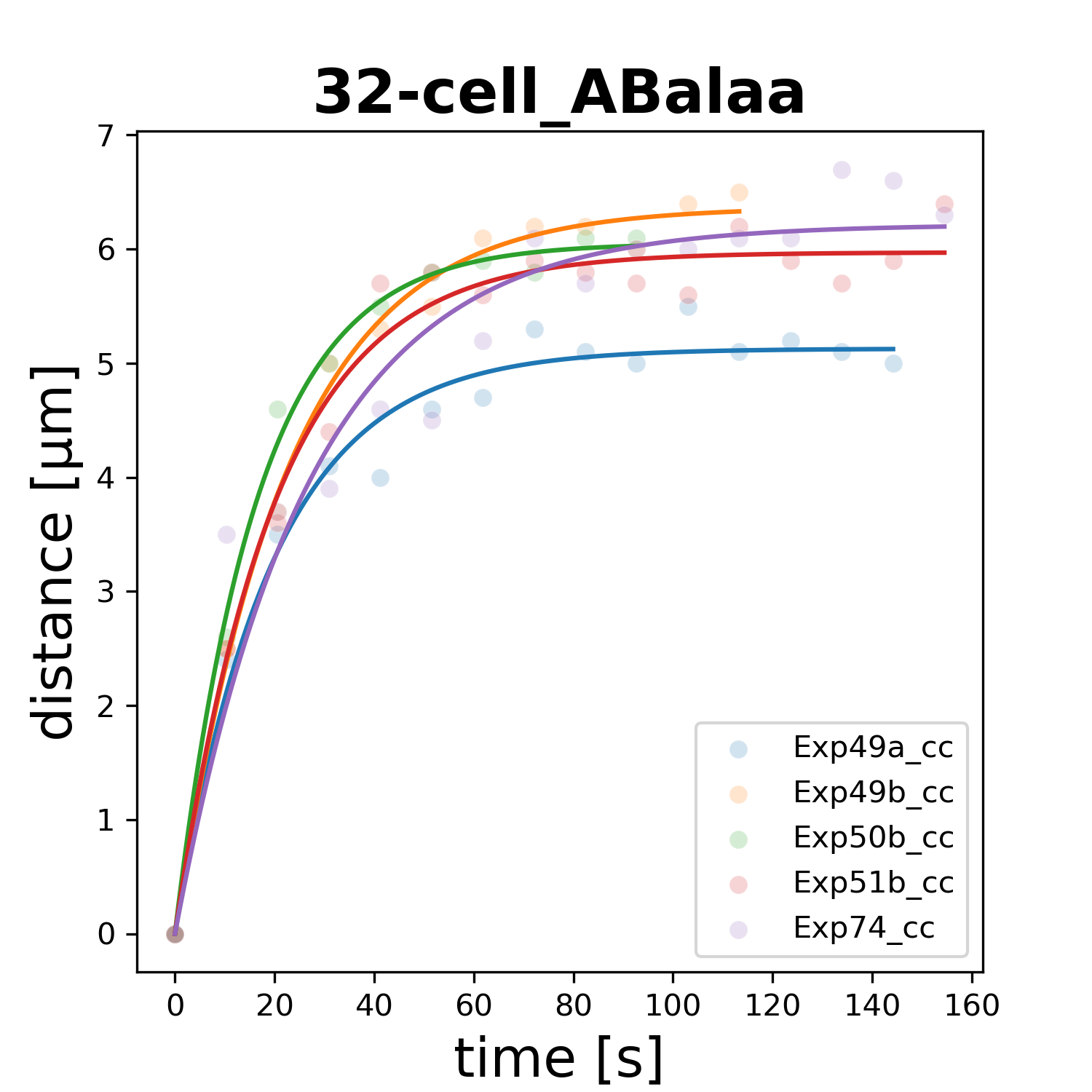

### 32-cell_ABalap.png

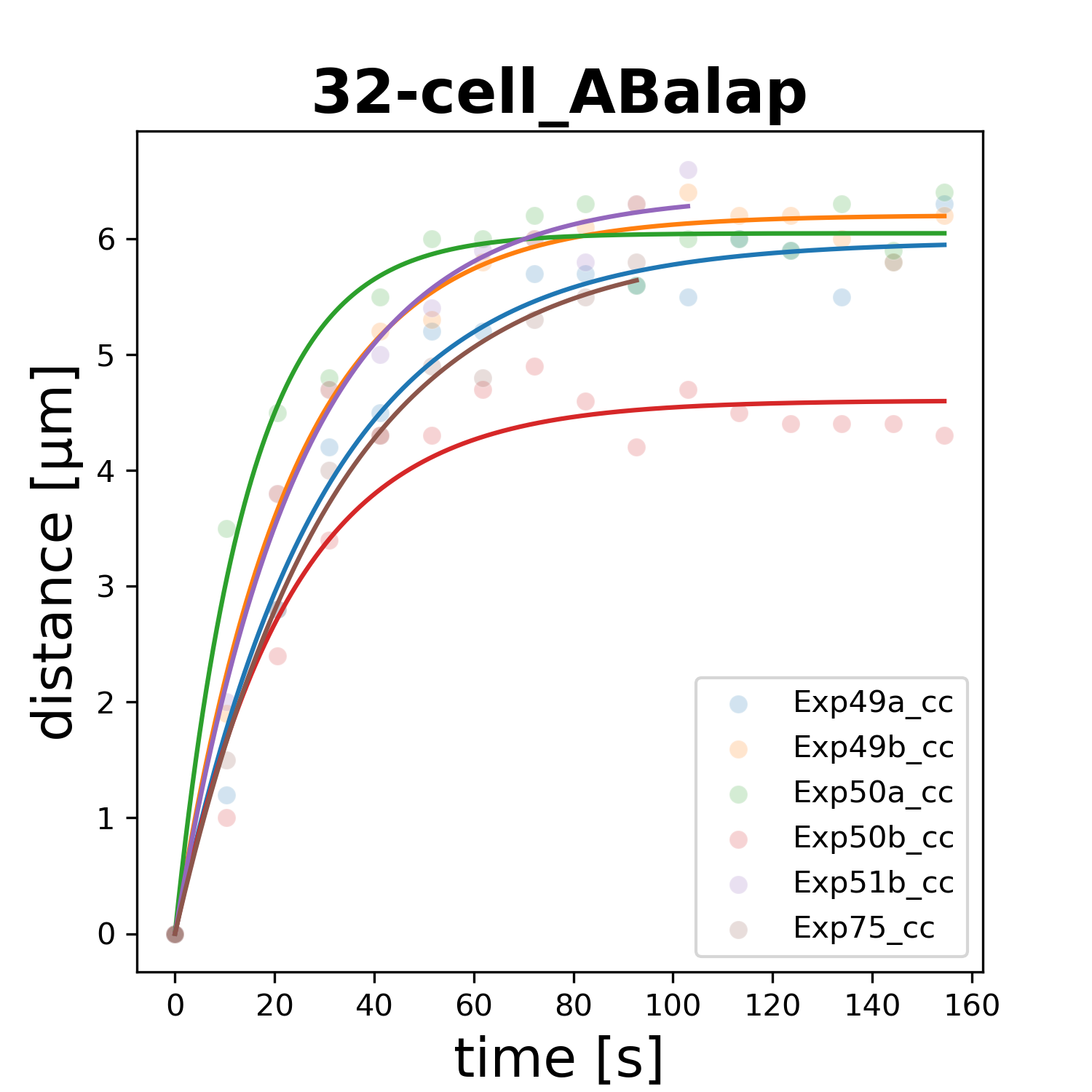

### 32-cell_ABalpa.png

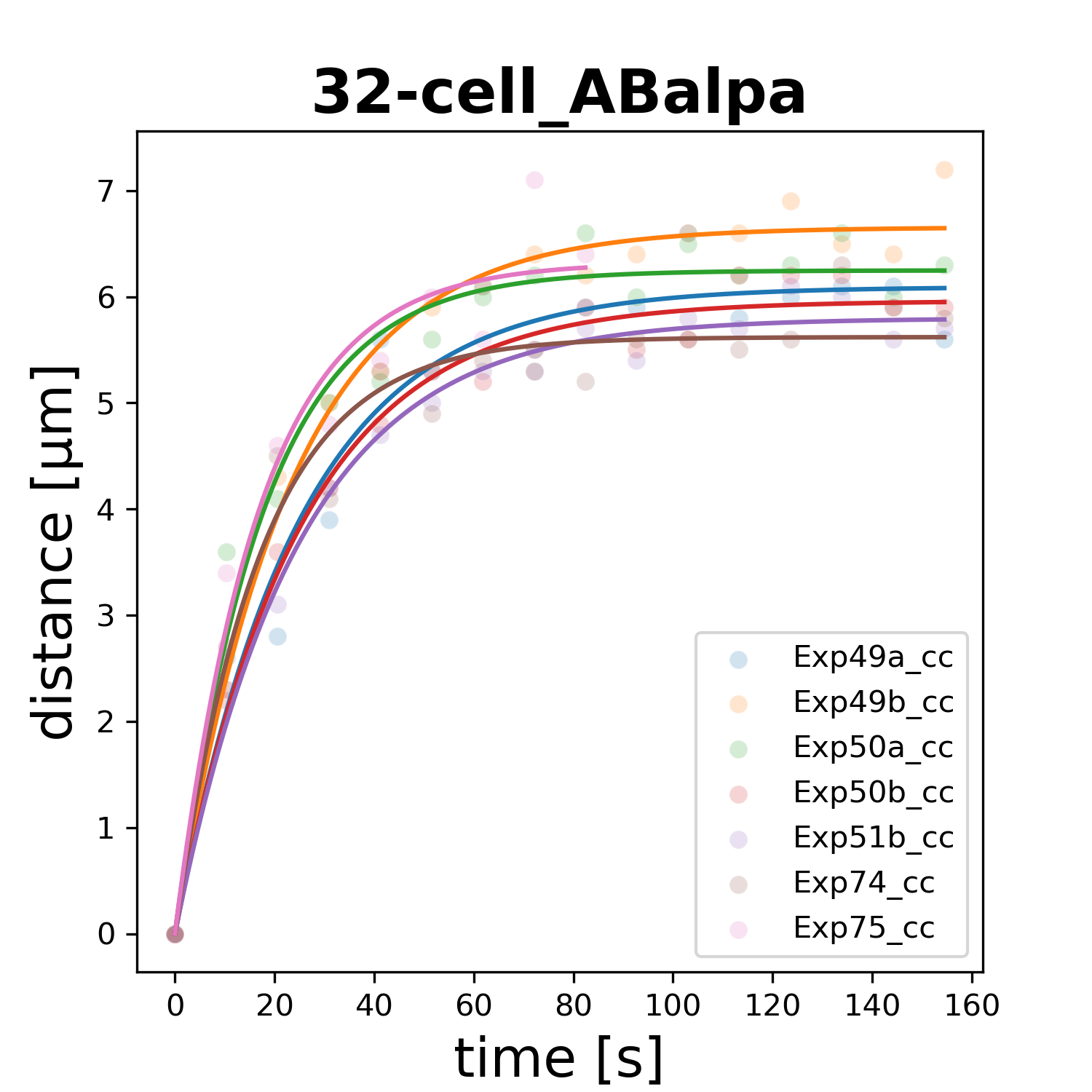

### 32-cell_ABalpp.png

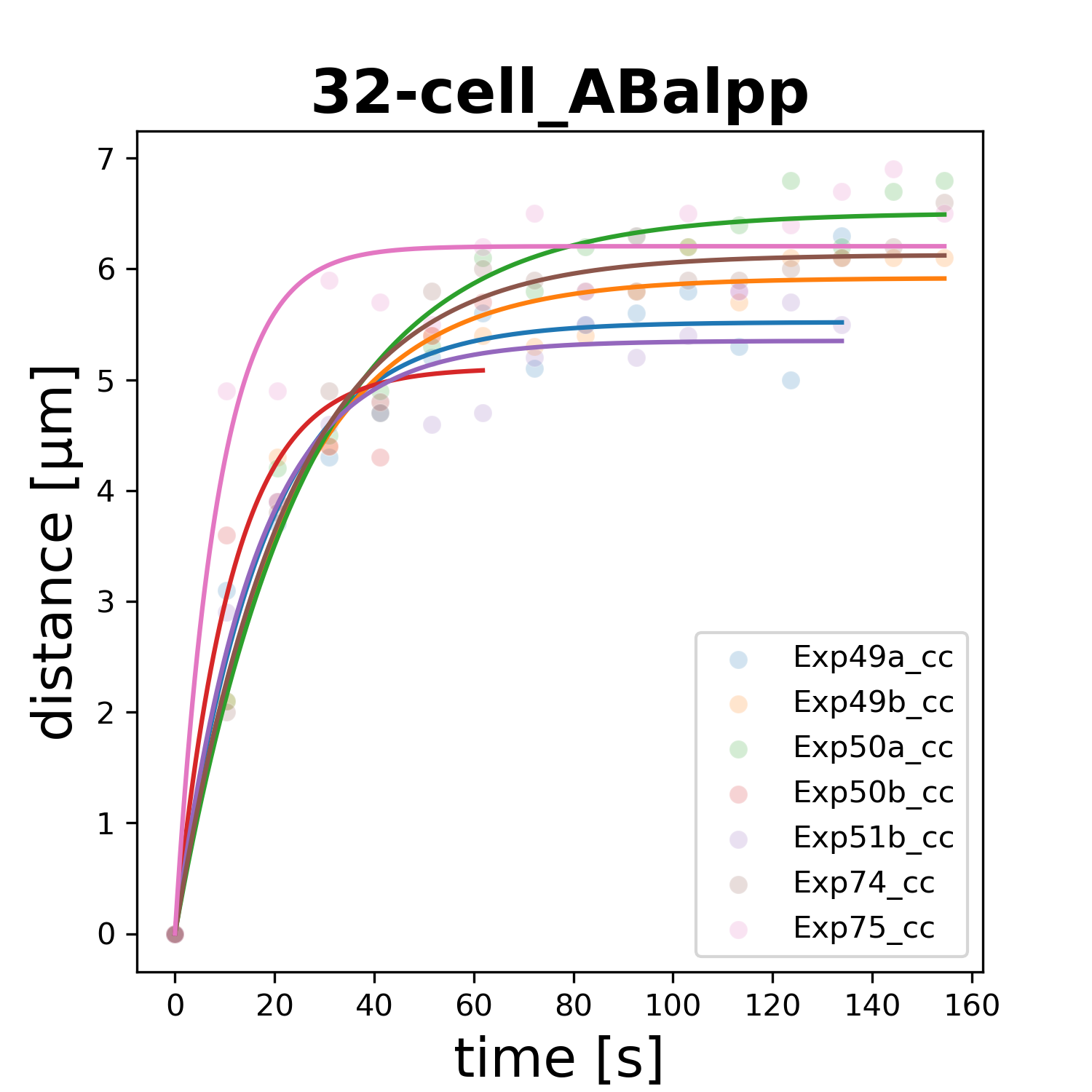

### 32-cell_ABaraa.png

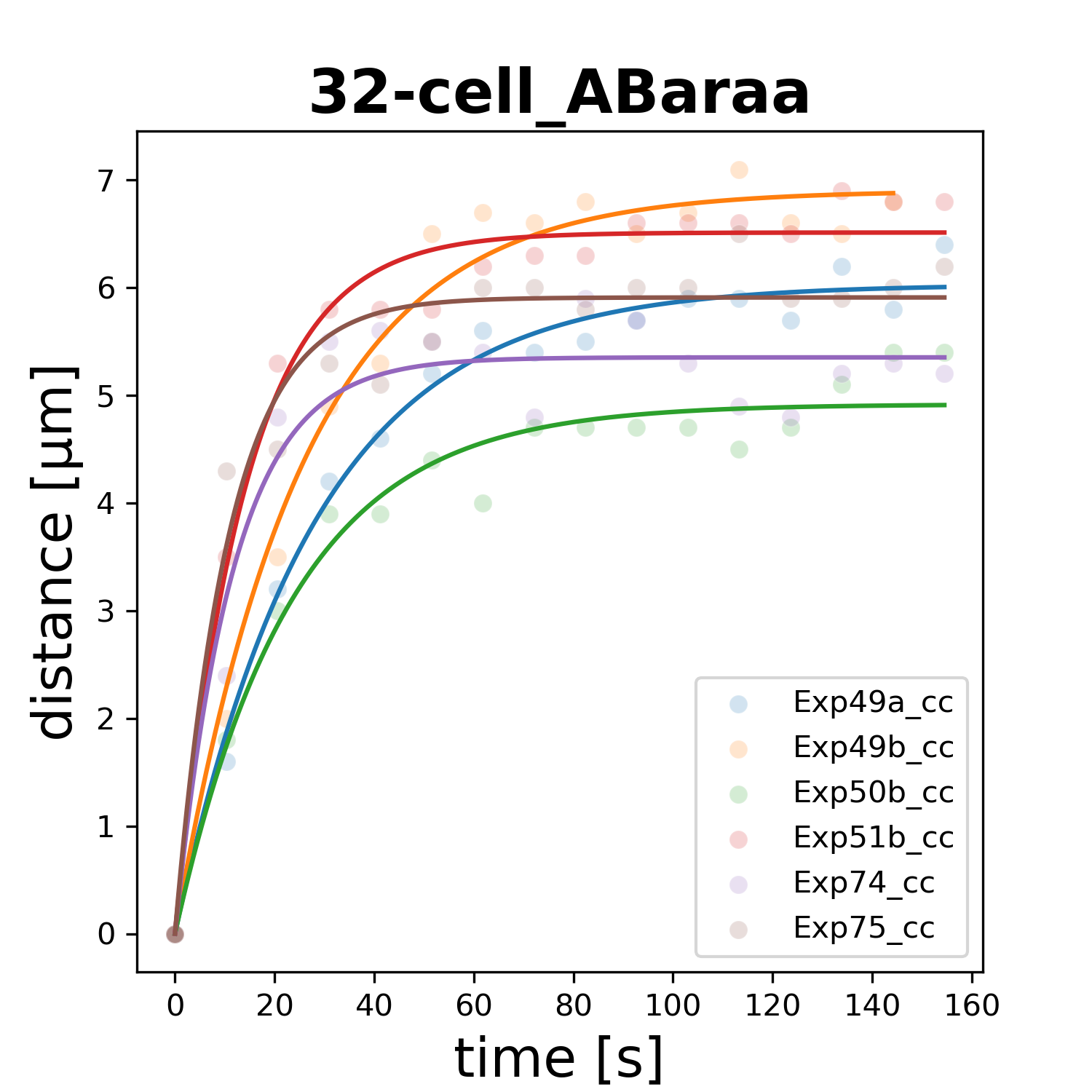

### 32-cell_ABarap.png

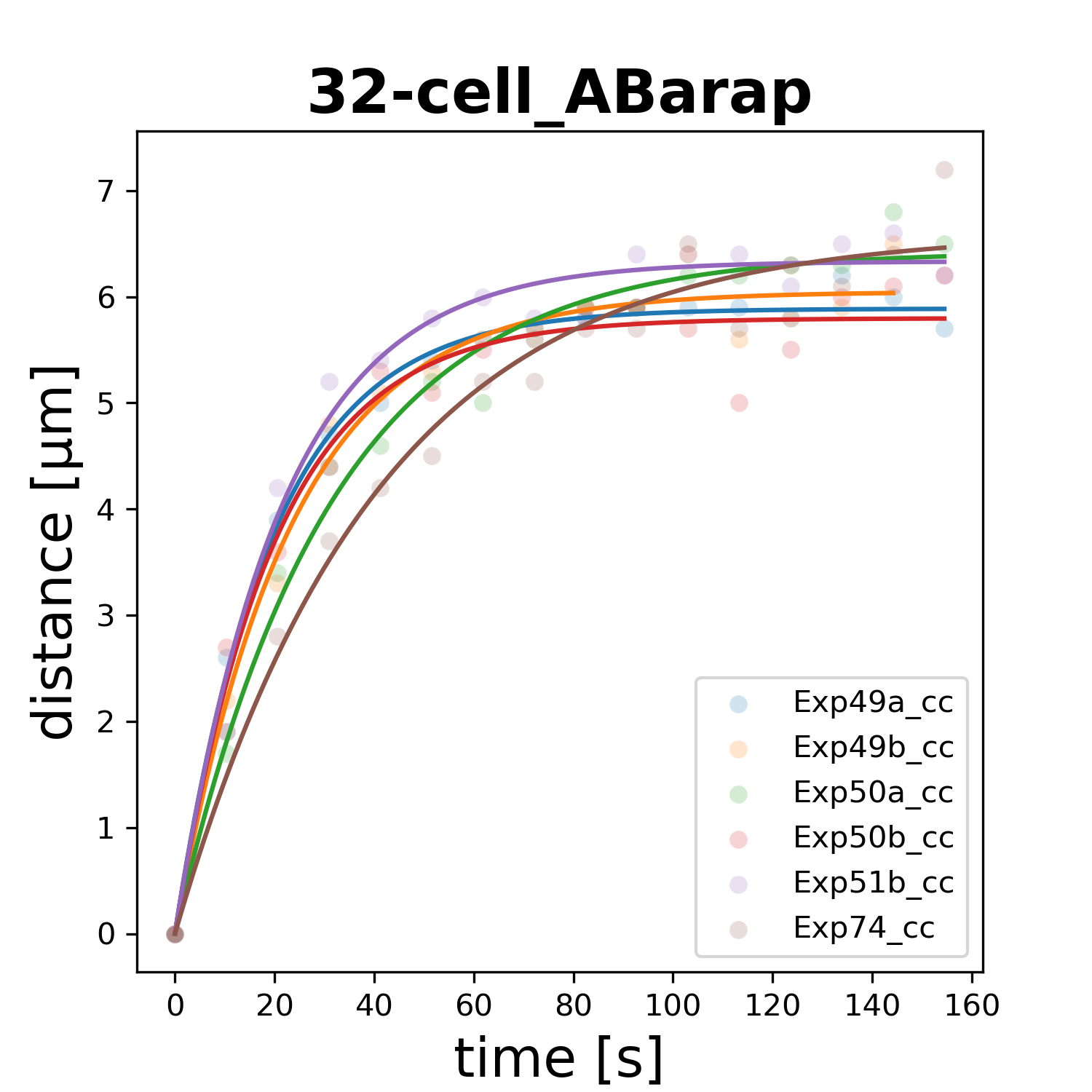

### 32-cell_ABarpa.png

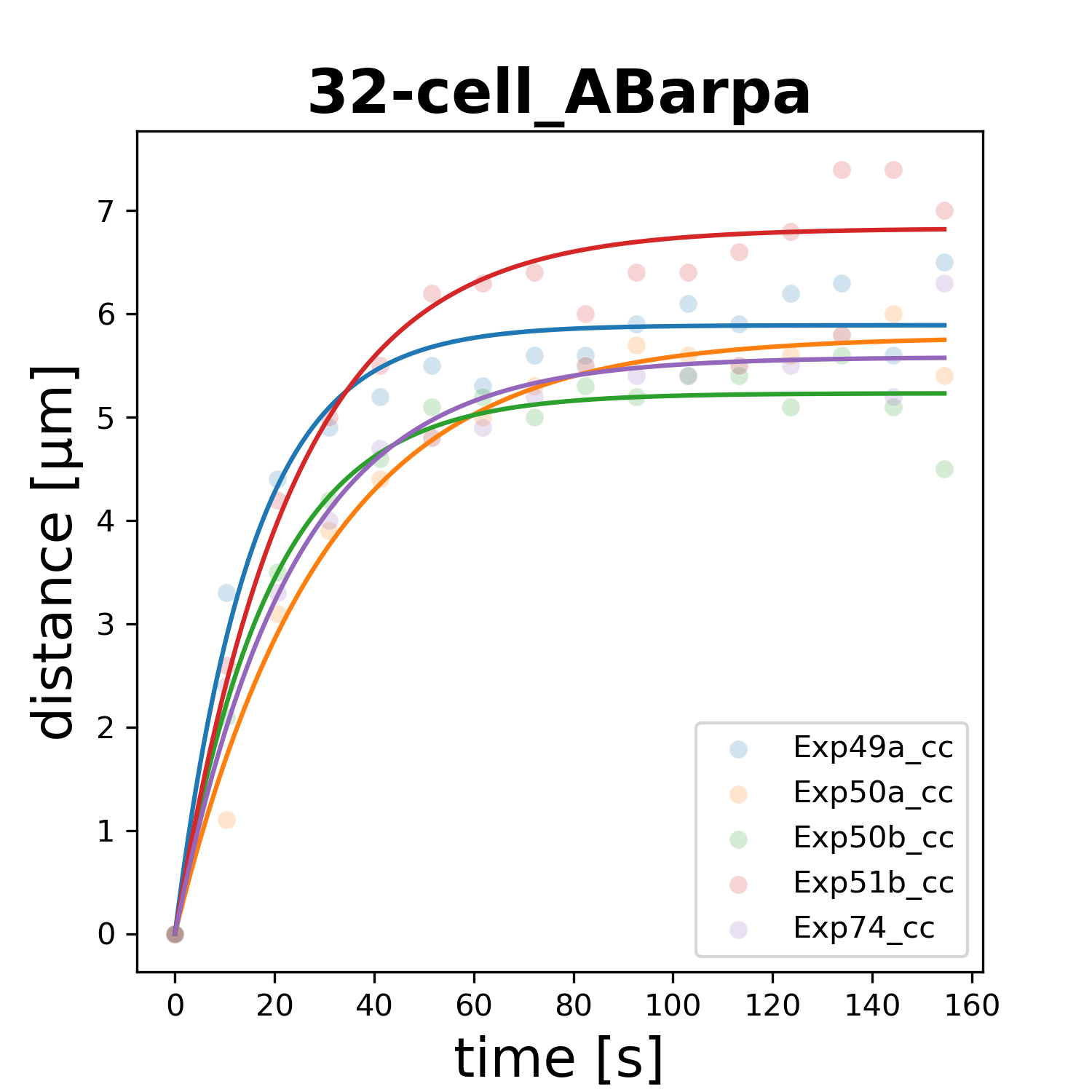

### 32-cell_ABarpp.png

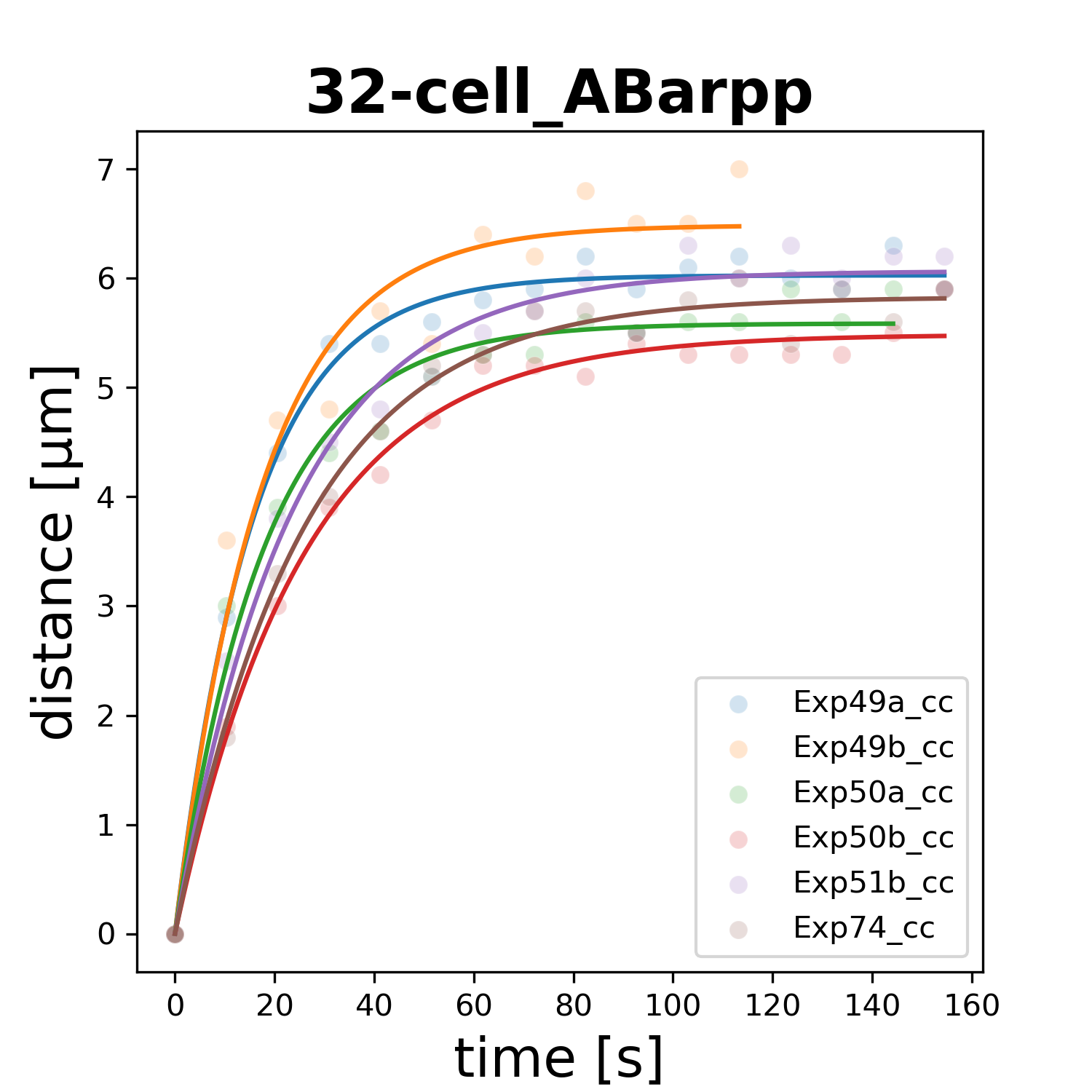

### 32-cell_ABplaa.png

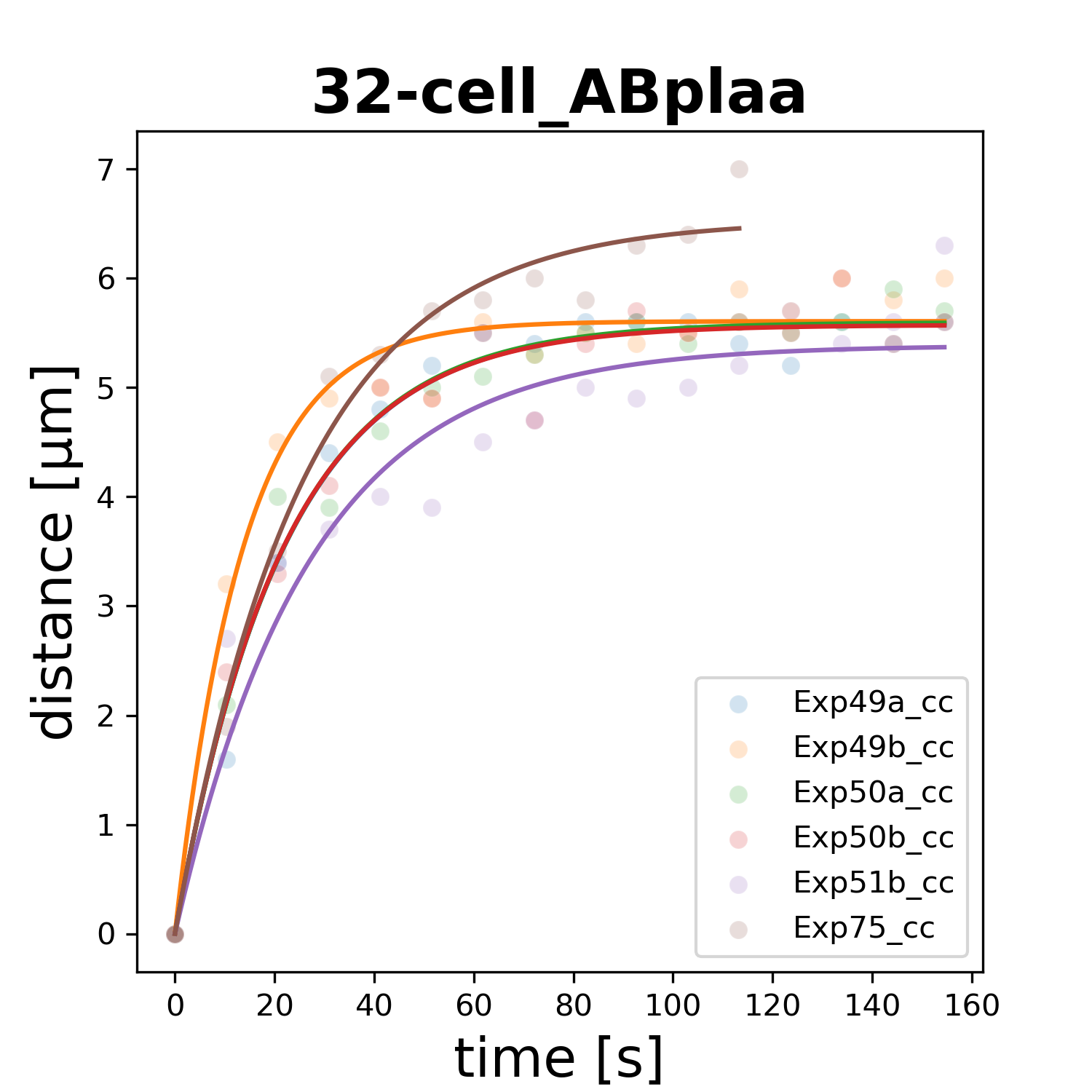

### 32-cell_ABplap.png

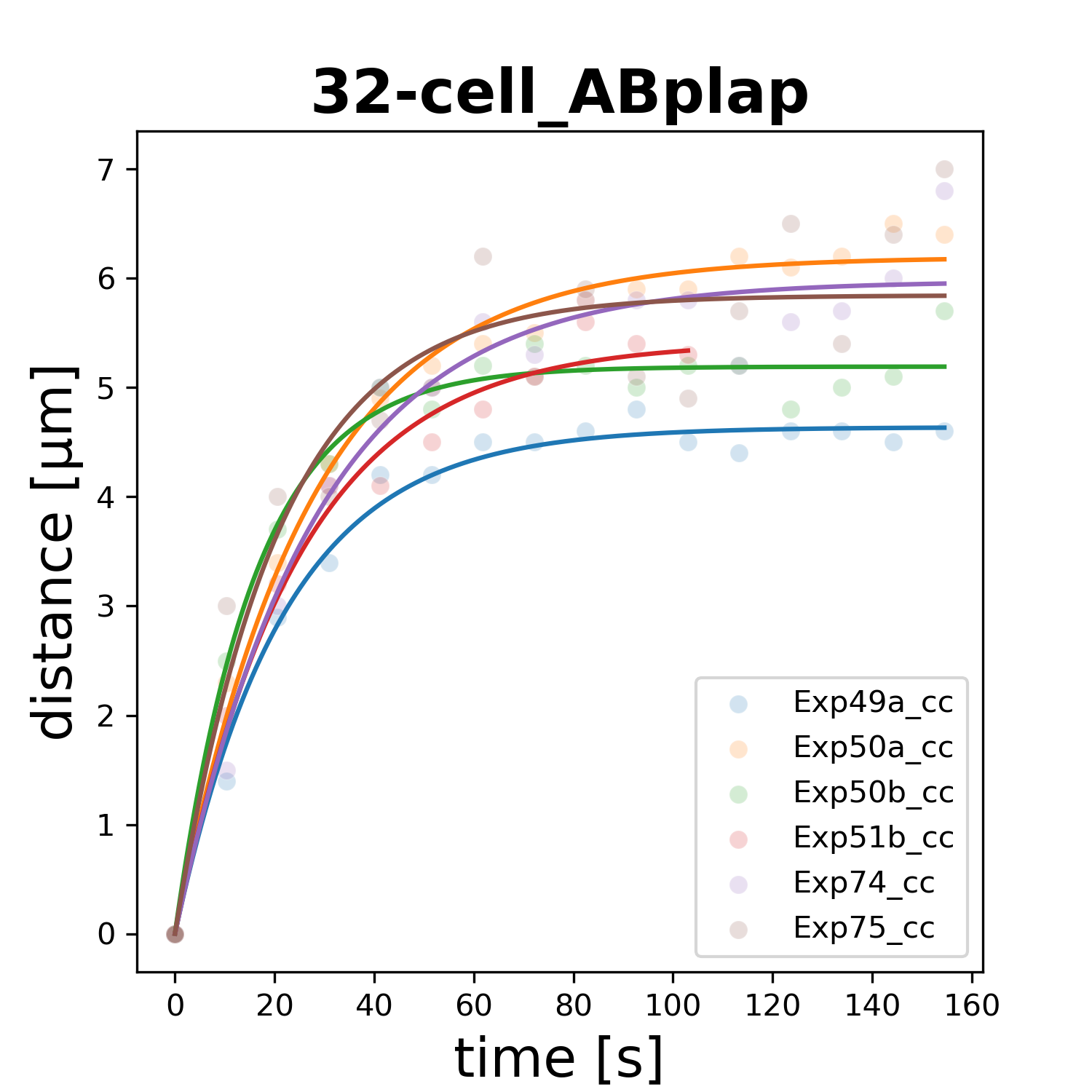

### 32-cell_ABplpa.png

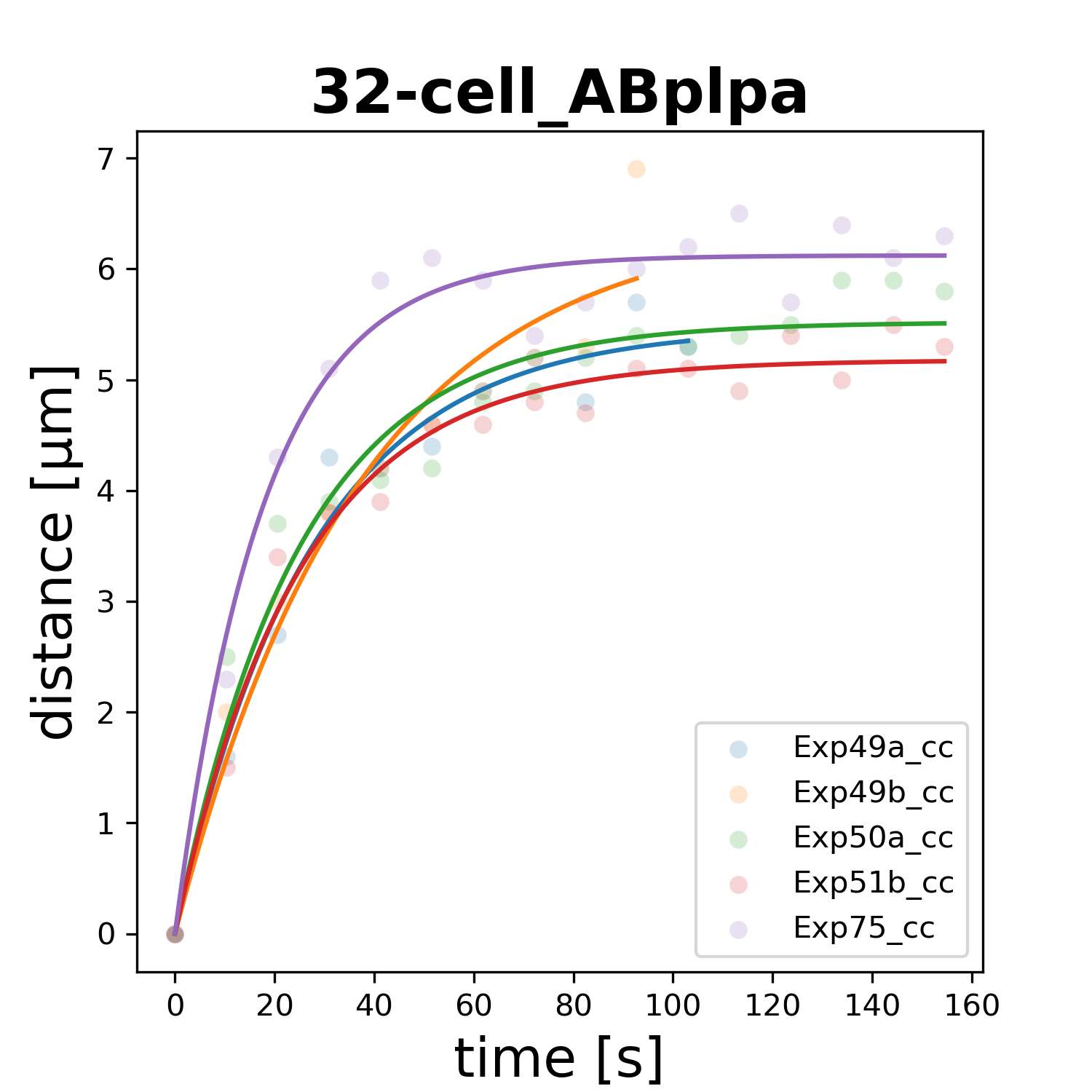

### 32-cell_ABplpp.png

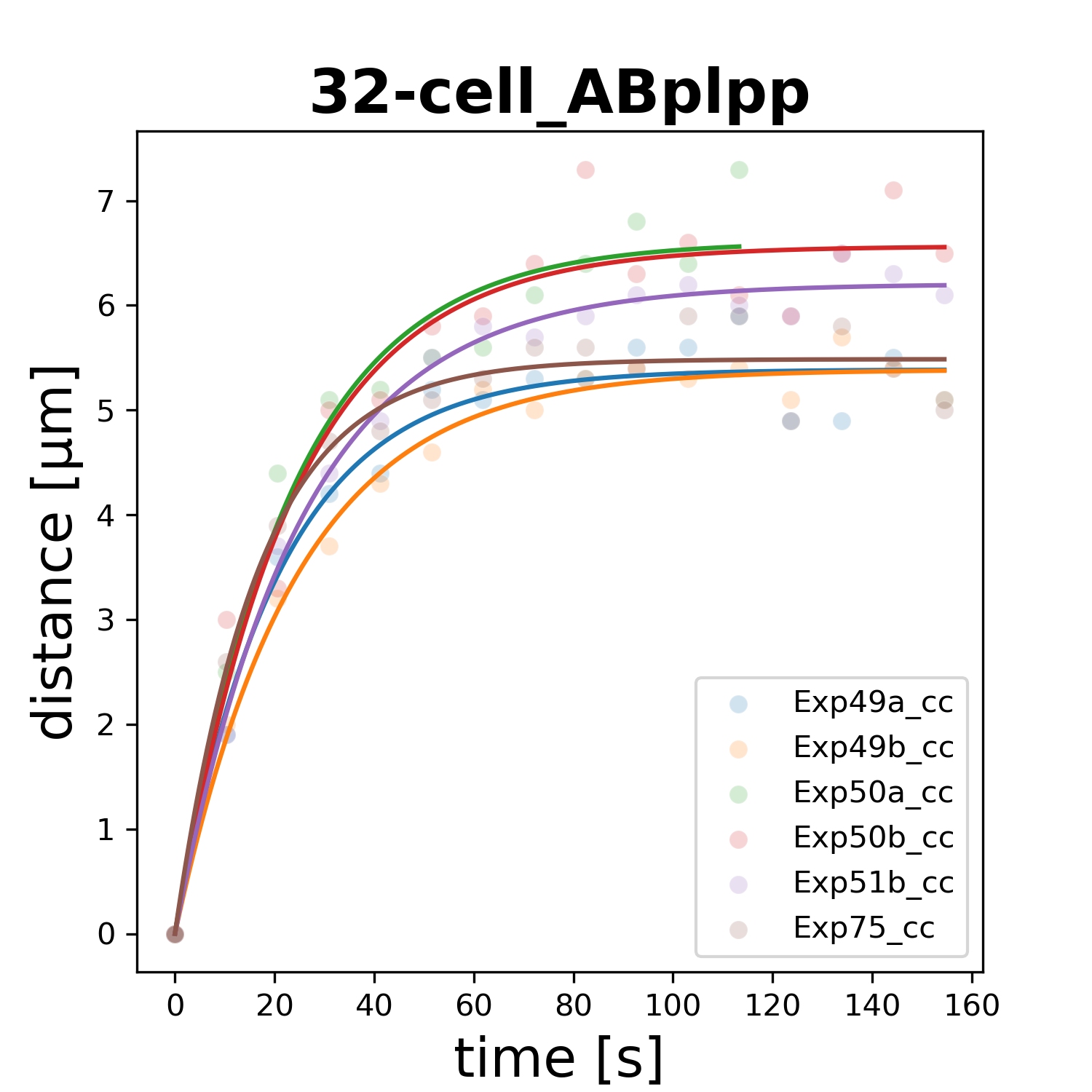

### 32-cell_ABpraa.png

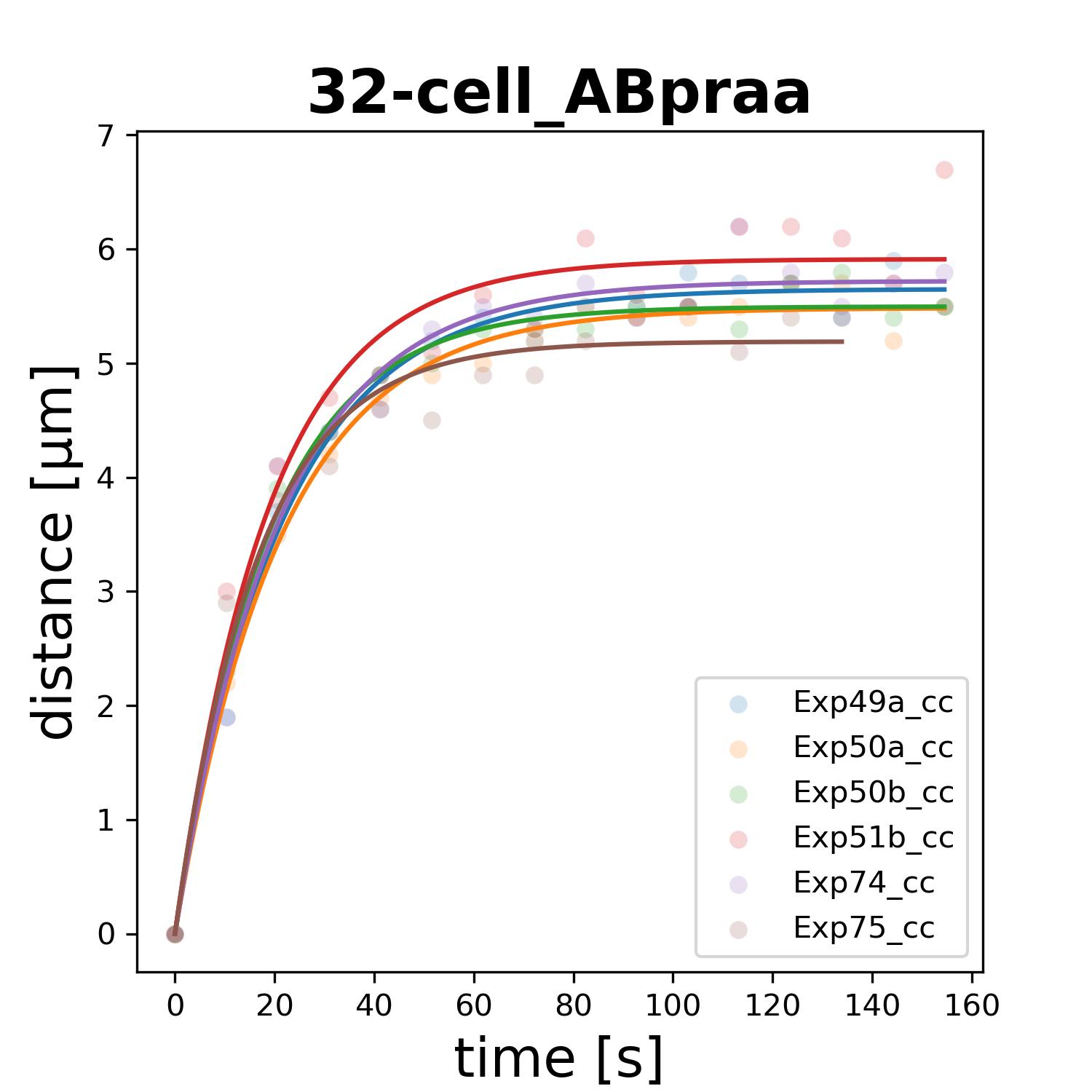

### 32-cell_ABprap.png

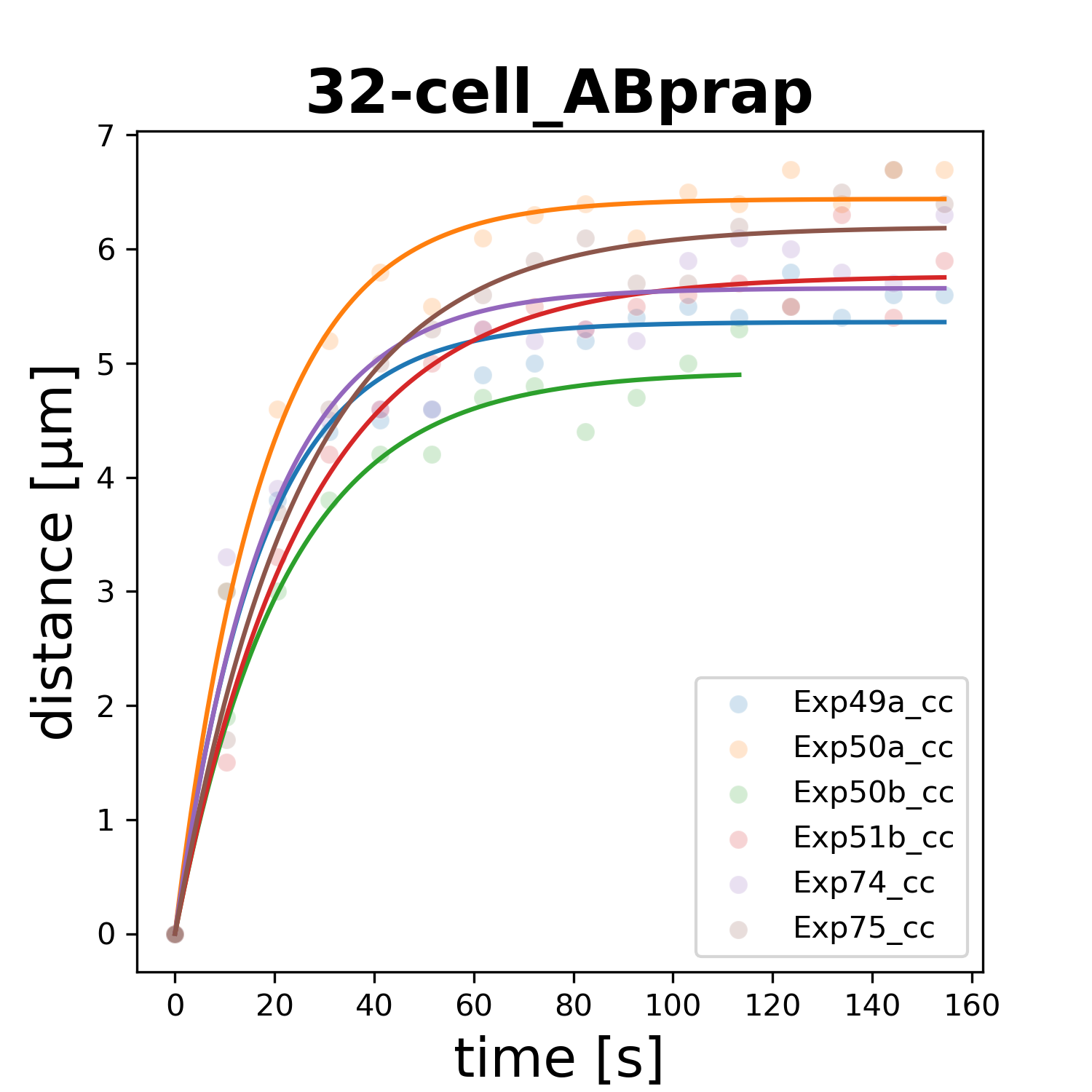

### 32-cell_ABprpa.png

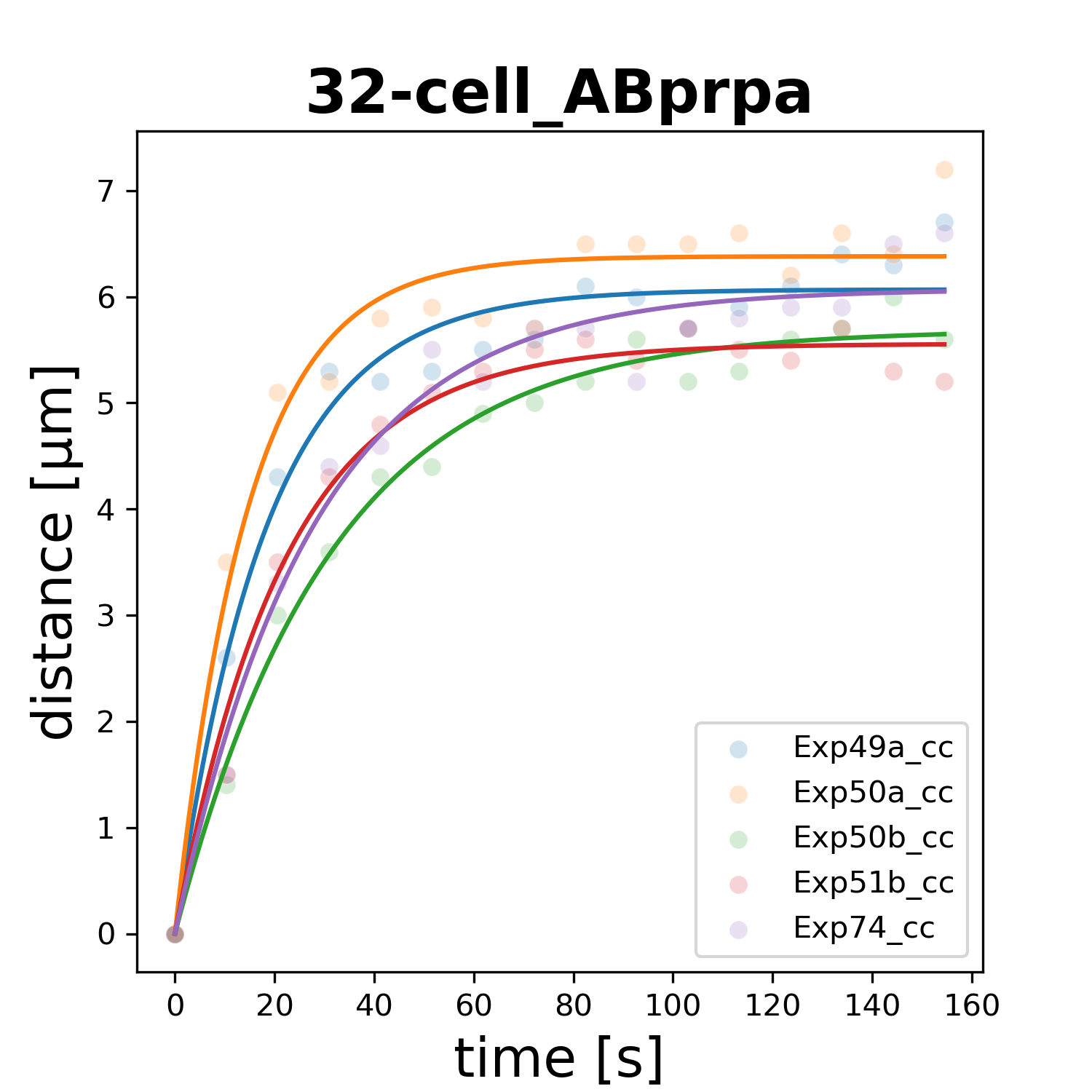

### 32-cell_ABprpp.png

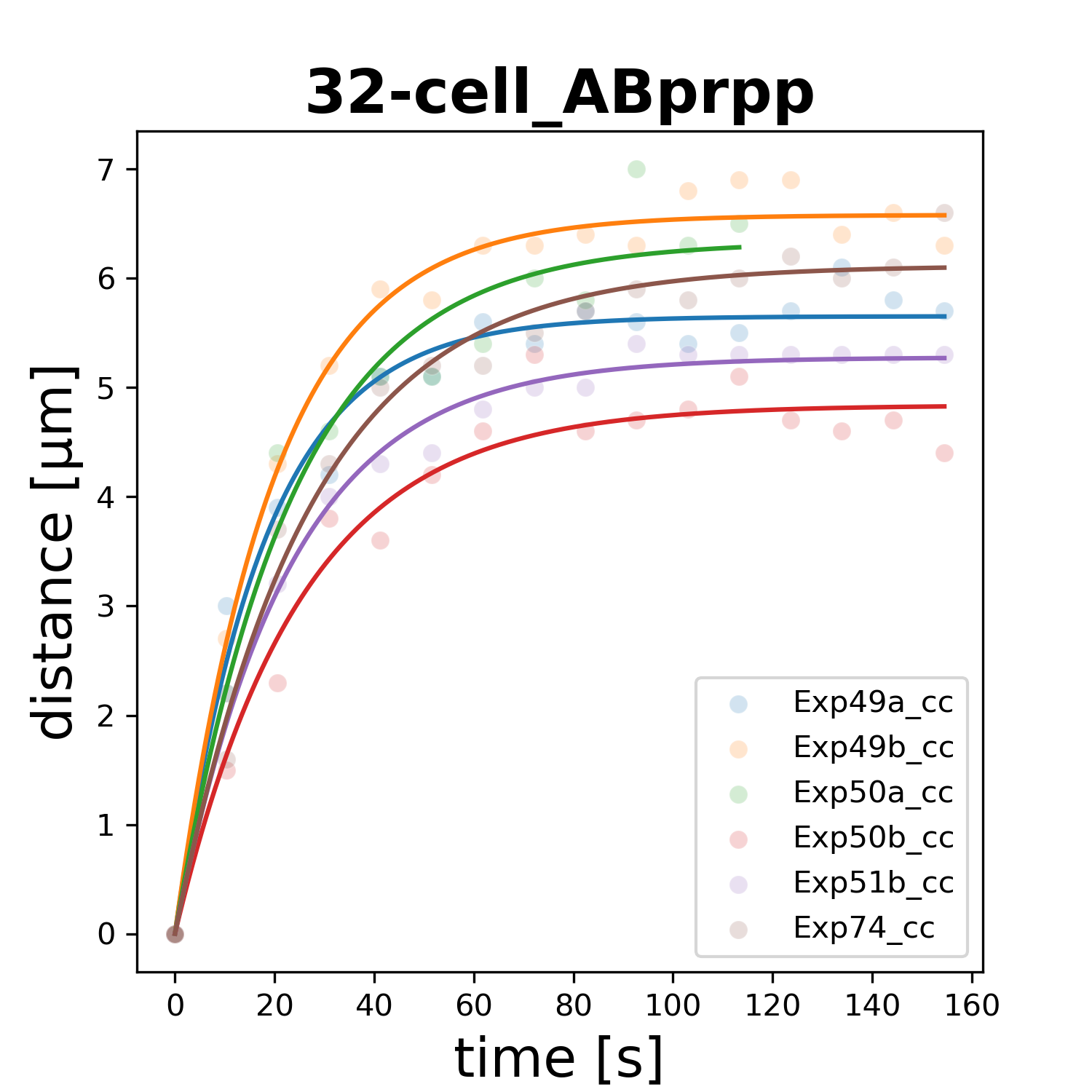

### 32-cell_Caa.png

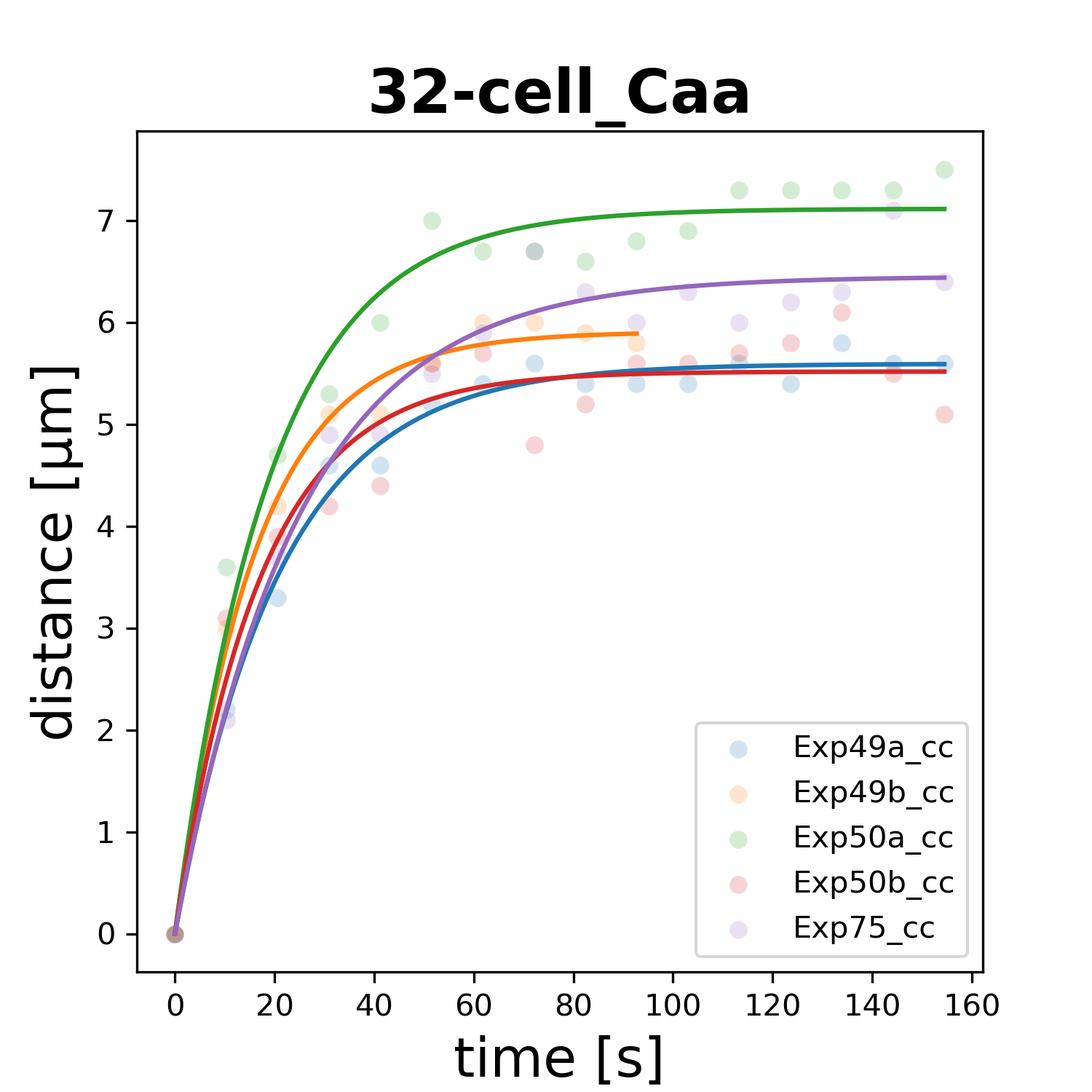

### 32-cell_Cap.png

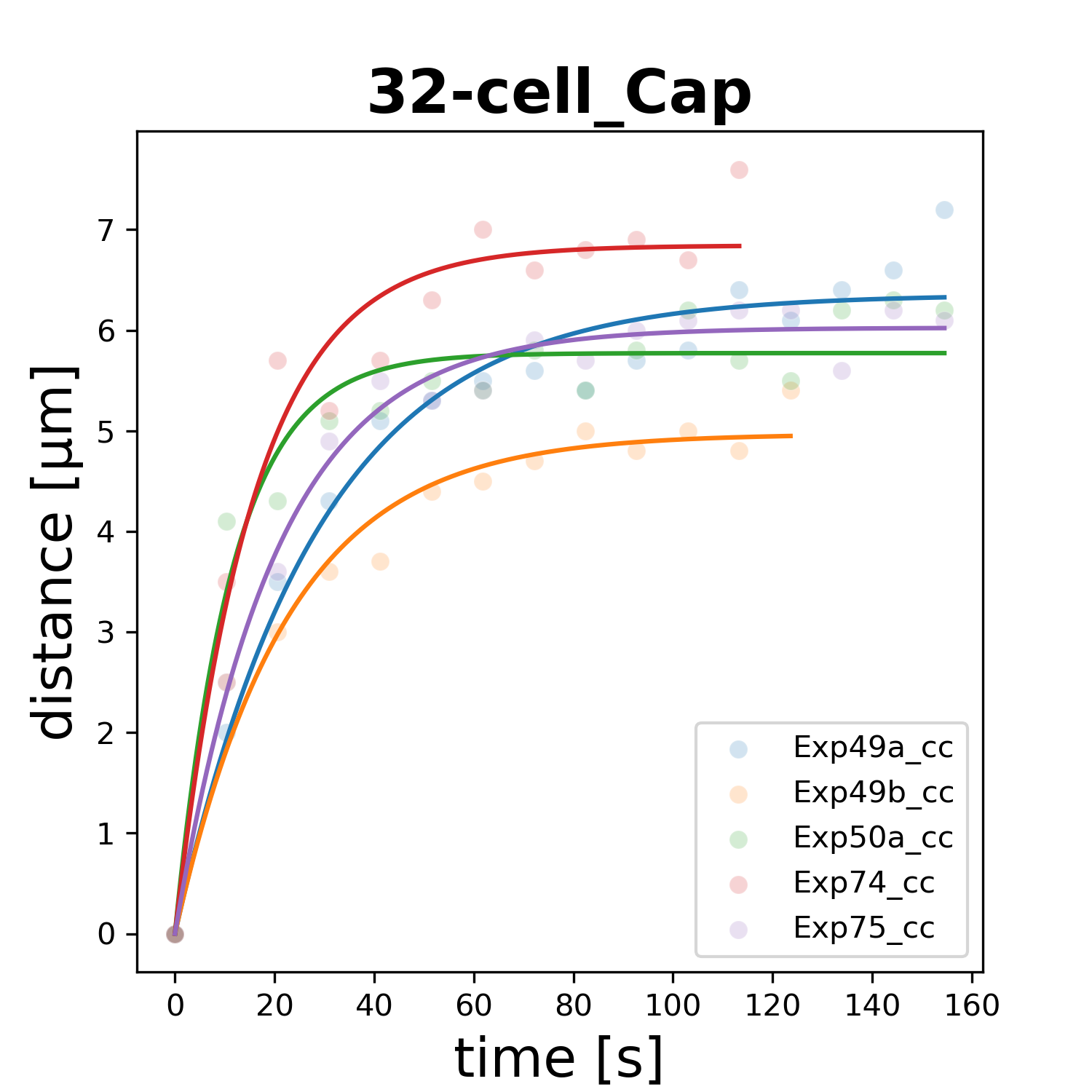

### 32-cell_Cpa.png

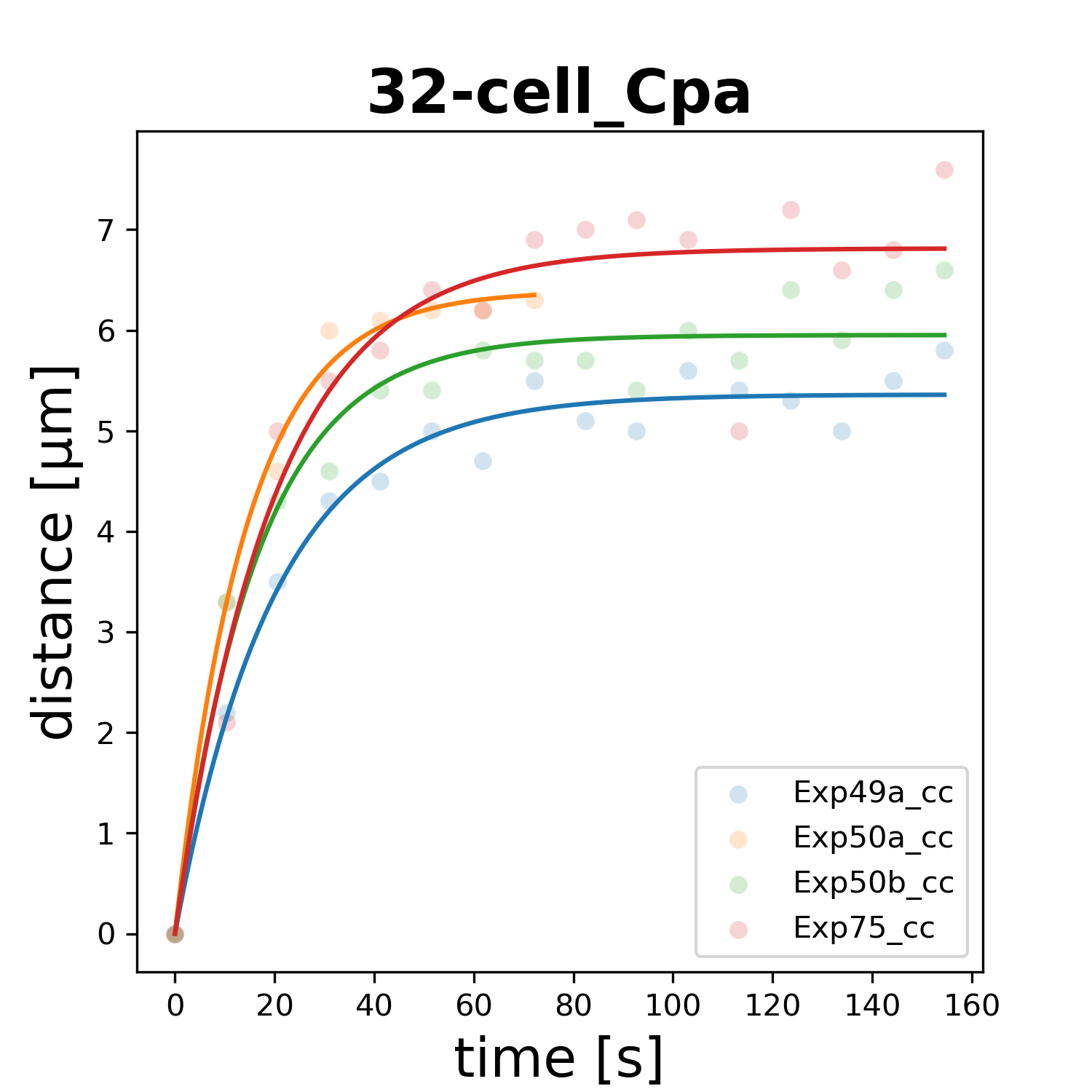

### 32-cell_Cpp.png

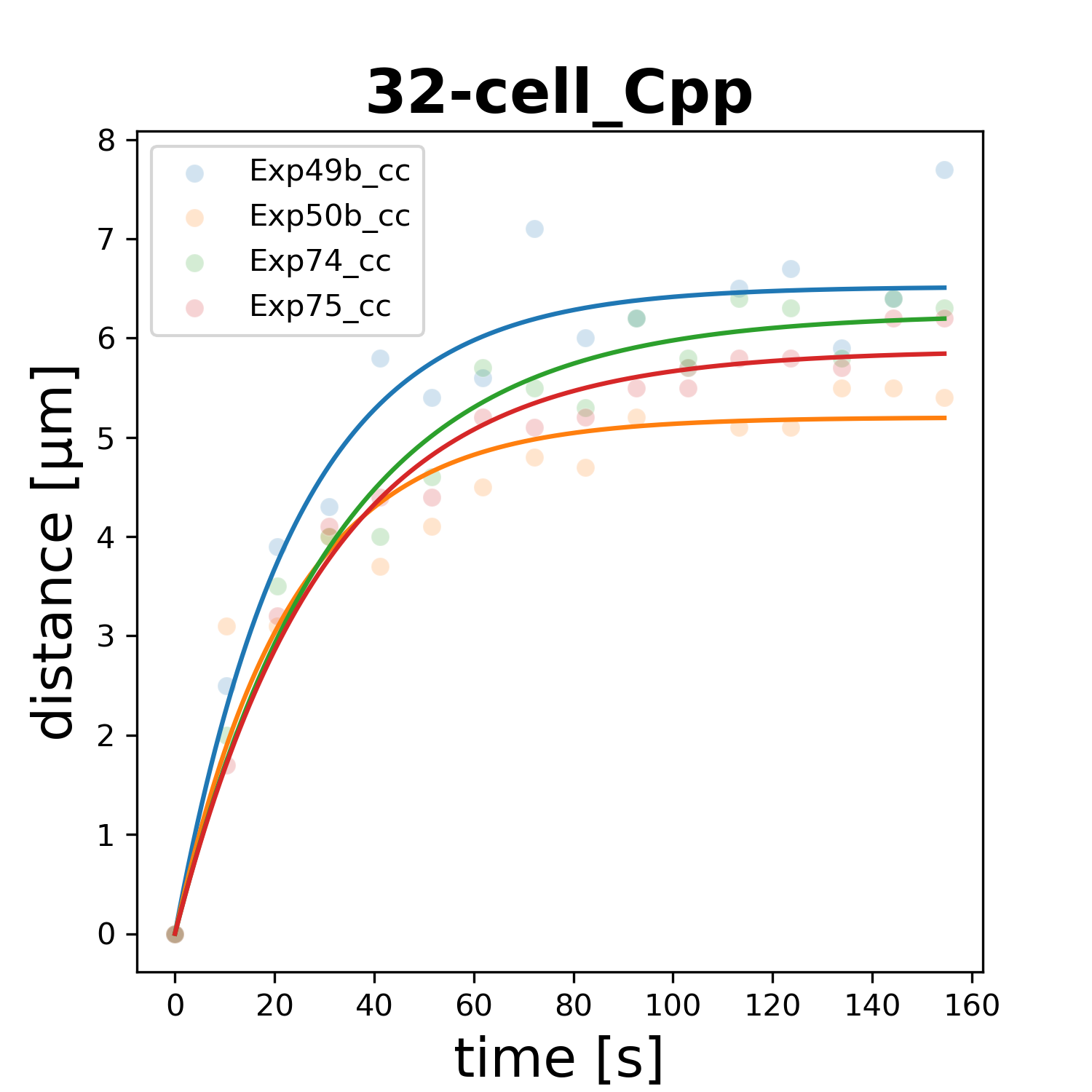

### 32-cell_Dp.png

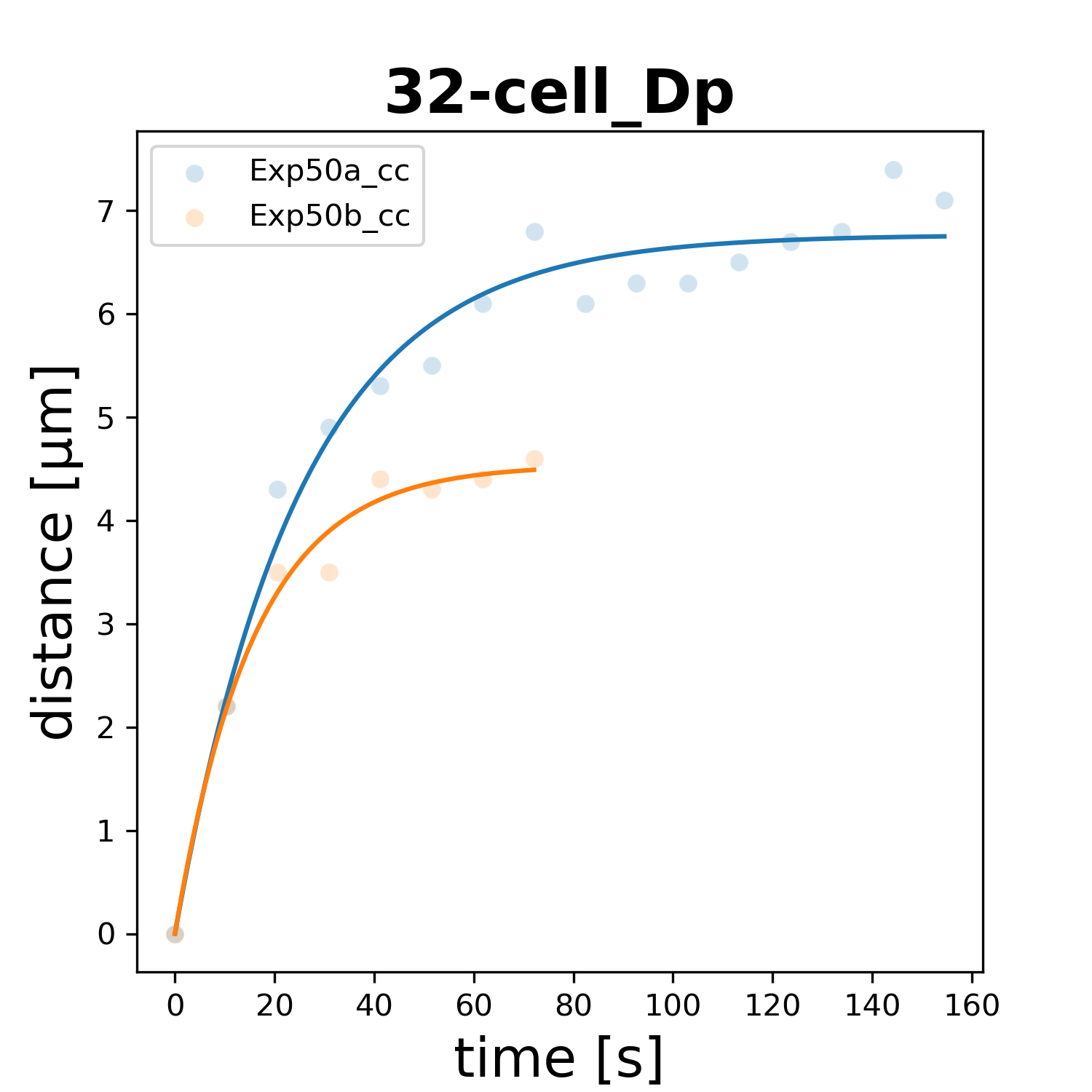

### 32-cell_Eal.png

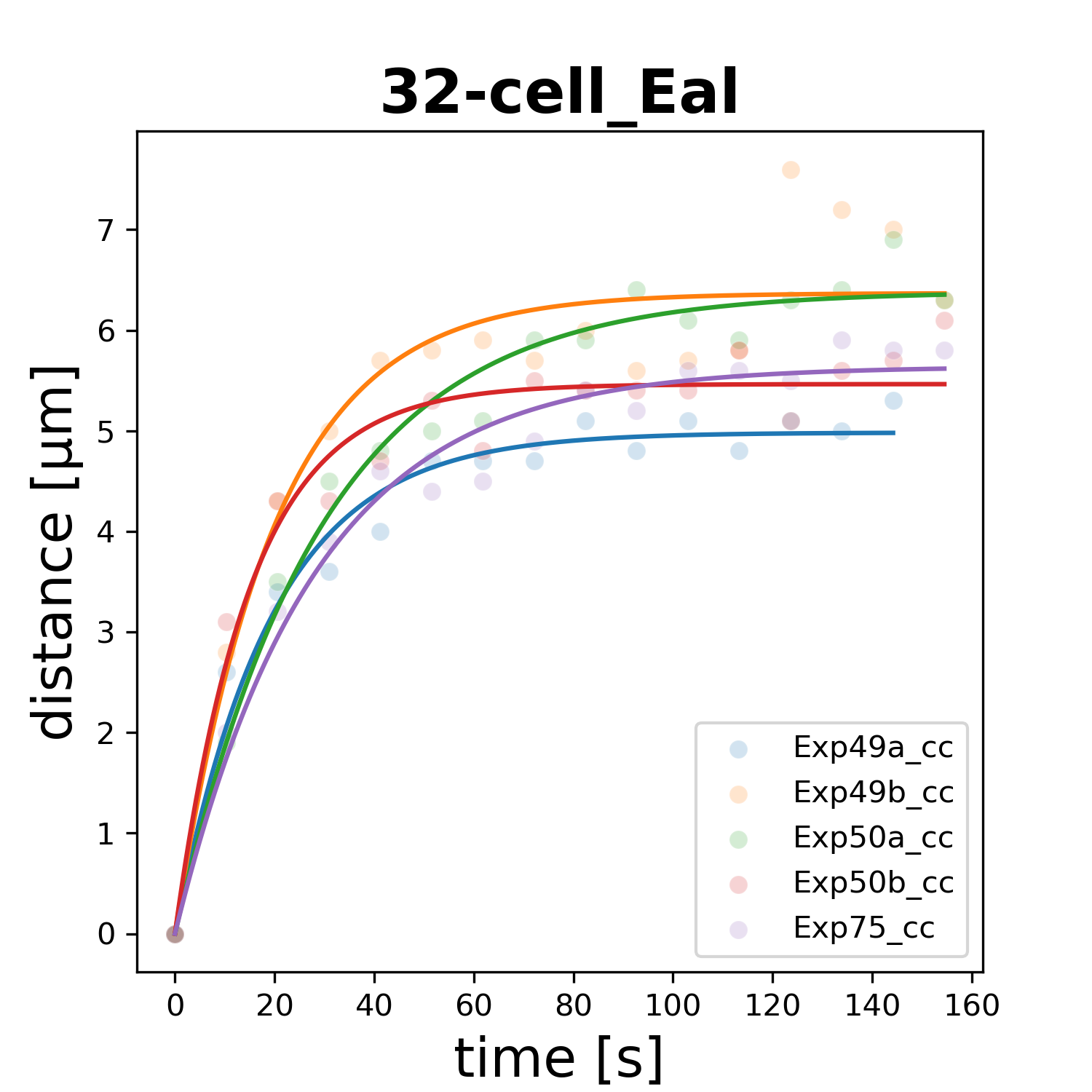

### 32-cell_Ear.png

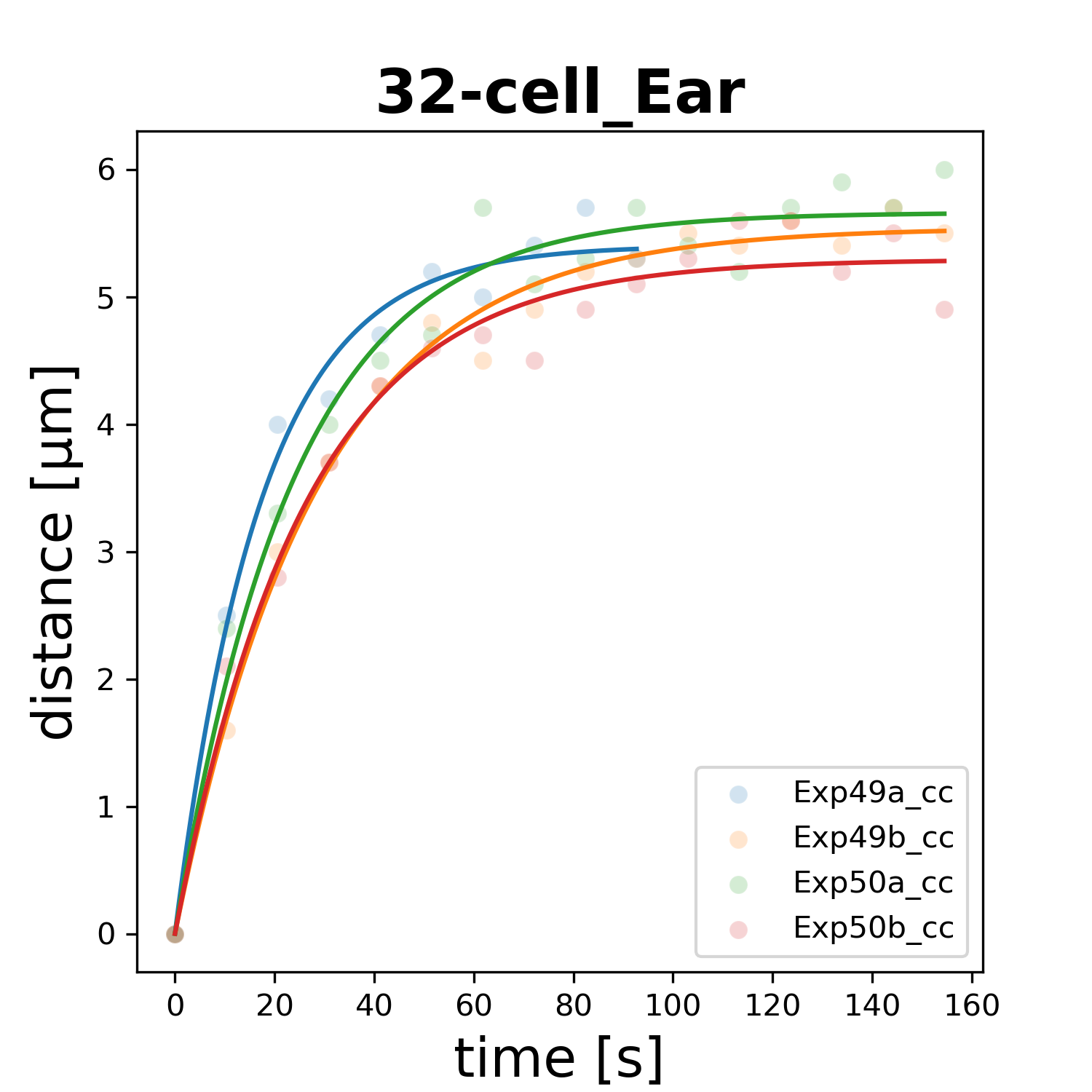

### 32-cell_Epl.png

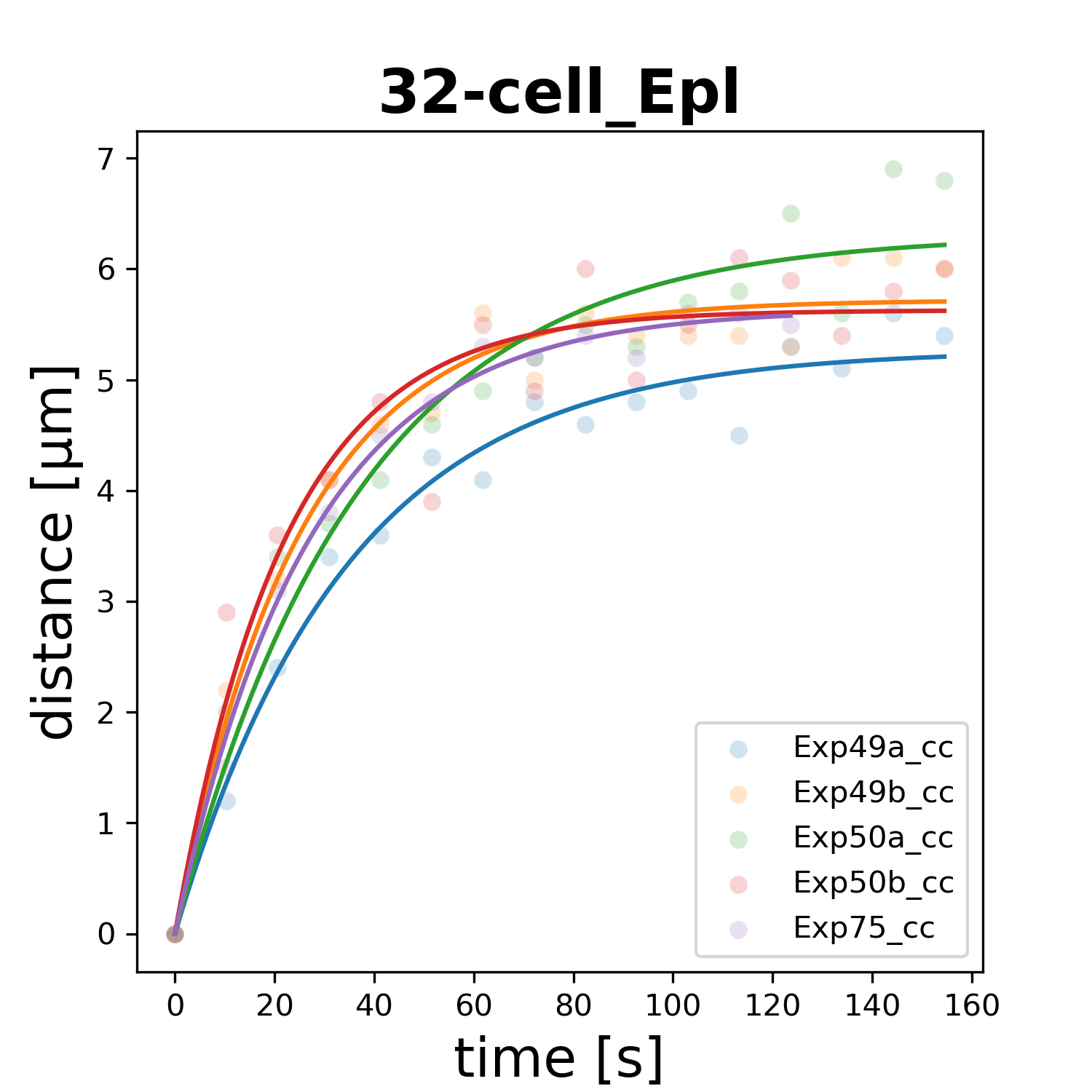

### 32-cell_Epr.png

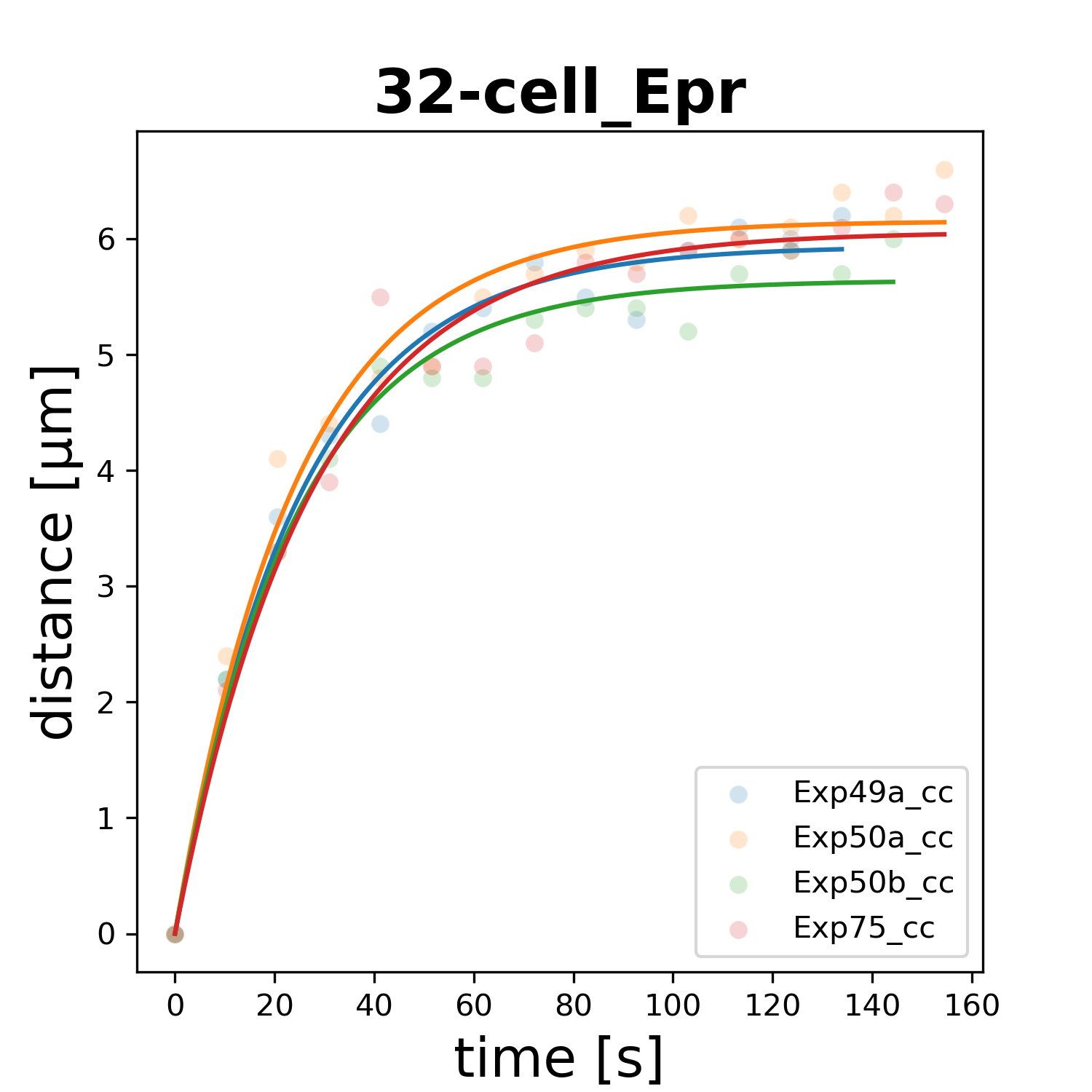

### 32-cell_MSaa.png

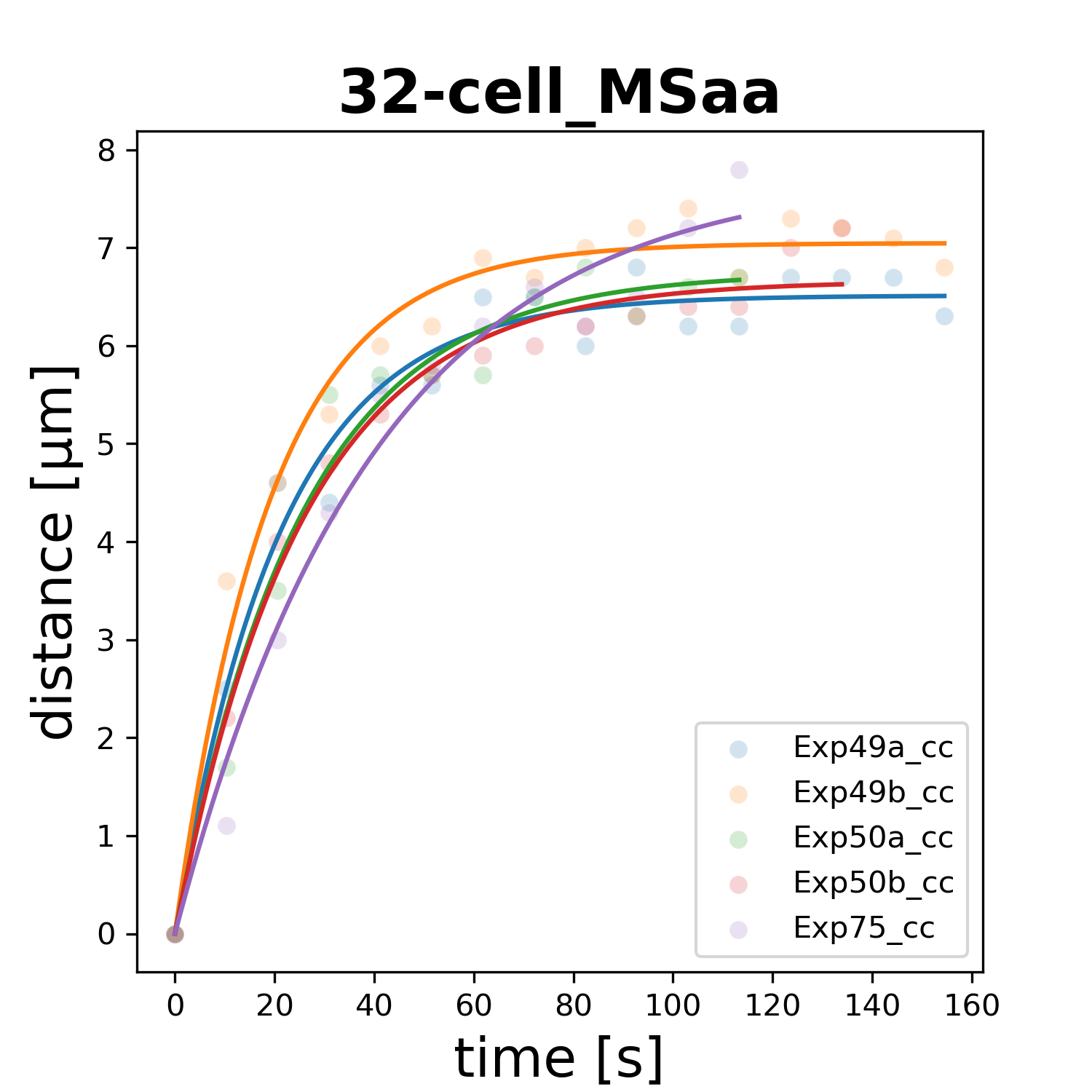

### 32-cell_MSap.png

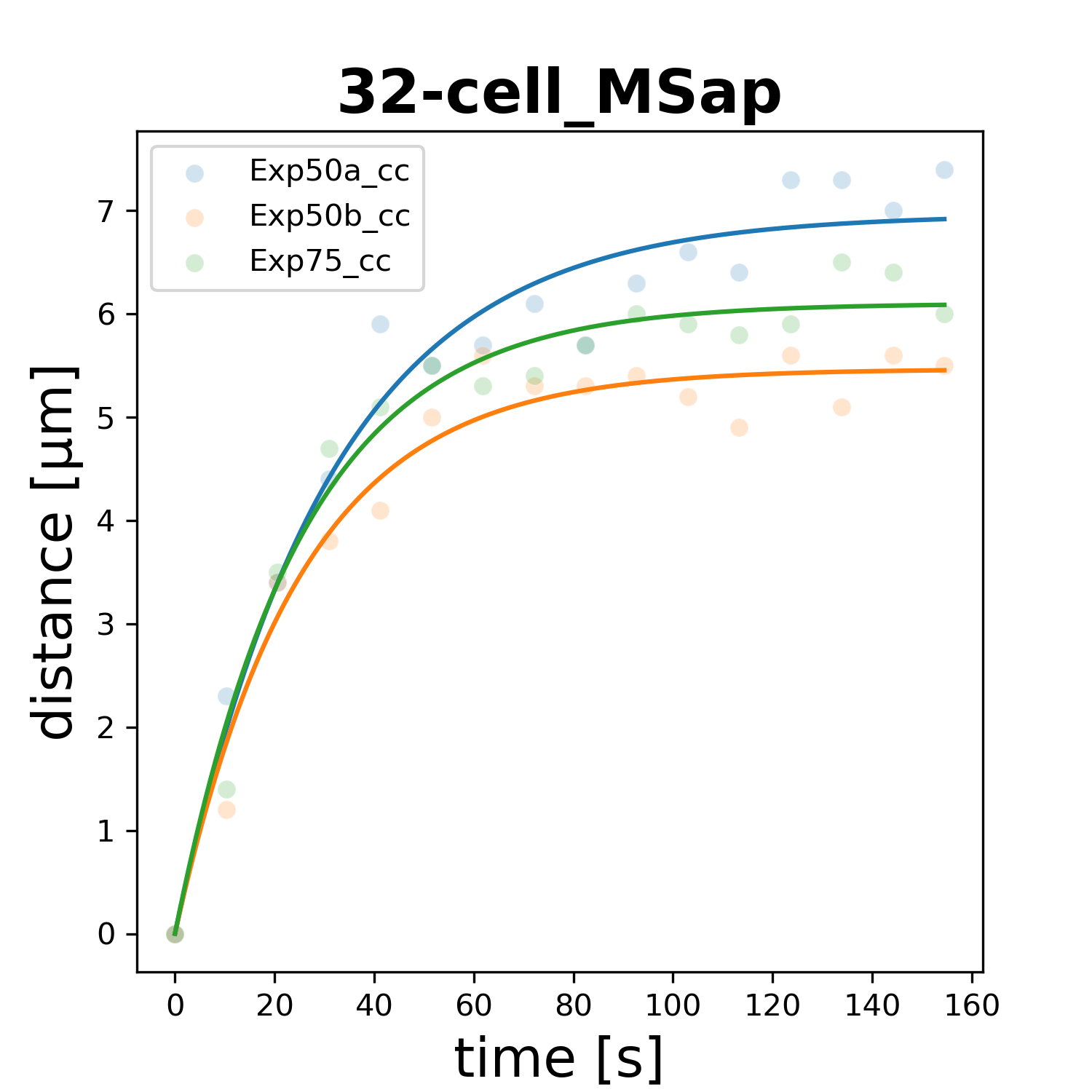

### 32-cell_MSpa.png

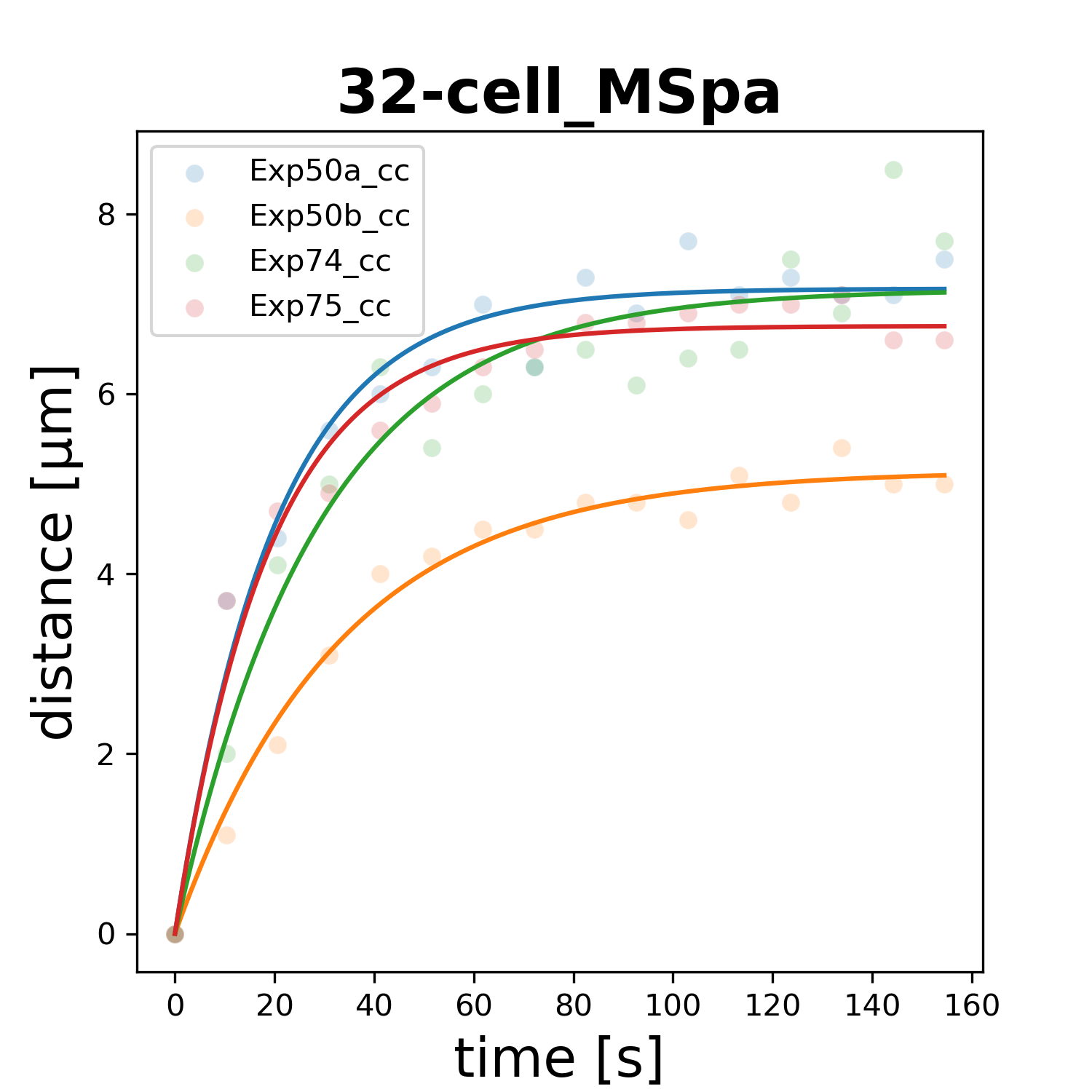

### 32-cell_MSpp.png

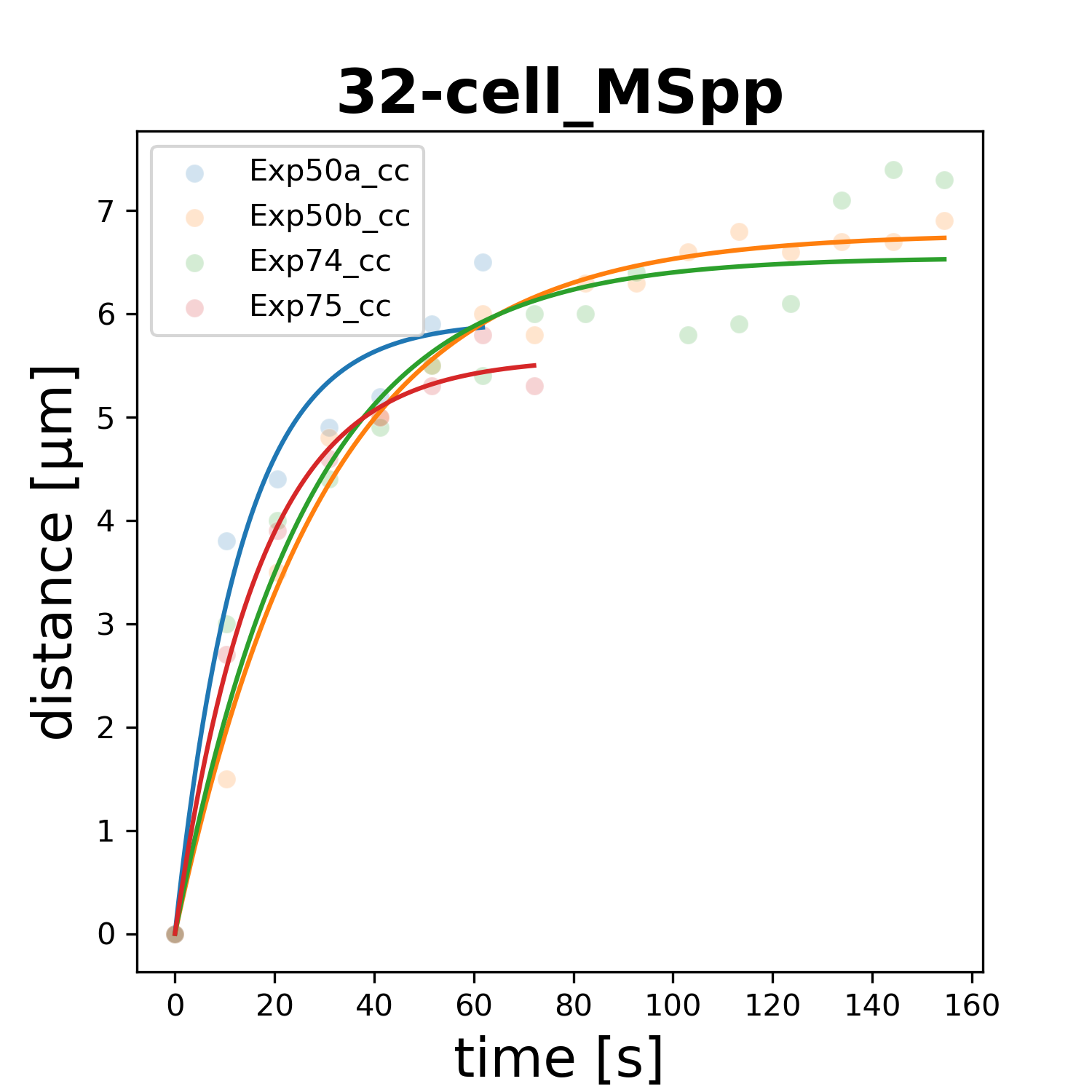

### 64-cell_ABalaaa.png

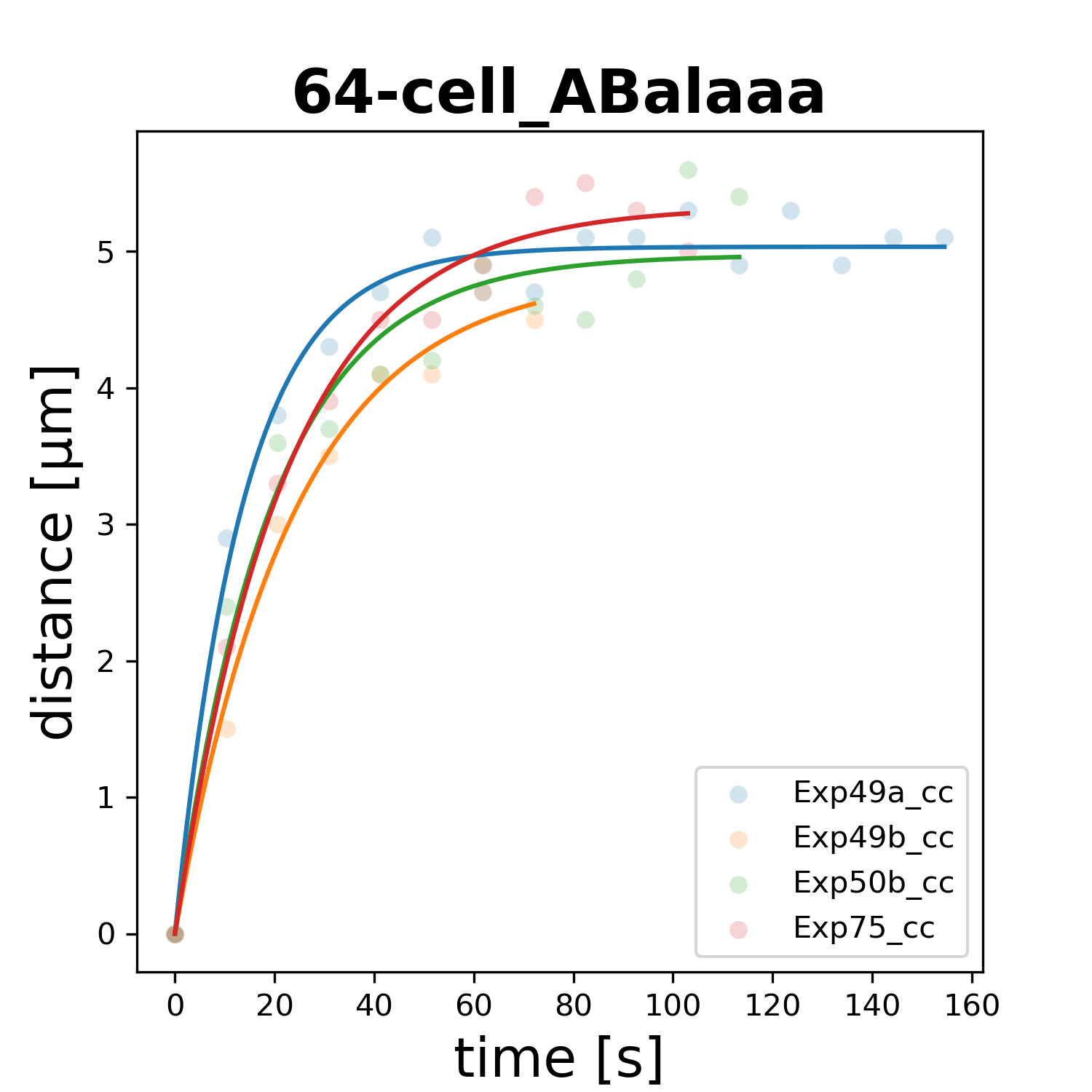

### 64-cell_ABalaap.png

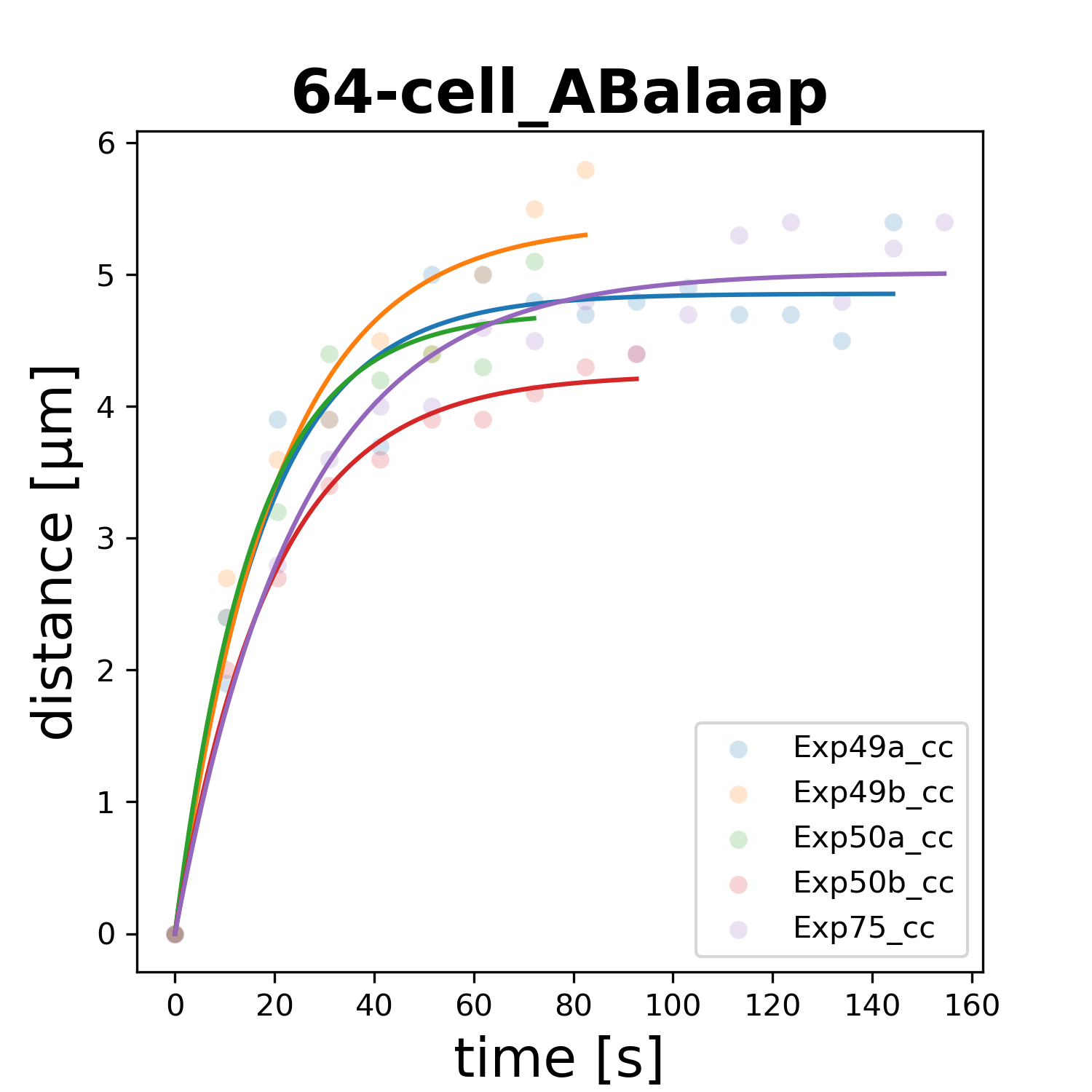

### 64-cell_ABalapa.png

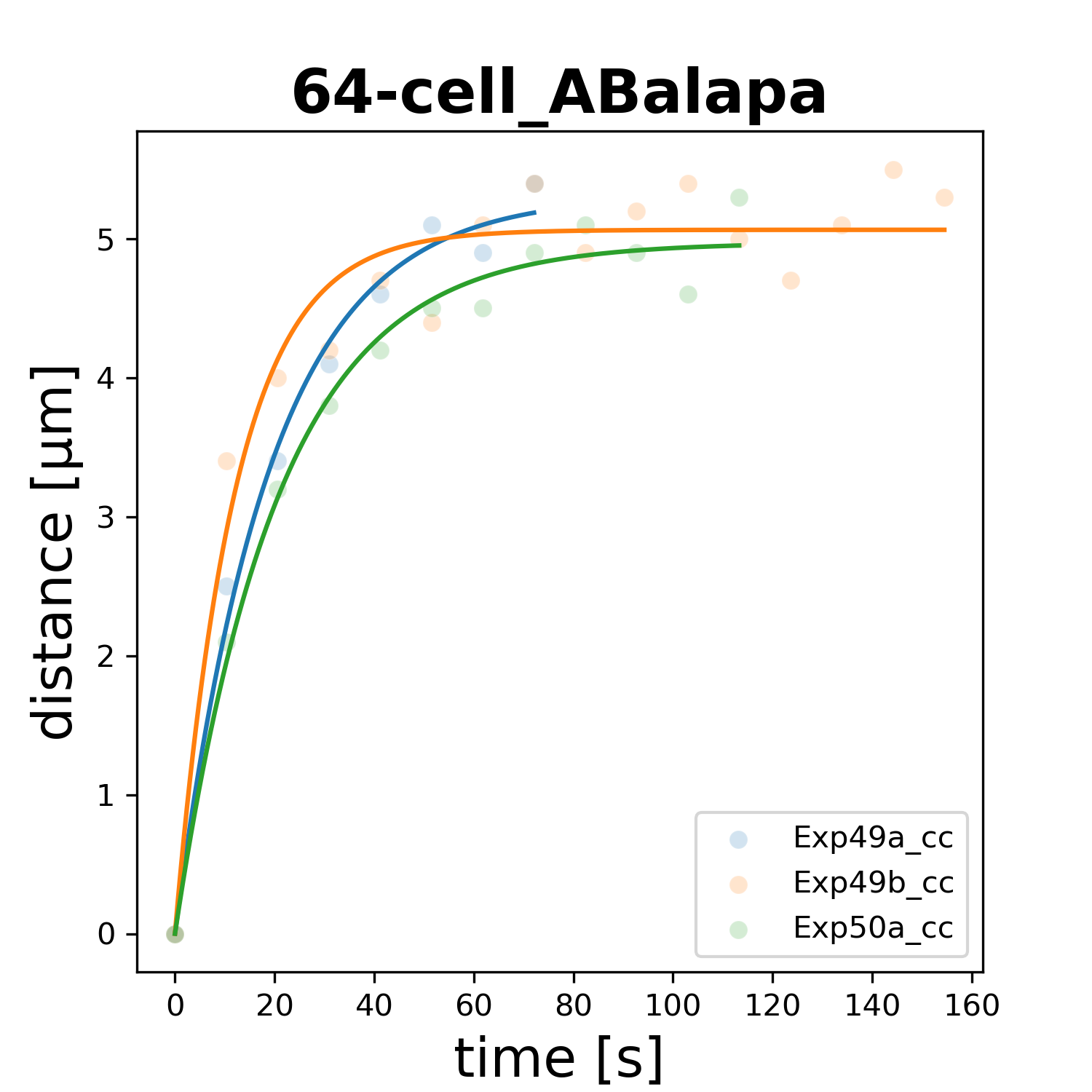

### 64-cell_ABalapp.png

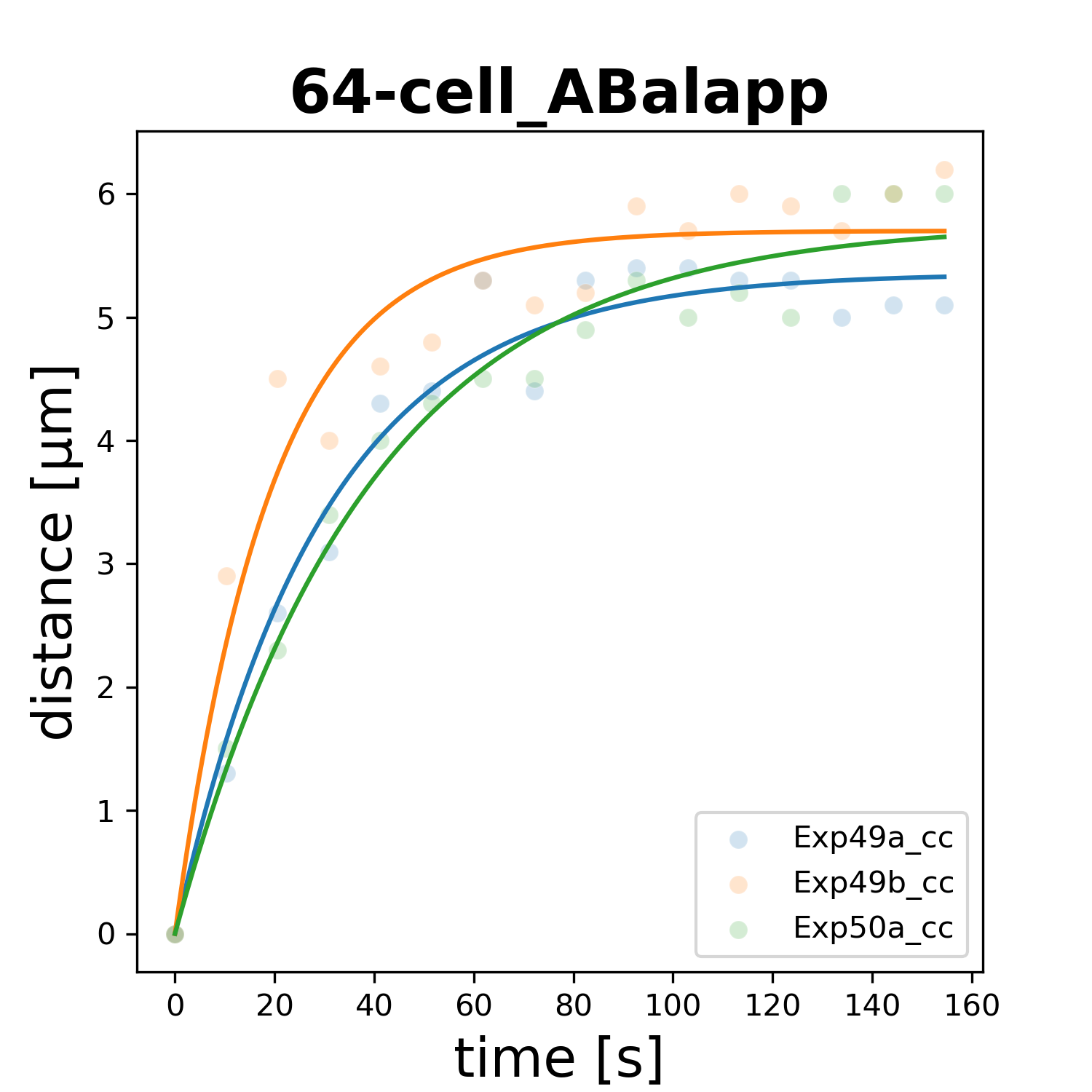

### 64-cell_ABalpaa.png

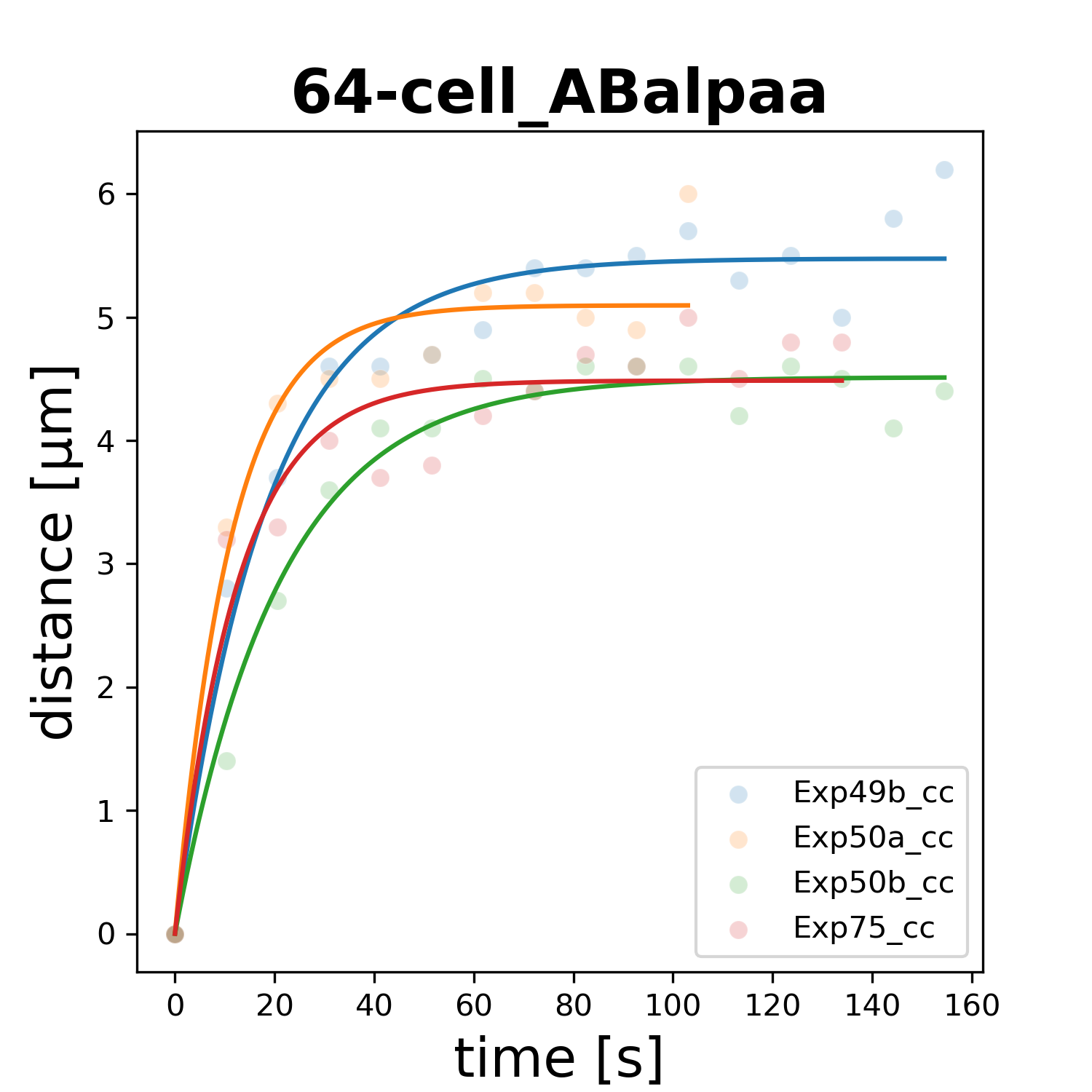

### 64-cell_ABalpap.png

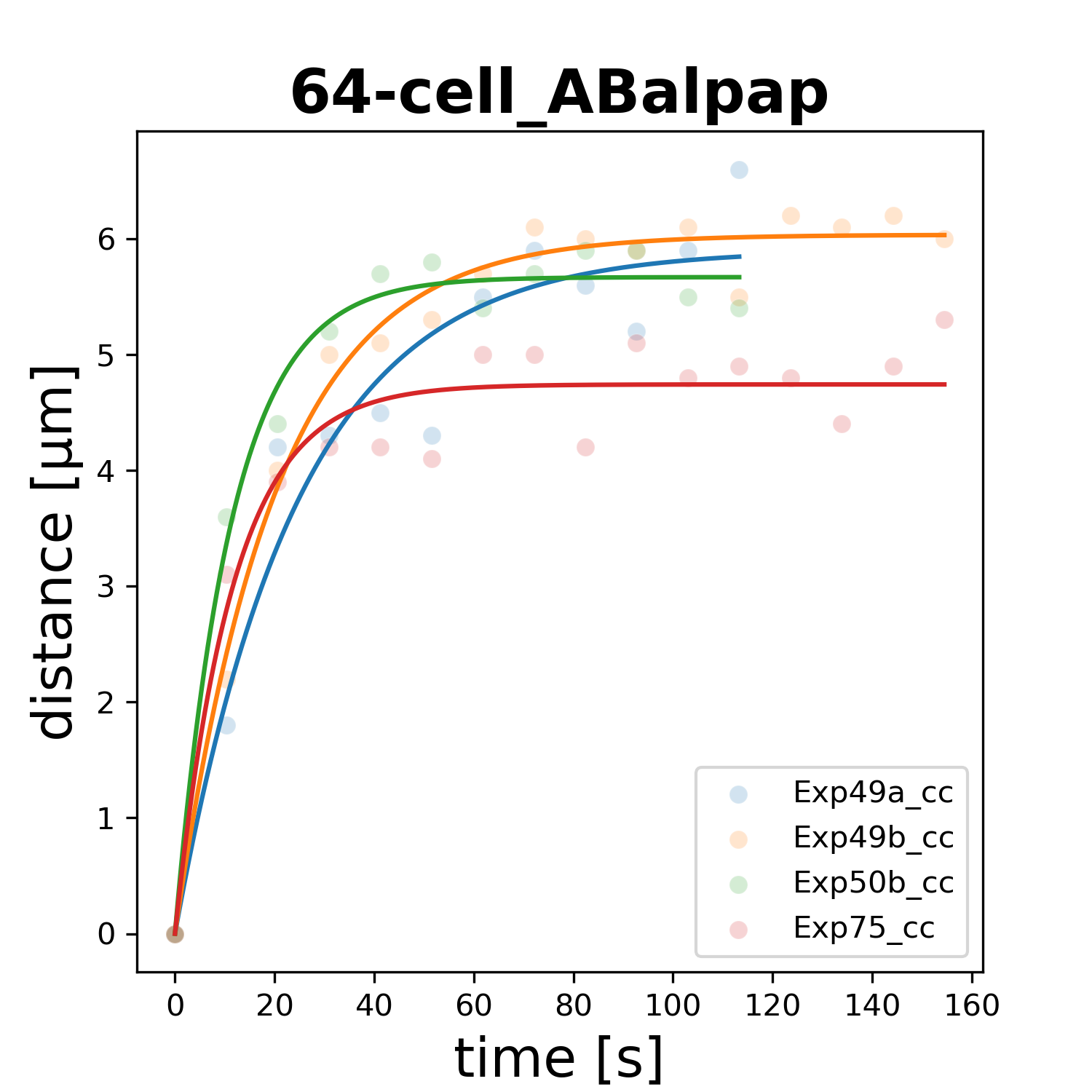

### 64-cell_ABalppa.png

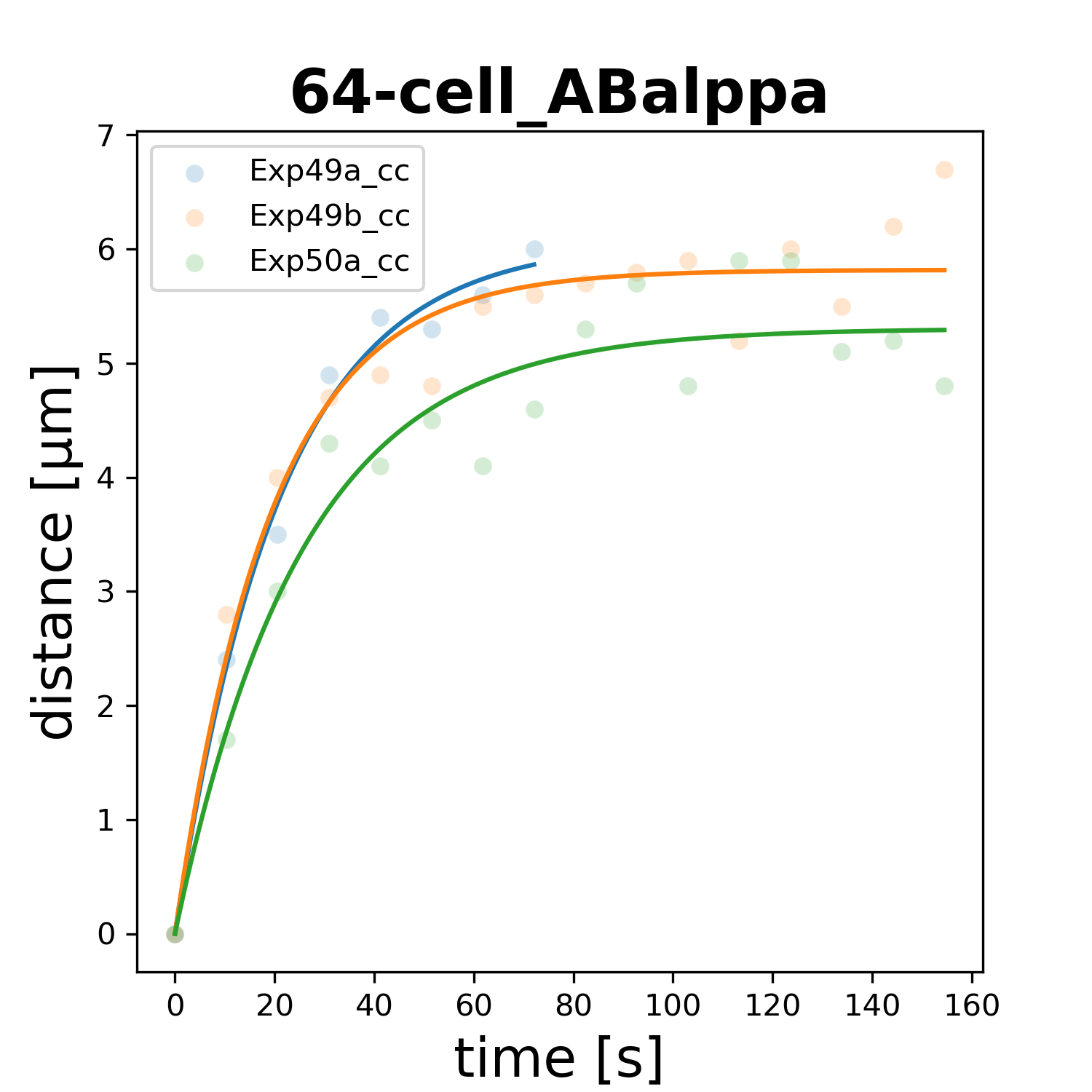

### 64-cell_ABalppp.png

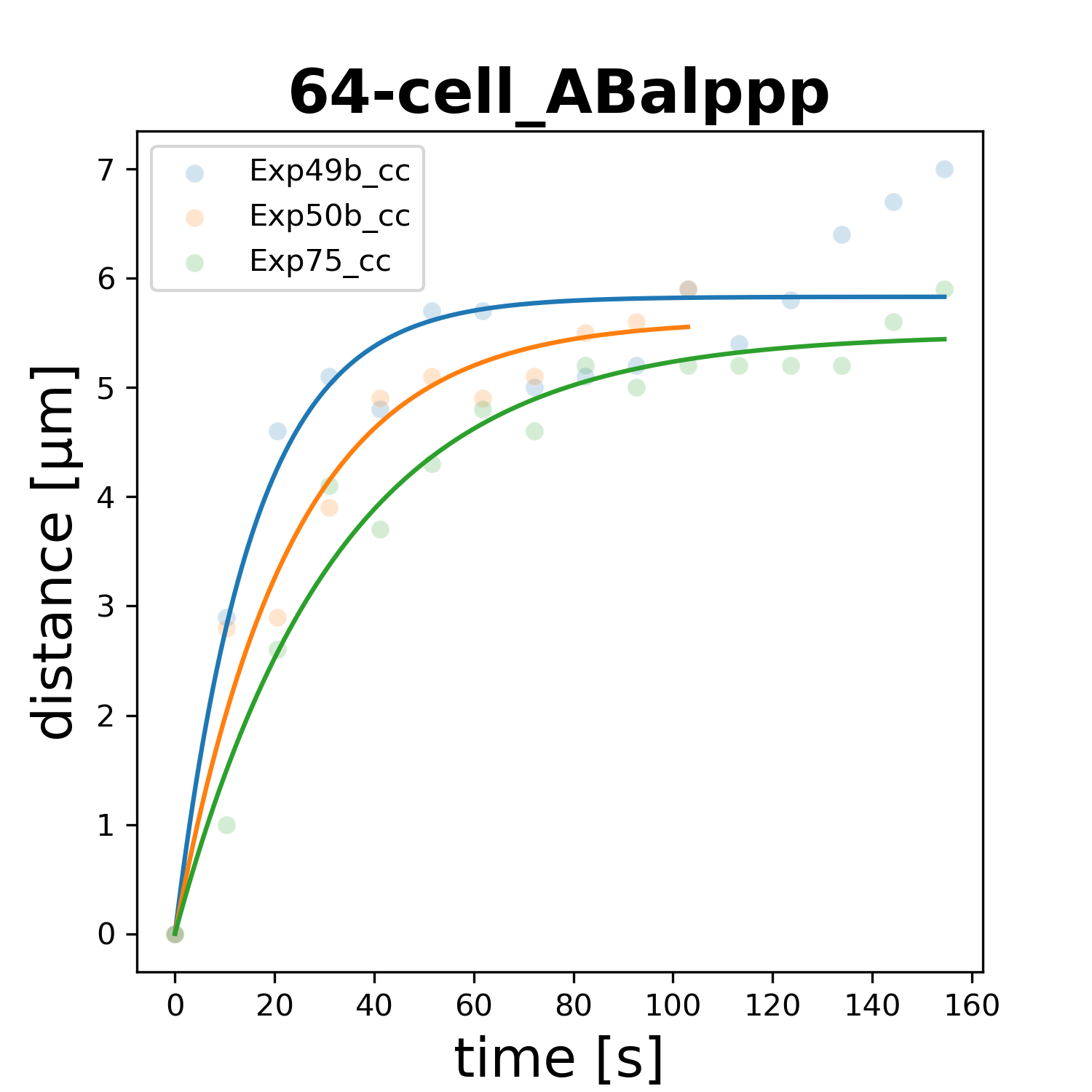

### 64-cell_ABaraaa.png

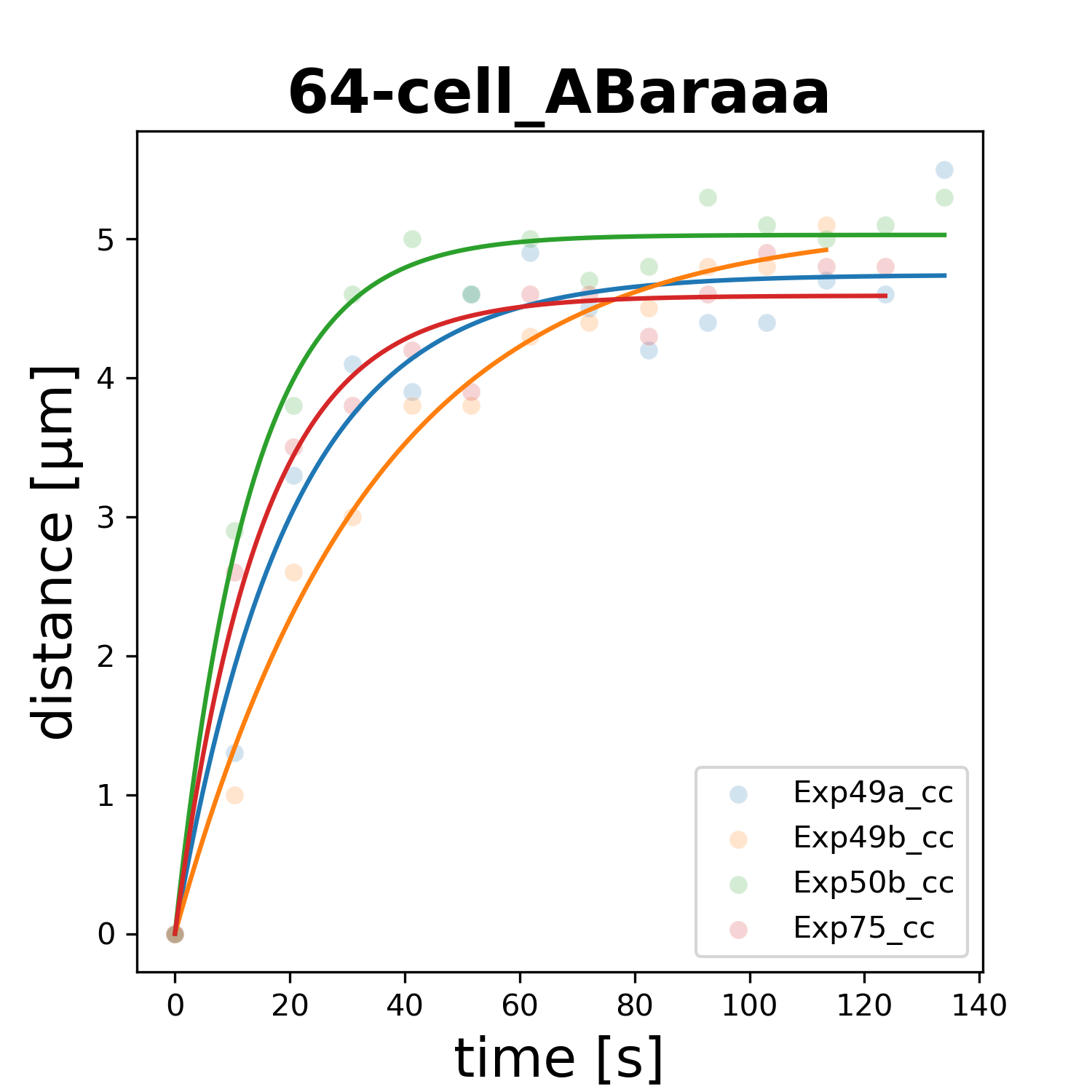

### 64-cell_ABaraap.png

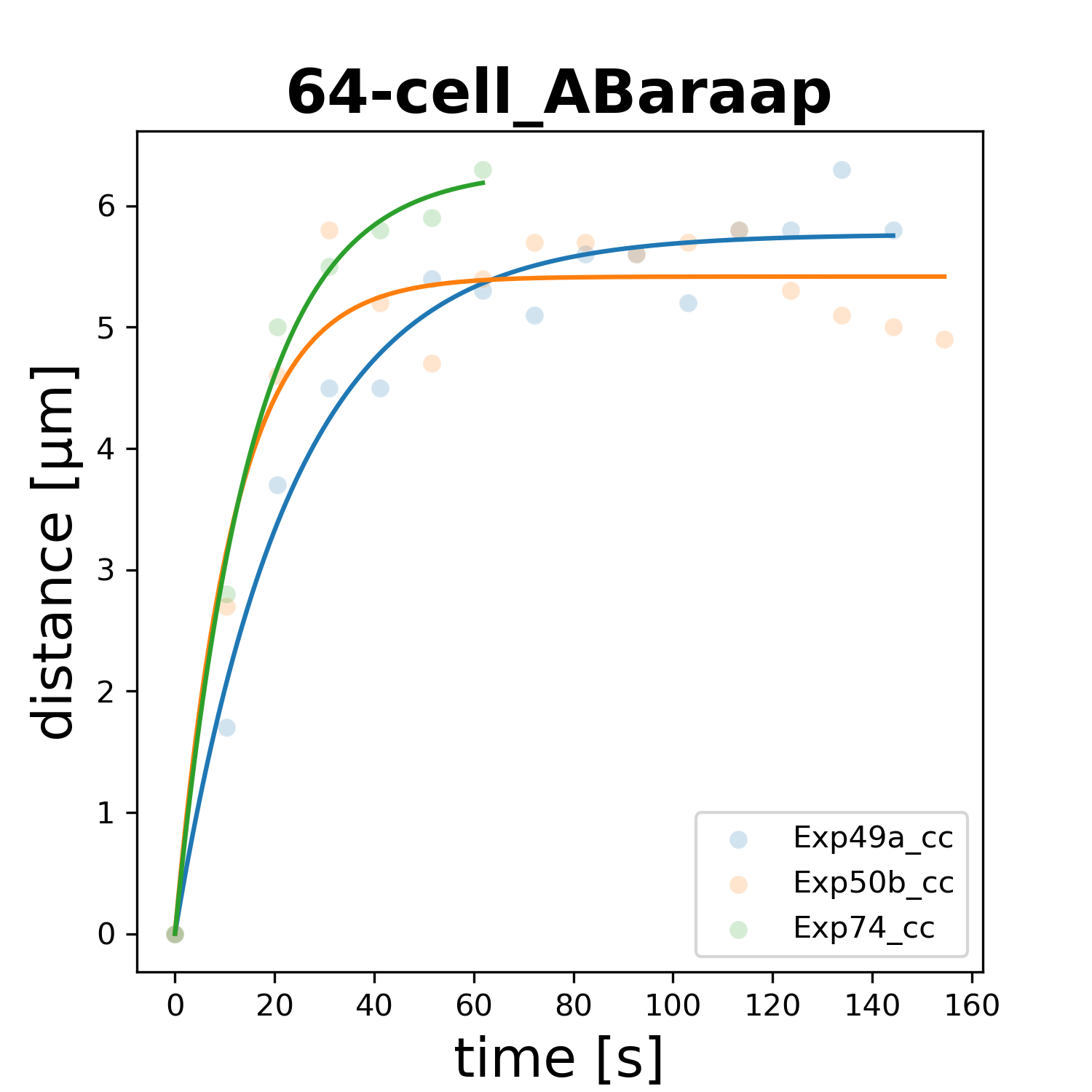

### 64-cell_ABarapa.png

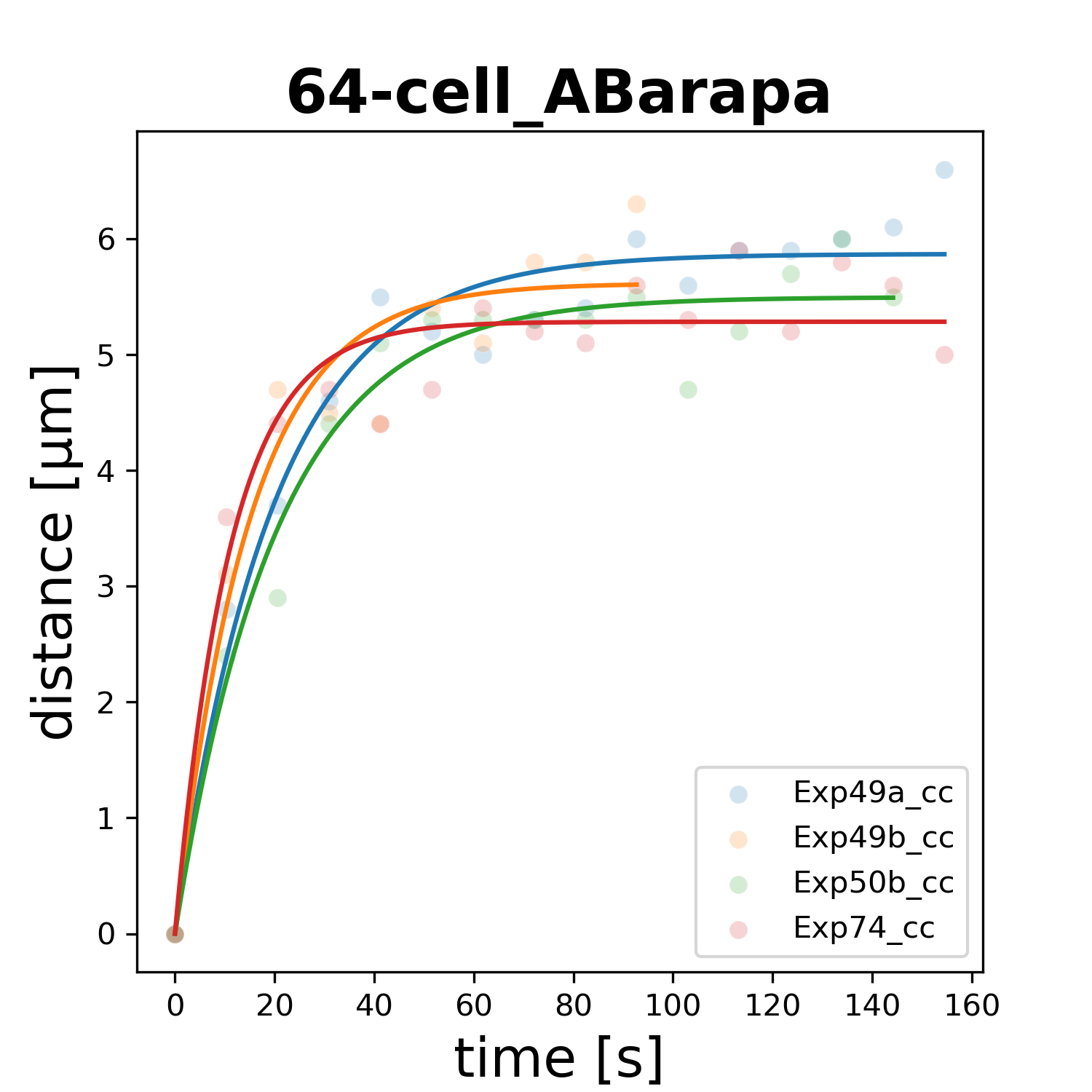

### 64-cell_ABarapp.png

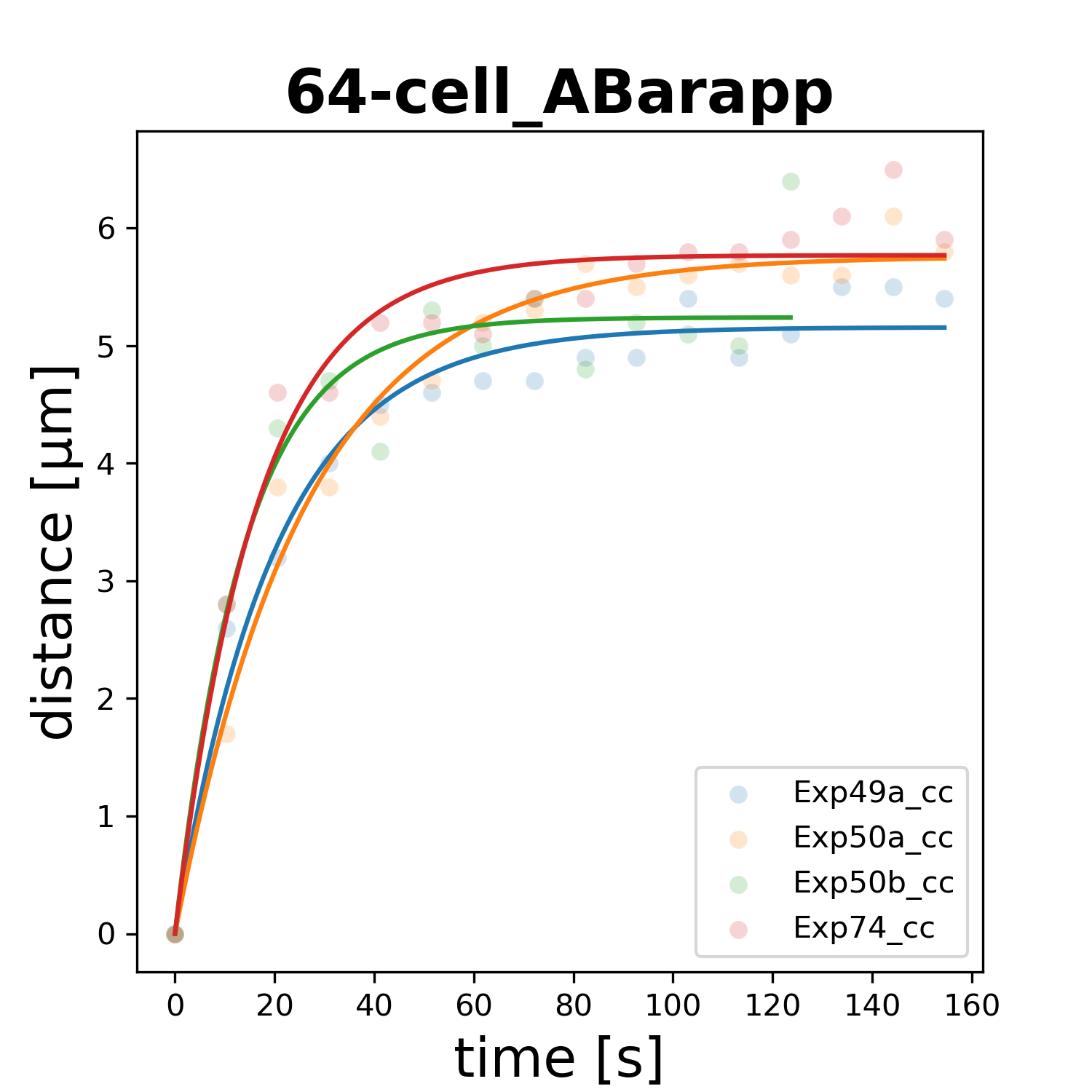

### 64-cell_ABarpaa.png

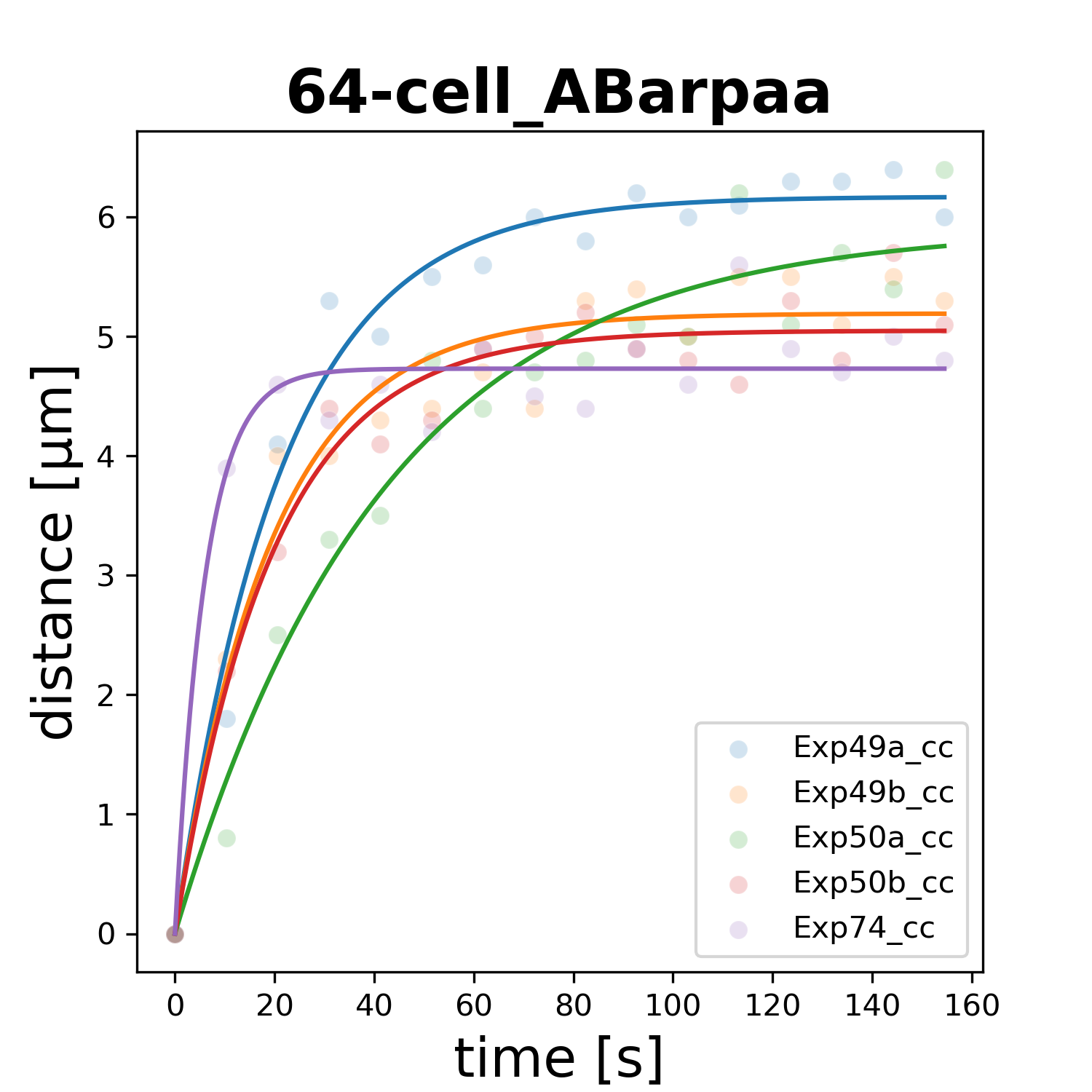

### 64-cell_ABarpap.png

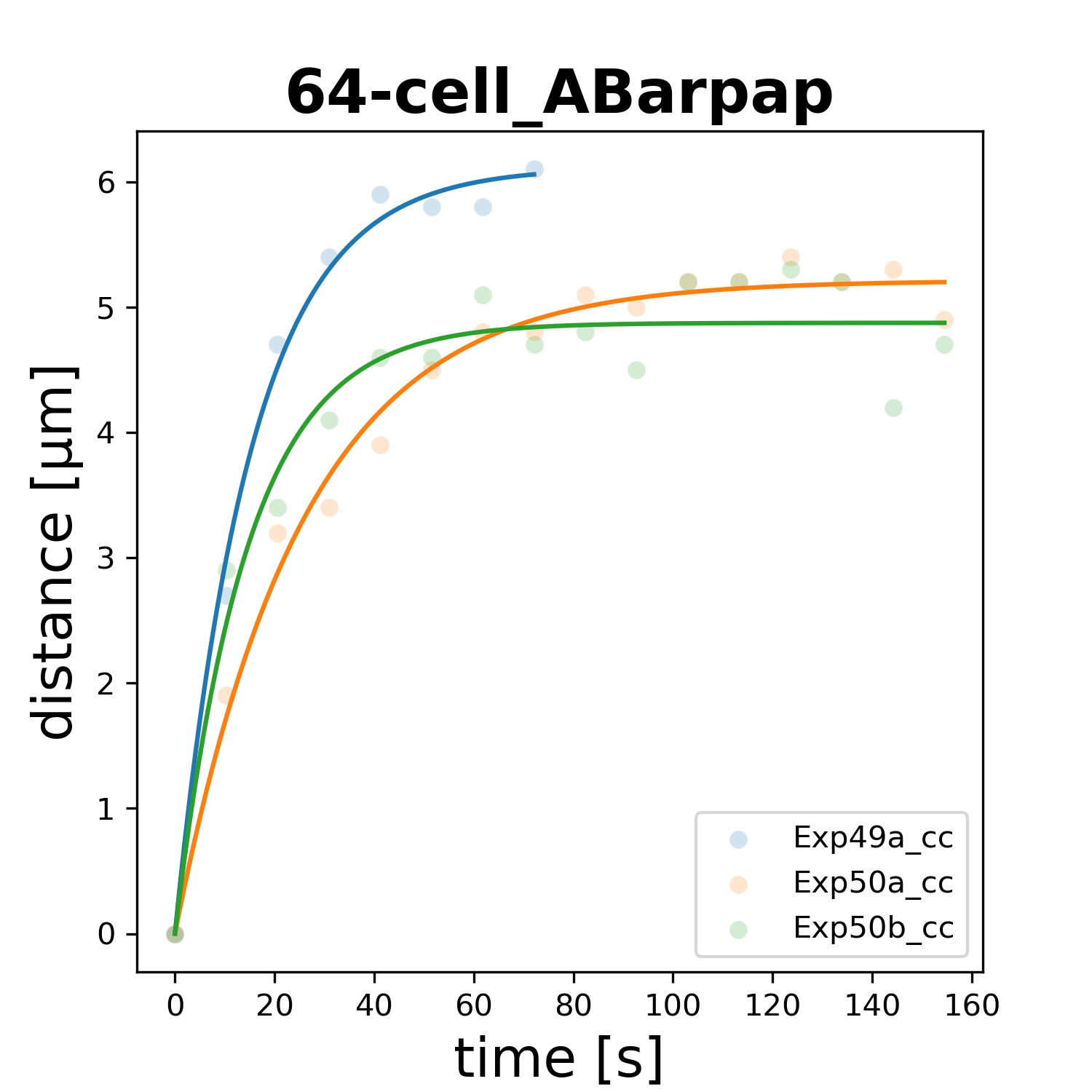

### 64-cell_ABarppa.png

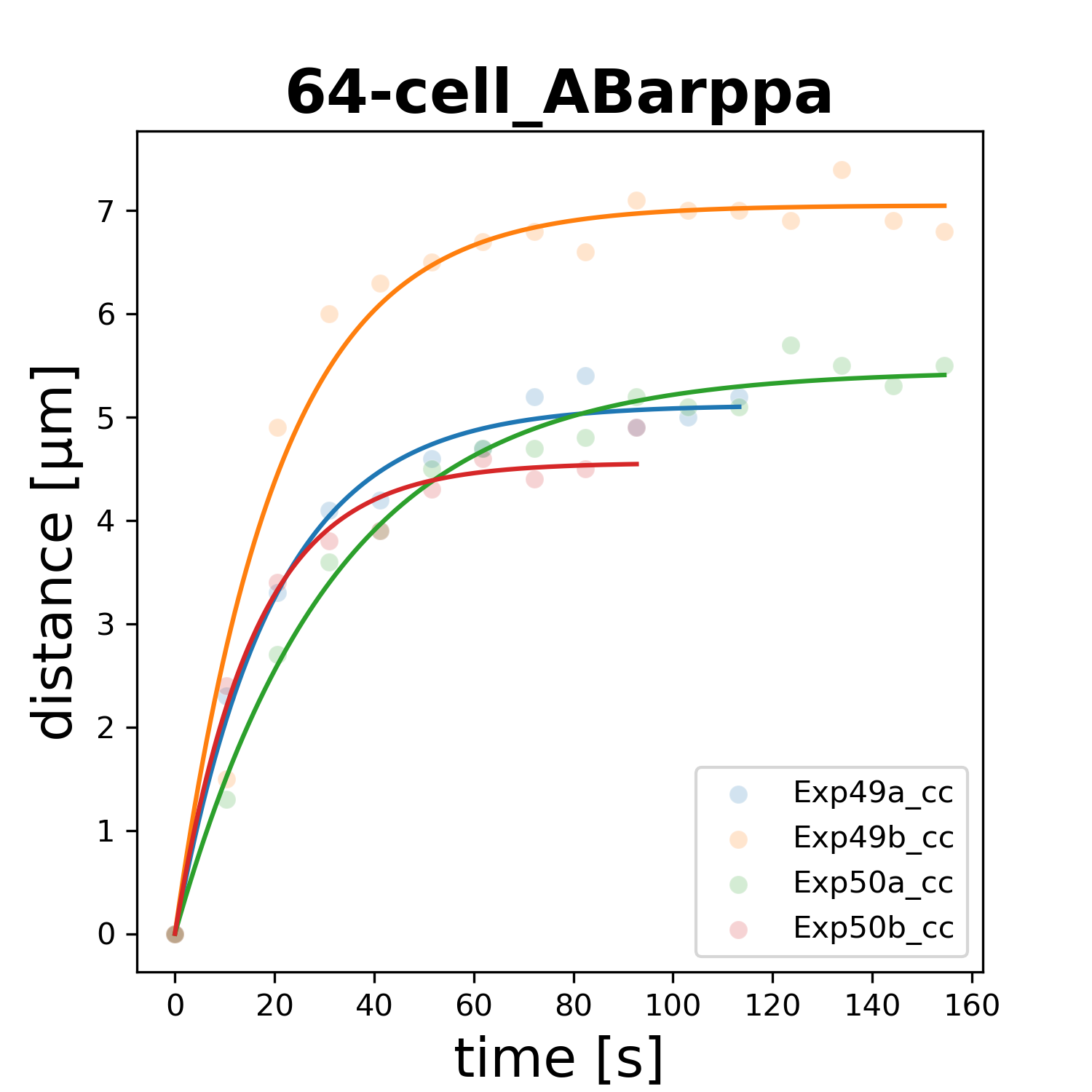

### 64-cell_ABarppp.png

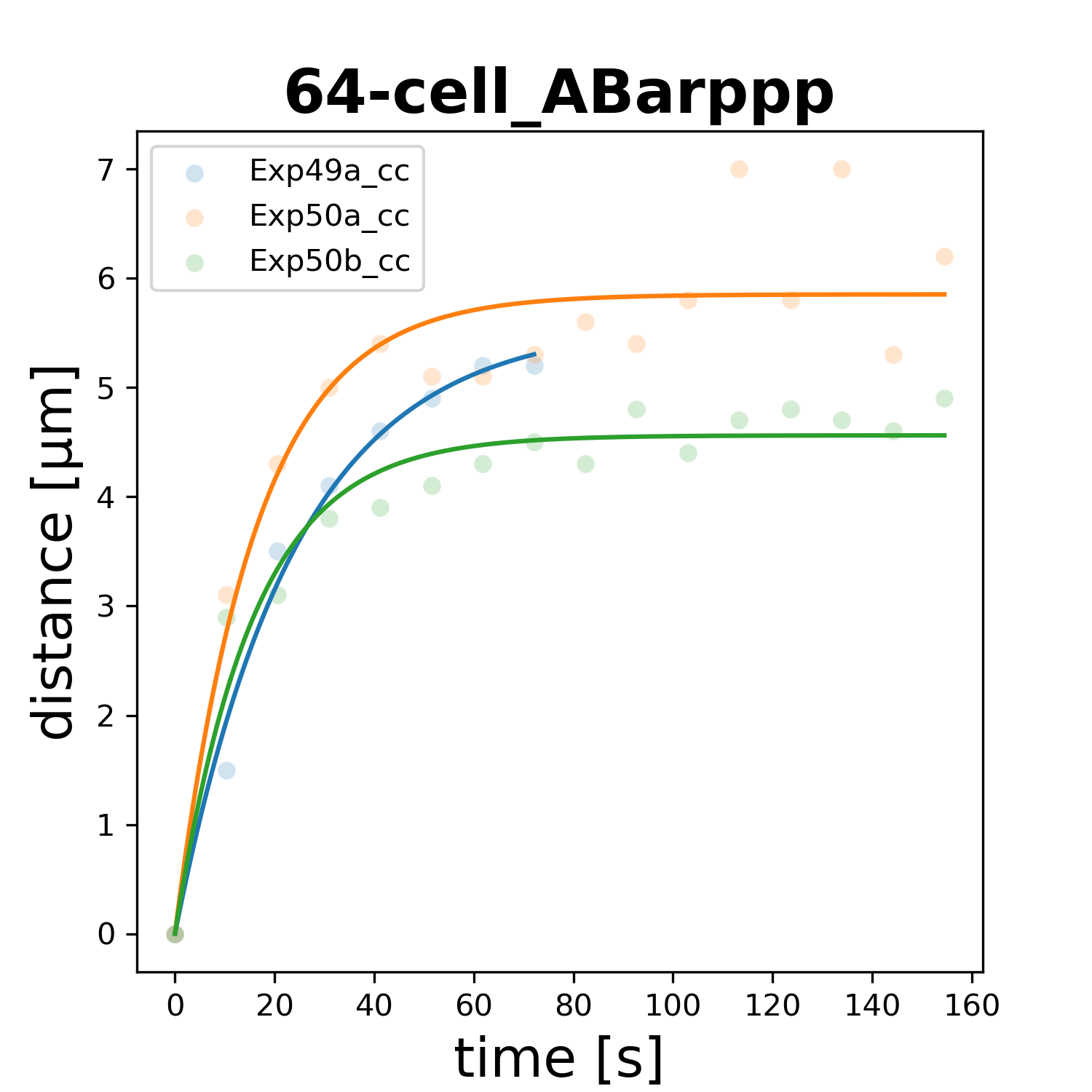

### 64-cell_ABplaaa.png

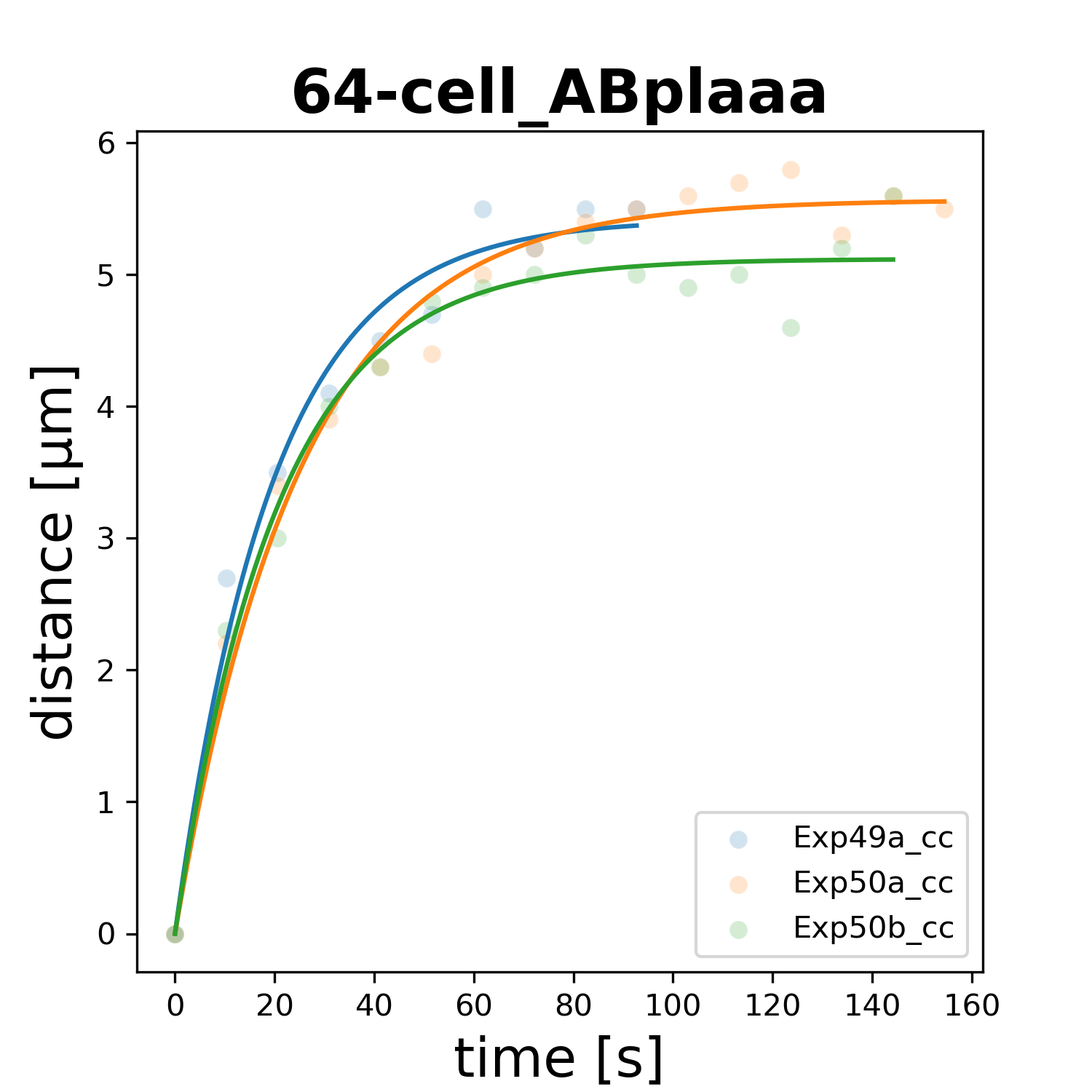

### 64-cell_ABplaap.png

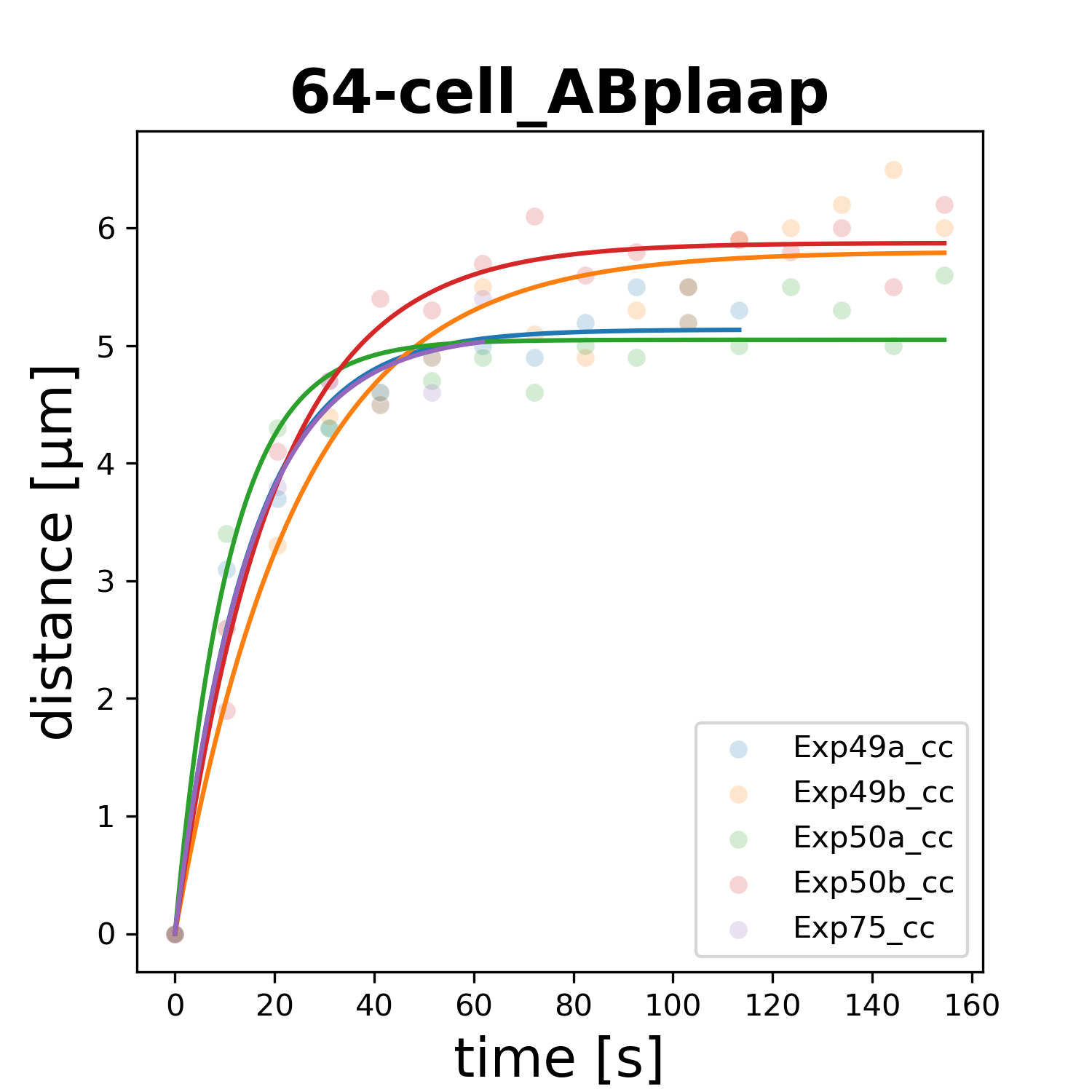

### 64-cell_ABplapa.png

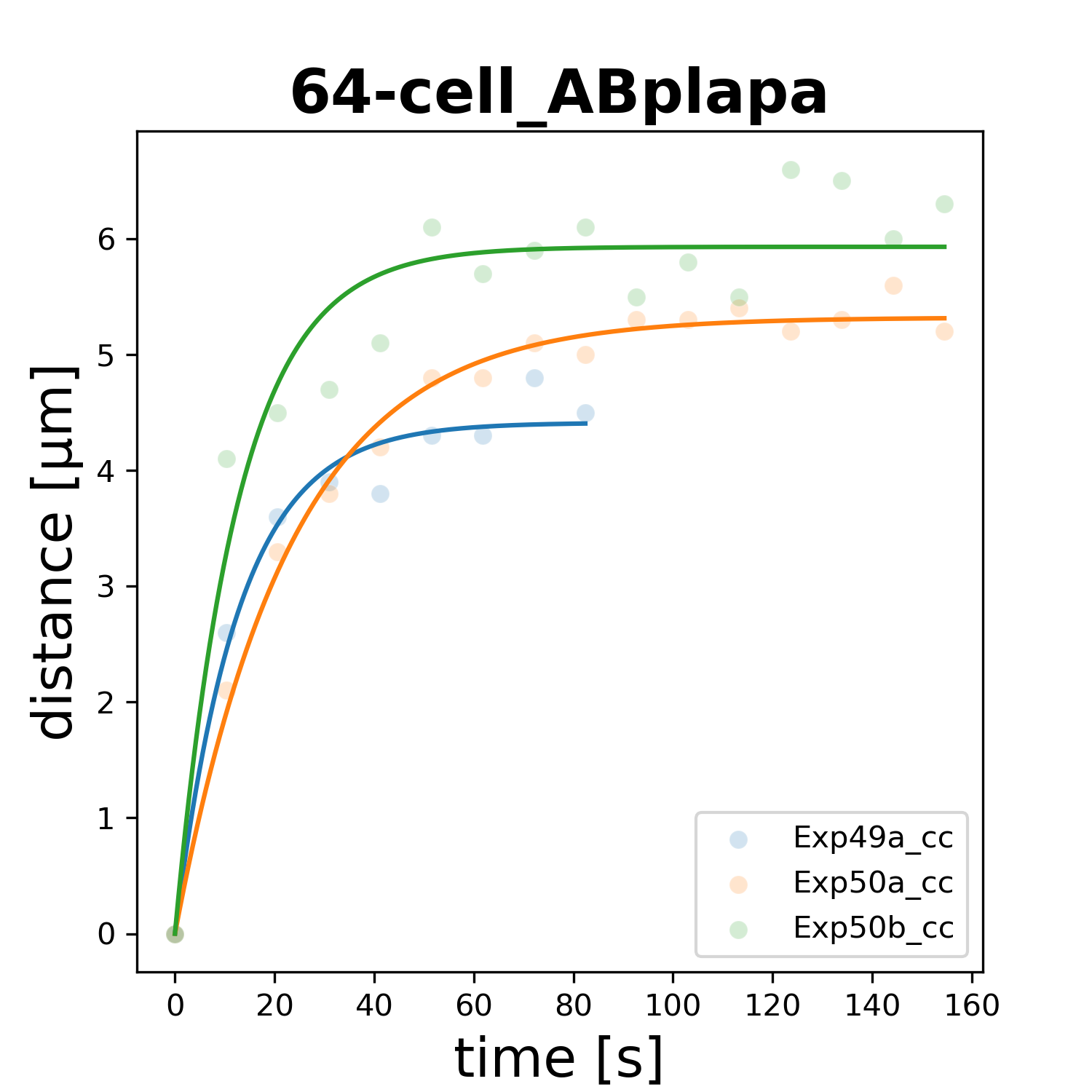

### 64-cell_ABplapp.png

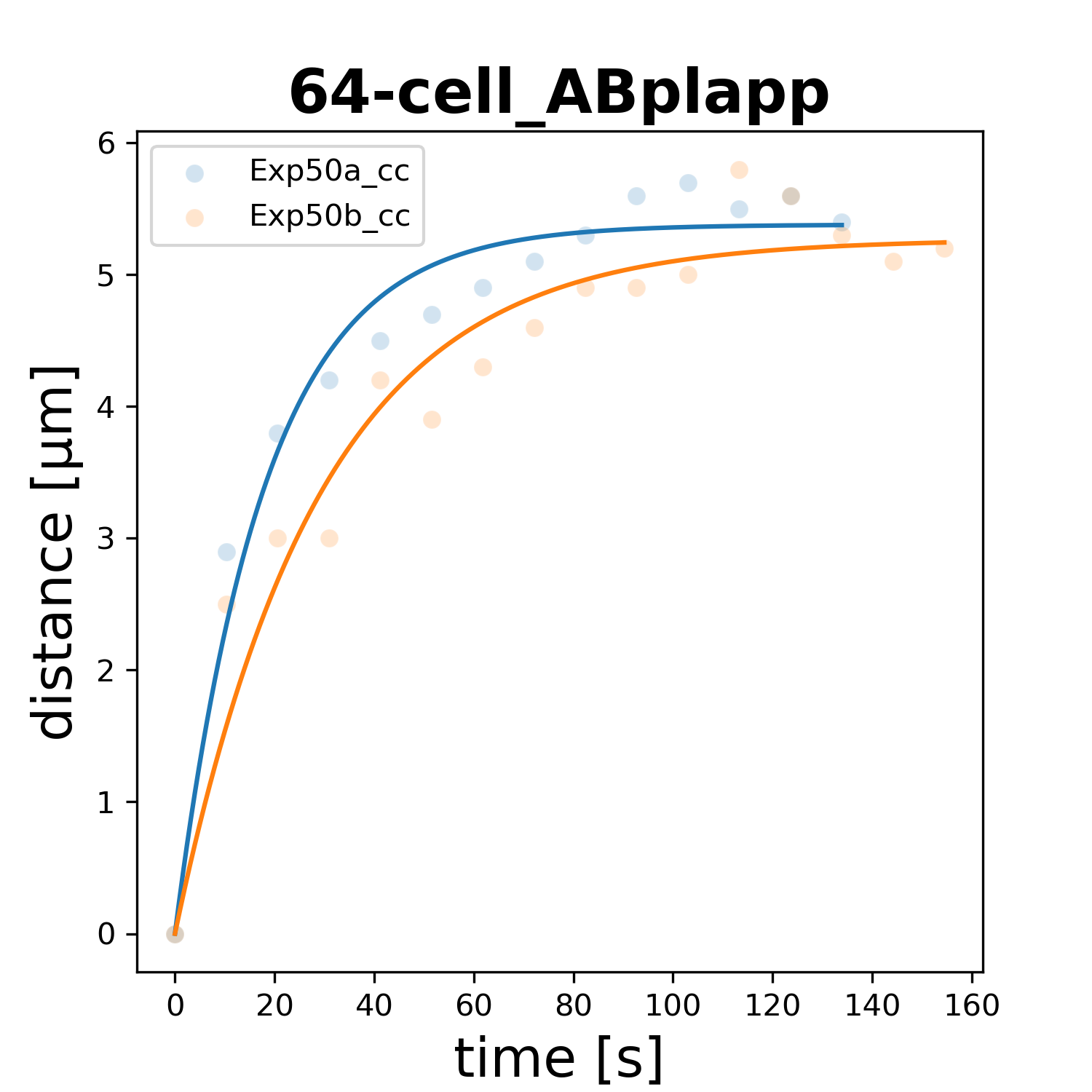

### 64-cell_ABplpaa.png

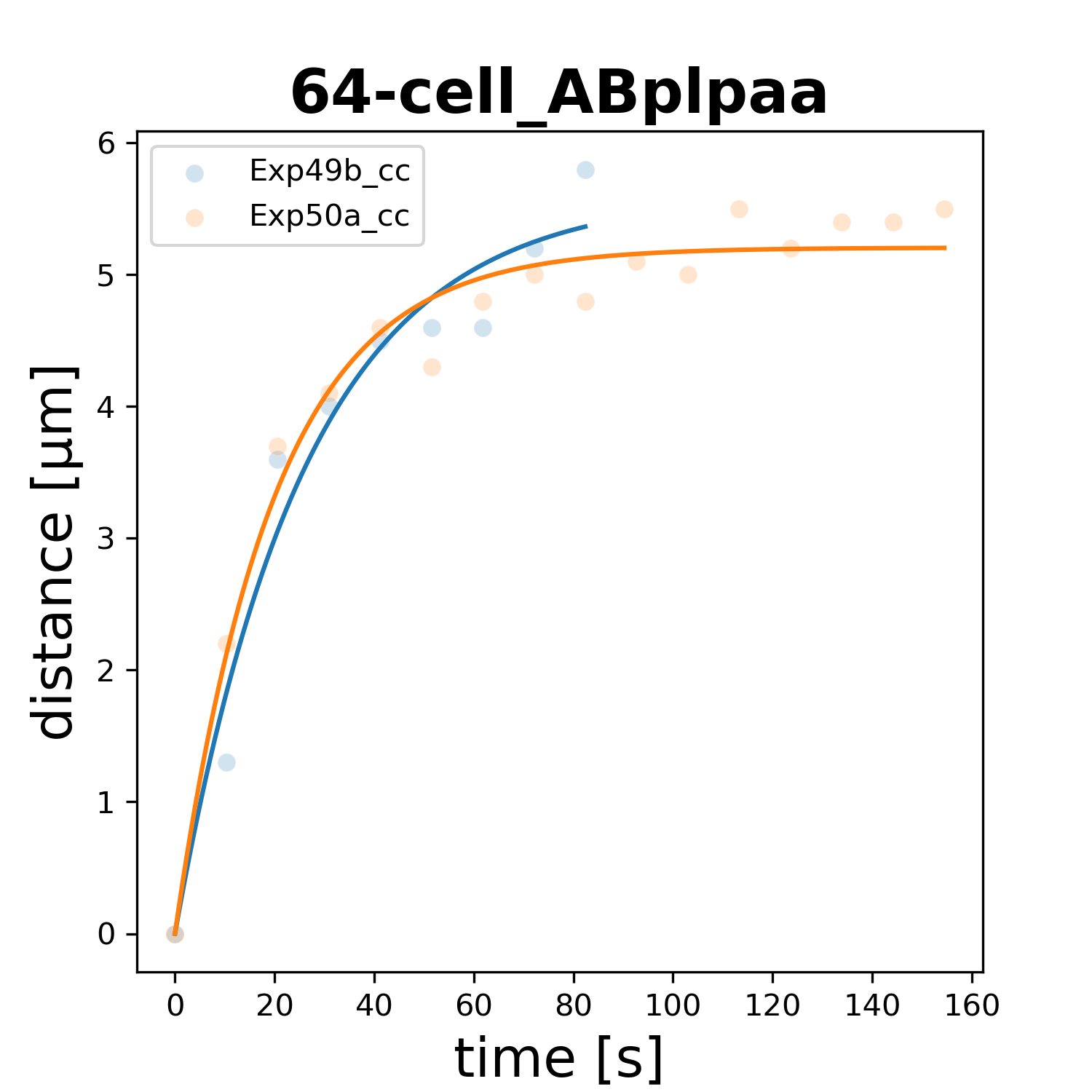

### 64-cell_ABplpap.png

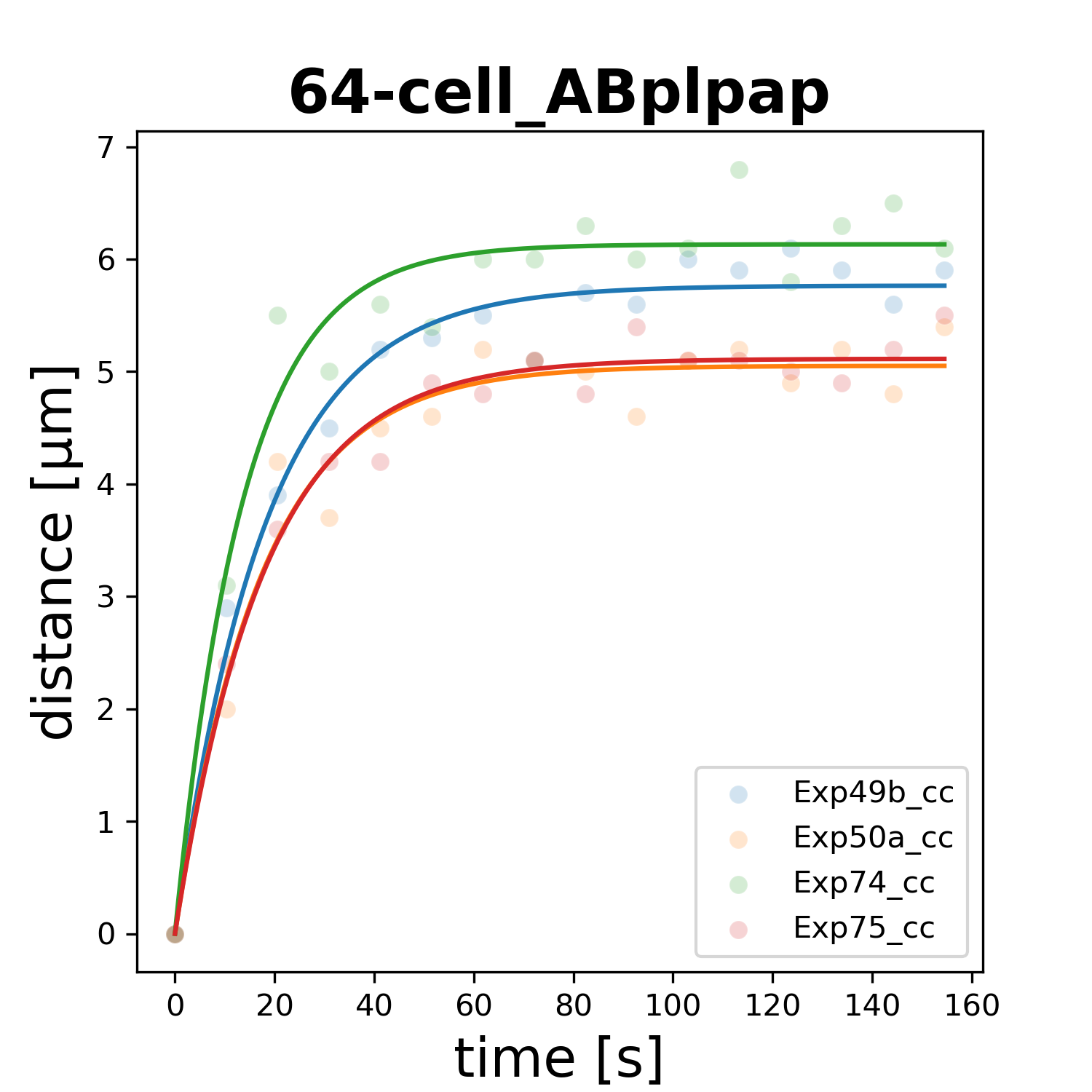

### 64-cell_ABplppa.png

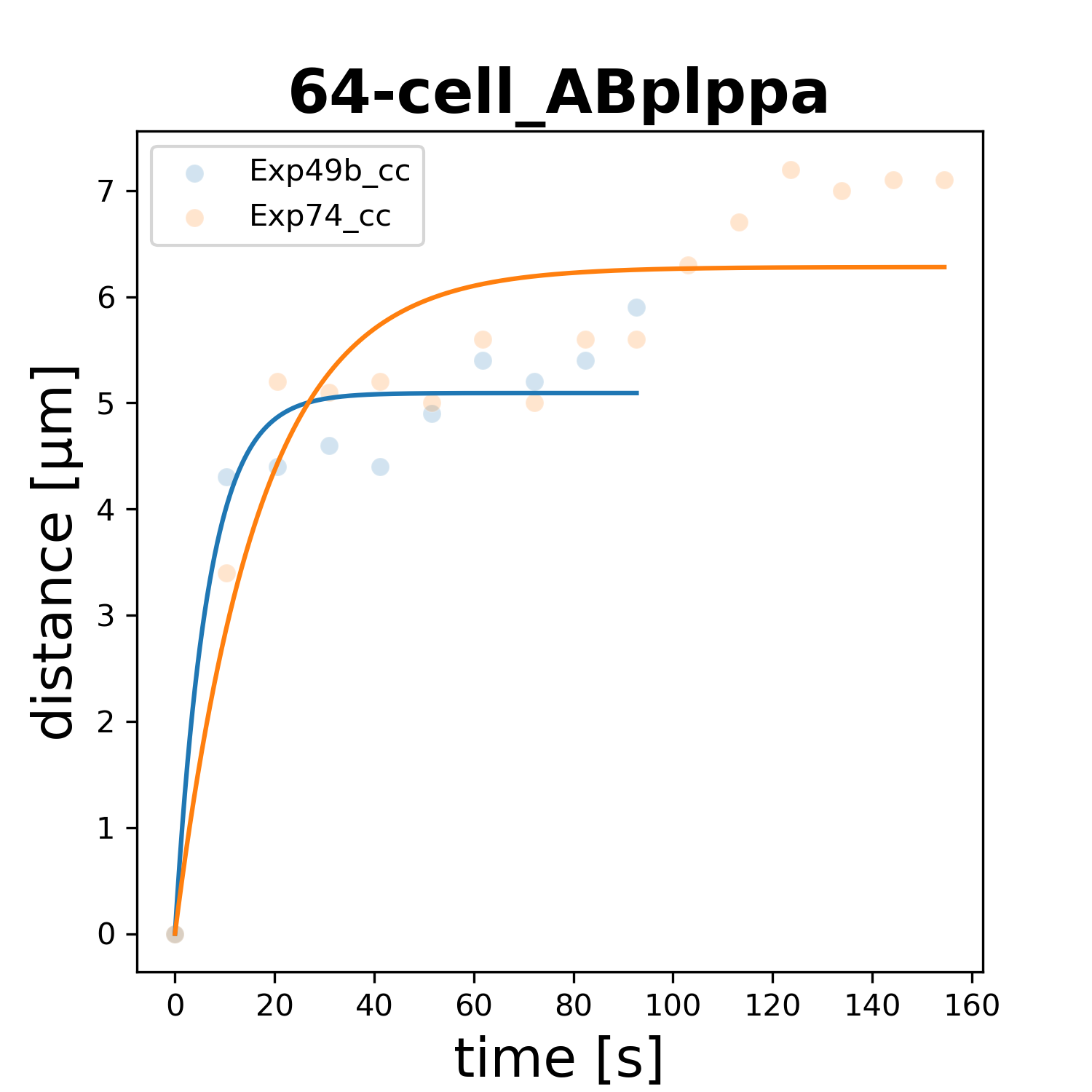

### 64-cell_ABpraaa.png

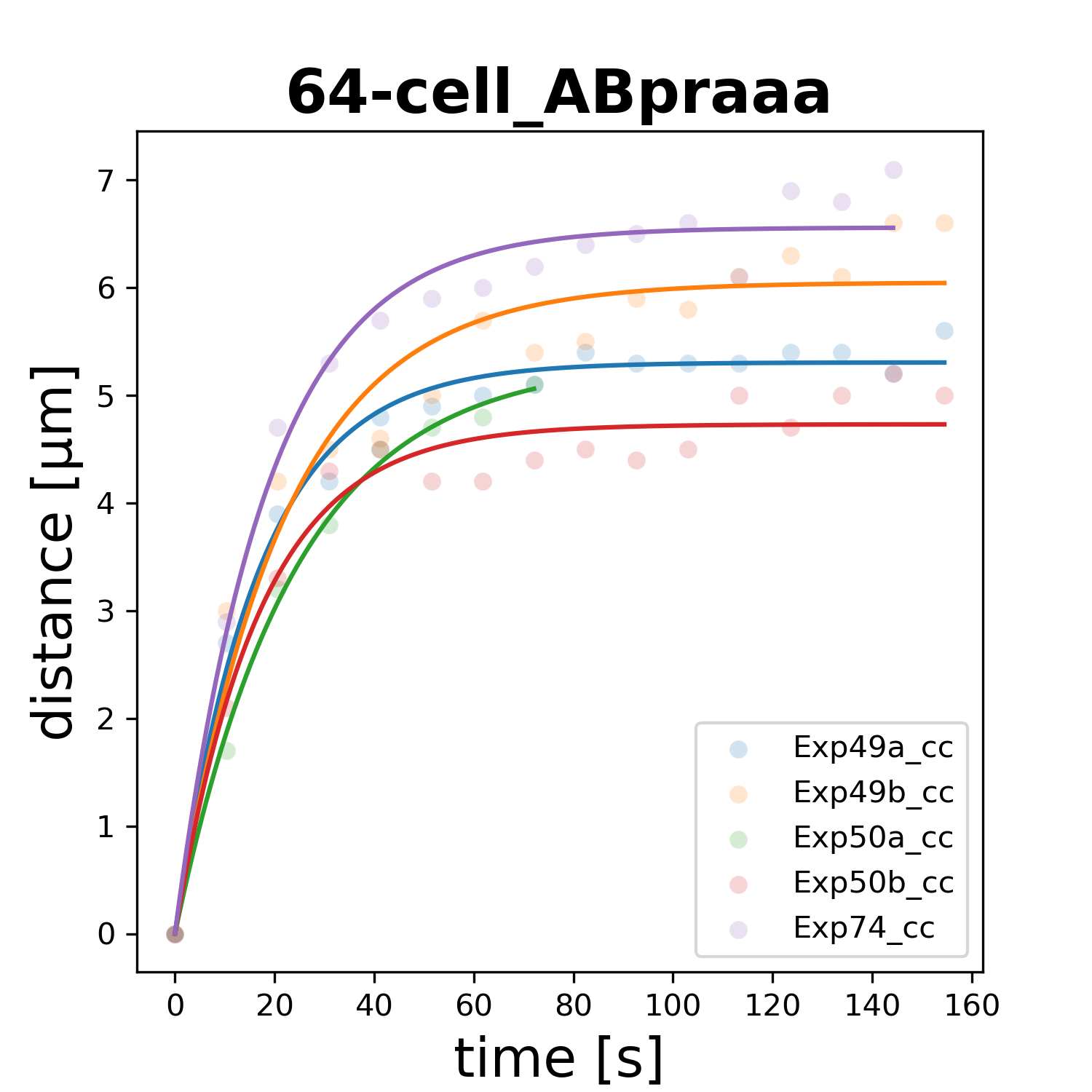

### 64-cell_ABpraap.png

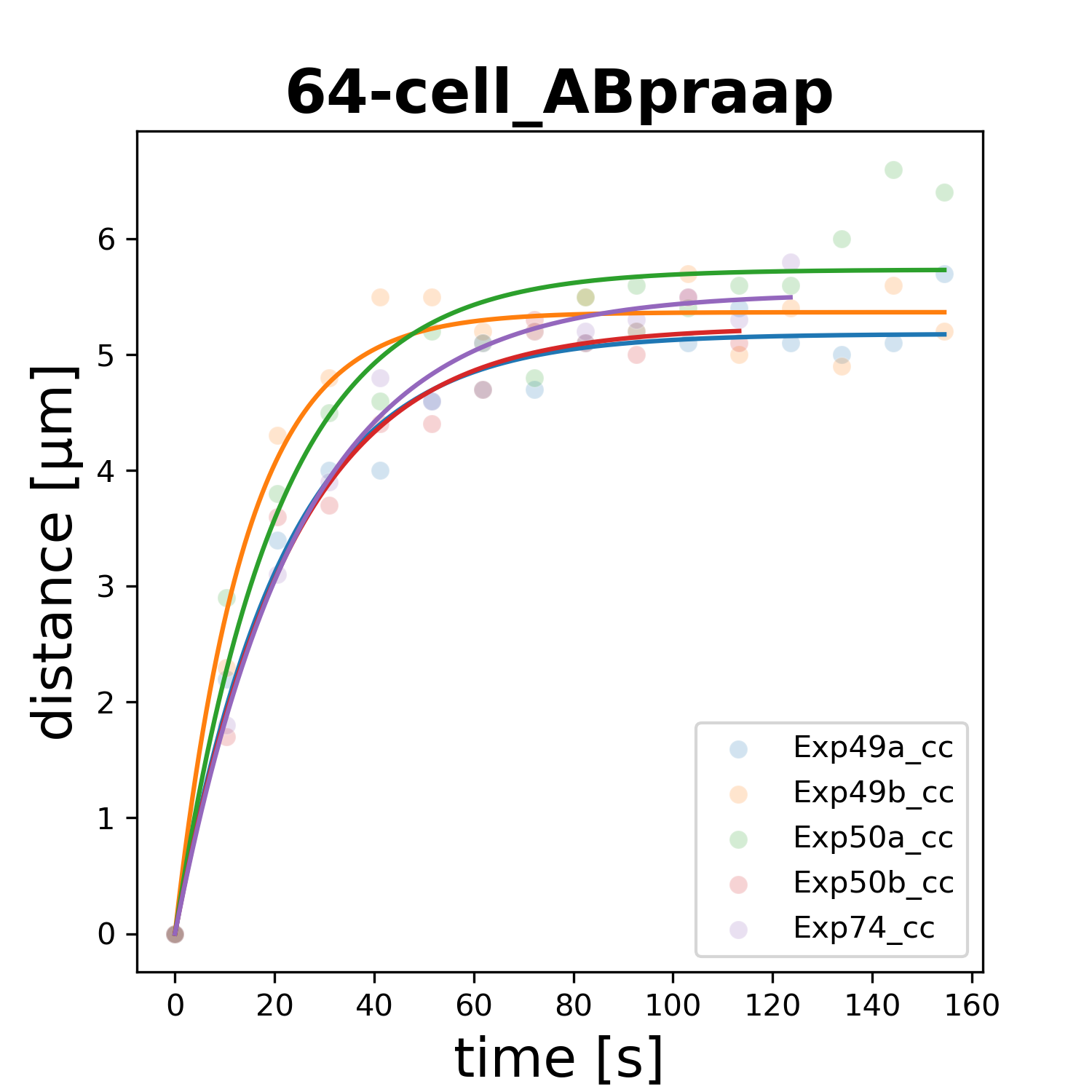

### 64-cell_ABprapa.png

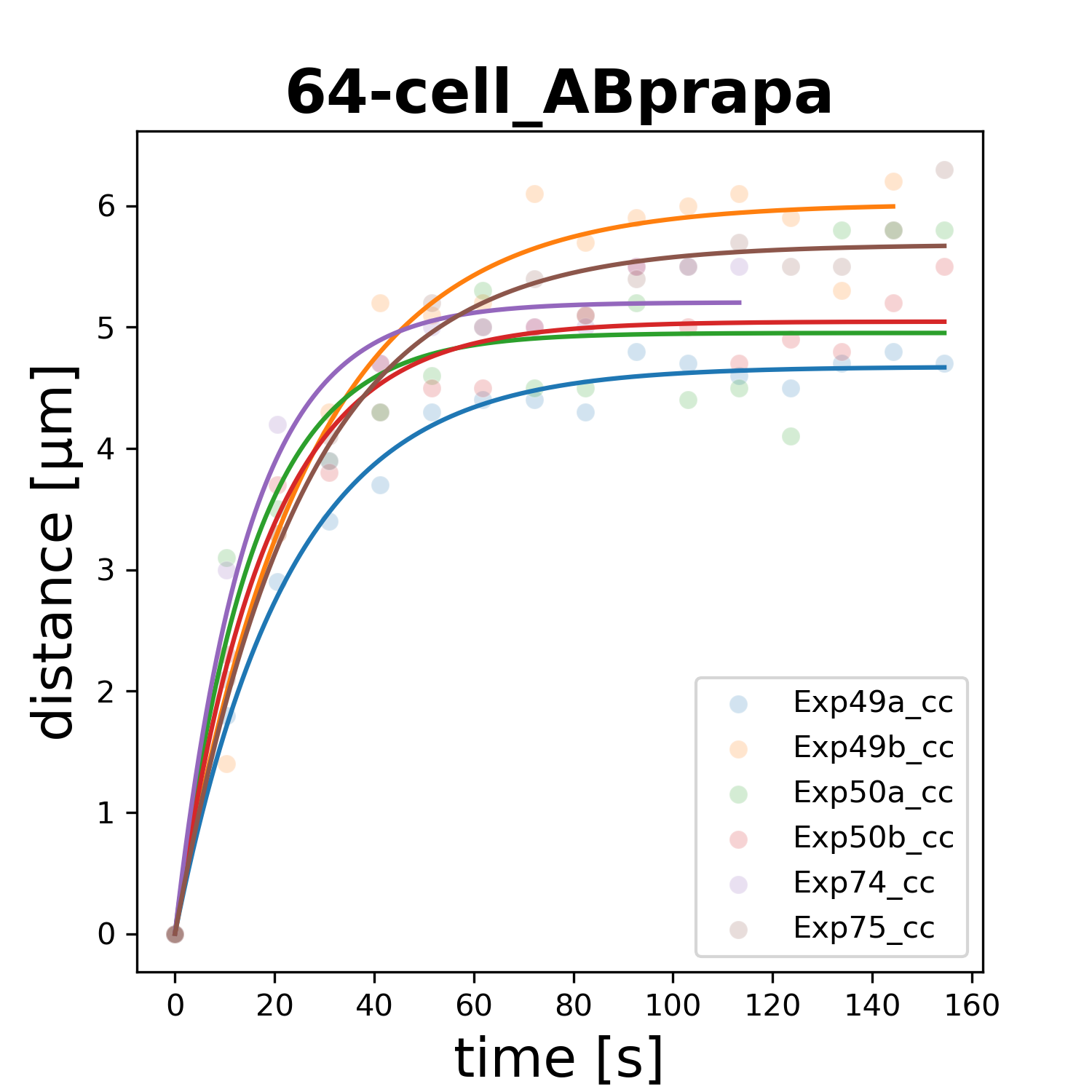

### 64-cell_ABprapp.png

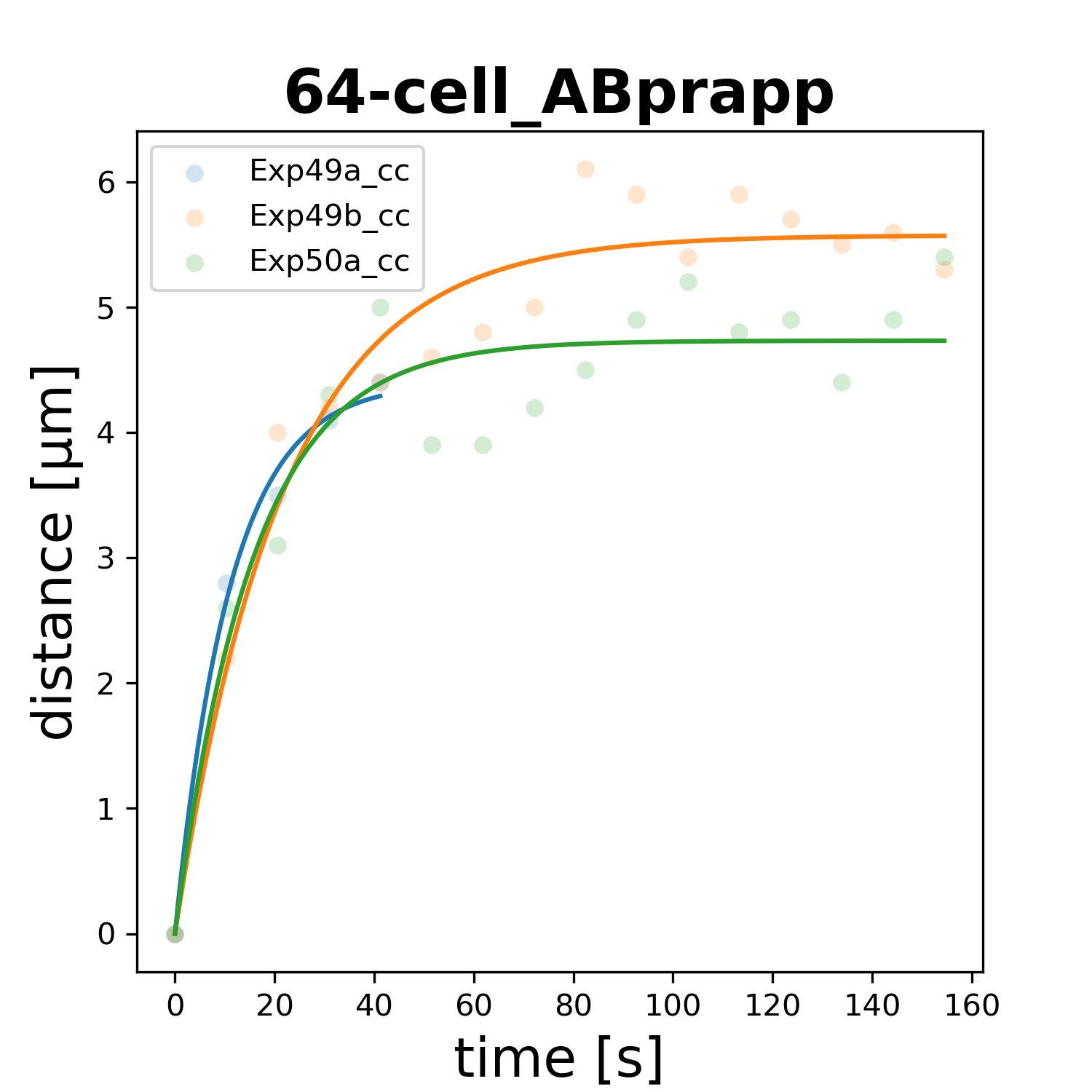

### 64-cell_ABprpaa.png

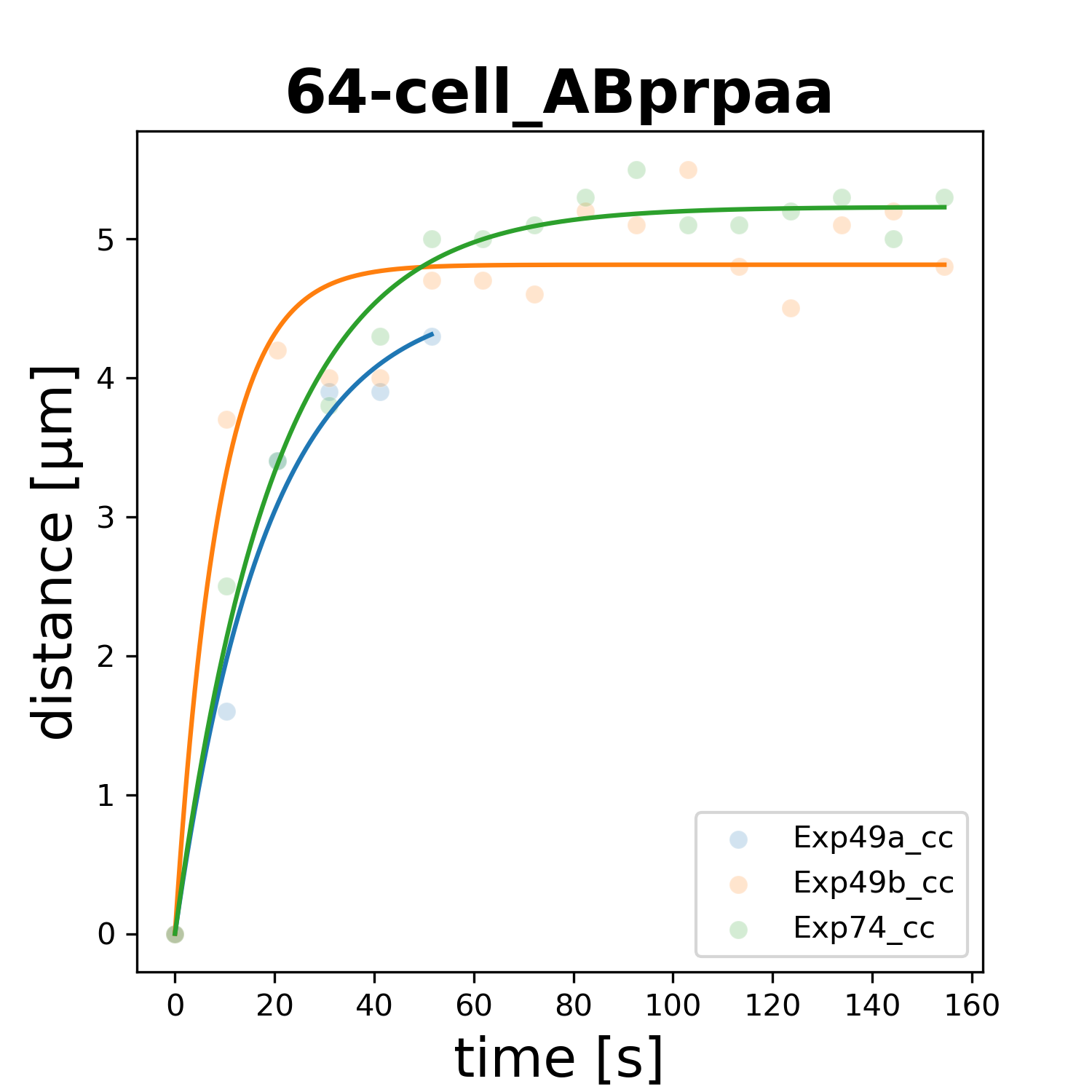

### 64-cell_ABprpap.png

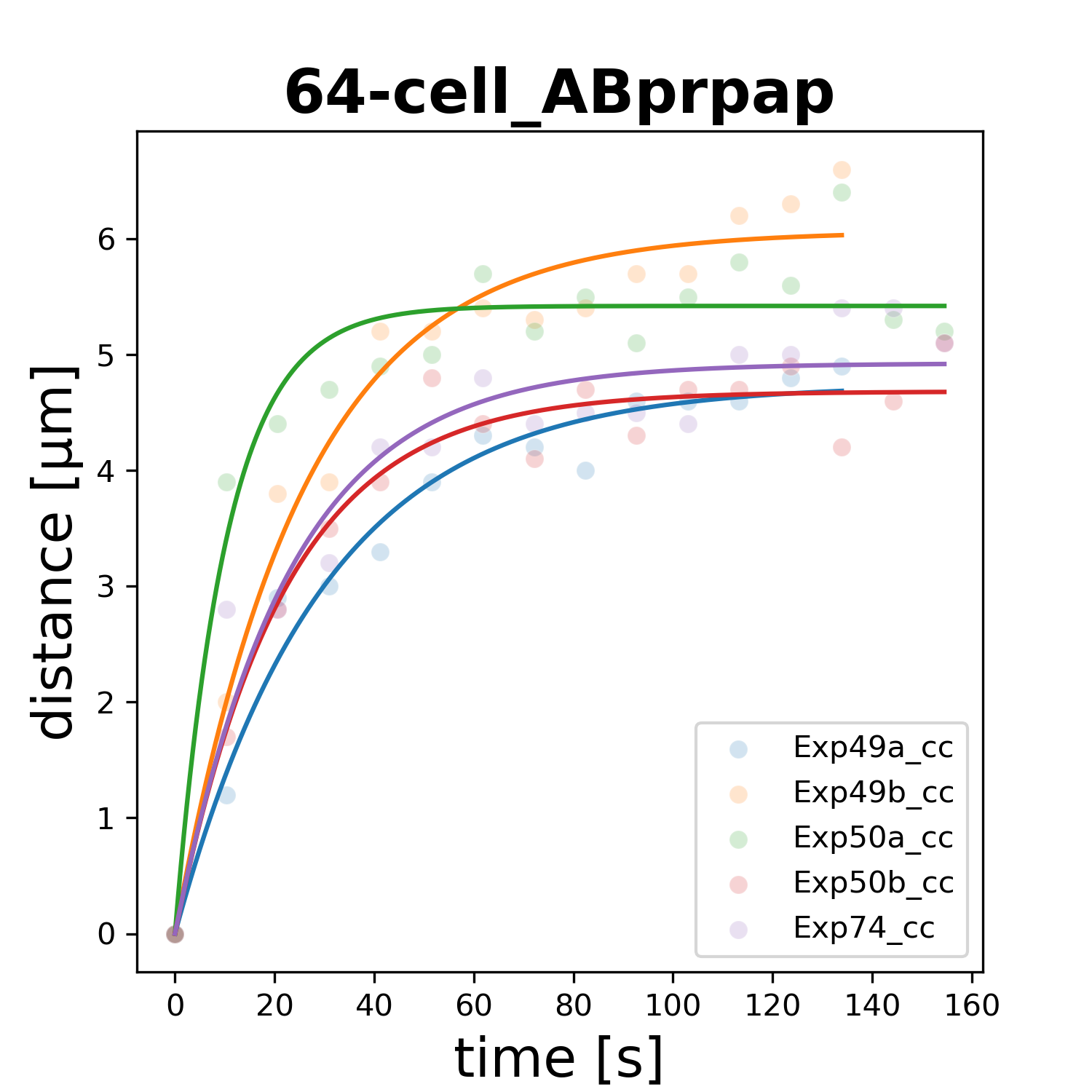

### 64-cell_ABprppa.png

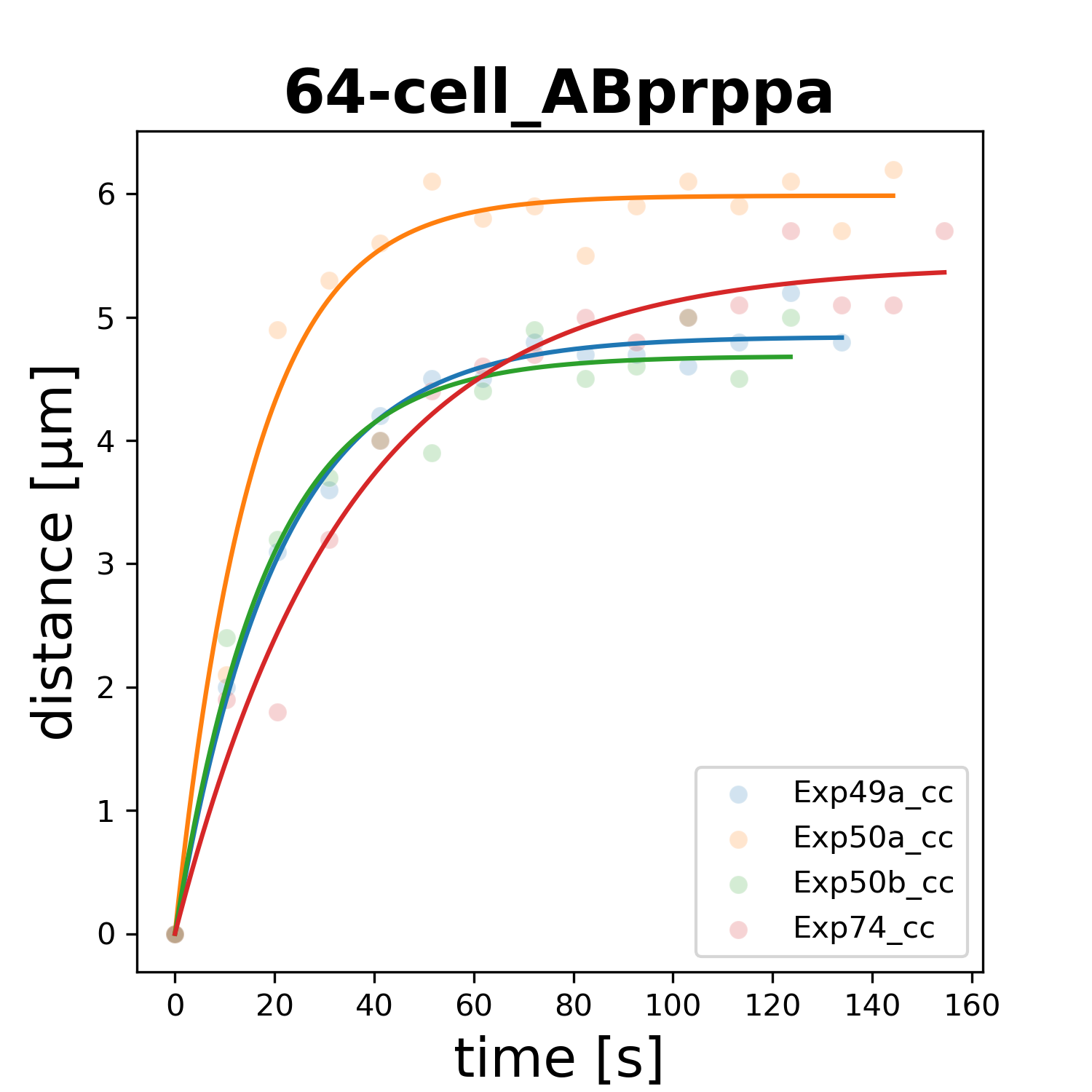

### 64-cell_ABprppp.png

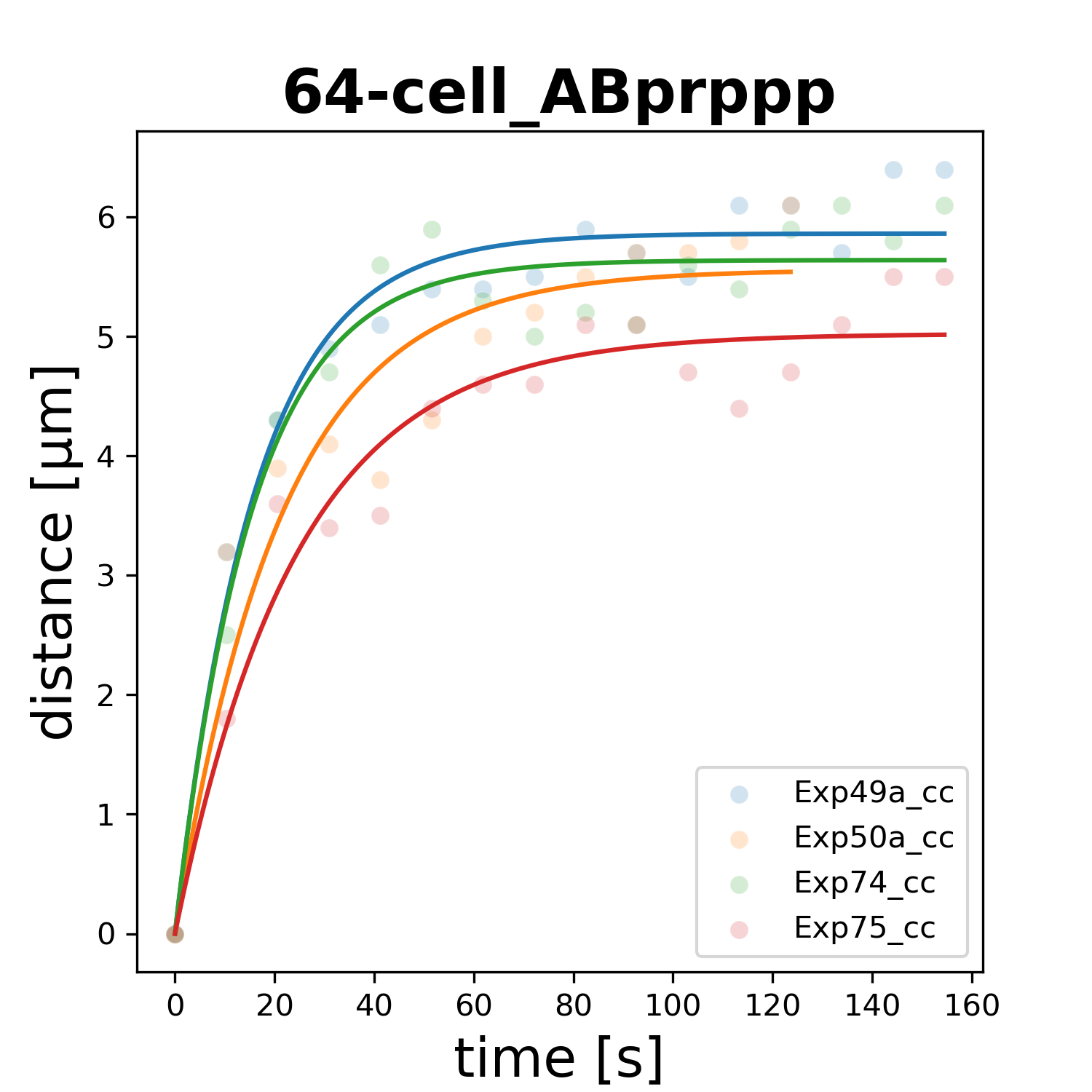

### 64-cell_MSaaa.png

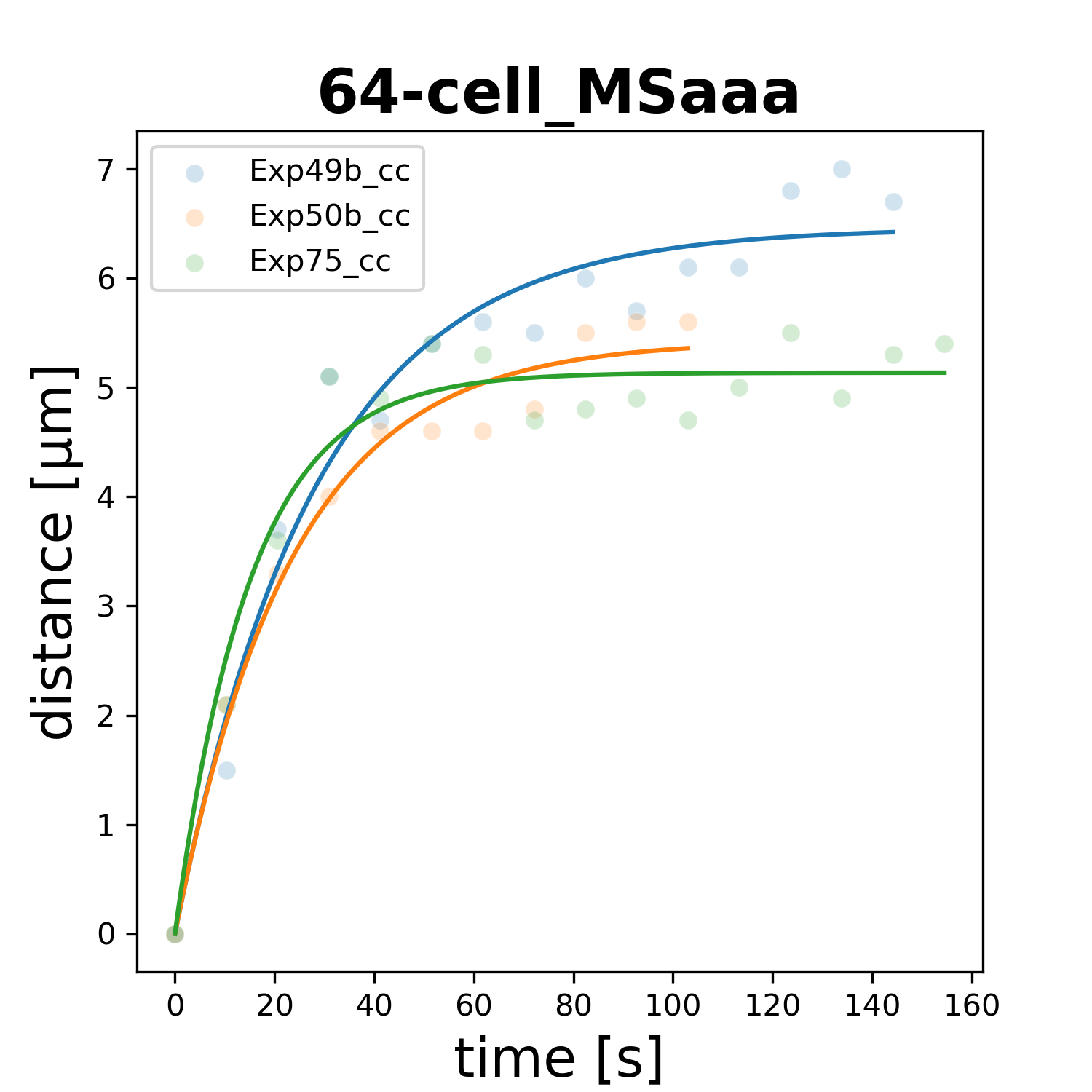

### 64-cell_MSaap.png

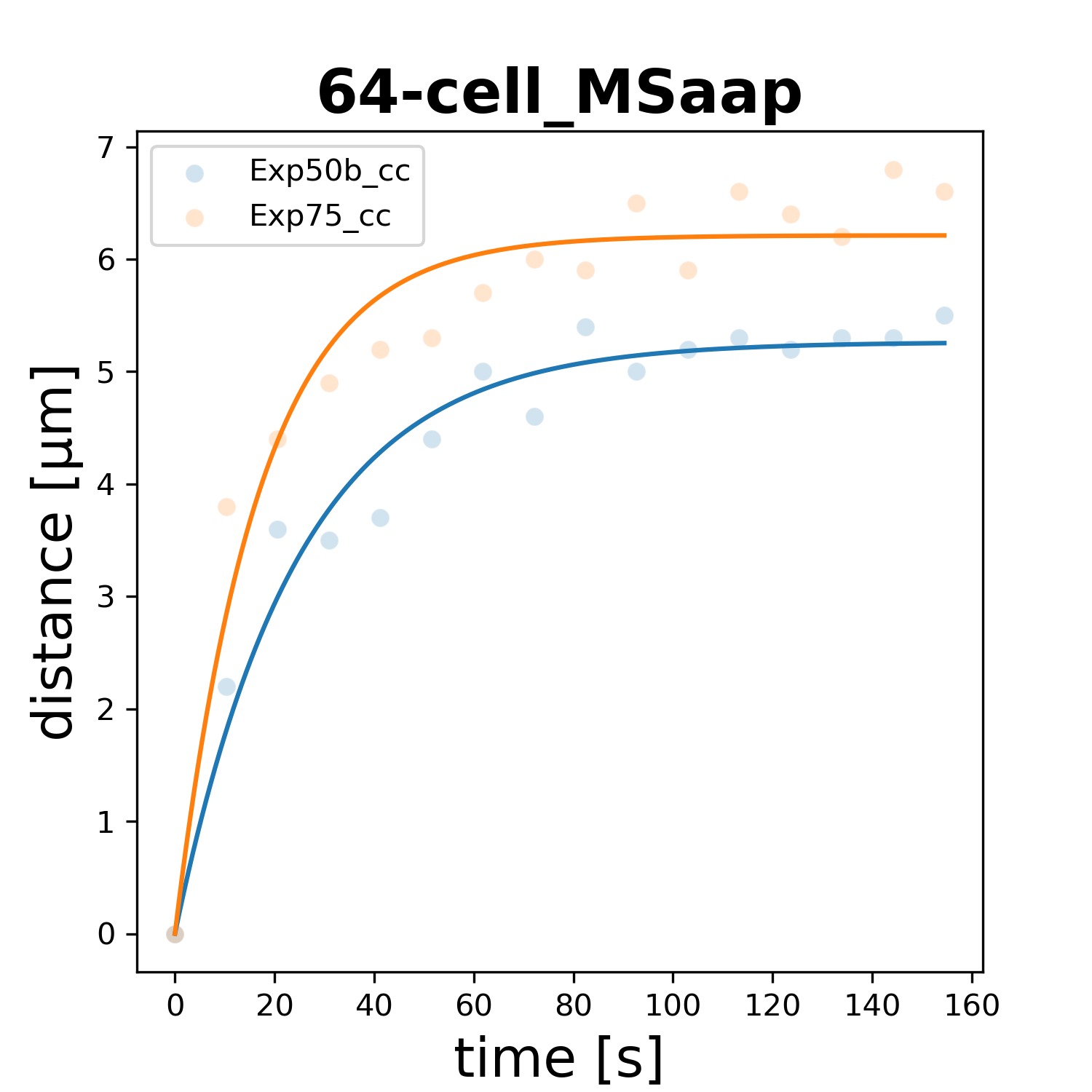

### 64-cell_MSapa.png

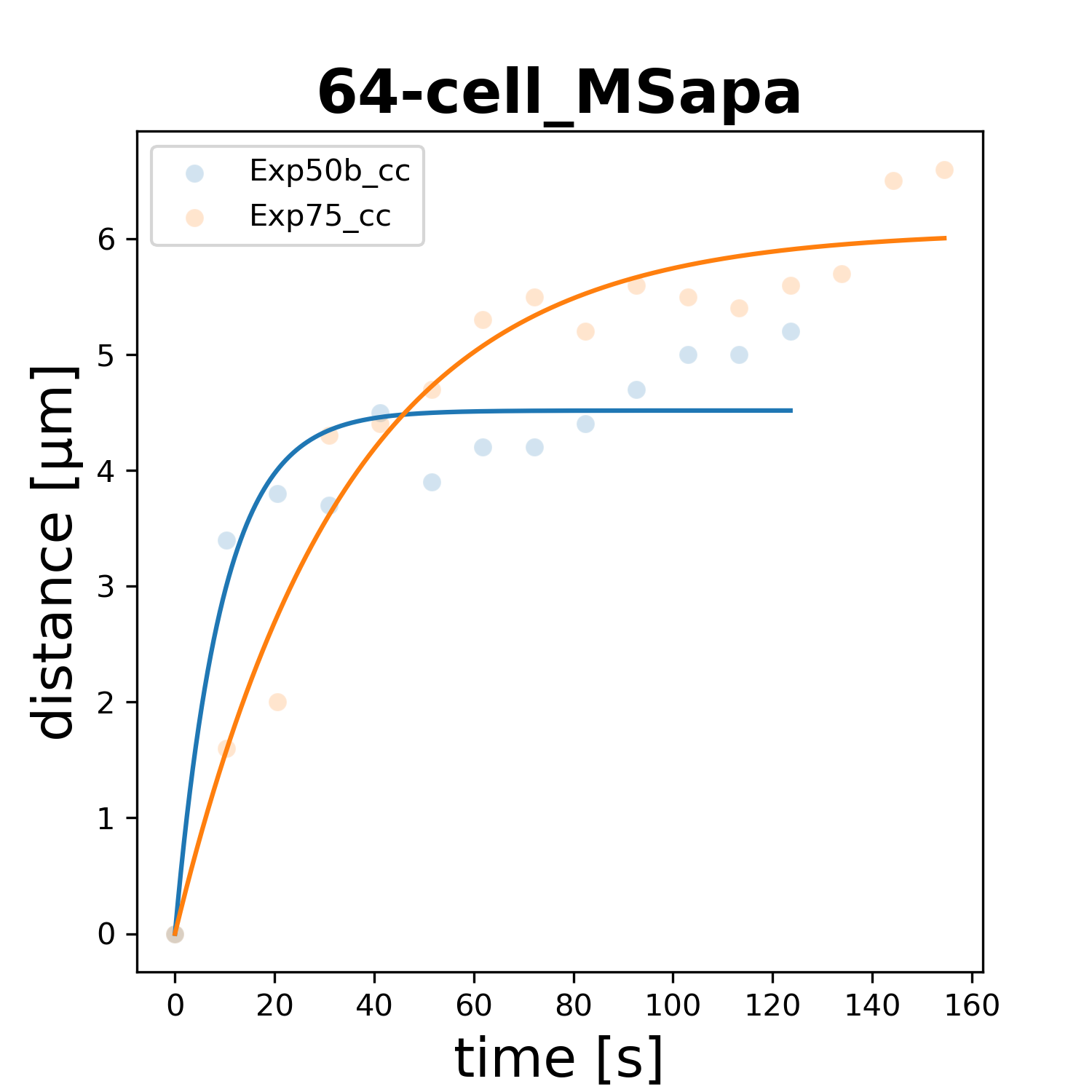

### 64-cell_MSpaa.png

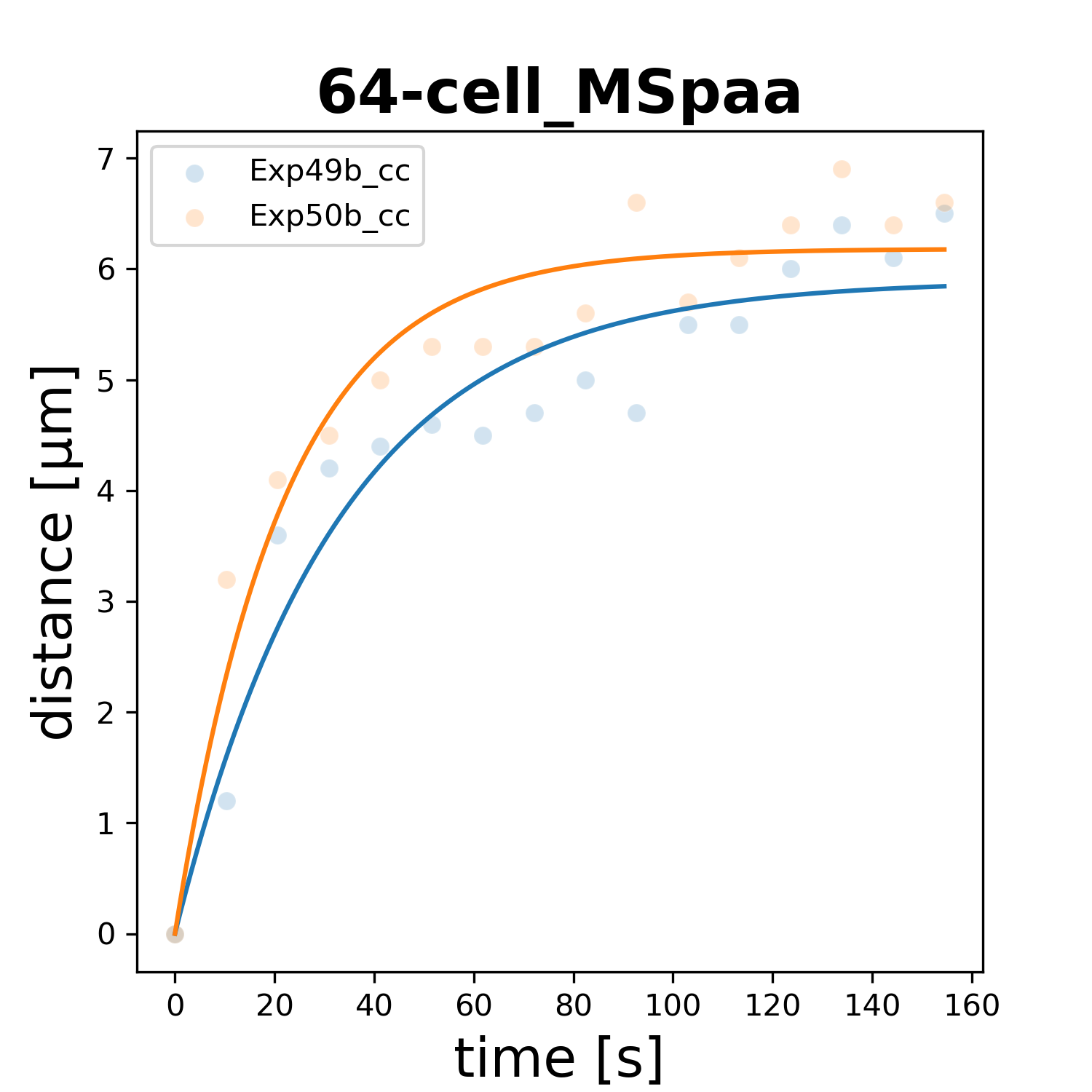

### 64-cell_MSpap.png

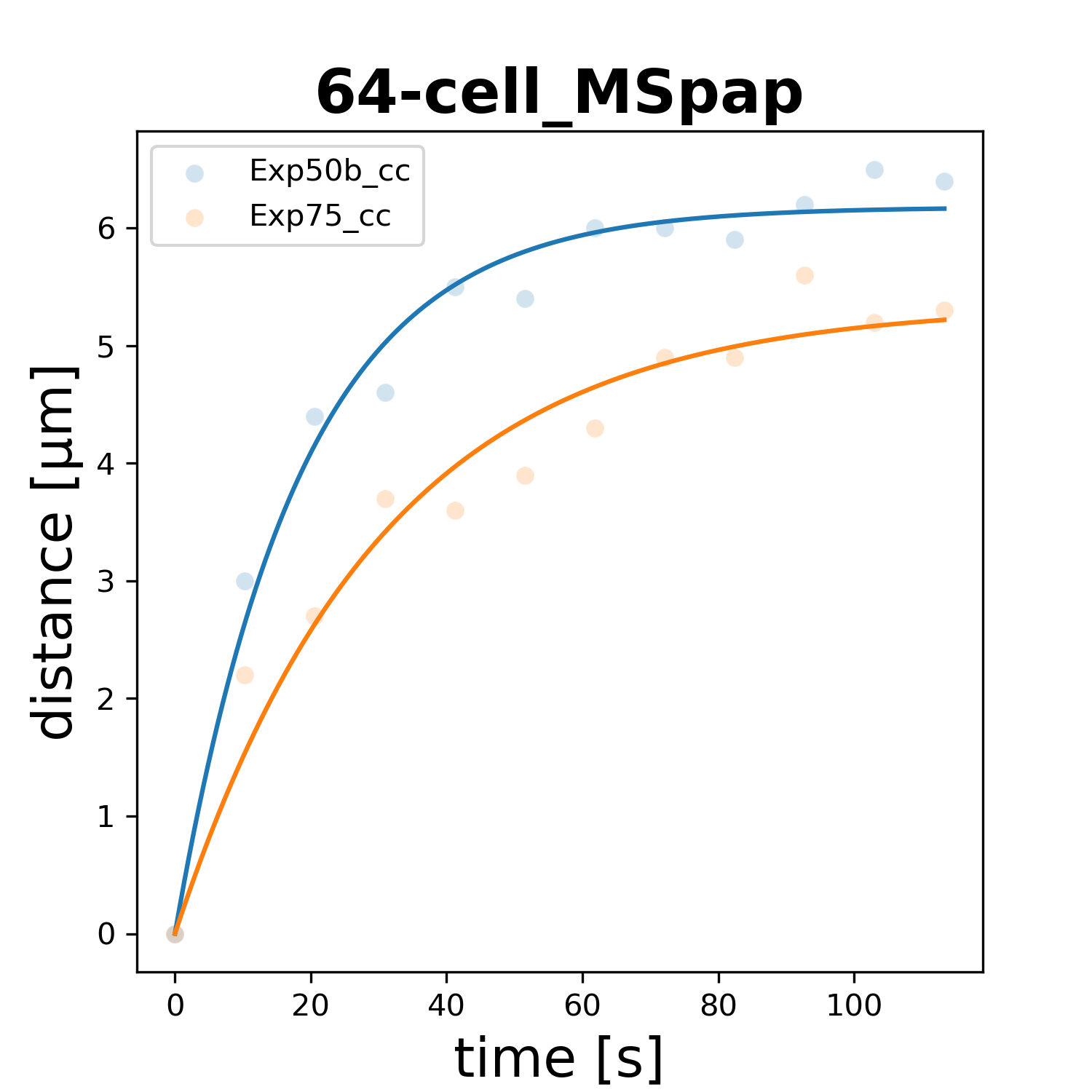
